# BioReason-Pro: Advancing Protein Function Prediction with Multimodal Biological Reasoning

**DOI:** 10.64898/2026.03.19.712954

**Authors:** Adibvafa Fallahpour, Arman Seyed-Ahmadi, Parsa Idehpour, Omar Ibrahim, Benedict M.H. Choi, Purav Gupta, Jack Naimer, Kevin Zhu, Abhinav Adduri, Arnav Shah, Shihao Ma, Talu Güloglu, Nuo Liu, Haotian Cui, Arihant Jain, Max de Castro, Amirfaham Fallahpour, Antonio Cembellin-Prieto, John S. Stiles, Filip Nemčko, Alexander A. Nevue, Hyungseok C. Moon, Lucas Sosnick, Olivia Markham, Haonan Duan, Michelle Y. Y. Lee, Andrea F. M. Salvador, Chris J. Maddison, Christoph A. Thaiss, Chiara Ricci-Tam, Brian S. Plosky, Dave P. Burke, Patrick D. Hsu, Hani Goodarzi, Bo Wang

## Abstract

Protein function annotation is fundamental to understanding biological mechanisms, designing therapeutics, and advancing biomedical research. Current computational methods either rely on shallow sequence similarity or treat function prediction as isolated classification tasks, failing to capture the integrative reasoning across sequence, structure, domains, and interactions that expert biologists perform to infer function. We introduce BioReason-Pro, the first multimodal reasoning large language model (LLM) for protein function prediction that integrates protein embeddings with biological context to generate structured reasoning traces. A key input into BioReason-Pro is the set of GO term predictions made by GO-GPT, our autoregressive transformer that captures hierarchical and cross-aspect dependencies of GO terms. BioReason-Pro is trained via supervised fine-tuning on synthetic reasoning traces generated by GPT-5 for over 130K proteins and further optimized through reinforcement learning. It achieves 73.6% *F*_max_ on GO term prediction and an LLM judge score of 8/10 on functional summaries, substantially outperforming previous methods. Evaluations with human protein experts show that BioReason-Pro annotations are preferred over ground truth UniProt annotations in 79% of cases. Remarkably, BioReason-Pro predicted a novel interaction partner for the renal cancer biomarker RCDG1, which we confirmed in the lab by co-immunoprecipitation. In other binding-partner predictions, its per-residue attention localized to the exact contact residues resolved in cryo-EM structures. Together, GO-GPT and BioReason-Pro establish a framework for protein function prediction that combines precise ontology modeling with interpretable biological reasoning.

## 1. Introduction

Proteins are the molecular machines of life, executing a vast repertoire of biochemical functions that underpin metabolism, signaling, and cellular organization (Fischer, 1894; Anfinsen, 1973). Advances in high-throughput sequencing have yielded an explosion of protein sequence data, with over 250 million entries in databases such as UniProt (Consortium et al., 2025). Yet functional characterization has not kept pace, as fewer than 0.1% of known proteins possess experimentally validated annotations due to the high cost and limited throughput of wet-lab assays (O’Donovan et al., 2002). This persistent annotation gap poses a major challenge for molecular biology and drug discovery, motivating the development of scalable computational approaches that infer protein function directly from sequence (Friedberg, 2006; Radivojac et al., 2013).

Protein function prediction is an integrative reasoning task. Biologists examining an uncharacterized protein analyze evidence from domain architecture, structural motifs, evolutionary context, organism biology, and interaction networks to construct a coherent functional hypothesis. Protein foundation models such as ESM3 (Hayes et al., 2024) and ProtT5 (Elnaggar et al., 2022) encode rich sequence-structure-function relationships from evolutionary data, yet this knowledge remains implicit, falling short of the deliberate reasoning that expert annotation demands. Conversely, advances in LLM reasoning have shown that explicit chain-of-thought dramatically improves performance on complex multi-step problems (Wei et al., 2023; OpenAI et al., 2024; Istrate and Karaletsos, 2025). To the best of our knowledge, no method yet combines the representational depth of protein foundation models with structured, multi-step reasoning to predict and explain protein function.

A complementary line of research focuses on free-text protein function prediction. These methods align protein representations with large language models (LLMs) to generate natural language functional summaries (Xiao et al., 2025b). Notable examples include Prot2Text (Abdine et al., 2023), ProtT3 (Liu et al., 2024), STELLA (Xiao et al., 2025a), ProtChatGPT (Wang et al., 2025b), ProteinCLIP (Wu et al., 2024), Evolla (Zhou et al., 2025), Prot2Chat (Wang et al., 2025d), and Prot2Text-V2 (Fei et al., 2025). Despite improved interpretability, these models remain descriptive rather than explanatory. They summarise what function a protein may have, but do not reason over relevant evidence or justify why that function is plausible.

Protein function is primarily described using the Gene Ontology (GO) (Ashburner et al., 2000), which organizes protein function into three aspects: Molecular Function (MF), Biological Process (BP), and Cellular Component (CC). GO terms are widely used to interpret large-scale experiments such as RNA-seq and proteomics (Consortium, 2021). Manual annotation requires expert curation and is time-consuming and costly. Early efforts transferred functions from homologous proteins identified by sequence alignment tools such as BLAST and DIAMOND (Altschul et al., 1990; Buchfink et al., 2015). Deep learning has since transformed this field, with methods such as NetGO 3.0 (Wang et al., 2023), InterLabelGO+ (Evans and Shen, 2024), DPFunc (Wang et al., 2025c), PhiGnet (Jang et al., 2024), ProtGO (Wang et al., 2025a), ProtNote (Char et al., 2025), and ProtBoost (Chervov, 2024) leveraging protein language model embeddings, structural representations (Fleming et al., 2025), and protein interaction networks, achieving strong performance on benchmarks such as the Critical Assessment of Functional Annotation (CAFA) (Radivojac et al., 2013). Despite this progress, these methods treat GO terms as isolated classification targets and their fixed ontology vocabulary constrains expressivity for proteins with novel or combinatorial functions (Kulmanov and Hoehndorf, 2020).

Here we present BioReason-Pro, the first multimodal reasoning LLM for protein function prediction. An important contribution to the reasoning of this model are predicted GO terms from GO-GPT, an autoregressive transformer for GO prediction that we built. GO-GPT treats GO annotation as a sequence generation task conditioned on protein representations, achieving a weighted 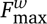 of 0.65–0.70 across inference strategies, surpassing the top accessible methods from the CAFA 5 competition (Friedberg et al., 2023b). By integrating output from GO-GPT, protein embeddings, and additional biological context, BioReason-Pro is able to generate structured traces that reason from domain analysis to functional hypotheses. We trained BioReason-Pro on over 130K synthetic reasoning traces spanning 3,135 organisms and refined it with reinforcement learning. Human experts preferred BioReason-Pro annotations over curated UniProt entries in 79% of evaluated cases, and an LLM judge scored its functional summaries 8/10 on average, substantially outperforming previous methods even for proteins with very low similarity to training data. Remarkably, BioReason-Pro de novo identified experimentally validated binding partners for individual test proteins, with per-residue attention localizing to the precise contact interfaces resolved in cryo-EM structures of those complexes. The model also performed structural reasoning that overrode misleading superfamily-level domain annotations, a capability beyond the reach of homology transfer or domain lookup alone.

To enable broad adoption, we release all model weights, code, and curated datasets, alongside a web interface and model predictions for over 240,000 proteins including the Human Protein Atlas. BioReason-Pro demonstrates that AI systems can reason about protein function at expert level, opening a path toward scalable functional characterization of the millions of uncharacterized proteins across all domains of life.

## 2. Results

### 2.1. Building a biology-language multimodal model for protein function prediction

Protein function is determined by the interplay of sequence, structure, domain architecture, evolutionary context, and molecular interaction networks, which are the same evidence sources that expert biologists integrate when annotating uncharacterized proteins. To test whether a language model augmented with these biological modalities could reason about protein function, we built BioReason-Pro (**Fig. 1A**), a multimodal LLM based on Qwen3-4B (Yang et al., 2025). BioReason-Pro reasons over biological context in natural language, leveraging the chain-of-thought capabilities that have emerged in recent LLMs (OpenAI et al., 2024; Guo et al., 2025). BioReason-Pro integrates residue-level embeddings from ESM3 (Hayes et al., 2024), which encodes protein function jointly with sequence and structure to provide a biologically grounded representation of protein context. Separately, a GO graph encoder captures the structure of the Gene Ontology as embeddings. Alongside these embeddings, the LLM receives target organism, domain annotations, protein-protein interactions, and initial GO term hypotheses from GO-GPT.

**Figure 1.**
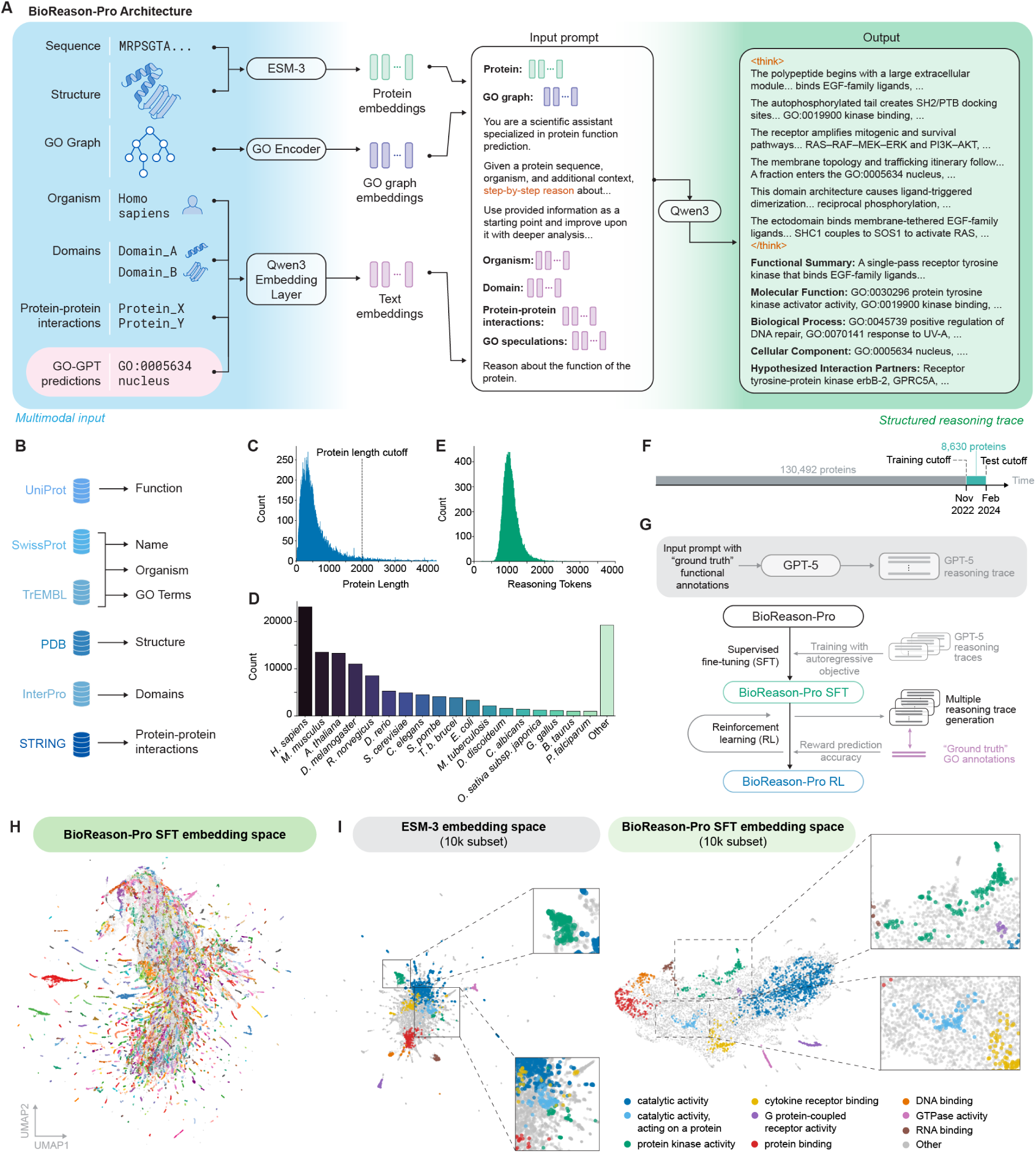
Overview of BioReason-Pro for protein function prediction. **(A) BioReason-Pro architecture.** A multimodal reasoning LLM that integrates ESM3 protein embeddings, a GO graph encoder, and biological context to generate structured reasoning traces and functional annotations. **(B) Dataset overview.** 133,492 proteins across 3,135 organisms curated from UniProt with experimental GO annotations, InterPro domains, STRING protein-protein interactions, and PDB protein structures. **(C) Protein length distribution.** Distribution of protein sequence lengths across the dataset with a 2,000 residue cutoff. **(D) Organism breakdown.** Taxonomic diversity of the training dataset. **(E) Reasoning trace distribution.** Token length distribution of synthetic reasoning traces across training proteins. **(F) Temporal split.** Training data through November 2022 and test data from March 2023 to February 2024. **(G) Training procedure.** SFT on GPT-5-generated reasoning traces followed by RL with GO term prediction accuracy as the reward. **(H) Learned embedding space.** UMAP projection of BioReason-Pro SFT LLM Layer 35 embeddings across all training proteins, colored by HDBSCAN clusters computed in PCA-reduced space. **(I) Embedding space comparison.** UMAP projections of ESM3 (left) and BioReason-Pro SFT (right) embeddings for a 10K protein subset. Cluster assignments were derived from BioReason-Pro embeddings and applied to both panels, with each cluster annotated by its most enriched GO Molecular Function term.

To ground our models in known biology, we collected domain annotations from InterPro (Blum et al., 2024), protein-protein interactions from STRING (Szklarczyk et al., 2024), protein structures from the PDB (Burley et al., 2018), and subcellular localization metadata (**Fig. 1B**). For training, we curated a broadly representative protein function dataset from UniProt (Consortium et al., 2025) comprising 133,492 proteins across 3,135 organisms (**Fig. S1A,B**). Following the CAFA community challenge (Friedberg et al., 2023b), which defines temporal holdout protocols and experimental evidence codes for benchmarking function prediction, we retained only proteins with experimental GO annotations (**Fig. S1C–F**). We excluded incomplete sequences and truncated proteins to 2000 amino acids to fit within the ESM3 context window (**Fig. 1C**). The resulting dataset spans broad taxonomic diversity (**Fig. 1D**), with per-protein domain, interaction, and localization statistics in **Fig. S1G–I**.

Training a model to reason about protein function requires step-by-step reasoning traces, yet no large-scale corpus of human-authored traces exists. Recent work has shown that models can learn to reason through supervised fine-tuning (SFT) on LLM-generated synthetic reasoning traces and be further improved via reinforcement learning (RL) to increase prediction accuracy (Guo et al., 2025; Zelikman et al., 2022; Fallahpour et al., 2025b). We adopted this approach by using GPT-5 (Singh et al., 2025) to generate over 130K synthetic traces for protein function prediction (**Fig. 1E**, Section C.1). We split the data temporally according to the CAFA framework (Zhou et al., 2019), where proteins annotated before November 2022 were used for training, and those that gained new experimental annotations between March 2023 and February 2024 but lacked prior annotations in the target GO aspect were held out for evaluation (**Fig. 1F**). BioReason-Pro was trained via SFT with next-token prediction on these traces (**Fig. 1G**), which taught BioReason-Pro to generate biological reasoning traces. To directly optimize for GO term annotation accuracy, we further applied reinforcement learning via Group Sequence Policy Optimization (GSPO) (Zheng et al., 2025), using 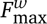 between predicted and ground truth GO terms as the reward signal. The protein embedding space learned by BioReason-Pro SFT is visualized across the full training set (**Fig. 1H**). We also compare ESM3 and BioReason-Pro SFT embeddings on a 10K protein subset, with clusters annotated by their most enriched GO Molecular Function term (**Fig. 1I**).

### 2.2. GO-GPT improves GO term prediction and captures biologically meaningful structure

As shown in **Fig. 1A**, BioReason-Pro operates on a set of likely GO terms for the given protein. For this, we use our GO-GPT model, which is an autoregressive transformer (Vaswani et al., 2023; Radford et al., 2019) that generates GO terms sequentially, with each prediction conditioned on residue-level protein embeddings from ESM2 (Lin et al., 2023), the user-specified organism, and all previously generated terms (**Fig. 2A**). We chose ESM2 over ESM3 as it performed comparably on GO term prediction while enabling more efficient batched embedding extraction (Appendix B.3). This approach captures both the hierarchical relationships within ontology aspects and the dependencies across them that independent classification approaches miss.

**Figure 2.**
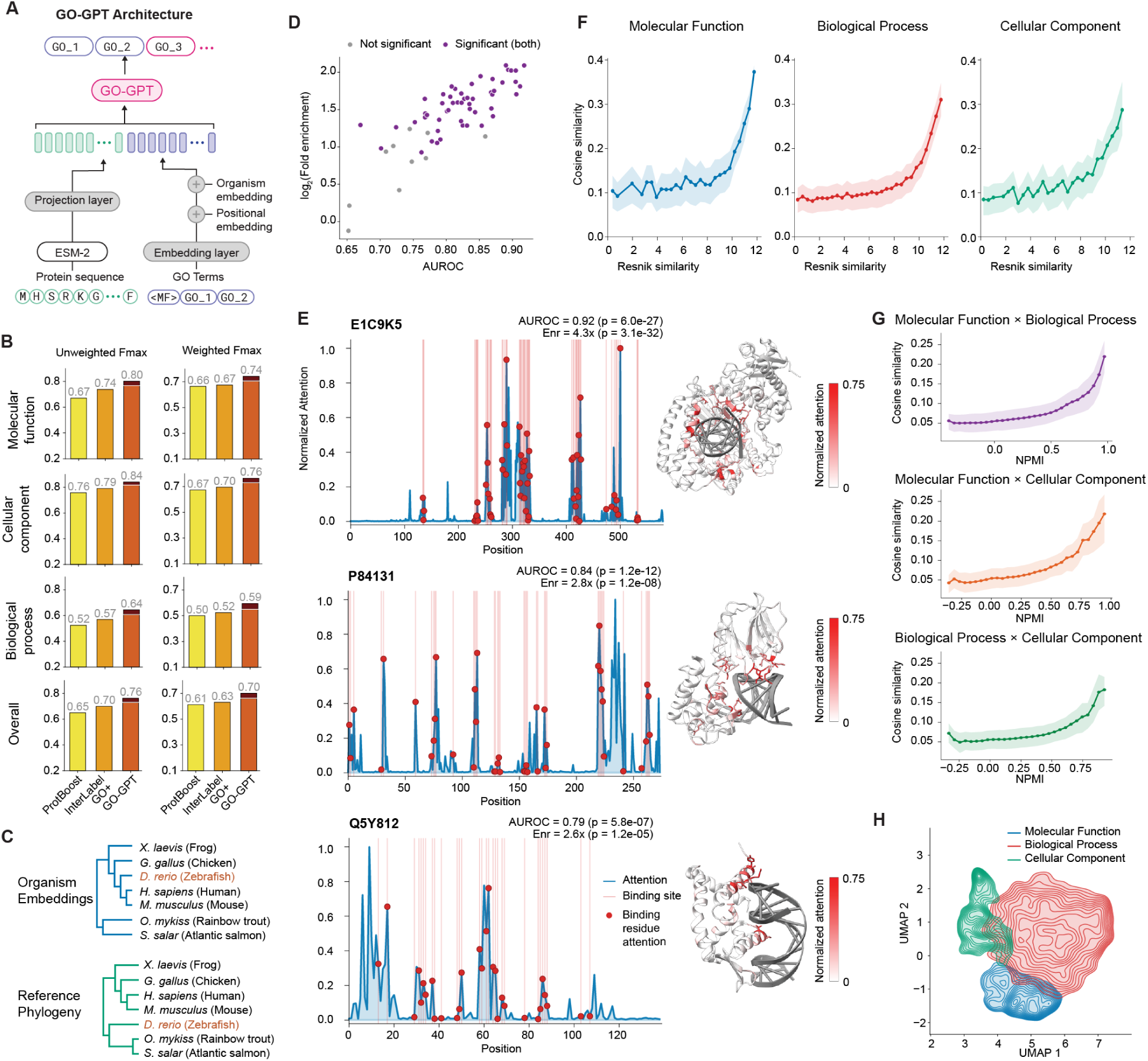
GO-GPT architecture, performance, and interpretability analysis. **(A) GO-GPT architecture.** Autoregressive Transformer encoding protein sequences via ESM2, generating GO terms conditioned on organism and previously generated terms. **(B) GO term prediction performance.** Weighted and unweighted *F*_max_ across all aspects (dark shading: best-of-10 upper bound; all pairwise differences significant except GO-GPT greedy vs. InterLabelGO+ on weighted MF, *p* = 0.073; Tables S3–S5). **(C) Organism embedding phylogeny.** Dendrogram from cosine similarity of learned organism embeddings (top) compared against a reference phylogeny (bottom). The learned tree recapitulates known phylogenetic relationships, with *D. rerio* (highlighted in red) as the only misplaced organism. **(D) Binding residue attention enrichment.** AUROC vs. log_2_(fold enrichment) for 63 non-training BioLiP proteins where GO-GPT predicted GO:0003677 (DNA binding). Purple: significant on both Mann–Whitney U and hypergeometric tests (*p* < 0.05); gray: not significant on at least one. **(E) DNA-binding attention analysis.** Per-residue attention when predicting GO:0003677 for three proteins (E1C9K5, P84131, Q5Y812) spanning high to low binding-site overlap. Left: attention profiles with binding sites shaded; right: protein–DNA structures with high-attention residues in red. **(F) Resnik vs. cosine similarity.** Cosine embedding similarity increases monotonically with Resnik semantic similarity (Resnik, 1995) across all aspects (all *p* < 10^−3^, Mantel permutation test). **(G) Cross-aspect NPMI vs. cosine similarity.** Co-annotation NPMI correlates with embedding similarity across aspect pairs (Spearman *ρ* = 0.10–0.17, *p* < 10^−3^, permutation test). **(H) GO term embedding landscape.** UMAP with KDE contours per aspect (MF blue, BP red, CC green), showing aspect-level spatial organization.

The Gene Ontology organizes protein function as a directed acyclic graph from general root concepts to highly specific leaf annotations (Ashburner et al., 2000; Aleksander et al., 2023). Existing methods predict each GO term independently, ignoring both this hierarchical structure and the dependencies between terms. GO-GPT captures these relationships by treating GO prediction as a sequential generation task, predicting increasingly specific terms conditioned on residue-level protein embeddings, the target GO aspect, more general GO terms, and organism. We evaluated GO-GPT against InterLabelGO+ (Evans and Shen, 2024) and ProtBoost (Chervov, 2024), the highest-ranked CAFA 5 methods with publicly available implementations, on a temporal holdout set following the CAFA framework (Friedberg et al., 2023a) (**Fig. 1F**; **Fig. 2B**).

We measure performance of function prediction models using *F*_max_, the maximum F-measure (harmonic mean of precision and recall) across all possible probability thresholds (Section 4.4.1) (Clark and Radivojac, 2013b). For each threshold, GO terms with probability above the threshold are considered predictions; precision and recall are computed against ground truth and averaged across proteins. *F*_max_ selects the threshold maximizing this protein-averaged F1, providing a single summary of predictive performance independent of threshold choice. We report both unweighted *F*_max_, where all terms contribute equally, and weighted 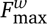, which emphasizes rare and specific terms that are harder to predict but more biologically informative (Zhou et al., 2019). *F*_max_ requires per-term probability estimates, but GO-GPT generates sequences of discrete GO tokens. Assigning probability 1 to all generated terms and 0 otherwise, GO-GPT achieves 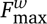 = 0.65, surpass-ing InterLabelGO+ at 0.63 (significant for BP and CC; MF difference not significant under greedy decoding, *p* = 0.073; Table S3). Since GO-GPT generates sequences stochastically, sampling multiple trajectories per protein allows estimating per-term probabilities. Generating 10 independent samples per protein and using term frequencies as probabilities improves performance to 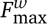 = 0.67. Selecting the best of 10 samples per protein reaches 0.70, indicating substantial headroom from improved decoding or selection strategies. GO-GPT shows a similar gap on unweighted *F*_max_, achieving 0.76 compared to 0.70 for InterLabelGO+ (**Fig. 2B**). These results demonstrate that autoregressive generation is a powerful framework for GO term prediction, capturing both hierarchical relationships within aspects and dependencies across them. Sampling-based confidence estimation emerges naturally from this generative approach, providing per-term probability estimates without requiring additional calibration.

Beyond predictive accuracy, we examined whether GO-GPT’s learned representations capture biologically meaningful structure. GO-GPT learns a unique embedding for each organism during training. Because evolutionary relatedness shapes the functional constraints acting on proteins, we asked whether these learned embeddings recover known phylogenetic structure. Computing cosine similarity between these embeddings and constructing a dendrogram produces a tree that recapitulates known phylogenetic relationships, with closely related taxa consistently grouping together (**Fig. 2C**; Mantel test, *p* = 4 ×10^−3^, for top 30 most frequent organisms) (Fallahpour et al., 2025a; Kumar et al., 2022; Mantel, 1967). This suggests that the model captures organism-specific biases in how protein sequence maps to function, and that these biases are more conserved between phylogenetically proximal species.

We next asked whether GO-GPT leverages functionally relevant regions of the protein when making predictions. To systematically evaluate this, we focused on DNA-binding proteins, whose direct interaction with DNA through specific residues provides a well-defined ground truth for assessing whether attention aligns with functional sites. We analyzed attention patterns when GO-GPT predicts the DNA-binding function term (GO:0003677), using proteins with annotated DNA-binding residues from BioLip (Yang et al., 2012) that were absent from the training set (Section 4.5.1). For each protein, we extracted per-residue attention scores when the model generates the DNA-binding term (GO:0003677), producing an attention profile across the sequence that can be compared against known binding sites. We then quantified the correspondence between model attention and known binding sites using two complementary metrics: AUROC, which measures how well attention scores distinguish binding from non-binding residues, and fold-enrichment, which measures the concentration of attention at binding sites relative to the sequence background.

Across all 63 evaluated proteins, attention is consistently enriched at annotated binding residues (mean AUROC = 0.81 ± 0.06, significant in 59/63; mean top-20% fold-enrichment = 2.8x ± 0.7x, significant in 55/63), with AUROC and fold enrichment positively correlated across proteins (**Fig. 2D**) and 60 of 63 exceeding AUROC 0.7 (full per-protein statistics in Table S1; **Fig. S2**). Three proteins spanning different AUROC values (E1C9K5, P84131, Q5Y812; selected as described in Section 4.5.1) are visualized in **Fig. 2E**, with AUROC ranging from 0.79 to 0.92 and top-20% fold-enrichment from 2.6x to 4.3x (all *p* < 10^−3^, Mann–Whitney U and hypergeometric tests). Projecting high-attention residues onto protein–DNA complex structures confirms that attended regions cluster proximal to bound DNA. To rule out that this enrichment merely reflects pre-existing structure in the frozen ESM2 embeddings, we compared against ESM2 residue embedding L2 norms as a baseline predictor of binding sites (Section 4.5.2; **Fig. S2C–D**). ESM2 norms show substantially weaker discrimination, suggesting that cross-attention captures function-specific signal not readily apparent in the input representations.

GO-GPT’s vocabulary consists of individual GO terms as discrete tokens, each with its own learned embedding. To test whether these embeddings capture the semantic organization of the Gene Ontology, we measured the relationship between cosine embedding similarity and Resnik semantic similarity (Resnik, 1995), an information-content measure defined over the GO directed acyclic graph, within each aspect (**Fig. 2F**). Cosine similarity increases monotonically with Resnik similarity across all three aspects, with a pronounced rise at high Resnik values, indicating that the embeddings particularly distinguish the most semantically related term pairs (all *p* < 10^−3^, Mantel permutation test; Section 4.5.3). Consistently, *κ*-nearest neighbors in embedding space overlap significantly with ontological neighbors in the GO graph across all aspects (**Fig. S6B**; *p* < 10^−3^, permutation test).

We next asked whether the embeddings also capture relationships that span GO aspects. Normalized Pointwise Mutual Information (NPMI) between cross-aspect GO term pairs (Section 4.5.3) correlates positively with cosine embedding similarity across all three aspect pairs (Spearman *ρ* = 0.10–0.17, all *p* < 10^−3^, embedding-permutation test; **Fig. 2G**), with an upturn at high NPMI values most visible for MF×CC. To test whether this correlation translates into practical retrieval, we ranked partner-aspect terms by cosine similarity for each GO term and evaluated retrieval of strongly co-annotated partners (NPMI > 0.5; Section 4.5.3). Embedding similarity retrieves correct cross-aspect partners well above chance across all three aspect pairs (AUROC = 0.66–0.80, all *p* < 10^−3^; Precision@1 = 3.5–6×above random baseline), with MF×CC showing the strongest retrieval (AUROC = 0.80; **Fig. S7B**). Notably, the model receives only numerical GO identifiers and never human-readable term names, so these cross-aspect associations are learned entirely from protein co-annotation patterns. UMAP projection (McInnes et al., 2020) of the embedding space (**Fig. 2H**) visually reflects both types of organization: MF and CC embeddings form more compact, localized clusters while BP is diffusely spread. This is consistent with quantitative cosine purity analysis showing MF and CC neighborhoods enriched 3–5× above their aspect-specific random baselines (*p* < 10^−3^) whereas BP shows no enrichment (**Fig. S7C**). Regions of inter-aspect overlap are also visible; representative cross-aspect pairs with high co-annotation frequency and embedding similarity that happen to co-localize in the projection are annotated in **Fig. S8** (Table S2).

Together, these analyses demonstrate that GO-GPT captures functional, evolutionary, and ontological structure within its learned representations. The model grounds its GO term predictions in biologically relevant sequence features and has internalized both phylogenetic relationships and ontological structure without explicit supervision on either.

### 2.3. BioReason-Pro demonstrates generalizable expert-level reasoning on protein function

While GO-GPT generates GO term predictions through internal mechanisms that are not directly accessible and require post hoc analysis to interpret, BioReason-Pro is designed to openly reason across multimodal biological evidence to produce interpretable functional annotations. It analyzes domain architecture, infers molecular function, predicts localization, identifies relevant biological processes, proposes mechanistic explanations, and hypothesizes interaction partners. Standard text generation metrics such as ROUGE and BERTScore are therefore inadequate for evaluating such reasoning, as they measure surface-level overlap rather than biological correctness (Ganesan, 2018; Zhang et al., 2020). We employed GPT-5.1, the most capable model available at the time of evaluation, as an expert judge, scoring predictions from BioReason-Pro SFT, BioReason-Pro RL, and the best prior method Prot2Text-v2 against composite ground truth including UniProt function summaries and GO terms across all three aspects (Fei et al., 2025).

The LLM judge scored each prediction on a 1 to 10 scale across five axes, following the prompt detailed in Section C.3. Molecular Function, Biological Process, and Cellular Component assessed correctness of core functional annotations, Specificity evaluated the level of mechanistic detail provided, and Reliability distinguished logically supported inference from hallucinated claims (**Fig. 3A**). We evaluated BioReason-Pro SFT, trained on synthetic reasoning traces, and BioReason-Pro RL, further trained with reinforcement learning to optimize GO term accuracy (Section 2.4). BioReason-Pro RL achieved an average score of 8.03 out of 10, with SFT at 7.65, both substantially outperforming Prot2Text-v2 at 4.15 (**Fig. 3B**). Across individual axes, Molecular Function yielded the highest scores (RL 8.50, SFT 8.11) while Biological Process scored the lowest (RL 7.51, SFT 6.94). Reinforcement learning improved Reliability from 6.92 to 7.56, indicating that optimizing for annotation accuracy also reduced hallucinations, while SFT scored marginally higher on Specificity (8.74 vs 8.59). All pairwise differences between BioReason-Pro and baselines are statistically significant, as are differences between RL and SFT on all axes (paired Wilcoxon signed-rank tests, *p* < 10^−15^; full distributions in **Fig. S3A**, detailed statistics in Section B.5.2, Table S10).

**Figure 3.**
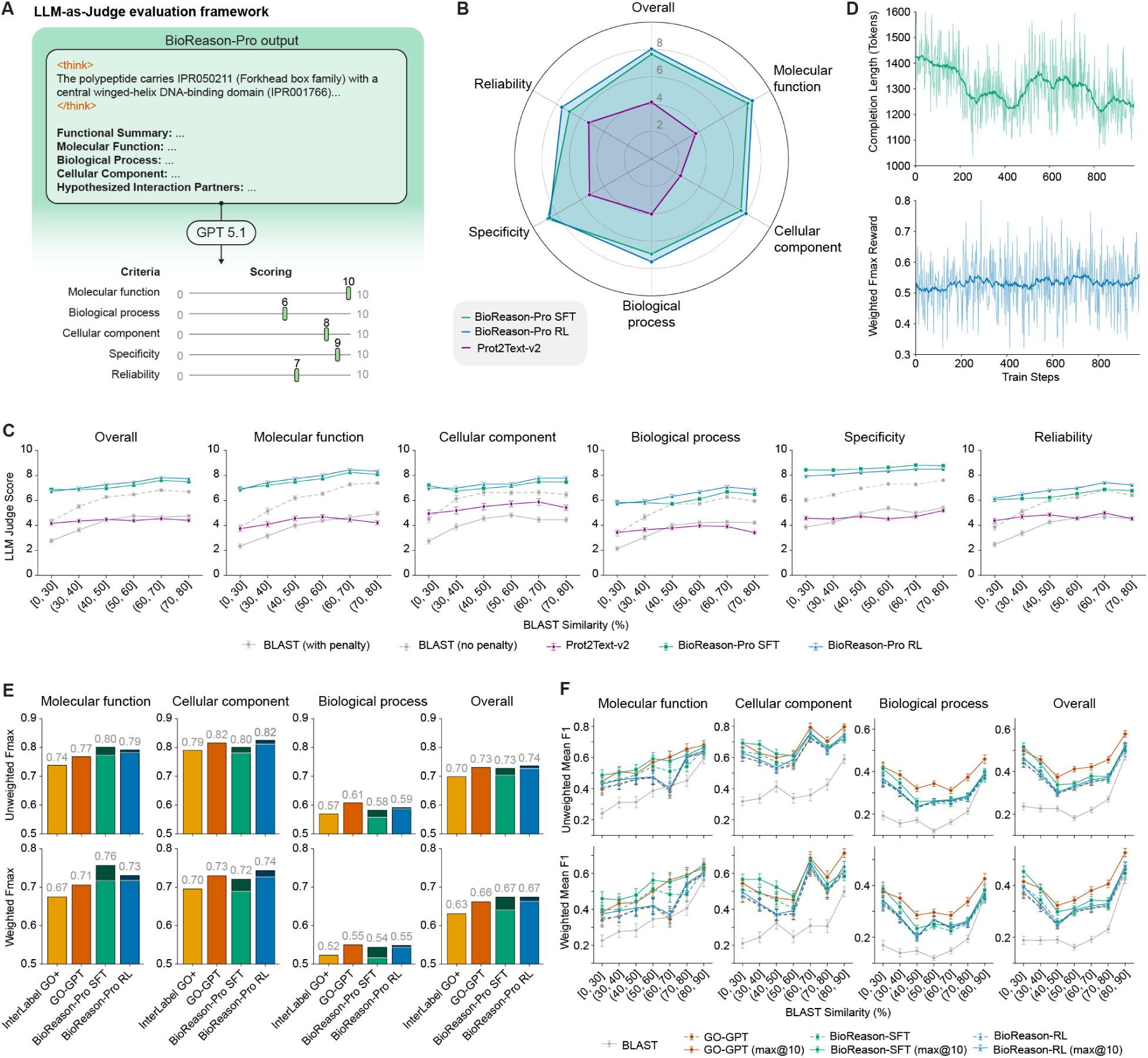
BioReason-Pro evaluation on protein function prediction. **(A) LLM-as-Judge framework.** GPT-5.1 evaluated model predictions against composite ground truth comprising UniProt summaries, GO terms, InterPro domains, protein-protein interactions, and subcellular localization. Each prediction was scored on a 1 to 10 scale across five axes: Molecular Function, Biological Process, Cellular Component, Specificity, and Reliability. **(B) LLM-as-Judge scores.** Radar plot comparing per-axis scores for BioReason-Pro RL, BioReason-Pro SFT, and Prot2Text-v2. BioReason-Pro RL achieved an average score of 8.03 across axes, SFT 7.65, and Prot2Text-v2 4.15. **(C) LLM judge generalization across sequence similarity.** LLM judge scores across BLAST similarity bins for BioReason-Pro SFT, BioReason-Pro RL, Prot2Text-v2, and BLAST baselines (with and without penalty), shown for all five evaluation axes and Overall. **(D) Reinforcement learning dynamics.** Completion length in tokens (top) and 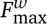 reward (bottom) over GSPO training steps. Reward increased while generation length decreased, indicating more focused reasoning. **(E) GO term prediction performance.** Weighted and unweighted *F*_max_ across Molecular Function, Cellular Component, Biological Process, and Overall for InterLabelGO+, GO-GPT, BioReason-Pro SFT, and BioReason-Pro RL (dark shading indicates best-of-10 selection upper bound). **(F) GO term prediction generalization across sequence similarity.** Unweighted (top) and weighted (bottom) mean per-protein F1 scores averaged within BLAST similarity bins for GO-GPT, BioReason-Pro SFT, BioReason-Pro RL, and BLAST, with best-of-10 selection variants shown for each generative model.

We next asked whether reasoning quality holds for proteins dissimilar from training data. We binned test proteins by best-hit BLAST sequence similarity to the training set and compared LLM judge scores across bins (**Fig. 3C**) (Altschul et al., 1990). As our BLAST baseline, we transferred the functional annotations of each test protein’s closest training set hit, scoring proteins whose closest hit lacked a UniProt function description as zero (BLAST with penalty) or excluding them entirely (BLAST without penalty). BLAST performance showed strong dependence on sequence similarity (Spearman *ρ* = 0.43), dropping sharply at low similarity. BioReason-Pro exhibited roughly half this dependence (RL *ρ* = 0.27, SFT *ρ* = 0.21), maintaining scores of 7–8.5 across the full similarity range, with OLS regression confirming that its advantage over BLAST grows significantly with decreasing similarity (*β*_1_ = − 0.019, *p* = 3.1 × 10^−30^; Section B.5.3, Table S11). Even at high similarity bins where close homologs including orthologs were readily available, BioReason-Pro outperformed BLAST.

Beyond sequence similarity, BioReason-Pro performance remained stable across protein lengths and GO annotation counts (**Fig. S3B,C**). Scores decreased with fewer InterPro domain annotations, but BioReason-Pro still outperformed all baselines even for proteins with zero or few annotated domains (**Fig. S3D**). Performance also generalized across diverse organisms and taxonomic classes rather than concentrating around well-studied model species (**Fig. S4**). Together, these results demonstrate that BioReason-Pro learned generalizable functional reasoning rather than retrieving annotations from similar training proteins.

### 2.4. Reinforcement learning produces more accurate and concise GO term predictions

While supervised fine-tuning taught BioReason-Pro to generate biological reasoning traces, reinforcement learning optimized those traces for GO term prediction accuracy. Over training, the 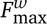 reward between predicted and ground truth GO terms increased while reasoning traces became shorter and more focused, suggesting the model learned to eliminate verbose reasoning that did not contribute to correct predictions (**Fig. 3D**). Per-protein comparison confirmed that RL produced significantly shorter traces than SFT for the same proteins (mean Δ = 60.0 words, paired Wilcoxon *p* < 10^−300^; **Fig. S3E,F**). Notably, RL increasingly outperformed SFT at shorter reasoning trace lengths, indicating that reinforcement learning produced more efficient reasoning rather than simply shorter output (**Fig. S3G**).

We evaluated GO term prediction using 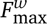, comparing BioReason-Pro against GO-GPT and InterLabelGO+ (**Fig. 3E**) (Evans and Shen, 2024). BioReason-Pro SFT achieved 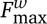 = 0.64 on single-run greedy decoding and 0.67 with best-of-10 sampling, revealing substantial generation diversity. Reinforcement learning closed this gap, with BioReason-Pro RL achieving 0.66 on single-run greedy decoding while best-of-10 sampling remained at 0.67, indicating that RL concentrated probability mass on correct predictions. The best unweighted *F*_max_ sits at 0.74. Both BioReason-Pro variants surpassed InterLabelGO+ at 0.63 and single-run GO-GPT at 0.65 on 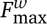, with consistent improvements across unweighted metrics.

We then examined how GO term prediction accuracy varied with sequence similarity by binning test proteins according to best-hit BLAST sequence similarity (**Fig. 3F**) (Altschul et al., 1990). Because *F*_max_ cannot be decomposed across subsets of proteins, we report mean per-protein F1 scores within each bin to enable direct comparison across similarity ranges. BLAST showed the strongest similarity dependence (Spearman *ρ* = 0.63), while BioReason-Pro and GO-GPT exhibited substantially weaker dependence (SFT *ρ* = 0.41, RL *ρ* = 0.46, GO-GPT *ρ* = 0.46), with OLS regression confirming that all generative models maintained significantly greater advantage over BLAST at low similarity (all *p* < 10^−130^; Section B.5.3, Table S12). Even at high similarity where close homologs were available, BioReason-Pro outperformed BLAST, demonstrating that biological reasoning captured functional relationships beyond sequence alignment alone.

### 2.5. Human experts prefer BioReason-Pro over ground truth annotations

Although the results from the automated metrics and LLM judges are promising, it is difficult to extrapolate from these to whether the annotations are genuinely useful to domain experts. We therefore conducted a blinded human evaluation in which 27 molecular biologists assessed BioReason-Pro predictions on 162 randomly selected test proteins (Section C.4). Evaluators were blinded to model identity (SFT or RL) and had access to external sources and recent literature. Evaluators performed pairwise preference comparisons between each model and ground truth UniProt annotations on a five-point scale from significantly falls short to significantly exceeds. BioReason-Pro SFT achieved a 79% tie-or-exceed rate against UniProt ground truth, with RL at 73%, meaning human experts found model predictions to match or exceed curated database entries in the majority of cases (**Fig. 4A**). This shows that RL improved GO term accuracy (Section 2.4) without significantly reducing the reasoning quality (McNemar’s exact test, *p* = 0.087).

**Figure 4.**
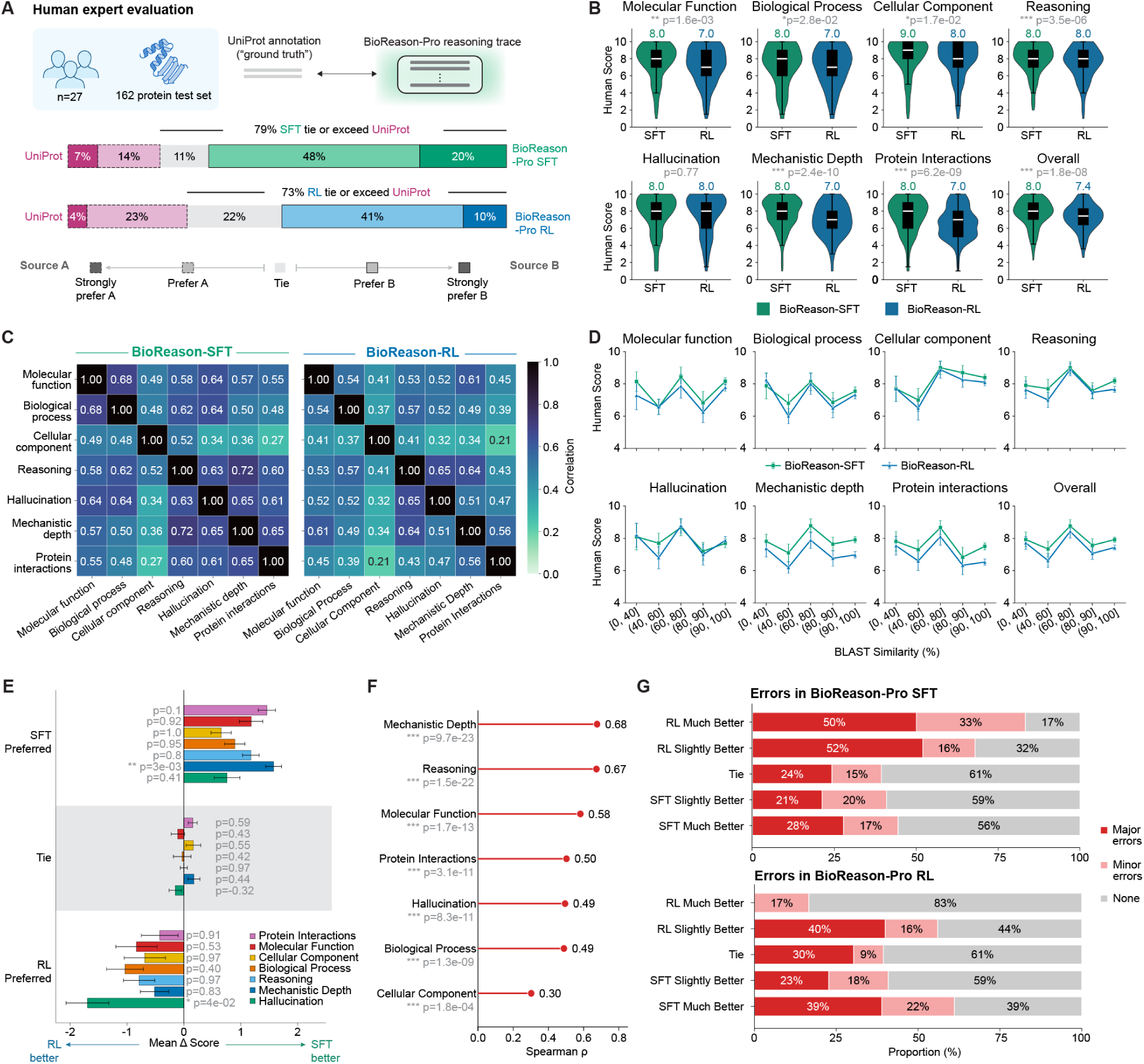
Human expert evaluation of BioReason-Pro predictions. **(A) Pairwise preference comparisons.** 27 protein experts compared BioReason-Pro SFT and RL predictions against UniProt ground truth on a five-point preference scale. SFT tied or exceeded UniProt in 79% of cases, RL in 73%. **(B) Per-axis score distributions.** Violin plots of expert scores for BioReason-Pro SFT and RL across Molecular Function, Biological Process, Cellular Component, Reasoning, Hallucination, Mechanistic Depth, Protein Interactions, and Overall, with median scores indicated. **(C) Correlation structure of evaluation axes.** Pairwise correlations between the seven evaluation axes for SFT and RL, showing that Reasoning, Mechanistic Depth, and Hallucination form a correlated cluster reflecting overall biological reasoning quality. **(D) Human evaluation across sequence similarity.** Expert scores across BLAST sequence similarity bins for BioReason-Pro SFT and RL across all seven evaluation axes and Overall, showing stable performance with no degradation at low similarity. **(E) Score deltas by preference group.** Mean per-axis score difference (SFT RL) stratified by expert preference. SFT advantage is driven by Mechanistic Depth (*p* = 3 × 10^−3^), while RL advantage is driven by Hallucination (*p* = 4 × 10^−2^). **(F) Axes driving expert preference.** Spearman correlation between per-axis score deltas and expert preference, identifying Mechanistic Depth and Reasoning as the primary axes driving preference decisions. **(G) Error attribution by expert preference.** Proportion of major, minor, or no errors (classified by GPT-5-mini from free-text responses) in each model, stratified by expert head-to-head preference (Section C.5).

Evaluators also scored each prediction on a 1 to 10 scale along seven axes. Molecular function, biological process, and cellular component assessed correctness of core functional annotations; reasoning evaluated evidence attribution; hallucination measured avoidance of fabricated biological claims; mechanistic depth measured accuracy of proposed molecular mechanisms; and protein-protein interactions evaluated plausibility of predicted binding partners. BioReason-Pro SFT averaged 8.0 out of 10 overall with RL at 7.4 (**Fig. 4B**; full distributions in **Fig. S5A**). Reasoning, Mechanistic Depth, and Hallucination scores were strongly correlated with each other across both models, indicating that predictions with well-supported reasoning also tended to avoid hallucinations and provide deeper mechanistic insight (**Fig. 4C**; **Fig. S5B**).

We examined whether human-perceived annotation quality depends on sequence similarity to training data by comparing scores across BLAST sequence identity bins (**Fig. 4D**) (Altschul et al., 1990). Neither model showed significant similarity dependence on any evaluation axis (Spearman *ρ* < 0.06 for overall scores, all *p* > 0.05; Section B.5.4, Table S13), and the score gap between RL and SFT was also stable across the similarity range (all *p* > 0.05; Table S14). Win rates against UniProt did not depend on similarity for either model (logistic regression, *p* = 0.71 for SFT, *p* = 0.19 for RL; Table S15; **Fig. S5H**).

To understand what drives expert preference between SFT and RL, we examined per-axis score differences stratified by preference group. When experts preferred SFT, the score advantage was concentrated in Mechanistic Depth (*p* = 3×10^−3^), while when experts preferred RL, the advantage was driven by fewer hallucinated claims (*p* = 4 × 10^−2^; **Fig. 4E**). Mechanistic Depth and Reasoning showed the highest overall correlation with expert preference decisions (**Fig. 4F**). Error attribution of free-text expert responses confirmed this pattern, with SFT-preferred cases showing richer mechanistic detail and RL-preferred cases containing fewer major errors (**Fig. 4G**). Where experts preferred RL, free-text error reports revealed that SFT made specific mechanistic errors such as enzyme misidentification and inverted pathway directionality, whereas RL’s more conservative outputs avoided these hallucinations. This suggests that RL trades some mechanistic depth for improved factual reliability.

Beyond sequence similarity, performance generalized across diverse organisms (**Fig. S5C**) and remained stable across protein lengths, GO annotation counts, InterPro domain counts, and reasoning trace lengths (**Fig. S5D,E,F,G**). Together, these results demonstrate that BioReason-Pro produces expert-level functional reasoning across a broad range of proteins regardless of similarity to training data.

### 2.6. BioReason-Pro generates de novo predictions validated by experimental structures

We asked whether BioReason-Pro could synthesize specific mechanistic predictions from its input context alone and whether such predictions are grounded in structurally meaningful features. We screened test set generations for cases in which the model proposed specific interaction partners or pathway-level mechanisms that could be validated against experimental structural data. One such case is eEFSec (P57772), the selenocysteine-specific elongation factor, a four-domain translational GTPase that delivers the 21st amino acid, selenocysteine, to ribosomes at recoded UGA stop codons. Unlike canonical elongation factors such as eEF1A that service all standard aminoacyl-tRNAs through a shared mechanism, eEFSec operates through a non-canonical pathway requiring coordination with a dedicated mRNA stem-loop element and an accessory protein that no other translation factor uses (Copeland et al., 2000; Hilal et al., 2022). The closest training set match shared 44% sequence identity, a regime where homology transfer may capture general family membership but cannot resolve the mechanistic specializations that distinguish eEFSec from other EF-Tu-like GTPases.

BioReason-Pro processed domains spanning the N-terminal GTPase core through the C-terminal RIFT module. The reasoning traced how the translational-type GTP-binding domain and EF-Tu-like beta-barrel establish canonical translational GTPase machinery, then identified the C-terminal selenocysteine-specific domains (IPR049393, IPR049394) as restricting this factor to a single substrate (Gonzalez-Flores et al., 2012). Rather than assigning generic tRNA binding, the model described selenocysteine-specific tRNA recognition as the defining molecular function (full traces in **Fig. 5A** and Section C.6), a functional distinction that separates eEFSec from all other EF-Tu family members but that the Gene Ontology has not formalized into a dedicated term. Biochemical studies support this specificity, showing that eEFSec binds only selenocysteinyl-tRNA(Sec) and rejects both the serylated precursor and canonical aminoacyl-tRNAs (Fagegaltier, 2000).

**Figure 5.**
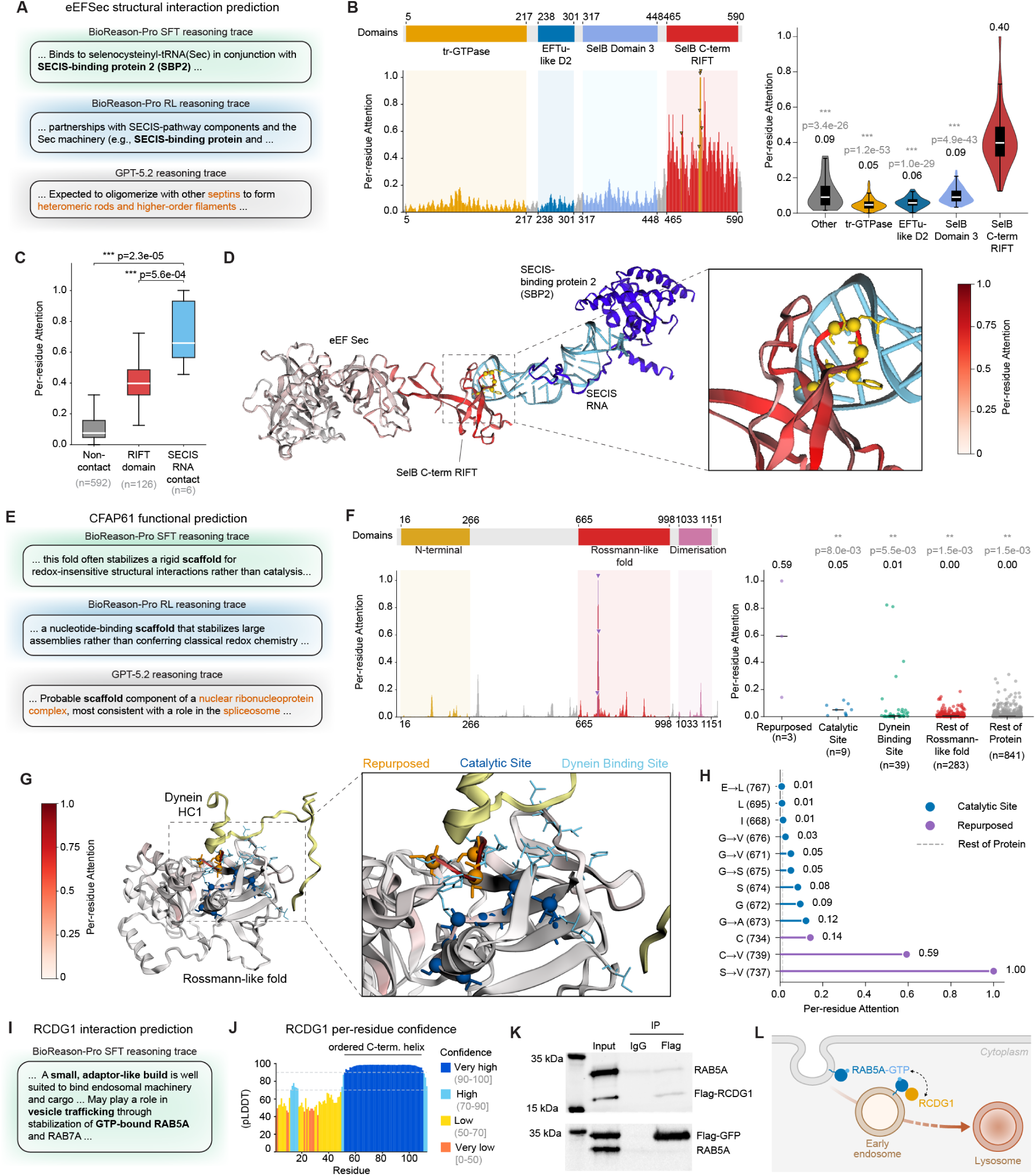
BioReason-Pro generates de novo functional predictions across three proteins. **(A–D) eEFSec. (A) Reasoning traces.** Key passages from BioReason-Pro SFT, BioReason-Pro RL, and GPT-5.2 Thinking High. **(B) Per-residue attention.** Attention is enriched at the C-terminal RIFT domain. **(C) Contact interface enrichment.** Per-residue attention across non-contact, RIFT domain, and SECIS RNA contact residues. **(D) Structural context.** Cryo-EM selenosome with eEFSec colored by per-residue attention, highest at the RIFT surface overlapping the SECIS RNA interface. **(E–H) CFAP61. (E) Reasoning traces.** Key passages from BioReason-Pro SFT, BioReason-Pro RL, and GPT-5.2 Thinking High. **(F) Per-residue attention.** Attention is enriched at the three repurposed active-site residues. **(G) Structural context.** Cryo-EM axoneme with the Rossmann domain colored by per-residue attention, highest at the repurposed residues forming the dynein interface. **(H) Per-residue attention at catalytic positions.** Attention for individual catalytic and repurposed positions. **(I–L) RCDG1. (I) Reasoning trace.** Key passage from the BioReason-Pro SFT generation. **(J) Per-residue confidence.** AlphaFold3 per-residue pLDDT for RCDG1, a disordered N-terminus with a single ordered C-terminal helix. **(K) Co-immunoprecipitation.** Flag-RCDG1 pulls down endogenous RAB5A, whereas Flag-GFP does not. **(L) Proposed model.** RCDG1 acts at the RAB5A node of the early endosome.

Selenocysteine incorporation requires coordination between the elongation factor, a dedicated mRNA stem-loop in the 3’-UTR (SECIS), and an accessory protein that bridges these elements at the ribosome (Caban and Copeland, 2006). When no protein interactions were provided in context, BioReason-Pro SFT reconstructed this three-component logic from architectural constraints alone, reasoning that the C-terminal RIFT domain implies an escorting factor that connects the SECIS element to the decoding site, and identified SECIS-binding protein 2 (SBP2) as this partner (**Fig. 5A**). Indeed, SBP2 is the most specific functional partner of eEFSec, confirmed by co-immunoprecipitation and visualized in the 2.8 Å cryo-EM selenosome structure, which revealed that the interaction is mediated by the SECIS element (Copeland et al., 2000; Hilal et al., 2022). BioReason-Pro RL also identified SECIS-binding protein as a partner (Section C.7). In contrast, GPT-5.2 Thinking High misidentified this protein entirely as a septin-family GTPase and predicted cytokinesis and cytoskeletal scaffolding functions (Section C.8).

To investigate whether this de novo prediction is grounded in the protein representation, we examined BioReason-Pro SFT’s per-residue attention from the token immediately preceding the predicted partner name to the protein sequence, since this variant produced the most specific partner prediction (**Fig. 5B**). Attention localizes sharply to the RIFT domain, with all other domains showing significantly lower scores (tr-GTPase *p* = 1.2 × 10^−53^, EFTu-like D2 *p* = 1.0 × 10^−29^, SelB Domain 3 *p* = 4.9 × 10^−43^; **Fig. 5B**). At residue resolution, attention scores at SECIS RNA contact residues are significantly elevated relative to non-contact residues (Mann–Whitney *U*, *p* = 2.3 × 10^−5^; **Fig. 5C**) (Mann and Whitney, 1947). Projection onto the cryo-EM selenosome structure (PDB 7ZJW) confirms that the highest-attention surface on the RIFT domain coincides with the SECIS RNA binding interface (**Fig. 5D**) (Hilal et al., 2022). The model thus predicted the principal functional partner of eEFSec from sequence alone, and its internal representation concentrated on the precise binding surface that structural biology identified through the selenosome reconstruction. The model further reconstructed the full selenocysteine incorporation cycle with mechanistic accuracy, describing GTP-coupled A-site docking at UGA codons, SECIS-programmed context-dependent decoding, and GTP hydrolysis triggering factor dissociation to commit the charged tRNA to peptide bond formation (Simonović and Puppala, 2018; Hilal et al., 2022).

This case demonstrates that BioReason-Pro can generate predictions that exceed existing curated annotations in both specificity and structural grounding. The model de novo identified SBP2 as the obligate functional partner, a prediction validated by the cryo-EM selenosome structure (Hilal et al., 2022; Dobosz-Bartoszek et al., 2016). The significant overlap between attention and the experimentally resolved contact interface provides direct evidence that these predictions are rooted in structurally meaningful protein features. These results suggest that explicit chain-of-thought reasoning over learned protein representations can produce functional insights that refine and extend expert-curated database entries.

### 2.7. BioReason-Pro performs structural reasoning beyond domain annotation transfer

A central question for BioReason-Pro is whether it performs genuine reasoning or merely restates domain annotations in natural language. To evaluate this, we systematically screened test set generations for cases in which naive domain interpretation would predict incorrect function but the model arrived at the correct annotation through contextual reasoning. One such case is the Cilia- and flagella-associated protein 61 (CFAP61, UniProt Q8NHU2), a 1,237-residue axonemal protein whose closest training set match shares only 29% sequence identity, well below the threshold where homology transfer provides reliable annotations (Rost, 1999). This protein presents a specific interpretive challenge because its domain architecture contains a Rossmann-like FAD/NAD(P)-binding superfamily fold (IPR036188) that would cue enzymatic or oxidoreductase function in standard domain-lookup pipelines (Heuser et al., 2012; Blum et al., 2024). Indeed, some databases incorrectly list oxidoreductase activity as a Gene Ontology annotation for this protein (Stelzer et al., 2016). However, experimental characterization has established CFAP61 as a non-enzymatic scaffold of the calmodulin- and radial-spoke-associated complex, essential for sperm flagellum formation (Liu et al., 2021; Ma et al., 2022).

BioReason-Pro resolves this ambiguity through contextual architectural reasoning rather than isolated domain interpretation. The model observes that the Rossmann-like fold is situated between an N-terminal axonemal targeting module (IPR032151) and a C-terminal dimerization domain (IPR056299), all within a cilia-specific protein family (IPR038884) (Blum et al., 2024). From this context, it infers that the Rossmann-like domain contributes a stable structural core for redox-insensitive scaffolding interactions rather than catalysis (**Fig. 5E**). This contextual override of a superfamily-level annotation, where interpretation is conditioned on flanking domains, overall protein architecture, and family membership, demonstrates reasoning that cannot be replicated by domain lookup or homology transfer alone (full trace in Section C.9). BioReason-Pro RL also correctly identified the non-catalytic scaffolding role but with less functional specificity (Section C.10). In contrast, GPT-5.2 Thinking High misidentified CFAP61 as a nuclear spliceosome-associated HEAT-repeat protein and predicted mRNA splicing and nuclear localization (Section C.11).

To determine whether there is a structural basis for BioReason-Pro’s inferences, we aligned the CFAP61 Rossmann domain against glutathione reductase (PDB 3GRS) (Karplus and Schulz, 1987) and identified 12 positions at catalytic and cofactor-binding sites. All 12 are degenerate in CFAP61. The GxGxxG dinucleotide-binding loop is disrupted (VGASSV at positions 671–676); the catalytic nucleophile CysI is replaced by Val at position 739; the FAD-stacking Glu is replaced by Leu at position 767; and the cofactor-binding Ser is replaced by Val at position 737. Three of these former catalytic positions (C734, V737, V739) now form direct contacts with dynein heavy chain 1 (DHC1) in the cryo-EM axoneme structure (PDB 8J07) (Walton et al., 2023), with the ancestral cofactor-binding pocket occupied by the dynein interface. We examined per-residue attention from the token at which the model predicts scaffolding over catalytic function (**Fig. 5F**). Attention is significantly enriched at the three repurposed residues relative to remaining catalytic positions (*p* = 8.0 × 10^−3^), dynein binding site residues (*p* = 5.5 × 10^−3^), and the rest of the Rossmann domain (*p* = 1.5 × 10^−3^; Mann–Whitney *U*; **Fig. 5F**) (Mann and Whitney, 1947). Projection onto the cryo-EM structure confirms that the highest-attention surface on the Rossmann domain coincides with the dynein binding interface (**Fig. 5G**). At single-residue resolution, S→V at position 737 and C→V at position 739 showed the highest attention scores, both of which are direct DHC1 contacts (**Fig. 5H**). This suggests that the model has learned structural reasoning, favoring protein interaction over catalytic activity at the precise positions where sequence variation supports this functional shift.

The model also identified CFAP61 as a non-enzymatic axonemal scaffold involved in cilium assembly (GO:0060271) and cilium movement (GO:0003341), localizing it to the axoneme (GO:0005930) and motile cilium (GO:0031514). It predicted protein dimerization activity (GO:0046983) from the C-terminal dimerization domain and structural molecule activity (GO:0005198), both consistent with the experimentally established role of CFAP61 as a radial-spoke-associated factor required for axonemal integrity (Liu et al., 2021; Ma et al., 2022). BioReason-Pro also predicted specific binding partners including axonemal dynein subunits, radial spoke proteins, and tubulins. Dynein heavy chains are direct structural contacts of the CFAP61 Rossmann domain in the cryo-EM axoneme reconstruction (PDB 8J07) (Walton et al., 2023), while RSPH9 and TUBB3 are validated co-complex members by co-immunoprecipitation (Liu et al., 2021).

This case demonstrates reasoning at two scales. At the architectural level, the model conditions domain interpretation on flanking context and family membership to override a misleading superfamily annotation. At the residue level, attention concentrates on degenerate catalytic positions that have been evolutionarily repurposed as a protein-protein interaction surface. These results suggest that explicit chain-of-thought reasoning over learned protein representations captures structural and evolutionary features sufficient to extend functional characterization beyond existing database annotations, even for proteins far outside the reach of homology-based methods.

### 2.8. Experimental validation of a proposed interaction nominates a function for renal cell carcinoma biomarker RCDG1

Renal cancer differentiation gene 1 (RCDG1, also C4orf46; UniProt Q504U0) is a biomarker of renal cell carcinoma, the most common form of kidney cancer, where it is downregulated in tumor tissue relative to normal kidney and localizes to the cytoplasm of proximal and distal tubule epithelium (Yu et al., 2014; Hsieh et al., 2017; Uhlén et al., 2015). RCDG1 is a small 113-residue protein with a single family-level signature (IPR031457), a disordered N-terminal region, and no catalytic or functional domain, leaving structure-based and homology-based pipelines with nothing to transfer. Despite its clinical association, RCDG1 has no experimentally assigned molecular function in curated resources (Kelleher et al., 2023; Oprea et al., 2018), and the only additional clue is a mouse knockout that is dispensable for spermatogenesis and fertility (Shah et al., 2021), which assigns no mechanism.

BioReason-Pro interpreted the compact, domainless architecture of RCDG1 as that of a non-enzymatic adaptor acting through protein interactions rather than catalysis (**Fig. 5I**; full trace in Section C.13). From this premise it predicted a specific molecular function, Rab GTPase binding (GO:0017137), proposing that RCDG1 is a cytosolic adaptor that binds and stabilizes GTP-loaded Rab GTPases on early endosomes to tune receptor and cargo routing, consistent with how Rab effectors engage the active, membrane-associated state of their targets (Zerial and McBride, 2001). It named RAB5A, the master regulator of early endosomes, as a primary interaction partner (Bucci et al., 1992; Zeigerer et al., 2012). There are no curated interactions between RAB5A and RCDG1 (del Toro et al., 2022), making this a genuine de novo prediction.

Since this interaction appears in no database, we first asked whether it was structurally plausible using AlphaFold3 (Abramson et al., 2024). AlphaFold3 modeled RCDG1 as a largely disordered monomer with a single ordered C-terminal helix (**Fig. 5J**) and confidently reconstructed the active, GTP- and Mg^2+^-bound state of RAB5A, but could not resolve a confident RCDG1 interface (high inter-chain PAE, **Fig. S10A**; interface ipTM 0.11, **Fig. S10B**), placing this interaction beyond what structure prediction alone can establish and motivating direct experimental testing.

We therefore tested the interaction experimentally by performing co-immunoprecipitation from a stable cell line expressing Flag-tagged RCDG1 with a reversible crosslinker to capture the potentially transient interaction (Lomant and Fairbanks, 1976). Immunoprecipitation of Flag-RCDG1 pulled down endogenous RAB5A. To control for non-specific binding, we also expressed Flag-GFP, and immunoprecipitation of it failed to pull down RAB5A (**Fig. 5K**). This demonstrates that RCDG1 associates with RAB5A, whether directly or through a shared complex, linking RCDG1 to the early endosome and the machinery that operates there, consistent with the role the model predicted.

The interactions already reported for RCDG1 place it in the vicinity of endosomal trafficking machinery, including the adaptor PICK1 (Fiuza et al., 2017; Hanley, 2008) and the endosomal sorting and lysosome-positioning proteins BLOC1S6 and KXD1 (Starcevic and Dell’Angelica, 2004; Pu et al., 2015; Setty et al., 2007). These partners were identified with no prior notion of what RCDG1 does (Luck et al., 2020; Rolland et al., 2014), and on their own they suggest only a broad association with this system. BioReason-Pro went further, singling out RAB5A, the master regulator of early endosomes, as a specific partner and placing RCDG1 at the RAB5A node of the early endosome (**Fig. 5L**). This sharpens a previously correlative link to renal cell carcinoma into a candidate mechanism, in which loss of RCDG1 could disrupt the endosomal control of receptor signaling and cargo fate that is recurrently broken in cancer (Mosesson et al., 2008; Mellman and Yarden, 2013), contributing to the differentiation and tumor phenotypes the gene was first associated with (Yu et al., 2014). More broadly, this case shows that BioReason-Pro can assign a precise, testable, and experimentally confirmed interaction to an uncharacterized protein (Kelleher et al., 2023; Oprea et al., 2018).

### 2.9. Pushing BioReason-Pro to its limits

To characterize the boundaries of BioReason-Pro’s reasoning capabilities, we evaluated both the SFT and RL models on short proteins and peptides. This represents a challenging setting, as fewer than 0.5% of training sequences fell below 50 amino acids. The evaluation comprised 22 sequences spanning 12 to 977 amino acids across metabolic, immune, antimicrobial, and venom categories, assessed by two independent domain experts. Model performance on full-length proteins including HLA-A*02:01, p53, full-length BRINP2, and IA-2 was consistent with the main benchmarks, accurately capturing domain architecture, molecular function, and biological process annotations. Performance on moderately sized sequences such as preproinsulin (110 aa), CCK (115 aa), and GDF15 (112 aa) was mixed. The model identified the correct protein families but consistently missed physiologically important functions such as feeding behavior, specific receptor partners, and appetite suppression. Below approximately 50 amino acids, performance degraded systematically.

Short peptide failures fell into three consistent categories. First, when short peptides lacked recognizable domains, the SFT model fabricated InterPro entries, assigning a myosin family domain to a GAD65 epitope, an SPFH domain to an IA-2 epitope, and ARID4B to a proinsulin junction peptide. In each case, the model built elaborate but baseless mechanistic narratives on these fabricated domains. The RL model never fabricated domains and instead explicitly acknowledged the absence of recognizable signatures. Second, the model confused closely related peptide family members when sequence length was too short. GLP-1 (30 aa) was conflated with glucagon, resulting in predictions of negative regulation of insulin secretion. Third, the model reproduced training-era annotations faithfully but could not identify functions characterized after the data cutoff. Full-length BRINP2 was accurately characterized as a nuclear transcriptional repressor, consistent with its training annotations, but its recently discovered role as a prohormone precursor for BRP (Coassolo et al., 2025) was not identified. When the 12-amino-acid BRP peptide was tested independently, it predicted generic transcriptional regulator annotations with no connection to appetite regulation or GPCR signaling.

A natural application of BioReason-Pro is functional annotation of proteins designed by generative models such as RFdiffusion (Watson et al., 2023) or the Evo series (Nguyen et al., 2024; Brixi et al., 2026). As a showcase, we selected two AI-generated anti-CRISPR (Acr) proteins, EvoAcr1 (169 aa) and EvoAcr2 (157 aa), designed by prompting Evo 1.5 with known Cas9-targeting Acr operons (Merchant et al., 2025). Five of the resulting designs blocked Cas9 activity in *E. coli*, and EvoAcr1 and EvoAcr2 were chosen because they lack sequence homology to any known protein by BLAST and yielded only low-confidence structural predictions from AlphaFold3 (Abramson et al., 2024). These proteins have no InterPro domains and no representation in any training database. Because they are synthetic and have no natural species, organism assignment is ambiguous. We used *E. coli* as the primary organism since both proteins are functional in that host, and additionally tested specific strains (K12, O157:H7) and *Homo sapiens* to assess organism sensitivity. Full reasoning traces are in Sections C.12–C.16.

Both models generated detailed reasoning traces with mechanistic narratives and testable hypotheses despite the complete absence of homology or domain evidence. As noted above for the peptides, the SFT model also fabricated InterPro entries. That said, several predictions by both models were biologically coherent. For EvoAcr2 in generic *E. coli*, the SFT model described a phage-encoded effector that binds the nucleoid-associated proteins H-NS and HU to remodel chromosomal architecture. For EvoAcr2 in O157:H7, the RL model predicted a soluble phage-encoded adaptor that operates in the host cytoplasm, using multivalent binding to sequester host factors and reorganize protein complexes governing transcriptional output, ultimately dampening host gene expression to favor a viral life cycle. Both predictions are consistent with phage-encoded effectors that modulate host biology (Pawluk et al., 2017; Hwang and Maxwell, 2023). Why these phage-related functions were predicted specifically for the pathogenic strain remains unclear.

For these AI-generated proteins, predictions varied substantially across organism labels for the same sequence. The same EvoAcr1 sequence was predicted as a ribosomal protein in generic *E. coli*, a DNA-binding transcriptional repressor in K12, a host-pathogen membrane effector in O157:H7, and a translation initiation factor in human. EvoAcr2 showed similar divergence, ranging from an acid stress resistance protein in K12 to a peroxiredoxin in human. This strain-level sensitivity has a biological basis, as *E. coli* O157:H7 expresses over 1300 proteins not found in K12, many related to virulence (Da Silva et al., 2018), likely shaping a bias toward pathogenesis-related predictions. GO-GPT showed the same organism-driven divergence upstream, indicating that the effect propagated through the full pipeline. While the strong influence of organism context on predictions for novel proteins remains a limitation, BioReason-Pro was clearly capable of nominating plausible and testable molecular functions.

## 3. Discussion

Here we introduce GO-GPT and BioReason-Pro, two models that together establish expert-level protein function prediction through structured biological reasoning. GO-GPT is the first autoregressive transformer for Gene Ontology prediction, achieving state-of-the-art 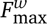 by capturing hierarchical and cross-aspect dependencies that discriminative methods miss. BioReason-Pro builds on GO-GPT as the first multimodal reasoning LLM for protein function prediction, deeply integrating protein representations with biological context to generate interpretable reasoning traces that progress from domain analysis to functional hypotheses. Human protein experts preferred BioReason-Pro annotations over curated UniProt entries in 79% of evaluated cases, and the model maintained strong performance even for proteins with low sequence similarity to training data, demonstrating generalizable functional reasoning. Together, these models open a path toward scalable characterization of the millions of proteins across all domains of life that lack functional annotation.

The computational biology community has invested heavily in foundation models that encode biological information, from protein language models such as the ESM family (Hayes et al., 2024) to structure predictors like AlphaFold (Fleming et al., 2025), yet comparatively little effort has addressed how to reason over these representations to perform the integrative synthesis that expert biologists conduct. BioReason-Pro reasons from structural and evolutionary principles across protein representations and biological context, enabling generalization across the full range of sequence similarity. The model’s advantage over BLAST widened with decreasing sequence identity, with stable performance across diverse organisms and protein lengths. Even at high similarity where close homologs were readily available, BioReason-Pro outperformed BLAST, indicating that biological reasoning captures functional relationships beyond sequence alignment alone.

Notably, reinforcement learning improved GO term accuracy and reduced hallucinated claims while producing shorter and more focused reasoning traces, without significantly reducing overall expert preference. Where individual preferences diverged, the separation was driven by mechanistic depth favoring SFT and hallucination avoidance favoring RL. Optimizing for ontology accuracy disciplines the model against fabricated biological claims but may limit the depth of mechanistic hypotheses the model generates. This tension likely reflects the reward signal itself, as 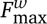 captures ontology-level correctness but does not measure the mechanistic detail or explanatory quality that distinguishes expert-level annotation. Designing reward signals and evaluation frameworks that capture biological reasoning quality beyond ontology metrics remains an important open problem for the field.

A key capability demonstrated by BioReason-Pro is structural and mechanistic reasoning that extends beyond existing database annotations, down to individual residue resolution. For eEFSec, the model predicted SBP2 as the obligate functional partner, a protein that bridges eEFSec to the SECIS mRNA element through the RIFT domain (Hilal et al., 2022). Notably, per-residue attention at the partner prediction token localized to this contact interface in the cryo-EM selenosome structure. The model also accurately reconstructed the full selenocysteine incorporation cycle (Simonović and Puppala, 2018; Hilal et al., 2022), predicting selenocysteine-specific tRNA recognition as the defining molecular function, a distinction from all other EF-Tu family members that the Gene Ontology has not yet formalized into a dedicated term (Ashburner et al., 2000). CFAP61 presents the complementary case, where the Rossmann-like fold contains a catalytic site that typically confers enzymatic activity (Karplus and Schulz, 1987), yet these positions are evolutionarily degenerate, with three now forming direct contacts with DHC1 in the cryo-EM axoneme structure (Walton et al., 2023). BioReason-Pro recognized this degeneration by specifically attending to the three repurposed dynein contact residues and predicted a non-enzymatic scaffolding role consistent with experimental characterization.

Furthermore, BioReason-Pro can predict biology that no database records and that holds up under direct experimental test. RCDG1 (C4orf46) is a biomarker of renal cell carcinoma, the most common kidney cancer, yet its molecular function has remained unknown and it carries no domain for annotation transfer to exploit. With no interactions supplied, the model inferred that RCDG1 is a cytosolic adaptor for early-endosomal Rab GTPases and nominated RAB5A (Bucci et al., 1992) as a partner. To the best of our knowledge, there is no curated record of an interaction between these proteins, and we verified this by co-immunoprecipitation. This prospective validation of a de novo prediction for a dark-genome protein supports the potential of BioReason-Pro as a hypothesis generator. It nominates a mechanism by which RCDG1 loss could contribute to renal cell carcinoma through dysregulated endosomal trafficking.

BioReason-Pro has several important limitations. The model was trained on synthetic reasoning traces generated by GPT-5 (Singh et al., 2025), which may contain subtle reasoning errors that propagate into the model. Furthermore, training requires proteins with experimental GO annotations, a resource that remains costly and limited in throughput to produce. Reasoning quality is heavily influenced by the availability of recognizable protein domains and degrades for proteins that lack identifiable InterPro annotations (Blum et al., 2024). Performance also degrades for extremely short peptides below 50 amino acids, where limited sequence information constrains the model’s ability to ground its functional predictions. For synthetic proteins that lack identifiable domains, such as the EvoAcr sequences (Merchant et al., 2025), predictions become heavily dependent on the organism label, producing divergent functional annotations and interaction partners across organisms for the same sequence. That said, several of these predictions were biologically coherent with known phage-encoded effector biology, suggesting that BioReason-Pro can nominate plausible hypotheses even in this challenging regime. LLM-based evaluation with GPT-5.1 may harbor systematic biases, and human expert evaluation covered 162 proteins, a sample size that limits statistical power for fine-grained comparisons. The model is also computationally expensive, requiring sequential inference through ESM3 (Hayes et al., 2024), GO-GPT, and the reasoning LLM. Finally, whether BioReason-Pro learns genuine biological reasoning or sophisticated imitation of reasoning patterns remains an open scientific question.

Protein databases contain millions of sequences with unknown or computationally inferred function, and experimental characterization remains slow and expensive. BioReason-Pro offers a path forward by taking a protein sequence and reasoning about its function, serving as a first-pass annotator for newly sequenced proteins, a hypothesis generator that proposes testable mechanisms, or a quality-check system for existing database annotations. This is particularly valuable for metagenomic proteins, disease-associated proteins that lack mechanistic characterization, and sequences lacking close homologs where traditional annotation transfer fails. The reasoning traces expose the logic underlying each prediction, allowing researchers to assess whether conclusions about protein function are biologically sound and to engage with model outputs critically rather than accepting or rejecting them wholesale.

In this work, we showed that AI systems can reason about protein function with sufficient fidelity that human experts prefer model outputs over curated database annotations. Proteins do not act in isolation, and extending this reasoning to cellular networks may enable models that predict how perturbations propagate through biological systems, how mutations cause disease, and how interventions restore health. The convergence of foundation models that encode biological knowledge with language models capable of reasoning over that knowledge establishes a new paradigm for computational biology, one where AI systems serve as genuine partners in biological discovery. We release all model weights, training code, and curated datasets alongside a web interface for broad accessibility (see Code and Model Availability for details). We also provide pre-computed predictions for over 240,000 proteins including the Human Protein Atlas to facilitate immediate application. We hope these materials enable the community to extend, critique, and build upon this work.

## 4. Methods

### 4.1. Datasets

#### 4.1.1. Dataset Pipeline

All protein annotations and auxiliary biological features used in this study were derived from publicly available sources and follow the same curation criteria as the CAFA5 benchmark (Friedberg et al., 2023b,a). The dataset integrates multiple biological modalities, including protein sequence, structure, domain, family, and interaction features, into a single coherent representation, with detailed statistics presented in **Fig. S1**.

Protein-level GO term annotations were obtained from the Gene Ontology Annotation (GOA) database (Consortium, 2021) (November 2022 release), which provides comprehensive mappings between UniProt accessions and GO terms under diverse evidence codes. Following CAFA standards, only annotations supported by experimental or curated evidence codes (IDA, IPI, EXP, IGI, IMP, IEP, IC, TAS) were retained to ensure high-confidence ground truth. Each annotation was propagated upward through the Gene Ontology hierarchy using is_a and part_of relations to maintain hierarchical completeness. GO term identifiers were standardized to the January 2023 ontology. Information Accretion (IA) weights from CAFA5 (Clark and Radivojac, 2013a; Friedberg et al., 2023b) were used unchanged to ensure consistency across all experiments.

Protein metadata, including amino acid sequences, organism, taxonomy identifiers, subcellular localization, and functional comments, were obtained from the UniProt KnowledgeBase (Consortium et al., 2025) (release 2023_01). Proteins with incomplete sequences or containing non-canonical residues were excluded. Structural information was retrieved from UniProt cross-references to the Protein Data Bank (PDB) (Burley et al., 2018). For proteins lacking experimentally resolved structures, AlphaFold2 (Fleming et al., 2025; Jumper et al., 2021) predicted models were used as substitutes. Domain annotations were integrated from InterPro (Blum et al., 2024), including InterPro identifiers, entry names, domain types, and residue coordinates. These domain and structure features provide complementary information on subunit organization and sequence-function relationships.

Protein-protein interaction (PPI) data were obtained from STRING v12.0 (Szklarczyk et al., 2024). UniProt accessions were mapped to corresponding STRING identifiers using the official mapping tables. Only high-confidence interactions (combined score ≥ 700) were retained, and for each protein the top 10 interaction partners by combined score were selected. Each protein’s interaction context was represented both as a compact list of interaction partners and as a structured table containing evidence-channel scores (neighborhood, coexpression, experimental, database, and text-mining). This preserves both topological and evidence-level information about the protein’s interaction environment.

Protein names in the PPI metadata were standardized by applying systematic pattern matching to extract only the primary protein name, removing all parenthetical content containing EC numbers, alternative names, synonyms, and organism-specific identifiers. Text preceding the first opening parenthesis was retained, preserving the primary functional descriptor while eliminating redundant database-specific annotations.

All modalities were merged into a unified dataset linking each protein to its sequence, organism, subcellular localization, structural features, domains, GO annotations, and interaction partners. Proteins with experimental annotations before November 2022 were randomly split 90/10 into training and validation subsets. For testing, we used a temporal holdout strategy consistent with CAFA (Zhou et al., 2019), retaining proteins that gained new experimental annotations between March 2023 and February 2024 but lacked annotations in the target aspect prior to the beginning of that period (Aleksander et al., 2023).

The resulting curated dataset provides a consistent and biologically grounded foundation for downstream modeling. It preserves the same protein and organism composition as the CAFA5 benchmark while enriching each protein instance with additional structural, domain, and network context.

#### 4.1.2. Reasoning Data

For each protein in the training data, we constructed a compact context comprising InterPro domains with identifier, name, and amino acid range, the UniProt protein description, organism, subcellular localization, PPI, and GO leaf terms across Molecular Function, Biological Process, and Cellular Component with their identifiers and names. This context was provided to GPT-5 (Singh et al., 2025) using the instruction prompt shown in Section C.1 to generate a structured reasoning trace that begins by identifying and analyzing the biology of protein domains, then reasoning over Molecular Function, Biological Process, and Cellular Component while naming relevant GO terms, next articulates a plausible mechanism for protein function, and finally proposes hypotheses about interaction partners grounded in the provided context. Each trace ends with a final answer containing a concise summary of protein function, the list of InterPro domains, the list of leaf GO terms for all three aspects, and an explicit hypothesis for PPI.

We generated traces for 130,492 training proteins, averaging over 1,100 tokens per trace for a total of approximately 140 million tokens (**Fig. 1E**). The model was instructed to rely only on the supplied context. Manual spot checks on several hundred randomly selected proteins showed adherence to this constraint with no observed hallucinations.

#### 4.1.3. Temporal Holdout Test Set

We constructed a temporal holdout test set following the CAFA experimental framework (Zhou et al., 2019) to evaluate model generalization on proteins with newly acquired functional annotations. Protein sequences were downloaded from UniProt KnowledgeBase (Consortium et al., 2025) and GO term annotations from the GOA database (Consortium, 2021; Aleksander et al., 2023) at two time points, November 2022 and February 2024. Using the same experimental evidence codes described in Section 4.1, we retained proteins that gained new experimental annotations between March 2023 and February 2024 but lacked annotations in the target aspect prior to the beginning of that period, encompassing both no-knowledge proteins (no prior experimental annotations in any aspect) and limited-knowledge proteins (annotations in other aspects only). We exclude limited-knowledge proteins from BioReason-Pro evaluation because the training reasoning traces are generated from annotations across all known aspects, and cross-aspect reasoning by GPT-5 can implicitly convey information about the held-out aspect, inflating test performance.

All experimental annotations were propagated through the GO hierarchy to the root using is_a and part_of relations to maintain ontological completeness. The final temporal holdout test set comprises 8,630 unique proteins with an average of 26.75 GO terms per protein (median 18.0) and 230,824 total annotations. By ontology aspect, the test set contains 5,819 proteins with Biological Process annotations (169,459 terms, 73.4%), 2,080 proteins with Molecular Function annotations (19,773 terms, 8.6%), and 3,440 proteins with Cellular Component annotations (41,592 terms, 18.0%).

### 4.2. GO-GPT Model

#### 4.2.1. Architecture

GO-GPT is a unified autoregressive architecture for protein function prediction that models all three GO aspects, Molecular Function (MF), Biological Process (BP), and Cellular Component (CC), within a single model (**Fig. 2A**) (Aleksander et al., 2023). Aspect delimiter tokens (<|MF_START|>, <|BP_START|>, <|CC_START|>) explicitly condition the model on the ontology branch being generated while allowing it to learn cross-aspect correlations within a shared embedding space.

The model combines a frozen protein language model encoder with a custom GPT-style transformer decoder. All experiments employ the ESM2 backbone (Lin et al., 2023), from which residue-level embeddings are extracted without gradient updates. Each residue embedding is projected into the decoder’s hidden dimension via a two-layer feed-forward projection with GELU activation (Hendrycks and Gimpel, 2023) and dropout (Srivastava et al., 2014), aligning the frozen ESM2 representation space with the autoregressive GO token space. We used ESM2 as it performed on par with ESM3 (Hayes et al., 2024) in GO-GPT, while enabling faster training due to native support for batch embedding. Architectural and training explorations are detailed in Appendix B.3.

The decoder is a 12-layer, 12-head transformer (Vaswani et al., 2023) adapted from the open-source nanoGPT implementation (Karpathy, 2022), substantially modified to support prefix-causal attention (Dong et al., 2019). This mechanism defines separate attention patterns for the protein and GO token streams. Protein residue embeddings attend bidirectionally to all other protein positions, enabling contextualized residue representations that capture long-range sequence dependencies. GO tokens attend causally to all preceding GO tokens and cross-attend to all protein residue positions. Protein tokens do not attend to GO tokens, ensuring the protein representations remain independent of the generation state. This asymmetric attention pattern is illustrated in Table 1. Each transformer block maintains separate query-key-value projections, output projections, layer normalizations, and feed-forward networks for the protein and GO streams, allowing the model to learn modality-specific transformations while sharing information through the cross-attention pathway (Bahdanau et al., 2016). Each attention layer also applies a sigmoid output gate **Y**^′^ = **Y ʘ** *σ* **XW**_*g*_ (Qiu et al., 2025), which provided minor improvements to training stability and validation metrics.

**Table 1.** Prefix-causal attention mask pattern. Checkmarks indicate allowed attention; crosses indicate masked positions. Protein residues (*r*_*i*_) attend bidirectionally to each other but are blocked from GO tokens. GO tokens (*t* _*j*_) cross-attend to all protein residues and causally attend to preceding GO tokens.

| Query ↓ \ Key → | Protein tokens | | | | GO tokens | | | |
| --- | --- | --- | --- | --- | --- | --- | --- | --- |
| | $r_1$ | $r_2$ | $r_3$ | $r_4$ | $t_1$ | $t_2$ | $t_3$ | $t_4$ |
| Protein $r_1$ | ✓ | ✓ | ✓ | ✓ | × | × | × | × |
| Protein $r_2$ | ✓ | ✓ | ✓ | ✓ | × | × | × | × |
| Protein $r_3$ | ✓ | ✓ | ✓ | ✓ | × | × | × | × |
| Protein $r_4$ | ✓ | ✓ | ✓ | ✓ | × | × | × | × |
| GO $t_1$ | ✓ | ✓ | ✓ | ✓ | ✓ | × | × | × |
| GO $t_2$ | ✓ | ✓ | ✓ | ✓ | ✓ | ✓ | × | × |
| GO $t_3$ | ✓ | ✓ | ✓ | ✓ | ✓ | ✓ | ✓ | × |
| GO $t_4$ | ✓ | ✓ | ✓ | ✓ | ✓ | ✓ | ✓ | ✓ |

The decoder vocabulary comprises the pruned GO term set (1,470 MF, 7,500 BP, 1,007 CC, totaling ≈9,977 tokens) plus seven structural delimiters. Organism embeddings are added to each GO token representation, enabling species-specific conditioning during decoding (Fallahpour et al., 2025a). Learnable positional embeddings encode the depthwise order of GO tokens in the ontology graph, preserving hierarchical ordering and facilitating generalization from shallow, well-annotated terms to deeper, sparsely annotated ones.

Generation is conditioned on per-residue embeddings concatenated with aspect-delimited GO tokens

[*r*_1_, *r*_2_,…, *r*_*L*_] <|ASPECT_START|> *t*_1_, *t*_2_,…, *t*_*n*_ <|ASPECT_END|>,

where *r*_*i*_ denotes the embedding of the *i*-th amino acid residue and *t* _*j*_ a GO token.

#### 4.2.2. Data Processing and Training

Each training instance consists of a protein sequence, its organism identifier, and associated GO term lists per aspect. Preprocessing tokenizes amino acid sequences using the ESM2 tokenizer (Lin et al., 2023), encodes GO terms with a custom aspect-aware tokenizer, and maps organisms to integer identifiers, retaining the top 200 most frequent species in the training set. GO vocabularies are pruned to include only terms appearing in at least 20 proteins per aspect, yielding ≈10K GO tokens across all aspects, ensuring sufficient training signal for each term. Within each aspect, GO terms are sorted by ontology depth (longest root-to-term path) so that the model generates terms from general to specific during autoregressive decoding.

Training uses a next-token prediction objective under teacher forcing, minimizing cross-entropy loss. Optimization follows the AdamW algorithm (Kingma and Ba, 2017; Loshchilov and Hutter, 2019) with learning rate 1×10^−4^, cosine decay (minimum ratio 0.1) (Loshchilov and Hutter, 2017), 10% warm-up, and weight decay 0.01. Gradients are clipped to 1.0. Mixed-precision (bf16) training and distributed data parallelism are applied across four H100 GPUs with gradient accumulation, yielding an effective batch size of 160. Training runs for up to 100 epochs with validation every epoch, and the best checkpoint is selected by validation performance (**Fig. S9**). Full hyperparameters are reported in Appendix B.6.

#### 4.2.3. Inference

During inference, protein embeddings are computed once and cached. Because GO-GPT is trained to predict one aspect at a time, each test protein is evaluated per aspect by prompting the model with the corresponding aspect start token and generating predictions for that aspect.

We consider two inference strategies. In greedy decoding, the model generates a single deterministic sequence per aspect, and all generated terms are assigned a probability of 1 while all others are assigned 0. In sampling-based inference, we generate 10 independent samples per protein using temperature-controlled sampling (*T*=0.7, top-*κ*=20), with generation terminated upon reaching the aspect-end token (Holtzman et al., 2020). Per-term probability estimates are then computed as the fraction of samples in which each term appears. We additionally report a best-of-10 oracle analysis that selects, for each protein, the sample yielding the highest downstream score, providing an upper bound on the headroom from improved decoding or selection. Implementation uses PyTorch (Paszke et al., 2019) and PyTorch Lightning (Falcon and team, 2025) with Flash Attention (Dao, 2023; Dao et al., 2022) and bf16 mixed precision.

### 4.3. BioReason-Pro Model

#### 4.3.1. Architecture

BioReason-Pro is a multimodal LLM that integrates protein sequence, structure, and GO graph within a unified reasoning framework, inspired by BioReason (Fallahpour et al., 2025b) (**Fig. 1**A). The architecture comprises three primary components: a protein encoder, a GO graph encoder, and a language model backbone.

Let *s* = *α*_1_,…, *α*_*L*_ denote a protein sequence of length *L* ≤ 2,000 over the amino acid alphabet *A*, let G=*(V’ ɛ),* denote the Gene Ontology directed acyclic graph where is the set of GO terms and the set of is_a and part_of edges, and let *c* denote the textual biological context comprising organism, InterPro domain annotations, protein-protein interactions, and GO-GPT predictions.

##### Protein Encoder

The protein encoder is based on ESM3-1B (Hayes et al., 2024), a state-of-the-art protein language model capable of jointly encoding sequence and structure. For each input protein, ESM3-1B generates residue-level embeddings that capture local and global structural context. These embeddings are extracted from layer 38 of the 48-layer model, yielding one vector per residue. Each residue embedding is independently projected into the LLM’s hidden dimension via a two-layer MLP with GELU activation (Hendrycks and Gimpel, 2023) and dropout (Srivastava et al., 2014):

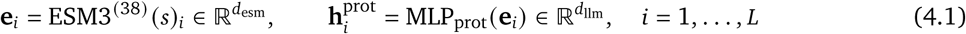

The full protein representation is the sequence of projected residue embeddings:

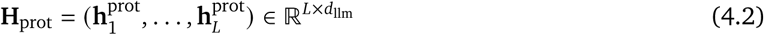

The protein encoder remains frozen throughout training, preserving its pretrained representations. Proteins longer than 2,000 amino acids are truncated.

##### GO Graph Encoder

The GO graph encoder processes the full GO graph (Consortium, 2021) to produce 200 fixed-length embedding vectors that capture hierarchical relationships and cross-namespace dependencies across all GO terms. Each embedding is projected into the LLM’s hidden dimension via a two-layer MLP:

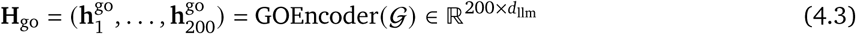

Full implementation details of the GOEncoder are provided in Section 4.3.2.

##### Language Model Backbone

The language model backbone uses Qwen3-4B-Thinking (Yang et al., 2025), a 4-billion parameter thinking model with built-in reasoning capabilities. The textual biological context *c* is tokenized and embedded through the LLM’s input embedding layer *E*_LLM_ (·):

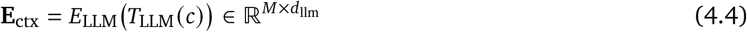

where *T*_LLM_ (·) is the language model tokenizer producing *M* tokens.

##### Multimodal Assembly

The *L* residue-level protein embeddings replace <|protein_pad|> placeholder tokens in the prompt, and the 200 GO graph embeddings replace <|go_graph_pad|> placeholder tokens (Liu et al., 2023). These are concatenated with the embedded textual context along the sequence dimension to form the full multimodal input:

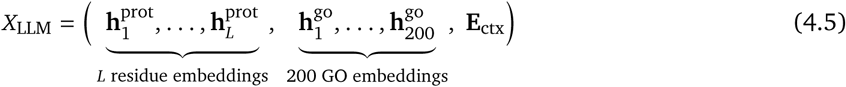

This formulation allows the LLM to attend to residue-level protein features, ontology-level knowledge, and biological context jointly when generating output. The model autoregressively generates a reasoning trace and functional annotations:

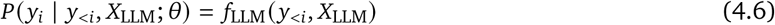

where the output sequence *Y* = *y*_1_,…, *y*_*K*_ contains structured biological reasoning followed by GO term predictions, functional summaries, and hypothesized interaction partners. Full architectural hyperparameters are reported in Table S17. Additional architectural design explorations that informed the final BioReason-Pro configuration are summarized in Section B.4.1.

#### 4.3.2. GO Encoder

The GO graph encoder is a hierarchical graph neural network designed to capture the semantic and structural properties of the GO directed acyclic graph *G = (V’ɛ)*, (Consortium, 2019). Hierarchical GNNs propagate information across parent-child relationships, enabling the model to share statistical strength between rare and common terms, capture multi-level dependencies, and preserve semantic consistency (Kim et al., 2021; Kabir and Shehu, 2024). The encoder processes all GO terms in a single unified graph without namespace separation, producing 200 fixed-length embeddings that are integrated into the LLM context.

##### Node Initialization

Each GO term *v* ɛ*V* is initialized by encoding its name and description into a single vector using the Qwen3-4B text embedding model (Zhang et al., 2025):

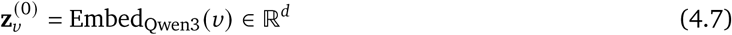

where *d* = 2560 is the embedding dimension. This initialization provides each node with a semantically rich starting representation grounded in the term’s functional definition.

##### GAT Propagation

The encoder applies three graph attention network (GAT) layers (Veličković et al., 2018) over the full ontology graph. At each layer *l* = 1,…, 3, the representation of term *v* is updated by attending to its neighbors *N(*v*)* defined by the is_a and part_of edges in *ɛ*. Each layer uses multi-head attention with *K* = 8 heads:

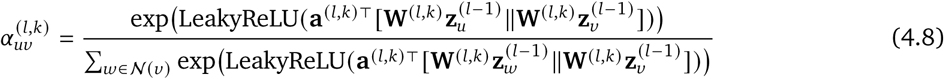

where **W**^(*l*,*κ*)^ and **a**^(*l*,*κ*)^ are the learnable weight matrix and attention vector for head *κ* at layer *l*, and || denotes concatenation. The multi-head outputs are concatenated and passed through GELU activation (Hendrycks and Gimpel, 2023) and dropout (Srivastava et al., 2014):

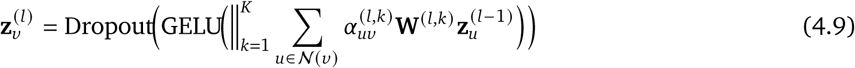

The first layer projects from the input dimension *d* to the hidden dimension (*d*_ℎ_ = 512, with *d*_ℎ_ *K* = 64 per head), while subsequent layers maintain the hidden dimension. After the final GAT layer, a two-layer MLP with GELU activation projects the representations back to the original embedding dimension:

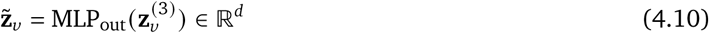

##### Cross-Attention Compression

To produce a fixed-length representation from the variable number of GO terms (|V| ≈ 43,000), we employ a cross-attention module inspired by the Perceiver architecture (Jaegle et al., 2021; Li et al., 2023). A set of 200 learnable query embeddings **Q** = (**q**_1_,…, **q**_200_) ∈ ℝ^200×*d*^ attend to the full set of processed node embeddings **Z̃** = (**z̃**_*v*_)_*v*∈V_ via multi-head cross-attention with *K* = 8 heads:

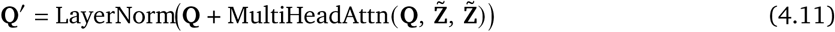

followed by layer norm (Ba et al., 2016) and a residual feed-forward block (He et al., 2015):

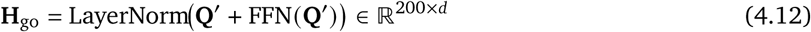

where FFN is a two-layer network with GELU activation and an expansion factor of 4. The resulting 200 embeddings 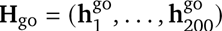 are then projected into the LLM’s hidden dimension via a two-layer MLP as described in Section 4.3.1 (Equation 4.3).

#### 4.3.3. Supervised Fine-tuning

BioReason-Pro is trained using a two-stage supervised fine-tuning (SFT) strategy that progressively integrates multimodal protein and ontology representations with the language model backbone. Training is performed on the reasoning traces generated by GPT-5 (Singh et al., 2025) as described in Section 4.1.2. Full hyperparameters for both stages are reported in Table S17 (Appendix B.7).

We denote the parameter groups as *θ*_esm_ for the protein encoder, *θ*_go_ for the GO graph encoder, *θ*_proj_ for the protein and GO projection layers, and *θ*_llm_ for the language model backbone. The protein encoder remains frozen throughout both stages, denoted *θ̄*_esm_. Both stages share the same training objective: causal language modeling with cross-entropy loss computed only over assistant tokens (Touvron et al., 2023), excluding user prompt tokens from the loss:

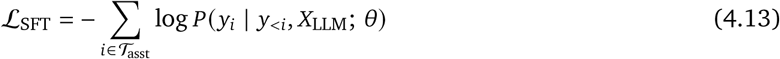

where *T*_asst_ is the set of token positions belonging to the assistant response and *X*_LLM_ is the multimodal input defined in Equation 4.5.

##### Stage 1: Modality Alignment

The first stage aligns the protein and GO graph representations with the LLM’s embedding space (Li et al., 2023). Only the GO graph encoder and projection layers are trainable, while both the protein encoder and the language model backbone remain frozen:

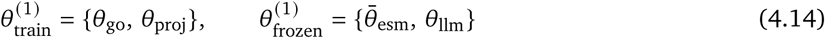

This stage runs for one epoch with a learning rate of 1 × 10^−4^, adapting the projection layers to map protein and ontology embeddings into the LLM’s representation space.

##### Stage 2: LLM Fine-tuning

The second stage unfreezes the LLM backbone and trains the complete model end-to-end for 10 epochs with a learning rate of 1 × 10^−4^. The LLM parameters are adapted using Low-Rank Adaptation (LoRA) (Hu et al., 2021), which decomposes each pretrained weight matrix **W**_0_ ∈ *θ*_llm_ as:

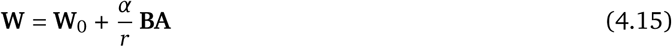

where **A ɛ** ℝ^*r*×*d*in^ and **B ɛ** ℝ^*d*out ×*r*^ are the learnable low-rank matrices, with rank *r* = 128 and scaling factor *α* = 256. LoRA is applied to all attention and MLP layers. The GO graph encoder and projection layers remain trainable, while the protein encoder and the pretrained LLM weights stay frozen:

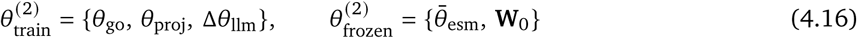

where Δ*θ*_llm_ = **A**, **B** denotes the set of all LoRA adapter parameters. The best checkpoint was selected at epoch 8 based on validation performance.

Both stages use the AdamW optimizer (Kingma and Ba, 2017; Loshchilov and Hutter, 2019) with cosine learning rate decay (Loshchilov and Hutter, 2017), a warmup ratio of 0.10 in Stage 1 and 0.05 in Stage 2, and weight decay of 0.01. Training is distributed across eight H100 GPUs on two nodes with a per-device batch size of 4, yielding an effective batch size of 32. The maximum protein length is 2,000 residues and the maximum text length is 10,000 tokens. The model was implemented using PyTorch (Paszke et al., 2019), PyTorch Lightning (Falcon and team, 2025), Hugging Face Transformers (Wolf et al., 2020), and Unsloth (Daniel Han and team, 2023). Additional supervised fine-tuning design explorations are summarized in Section B.4.2.

#### 4.3.4. Reinforcement Learning

Our RL algorithm is a hybrid variant of recent group-based policy optimization methods. We build on Group Relative Policy Optimization (GRPO) (Shao et al., 2024), which avoids training a value model by using group-wise reward statistics. To address the instability of token-level importance sampling weights identified by Zheng et al. (2025), we follow the core principle of Group Sequence Policy Optimization (GSPO), which aligns the unit of importance correction with the unit of reward at the sequence level. We additionally incorporate modifications inspired by Dr. GRPO (Liu et al., 2025) to correct optimization bias and mitigate length-dependent reward artifacts, and adopt the Clip-Higher strategy from DAPO (Yu et al., 2025) to enhance exploration. RL is initialized from the SFT epoch 8 checkpoint and trained with LoRA (Hu et al., 2021) at rank 16 and alpha 32, a learning rate of 3 × 10^−5^ with cosine decay, and a small KL penalty (*β* = 1 × 10^−4^). Full hyperparameters are reported in Table S18.

##### Algorithm Formulation

Let *π*_*θ*_ be an autoregressive policy model parameterized by *θ*, and let D denote the set of training queries (proteins). For each query *x ∈ D*, the old policy *π*_*θ*old_ generates a group of *G* responses:

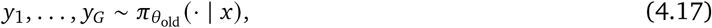

where each response *y*_*i*_ = *y*_*i*,1_,…, *y*_*i*, *yi*_ is a sequence of tokens and receives a scalar reward *r x*, *y*_*i*_ described below. We use a group size of *G* = 24 with rollouts generated at temperature *T* = 1.0. We process data in batches of *B* queries. Let denote the set of all *x*, *y* pairs in the current batch, so = *B G*, and let *L*_max_ denote the maximum response length in tokens.

##### Reward Computation

For each response *y*_*i*_, GO term identifiers are extracted from the reasoning trace using regular expressions, yielding a predicted term set *T*^_*i*_. These terms are propagated to parent terms through the GO hierarchy using is_a and part_of relations following the CAFA5 evaluation framework (Friedberg et al., 2023b). Since the extracted predictions are binary, the weighted 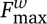 reduces to the IA-weighted F1 between the propagated predictions and the ground-truth term set *T*_*x*_ for protein *x*:

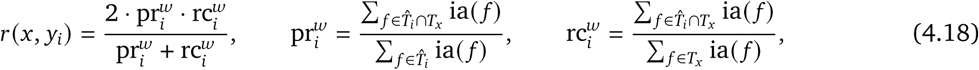

where ia(*f*) denotes the Information Accretion weight for GO term *f* (Clark and Radivojac, 2013b). This directly optimizes for the same weighted metric used in evaluation (Equation 4.28).

##### Advantage Estimation

Following GRPO and GSPO, we compute group-centered rewards. To improve stability in biological tasks with low-variance rewards, we normalize by the global batch standard deviation rather than per-group variance. For a fixed query *x* with responses *y*_1_,…, *y*_*G*_, we first compute the group mean reward:

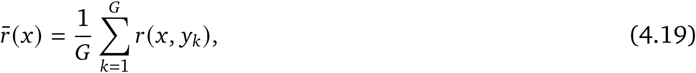

and then define the advantage for response *y*_*i*_ as:

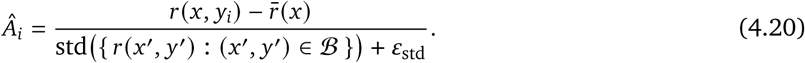

Here, std(·) denotes the standard deviation over all *B · G* rewards in the batch, and *ε*_std_ > 0 is a small constant for numerical stability. All tokens in *y*_*i*_ share the same advantage *Â*_*i*_, consistent with sequence-level rewards.

##### Sequence-Level Importance Ratio

As in GSPO, the off-policy correction is applied at the sequence level. The sequence likelihood of *y*_*i*_ under *π*_*θ*_ is:

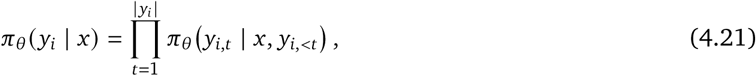

and similarly for *π*_*θ*old_. The sequence-level importance ratio is therefore:

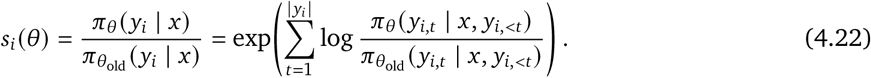

This ratio is controlled by asymmetric clipping parameters, *ε*_low_ and *ε*_high_, which implement the Clip-Higher strategy. This allows for a larger update range (1 + *ε*_high_) when advantages are positive to encourage exploration, while maintaining a strict clamp (1 − *ε*_low_) when advantages are negative for stability.

##### Clipped Surrogate Objective

We adopt the GSPO sequence-level clipped objective, with additional length regularization to counteract the length bias described in Dr. GRPO. The optimization objective is:

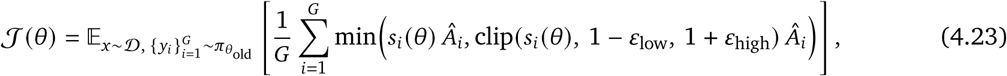

where clip(*s*, *α*, *b*) truncates the importance ratio *s* into the interval [*α*, *b*].

##### Training Loss

We minimize the negative of the above objective, normalized by a constant factor to produce a stable, unbiased learning signal:

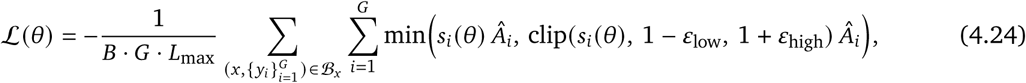

where *B_x_* ranges over the *B* queries in the batch. The normalization by *B G L*_max_ removes structural biases that could otherwise favor responses of a particular length.

Training runs for 1,200 steps with an effective batch size of 192 across eight H100 GPUs on two nodes, using DeepSpeed (Rasley et al., 2020) with vLLM (Kwon et al., 2023) in colocate mode for rollout generation. Additional RL explorations, including stability issues and reward-design choices, are summarized in Section B.4.3.

#### 4.3.5. Inference

At inference time, BioReason-Pro assembles a multimodal context for each query protein (**Fig. 1**A). Residue-level embeddings from ESM3-1B (Hayes et al., 2024) and the 200 GO graph embeddings are computed and inserted into the LLM context via their respective placeholder tokens. Textual context is constructed by concatenating the organism, InterPro domain annotations (identifiers, names, and residue ranges), and greedy-decoded GO-GPT predictions. Protein-protein interaction partners are optionally included. This assembled context serves as the user prompt, and the model generates a structured reasoning trace followed by a final answer containing functional summaries, GO term predictions, and hypothesized interaction partners. The exact prompt template used to assemble this multimodal context is provided in Section C.2.

We consider two inference strategies analogous to those used for GO-GPT. In greedy decoding (*T* = 0), the model generates a single deterministic trace per protein, and all GO terms appearing in the final answer are assigned probability 1. In sampling-based inference, we generate 10 independent traces per protein using temperature-controlled sampling (*T* = 1.0) and select the trace yielding the highest weighted 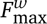 score as the best-of-10 oracle, providing an upper bound on performance from improved selection strategies. Per-term probability estimates can also be computed as the fraction of samples in which each GO term appears.

GO term identifiers are extracted from the final answer block of each generated trace using regular expressions and propagated to parent terms through the GO hierarchy following the CAFA5 framework (Friedberg et al., 2023b), as described in Section 4.4.1. The maximum generation length is 10,000 tokens. Inference is served using vLLM (Kwon et al., 2023) and takes approximately a few seconds per protein.

### 4.4. Evaluation

#### 4.4.1. GO Evaluation Metrics

GO term prediction performance is measured with the CAFA *F*_max_ metric (Zhou et al., 2019). For a set of *N* proteins, let *P*_*i*_ (*τ*) denote the set of GO terms predicted for protein *i* at score threshold *τ*, and let *T*_*i*_ denote the set of true GO terms. Protein-centric precision and recall at threshold *τ* are averaged across all proteins as

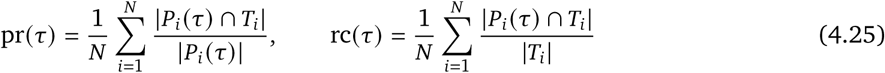

The unweighted *F*_max_ selects the threshold that maximizes the harmonic mean of these quantities

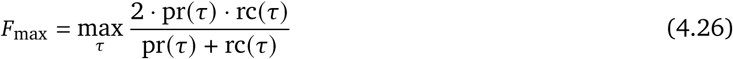

For the IA-weighted variant, each GO term *f* is assigned an Information Accretion weight ia(*f*) (Clark and Radivojac, 2013b) that increases with term specificity and rarity. Weighted precision and recall replace uniform term counts with IA-weighted sums

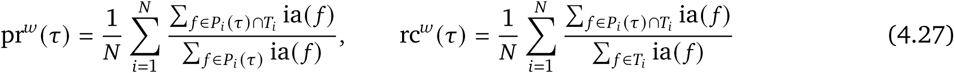

The weighted 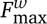 is then computed analogously to Equation 4.26 using pr^*w*^ and rc^*w*^

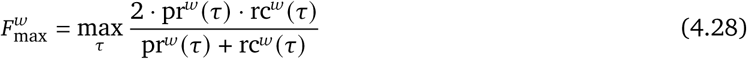

Both metrics are computed per aspect and reported separately. All evaluations use the official CAFA 5 evaluation toolkit (Piovesan et al., 2024), which applies IA weighting and hierarchy propagation to both predictions and ground truth prior to scoring (Clark and Radivojac, 2013b).

Because *F*_max_ is computed by sweeping a global threshold across all test proteins, it cannot be decomposed into per-protein contributions within individual subsets. To enable comparison across sequence similarity ranges (Section 4.4.5), we additionally report mean per-protein F1 within each bin. For a single protein *i* at the optimal global threshold *τ*^∗^, the per-protein F1 is

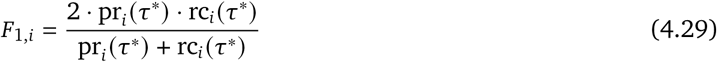

#### 4.4.2. Baselines

We evaluate GO-GPT and BioReason-Pro against multiple baselines spanning homology-based transfer, discriminative models, and generative language models. All baselines were trained or applied on the identical pre-November 2022 dataset and evaluated on the same temporal holdout test set.

For GO term prediction evaluated via *F*_max_ and 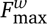, we compare against three baselines: InterLabelGO+ (Evans and Shen, 2024), which integrates deep learning predictions with sequence alignment and represented the previous state of the art; ProtBoost (Chervov, 2024), which combines protein language model features with gradient-boosted trees and graph neural networks; and BLAST-KNN, a homology-based baseline that transfers GO annotations from the closest training set hit as described below.

For functional summary evaluation via LLM judge and human expert scoring, we compare against Prot2Text-v2 (Fei et al., 2025), a multimodal model that generates natural language protein function descriptions from sequence and structural features, evaluated on the same test set using their released checkpoint with default inference settings. We also compare against BLAST free-text transfer as described below, reported in two variants. BLAST with penalty scores test proteins whose closest hit lacks a UniProt function description as zero. BLAST without penalty excludes such proteins from evaluation entirely.

All BLAST-based baselines and sequence similarity analyses use a single blastp (Altschul et al., 1990) run with standard parameters. The training set sequences serve as the database and the test set sequences as queries. For each test protein, the hit with the highest percent identity to the training set is selected. For GO term transfer, all GO terms from the best-matching training protein are assigned to the test protein with a score equal to the percent identity. The resulting predictions are evaluated using the CAFA evaluation framework (Friedberg et al., 2023b). For free-text transfer, the UniProt function summary and subcellular localization of the best-matching training protein are transferred directly.

#### 4.4.3. LLM-as-Judge Evaluation

We employ GPT-5.1 (Singh et al., 2025) as an automated expert judge to score model predictions against composite ground truth. For each test protein, the composite ground truth comprises the UniProt function summary, GO terms across all three aspects, InterPro domain annotations, known protein-protein interaction partners, organism, and subcellular localization. The judge receives this ground truth alongside a single model generation and produces a structured evaluation via a Pydantic schema-constrained output.

The output schema enforces five integer scores on a 1 to 10 scale and a short critique. Molecular Function, Biological Process, and Cellular Component each assess annotation correctness against their respective GO terms and supporting evidence. Specificity evaluates the depth and granularity of mechanistic detail. Reliability distinguishes logically grounded inference from hallucinated or contradicted claims. A score of 1 is returned for a functional axis only when all evidence sources are silent on that aspect. The critique field provides up to two sentences identifying specific mismatches between the prediction and ground truth.

The judge is called with deterministic decoding (temperature 0), a maximum output length of 512 tokens, default reasoning effort, and default verbosity. Evaluation is performed on the full temporal holdout test set for BioReason-Pro SFT, BioReason-Pro RL, Prot2Text-v2, and BLAST baselines (Section 4.4.2). The overall score for each model is the mean across the five axes, excluding any axis scored as 1. The full judge prompt and scoring rubric are provided in Appendix C.3.

#### 4.4.4. Human Expert Evaluation

To assess the scientific quality of model outputs beyond automated metrics, we conducted a blinded human expert evaluation. We recruited molecular biologists to independently evaluate model generations against curated ground truth annotations across 192 unique proteins randomly sampled from our test set. Both members of the BioReason team and external evaluators participated. To remove potential bias, we report results of the 162 proteins evaluated by the 27 external evaluators who were not members of the BioReason team.

Each evaluator was presented with a UniProt-derived Ground Truth dossier for a given protein, comprising the protein function summary, GO annotations, InterPro domains, known interaction partners, organism, and subcellular location. Alongside this dossier, two anonymized model generations were displayed: one from BioReason-Pro SFT and one from BioReason-Pro RL, randomly assigned as Model A and Model B. Evaluators were not informed which model produced which output.

The evaluation instrument comprises four parts, summarized below (Complete form in Supplementary C.4):

i. **Per-axis quantitative scoring** (Q1–Q10): ten dimensions rated on a 0–10 Likert scale applied independently to each model, covering molecular function accuracy, biological process accuracy, cellular component accuracy, reasoning and evidence attribution, plausibility of novel predictions, hallucination prevalence, hypothesis generation, mechanistic depth, protein–protein interaction predictions, and database-ready annotation quality.
ii. **Ordinal comparative judgments** (Q11–Q13): each model rated against the Ground Truth on a five-level ordinal scale, followed by a direct head-to-head preference judgment.
iii. **Free-text qualitative critique** (Q14–Q19): open-ended prompts eliciting commentary on key strengths, weaknesses, specific errors or hallucinations, reasoning quality, and comparison to expert expectations.
iv. **Meta-evaluation** (Q20, Evaluator Confidence): a case-study nomination flag and evaluator self-reported confidence on a 1–10 scale.

To quantify error prevalence from the free-text critiques, we used GPT-5-mini (Singh et al., 2025) to classify each evaluator’s written response into major error, minor error, or no error categories for each model independently. The classification prompt is provided in Appendix C.5 and the resulting error attributions stratified by expert preference are reported in **Fig. 4**G.

#### 4.4.5. Sequence Similarity Analysis

To evaluate model generalization as a function of homology to the training set, we stratified test proteins by best-hit BLAST sequence identity. For each test protein, the highest sequence identity against any training set sequence was obtained from the blastp search described in Section 4.4.2. Proteins were then binned into identity ranges. Proteins with no detectable BLAST hit were assigned to the lowest bin.

This stratification is applied consistently across all three evaluation modalities. For GO term prediction, we report mean per-protein F1 (Equation 4.29) within each bin, since *F*_max_ is a global metric that cannot be decomposed across protein subsets. For LLM-as-judge evaluation (Section 4.4.3), we report mean judge scores per bin. For human expert evaluation (Section 4.4.4), we report mean expert scores per bin.

#### 4.4.6. Statistical Comparison of GO Term Prediction Models

To assess statistical significance of performance differences between GO-GPT and baseline models, we used a paired bootstrap test. For each comparison, per-protein precision and recall arrays were computed for both models on the shared test set using CAFA-standard normalization (precision averaged over proteins with at least one prediction above threshold; recall averaged over all proteins). In each of 10,000 bootstrap iterations, proteins were resampled with replacement and the *F*_max_ difference was recomputed; the two-sided *p*-value was estimated as the fraction of bootstrap deltas with sign opposite to the observed difference. Both *F*_max_ and 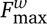 were tested. Because each inference mode involves 12 simultaneous tests (2 comparisons × 3 aspects × 2 metrics), *p*-values were corrected using the Holm–Bonferroni step-down procedure to control the family-wise error rate. Full bootstrap comparison tables are reported in Section B.5.1.

### 4.5. Interpretability

#### 4.5.1. GO-GPT Attention at Functional Sites

To evaluate whether GO-GPT grounds its predictions in functionally relevant sequence regions, we assessed the correspondence between model attention and experimentally determined DNA-binding residues. We identified proteins with experimentally determined DNA-binding sites from the BioLiP database (Yang et al., 2012) that were not present in the GO-GPT training set. DNA-binding sites were defined as BioLiP entries with ligand types DNA, DA, DC, DG, DT, DU, or DI. PDB binding residue positions were mapped to UniProt coordinates using per-residue SIFTS annotations (Burley et al., 2018; Consortium et al., 2025; Dana et al., 2019) extracted from mmCIF files, which correctly handles numbering gaps and insertion codes that segment-level SIFTS interpolation can miss. We retained proteins with at least 5 mapped binding residues and fetched their sequences from UniProt, yielding 304 candidate proteins. We then ran GO-GPT inference on each protein and retained the subset for which the model autoregressively generated the GO:0003677 (DNA binding) term, resulting in 63 evaluation proteins.

For each retained protein, we extracted cross-attention weights from the GO:0003677 output token to all input residue positions, averaged across all attention heads and transformer layers, producing a per-residue attention score vector. Discrimination between binding and non-binding residues was quantified using three complementary metrics. AUROC measures how well attention scores distinguish binding from non-binding residues, with its significance assessed via Mann–Whitney U test (Mann and Whitney, 1947). AUPRC Gain normalizes the area under the precision–recall curve by the positive class prevalence: AUPRC Gain = (AUPRC – *π*)/(1 – *π*), where *π* is the fraction of binding residues, yielding 0 at chance level and 1 for perfect discrimination; its significance is assessed via the same Mann–Whitney U test as AUROC. Fold-enrichment measures the concentration of attention at binding sites relative to the sequence background, with its significance assessed via hypergeometric test on the top 20% highest-attention residues. Full per-protein statistics are reported in Section B.1, and a subset is visualized in **Fig. S2**. Three proteins spanning different AUROC values (E1C9K5, P84131, Q5Y812) were selected for visualization in **Fig. 2E**. Three-dimensional structure visualizations with projected attention scores were rendered using UCSF ChimeraX (Pettersen et al., 2021).

#### 4.5.2. ESM2 Embedding Norm Control

To test whether the binding-site discrimination observed in GO-GPT’s cross-attention is already present in the frozen ESM2 representations, we computed L2 norms of ESM2 output embeddings at each residue position and evaluated them as a baseline predictor of DNA-binding residues using the same metrics (AUROC, AUPRC Gain, fold-enrichment) and statistical tests applied to attention scores. If ESM2 embeddings encode binding-site information in their magnitude—for example, through higher norms at structurally important positions— then GO-GPT’s attention could simply track this pre-existing signal rather than learning a GO-function-specific one. Per-protein comparisons between attention-based and L2-norm-based metrics are reported in **Fig. S2**.

#### 4.5.3. GO-GPT Embedding Analysis

We characterized the structure of GO-GPT’s learned representations through a suite of analyses spanning organism embeddings, GO term embeddings, and their relationship to external biological knowledge.

GO-GPT learns a unique embedding vector for each organism in the training vocabulary. We computed pairwise cosine distances between all organism embeddings and constructed a hierarchical clustering dendrogram using single linkage to visualize the resulting structure (**Fig. 2C**). To quantify the correspondence between embedding geometry and evolutionary relationships, we retrieved pairwise divergence times (in Mya) for the 30 most frequent organisms from the TimeTree 5 API (Kumar et al., 2022) and performed a Mantel test (Mantel, 1967) comparing the cosine distance matrix to the divergence time matrix (Spearman correlation, 9,999 permutations).

Each GO term in the vocabulary has a learned embedding. We assessed aspect-level organization in the original embedding space using cosine purity: for each GO term, we computed the fraction of its *κ* nearest neighbors (*κ* = 1, 3, 5, 10, 20) in cosine space that share the same GO aspect, averaged across all terms within each aspect. The per-aspect random baseline equals the aspect’s proportion of the full vocabulary (MF: 14.9%, BP: 75.0%, CC: 10.1%). Statistical significance was assessed via 1,000-permutation test (aspect label shuffles). To characterize the baseline distribution of embedding similarities, we computed pairwise cosine similarities between all GO terms within each aspect (**Fig. S6**). To visualize embedding structure, we projected GO term embeddings to two dimensions using UMAP (McInnes et al., 2020) with cosine distance, *n*_neighbors_ = 15, and min_dist = 0.1 (**Fig. 2H**; **Fig. S8**).

To test whether embedding geometry reflects ontology-based semantic relationships, we computed Resnik similarity (Resnik, 1995) for all within-aspect GO term pairs. Resnik similarity is defined as Resnik (*t*_*i*_, *t*_*j*_) = IC(MICA), where IC (*t*) = − log_2_ *P*(*t*) and *P*(*t*) is the fraction of training proteins annotated with term *t* or any of its descendants. IC-based measures naturally account for the uneven depth and annotation density across branches of the ontology, and have been shown to correlate with protein sequence similarity better than graph-distance-based alternatives (Lord et al., 2003). Crucially, IC is derived from the same annotation frequencies that the model observes during training, making it the appropriate ground truth for assessing whether the model has internalized the statistical structure of protein–function associations. We measured the Spearman correlation between Resnik similarity and cosine embedding similarity per aspect, with significance assessed via Mantel permutation test (1,000 permutations of term-to-embedding assignments; **Fig. 2F**).

To assess whether embeddings capture functional relationships that span ontology aspects, we computed Normalized Pointwise Mutual Information (NPMI; ranges from 1 for terms that never co-occur, through 0 for statistically independent terms, to 1 for perfectly co-occurring terms) for all cross-aspect term pairs (MF×BP, MF×CC, BP×CC) from protein co-annotation frequencies, restricting to terms with annotation frequency ≥ 20 and pairs with co-occurrence ≥ 5. Only proteins annotated in both relevant aspects contributed to each comparison. Significance was assessed by permuting the term-to-embedding assignment independently per aspect (1,000 permutations), recomputing cosine similarities for all valid pairs, and comparing the resulting Spearman *ρ* against the observed value (**Fig. 2G**). To evaluate whether this correlation enables practical retrieval, we formulated a cross-aspect retrieval benchmark: for each GO term in one aspect, we ranked all terms in the partner aspect by embedding cosine similarity and measured retrieval performance against ground-truth partners defined by NPMI > 0.5 (strong co-annotation). Only candidate pairs with co-occurrence ≥ 5 were evaluated, ensuring that positive and negative labels are supported by sufficient annotation evidence. We report AUROC, Mean Reciprocal Rank (MRR), and Precision@*κ* (*κ* = 1, 5, 10, 20) macro-averaged across query terms, with per-query random baselines computed from positive class prevalence. Both retrieval directions (A→B and B→A) are evaluated separately and combined into a symmetric AUROC. Statistical significance was assessed by the same embedding-permutation procedure (1,000 permutations; **Fig. S7**). To visualize representative cross-aspect associations in the embedding landscape, we selected pairs ranked by the product of NPMI and cosine similarity (top 200 per aspect combination) and filtered by UMAP Euclidean distance < 0.3, retaining 18 pairs that co-localize in the two-dimensional projection (**Fig. S8**; Table S2).

We evaluated whether embedding neighborhoods reflect the GO directed acyclic graph by computing *κ*-nearest-neighbor overlap with ontological neighbors (parents, children, and siblings within 2 hops via is_a and part_of relationships) for the top 1,000 most frequent terms per aspect. We report Precision@*κ* and Recall@*κ* at *κ* = 1, 3, 5, 10, 20, compared to a random baseline derived from vocabulary size and mean neighbor count, with significance assessed via 1,000-permutation test (**Fig. S6**).

#### 4.5.4. BioReason-Pro Embedding Space Analysis

To characterize how BioReason-Pro reorganizes protein representations relative to its input encoder, we extracted mean-pooled protein embeddings from the BioReason-Pro SFT model at two stages of the pipeline, following the approach of Hie et al. (2021) for visualizing learned protein representations. ESM3 pre-projection embeddings (1536-dimensional) capture the frozen protein language model representations before they enter the LLM (Hayes et al., 2024), while LLM Layer 35 post-RMSNorm embeddings (2560-dimensional) capture the final internal representations learned by the reasoning backbone.

For the full training set visualization (**Fig. 1H**), LLM Layer 35 embeddings were reduced from 2560 to 50 dimensions using PCA (Pearson, 1901) and then projected to two dimensions using UMAP (McInnes et al., 2020) with *n*_neighbors_ = 30 and min_dist = 0.3. Clusters were identified by applying HDBSCAN (Campello et al., 2013; McInnes et al., 2017) to the PCA-reduced LLM Layer 35 embeddings with min_cluster_size = 20, min_samples = 5, and Euclidean distance. The top 9 clusters by size were retained for annotation.

For the comparative visualization (**Fig. 1I**), a 10K protein subset was selected and both ESM3 and LLM Layer 35 embeddings were independently projected to two dimensions using UMAP with the same hyperparameters. Cluster assignments were fixed from the LLM Layer 35 clustering and applied identically to both projections so that the same proteins receive the same colors across panels. To functionally annotate each cluster, we computed GO Molecular Function term enrichment using one-sided Fisher’s exact test (Fisher, 2018) comparing term frequency within each cluster against all proteins as background. All MF terms at GO depth ≥ 1 were included, with a minimum of 5 occurrences per term per cluster. Each cluster is annotated by its most enriched MF term.

#### 4.5.5. BioReason-Pro Attention Analysis

To investigate whether BioReason-Pro grounds its reasoning in structurally meaningful protein features, we analyzed attention from specific tokens in the generated reasoning trace back to the protein embeddings.

For each analysis, a probe phrase is manually identified in the generated reasoning trace at the point where the model articulates a functionally relevant prediction. For example, in the eEFSec analysis the probe phrase is “with” preceding the predicted partner name “SECIS-binding protein 2”, and in the CFAP61 analysis it is “scaffold” at the point where the model infers non-enzymatic function (**Fig. 5A**, **Fig. 5E**).

Attention weights are extracted from all layers and all heads of the Qwen3-4B backbone. The best layer is selected automatically as the layer whose single best head achieves the highest mean attention to protein positions. At the selected layer, heads are ranked by mean attention to protein positions. Different attention heads capture different structural and functional features, so the head used for each analysis is selected based on specificity to the protein region of interest. For the eEFSec analysis the top-ranked head was used, while for the CFAP61 analysis the second-ranked head was selected. Per-residue attention scores are computed as the mean attention across all query token positions within the probe phrase to each protein residue position. For structural visualization, scores are min-max normalized to [0, 1].

Residue-to-domain mappings are obtained directly from the InterPro annotations. Only domain-type entries are retained, and start/end residue positions define each domain boundary. Residues not covered by any annotated domain are labeled as non-domain. This enables domain-level aggregation of per-residue attention scores for the plots in **Fig. 5B** and **Fig. 5F**.

Discrimination between functionally relevant and background residues is assessed using one-sided Mann–Whitney U tests (Mann and Whitney, 1947), testing whether attention at relevant residues exceeds that at non-relevant residues. Fold enrichment is computed as the ratio of mean attention at relevant residues to mean attention at non-relevant residues. Relevant residue sets are defined by structural contact analysis as described in Section 4.5.6.

#### 4.5.6. BioReason-Pro Structural Contact Analysis

To assess whether BioReason-Pro attention concentrates on experimentally validated functional interfaces, we computed structural contact maps from published cryo-EM structures and compared them against per-residue attention scores extracted as described in Section 4.5.5.

Contact residues were identified from mmCIF coordinate files (Westbrook et al., 2022) using KD-tree distance queries over all heavy atoms (Bentley, 1975). A protein residue was classified as contacting a partner molecule if any of its heavy atoms fell within 5.0 Å of any heavy atom in the partner (Méndez et al., 2003). Statistical significance was assessed using one-sided Mann–Whitney U tests (Mann and Whitney, 1947) comparing attention scores at contact versus non-contact residues. Fold enrichment was computed as the ratio of mean attention at contact residues to mean attention at non-contact residues.

##### eEFSec (P57772)

SECIS RNA contact residues were identified from the cryo-EM selenosome structure (PDB 7ZJW, 2.8 Å resolution) (Hilal et al., 2022), with Chain E corresponding to eEFSec and Chain S to the SECIS RNA element. All eEFSec residues with any heavy atom within 5.0 Å of any SECIS RNA atom were classified as contact residues. Per-residue attention scores from the probe token analysis (Section 4.5.5) were compared between contact and non-contact residues. Attention scores were projected onto the structure using a white-to-red colormap for visualization in PyMOL (Schrödinger, LLC, 2015).

##### CFAP61 (Q8NHU2)

The CFAP61 Rossmann-like domain (residues 665–998) was aligned to glutathione reductase (PDB 3GRS) (Karplus and Schulz, 1987) using topology-guided manual mapping. The canonical Rossmann *β*-*α*-*β*-*α*-*β* fold (Rao and Rossmann, 1973) was used to identify equivalent positions between 3GRS and the CFAP61 AlphaFold predicted structure (Fleming et al., 2025), matching *β*-strand/*α*-helix junctions, specifically anchoring on the GxGxxG loop at the *β*1-*α*1 junction (Wierenga et al., 1986). This alignment identified 12 positions in CFAP61 topologically equivalent to catalytic and cofactor-binding residues in 3GRS (Table 2). All 12 positions show amino acid substitutions inconsistent with catalytic function.

**Table 2.** Catalytic and cofactor-binding positions mapped from glutathione reductase (PDB 3GRS) to the CFAP61 Rossmann domain by topology-guided alignment. Repurposed residues (marked with †) are positions where the degenerate catalytic site directly contacts Dynein Heavy Chain 1 in the cryo-EM axoneme structure (PDB 8J07) (Walton et al., 2023).

| Functional role | 3GRS reference | CFAP61 position | Substitution |
| --- | --- | --- | --- |
| GxGxxG loop | G27–G32 | 671–676 | VGASSV |
| Redox cysteine | C63 | C734<sup>†</sup> | Conserved (interface) |
| Cofactor-binding | S | V737<sup>†</sup> | Ser → Val |
| Catalytic nucleophile | C58 | V739<sup>†</sup> | Cys → Val |
| FAD stacking | E50 | L767 | Glu → Leu |

Dynein Heavy Chain 1 contact residues were identified from the cryo-EM axoneme structure (PDB 8J07) (Walton et al., 2023), with Chain d0 corresponding to CFAP61 and Chain g6 to Dynein Heavy Chain 1. All CFAP61 residues within 5.0 Å of any Dynein Heavy Chain 1 heavy atom were classified as interface residues. Three positions (C734, V737, V739) were identified as repurposed residues by intersection of the catalytic position set from the 3GRS alignment with the Dynein Heavy Chain 1 interface set from the 8J07 structure.

For the attention enrichment analysis, Rossmann domain residues were partitioned into four groups: repurposed active-site residues (catalytic interface), remaining catalytic-site residues (catalytic only), Dynein Heavy Chain 1 interface residues (interface only), and other Rossmann-domain residues. Per-residue attention scores were compared across these groups using Mann–Whitney U tests (Mann and Whitney, 1947). Attention scores were projected onto the Rossmann domain for visualization in PyMOL (Schrödinger, LLC, 2015).

### 4.6. Experimental validation of the RCDG1–RAB5A interaction

#### 4.6.1. Wet-lab candidate selection

We selected the validation candidate through a staged filter. We restricted the pool to human proteins lacking a functional description in the latest UniProt release, and drew predictions from the BioReason-Pro SFT model, which proposes more hypotheses per protein than the RL model (Section 2.4). GPT-5 (Singh et al., 2025) flagged generations containing a specific, plausible, previously unreported hypothesis such as a named interaction partner, and a domain expert prioritized these by experimental tractability, favoring directly testable predictions such as a co-immunoprecipitation-assayable interaction, and biological value, favoring clinically relevant proteins such as cancer biomarkers. RCDG1, a renal cell carcinoma biomarker with no assigned function and a specific de novo prediction of RAB5A binding, met these criteria.

#### 4.6.2. RCDG1 co-immunoprecipitation

To test the predicted interaction between RCDG1 and RAB5A, we generated a HEK293T line stably expressing Flag-tagged RCDG1 or GFP, captured protein complexes under reversible crosslinking to preserve transient interactions, and assayed co-immunoprecipitation by immunoblot. The full procedure is detailed below.

##### Plasmid construction and lentiviral production

A Twist gBlock fragment for Flag-RCDG1 or Flag-GFP was cloned into pLV-EF1a-IRES-Puro (Addgene #85132) (Hayer et al., 2016) by Gibson assembly (Gibson et al., 2009). For viral production, the cloned plasmid was co-transfected with the plasmids pMD.2G and pCMV-dR8.91 (Naldini et al., 1996; Zufferey et al., 1997) using TransIT-Lenti (Mirus) into HEK293T cells following the manufacturer’s manual. Viral supernatant was collected 48 h after transfection and used to transduce HEK293T cells, which were selected with puromycin (2 *μ*g/mL) to generate a stably expressing line.

##### Crosslinking and lysis

To preserve transient interactions, cells were crosslinked with 1 mM of the reversible crosslinker DSP (Thermo Fisher) (Lomant and Fairbanks, 1976) at room temperature for 30 min, and quenched with 20 mM Tris, pH 7.5, for 15 min at room temperature. The cells were then lysed in Pierce IP lysis/wash buffer supplemented with 1× Halt protease inhibitor (Thermo Fisher).

##### Co-immunoprecipitation

For each immunoprecipitation reaction, 6 *μ*g of anti-Flag (Proteintech #20543-1-AP) or rabbit IgG (Proteintech #30000-0-AP) antibody was incubated with 1 mg of protein lysate at room temperature for 2 h with rotation. Subsequently, 25 *μ*L of washed Pierce Protein A/G beads was added to each immunoprecipitant reaction and incubated at room temperature for a further 1 h with rotation. The beads were then washed three times with Pierce IP lysis/wash buffer and once with DPBS. For elution, the beads were resuspended in 2× NuPAGE LDS sample buffer and incubated at 95 ^◦^C for 10 min together with the input. To reverse-crosslink, 62.5 mM DTT was added to each eluate, followed by incubation at 37 ^◦^C for 30 min and at 95 ^◦^C for 5 min.

##### Immunoblotting

For the western blot, the following antibodies were used for probing: anti-Flag M2 antibody (Merck #F3165), anti-RAB5A antibody, and anti-mouse IgG (H+L) antibody conjugated with DyLight 680 (Cell Signaling Technology #46449 and #5470, respectively). The blot was imaged with a Bio-Rad ChemiDoc MP imaging system.

## Data Availability

All datasets used to train and evaluate models are available at https://bioreason.net/data.

## Code and Model Availability

We make code and tools available at the following links:

- Web-based inference server: https://bioreason.net
- Training and evaluation code repository: https://bioreason.net/code
- BioReason-Pro function predictions for over 240,000 proteins, including the Human Protein Atlas: https://bioreason.net/atlas

We make the model parameters available on Hugging Face:

- GO-GPT: https://bioreason.net/gogpt
- BioReason-Pro SFT: https://bioreason.net/sft
- BioReason-Pro RL: https://bioreason.net/rl

## Acknowledgements and Disclosure of Funding

We thank Bryan Perozzi and Michael Galkin for insights on encoding Gene Ontology graph structure and learning hierarchical ontologies. We thank Jure Leskovec for insightful discussions on multimodal biological reasoning. We thank Guillaume Filion for advising and insights on protein function reasoning and training suggestions. We thank Shayan Pardis for valuable input on GO term representation and graph learning approaches. We thank Li Erran Li for insights on biological reasoning. We thank Anshul Kundaje for feedback on evaluation methodology and suggestions for assessing biological reasoning. We thank Iddo Friedberg for guidance on CAFA evaluation. We thank Andrew Magnuson and Ronald Xie for their valuable insights on model architecture. We thank Jacob Junqi Tian and John Willes for helpful discussions on training infrastructure and reinforcement learning strategies. We thank Amol Punjabi for discussions on the impact and release strategy of this work. We thank Kuan Pang for fruitful discussions about future work. We thank Joseph Caputo for media and communications support. We thank Clem Delangue for his support and encouragement in making our models publicly accessible through the Hugging Face platform. Compute resources were provided by the Arc Institute, Vector Institute, the Digital Research Alliance of Canada, the Government of Canada, and a compute grant from Modal Labs. H.G. is an Arc Core Investigator and acknowledges funding support from the Arc Institute.

## Author Contributions

A.F., A.S-A., and P.I. conceived the project and co-led its development. H.G. and B.W. supervised the project. D.P.B. and P.D.H. advised throughout the project. A.F. and P.I. designed the BioReason-Pro architecture. A.S-A. designed the GO-GPT architecture. K.Z. designed the GO graph encoder. A.F. developed the SFT training pipeline and ran all SFT experiments. P.I. developed the RL training pipeline and ran all RL experiments. A.S-A. developed the GO-GPT training code and ran all GO-GPT experiments. P.I. developed the vLLM-based fast inference stack. T.G. contributed to the GO-GPT inference code. A.Shah contributed to high-throughput inference for RL. A.A contributed to training and scaling BioReason-Pro. O.M. contributed and provided guidance on RL. H.D. and C.J.M. advised RL training. P.G., O.I., A.S-A, S.M., and J.N. curated and processed the training dataset, including protein sequences, domain annotations, interaction data, and GO terms. A.F. and O.I. generated the synthetic reasoning data. P.I. advised the synthetic data generation. O.I. and P.G. constructed the test dataset. A.F. developed the LLM judge and human expert evaluation frameworks and conducted all BioReason-Pro evaluations. P.I. advised the LLM judge and human expert evaluations. A.S-A. set up GO term evaluation pipelines and conducted all GO-GPT evaluations. B.M.H.C. conducted all wet lab experiments and validations. A.F., H.G., and B.W. recruited human expert evaluators. P.G. and T.G. conducted GO-based baseline evaluations. K.Z., J.N., and A.F. conducted free-text baseline evaluations. A.F. and P.I. conducted the BioReason-Pro interpretability analysis. A.S-A. conducted the GO-GPT interpretability analysis. A.F. wrote up the eEFSec and CFAP61 cases. O.I. contributed to case study selection. C.A.T., M.Y.Y.L., and A.F.M.S. conducted independent model evaluation. N.L., H.Cui, A.A., A.J., A.Fa., A.C-P., J.S.S., F.N., A.A.N., H.C.M., and L.S. served as human evaluators. B.S.P. provided scientific advising and feedback on the manuscript. C.R-T. provided scientific advising and designed figures. M.d.C. and P.I. designed the web interface and inference server. A.F., A.S-A., and T.G. generated predictions for the released atlas. A.F., A.S-A., and P.I. managed the open-source release of models, code, and datasets. A.F. and A.S-A. created all plots. A.F., A.S-A., P.I., P.G., and O.I. wrote the first draft of the manuscript. H.G. and B.W. provided guidance throughout.

## A. Supplementary Figures

**Figure S1.**
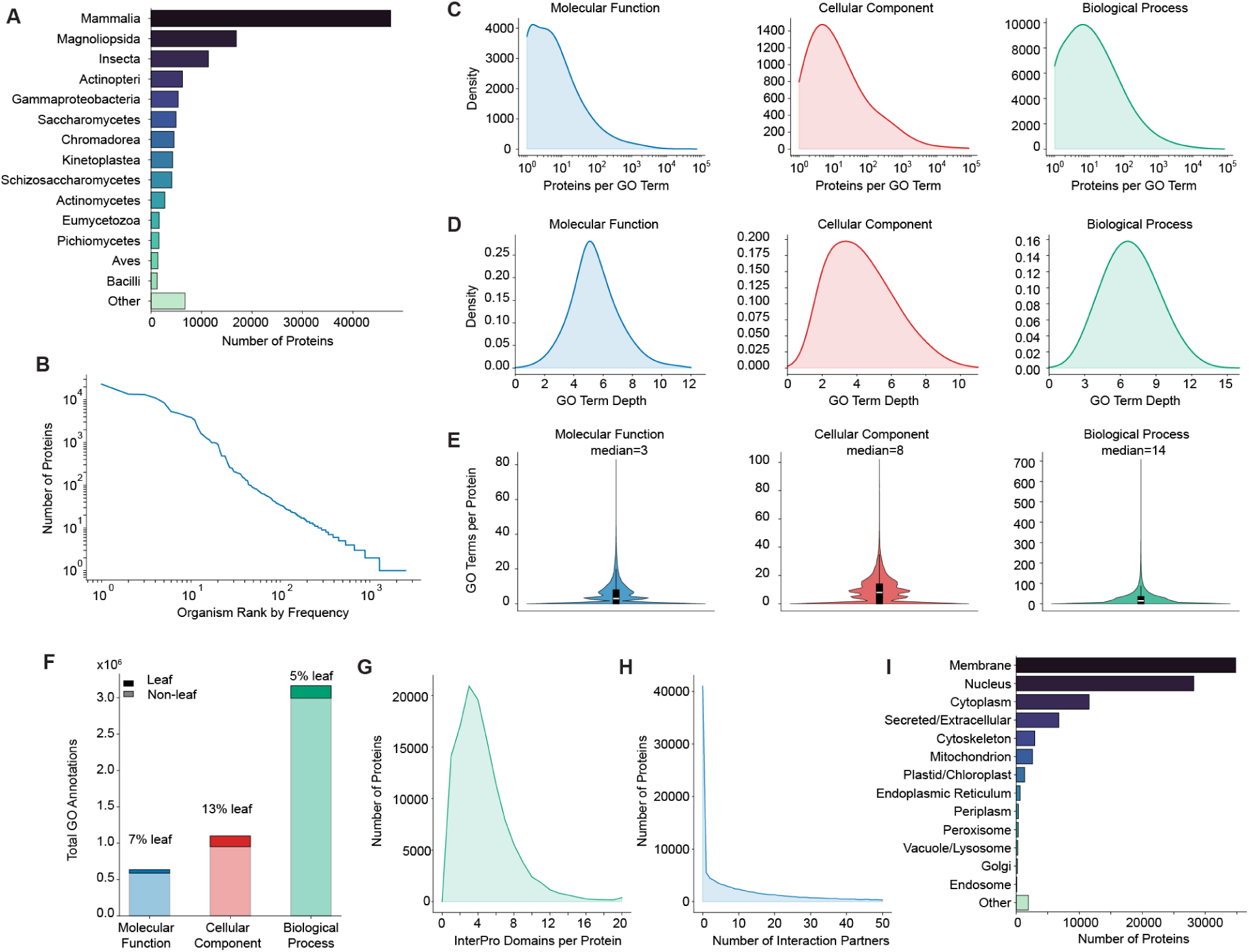
Training dataset statistics. **(A) Taxonomic distribution.** Proteins per taxonomic class showing strong mammalian bias with broad coverage across vertebrates, insects, plants, fungi, and bacteria. **(B) Organism frequency.** Rank-frequency plot on log-log scale revealing a long-tail distribution, with a small number of model organisms dominating protein counts. **(C) GO term frequency per aspect.** Kernel density estimates of proteins annotated per GO term on a log scale, showing most terms are rare with few appearing across many proteins. **(D) GO term depth per aspect.** Ontology depth distribution for unique GO terms in each aspect. **(E) GO terms per protein.** Violin plots of GO annotation counts per protein for each aspect, with median values of 3 (MF), 8 (CC), and 14 (BP). **(F) Leaf versus non-leaf annotations.** Proportion of annotations corresponding to leaf terms (no children in the ontology) versus non-leaf terms per aspect. Leaf terms constitute 7% of MF, 13% of CC, and 5% of BP annotations. **(G) InterPro domain coverage.** Distribution of InterPro domain annotations per protein, with most proteins having fewer than 10 annotated domains. **(H) Protein-protein interaction coverage.** Distribution of interaction partners per protein from STRING, showing a long-tail distribution. **(I) Subcellular localization.** Proteins per broad subcellular compartment, with membrane, nucleus, and cytoplasm as the dominant localizations.

**Figure S2.**
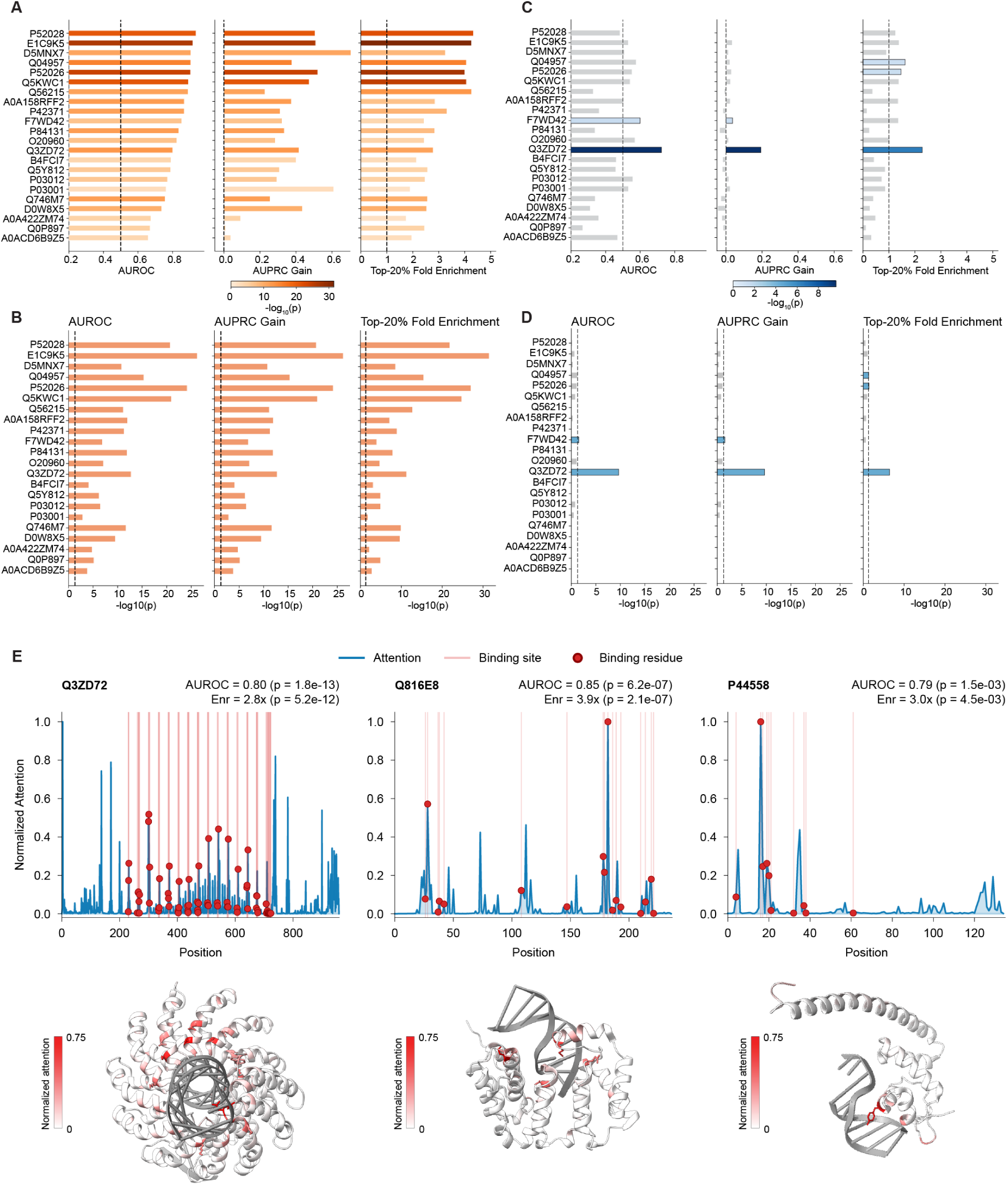
GO-GPT attention enrichment at DNA-binding residues. **(A)** AUROC, AUPRC Gain, and top-20% fold enrichment for a representative subset of the 63 non-training BioLiP proteins where GO-GPT predicted DNA binding, sorted by fold enrichment. Full per-protein statistics in Table S1. Bar color encodes − log_10_ ( *p*); gray bars are non-significant (*p* ≥ 0.05). Dashed lines: chance level. Across all 63 proteins, 59/63 are significant (Mann–Whitney U); 55/63 (hyper-geometric). **(B)** log_10_ *p* values for (A). AUROC and AUPRC Gain: Mann–Whitney U; fold enrichment: hypergeometric. Dashed line: *p* = 0.05. **(C, D)** ESM2 residue embedding L2 norm baseline (same metrics and protein ordering), testing whether binding-site discrimination is present in input representations before cross-attention (Section 4.5.2). **(E)** Per-residue attention for three additional proteins (Q3ZD72, Q816E8, P44558) with protein–DNA structures colored.

**Figure S3.**
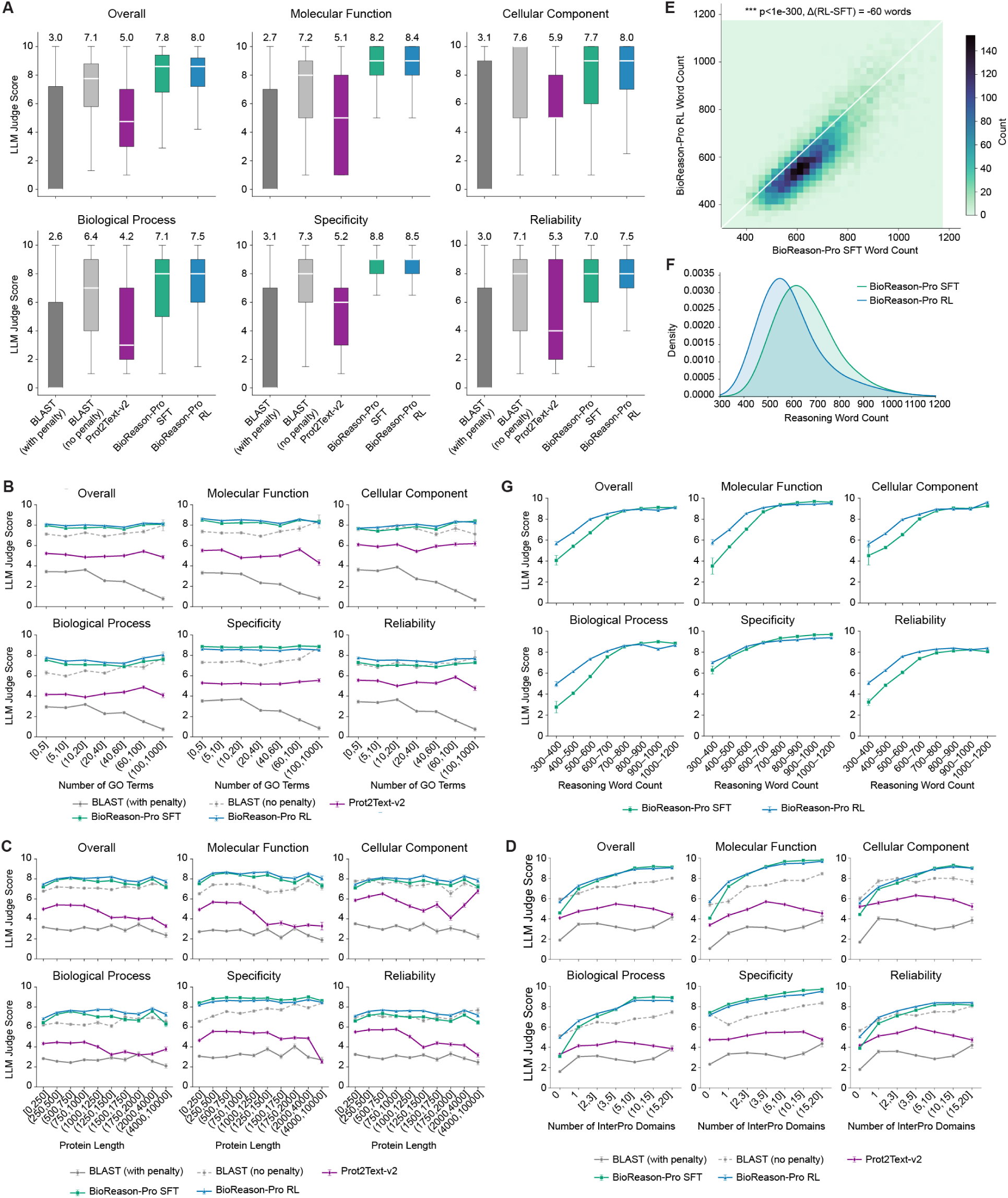
LLM-as-judge evaluation across models and protein properties. **(A) Score distributions per evaluation axis.** Box plots of LLM judge scores for BLAST (with or without penalty), Prot2Text-v2, BioReason-Pro SFT, and BioReason-Pro RL across all five evaluation axes. Mean values shown above each box. **(B) Performance by GO term count.** Mean scores binned by total number of GO annotations per protein. **(C) Performance by protein length.** Mean scores binned by protein sequence length. **(D) Performance by InterPro domain count.** Mean scores binned by number of InterPro domain annotations per protein. **(E) SFT versus RL trace length.** 2D density heatmap of per-protein reasoning word counts showing that SFT consistently produces longer reasoning traces than RL for the same proteins (mean Δ = −60 words, paired Wilcoxon *p* < 10^−300^). **(F) Trace length distributions.** Kernel density estimates of reasoning word counts for SFT and RL. **(G) Judge score by reasoning length.** Mean scores binned by word count for SFT and RL.

**Figure S4.**
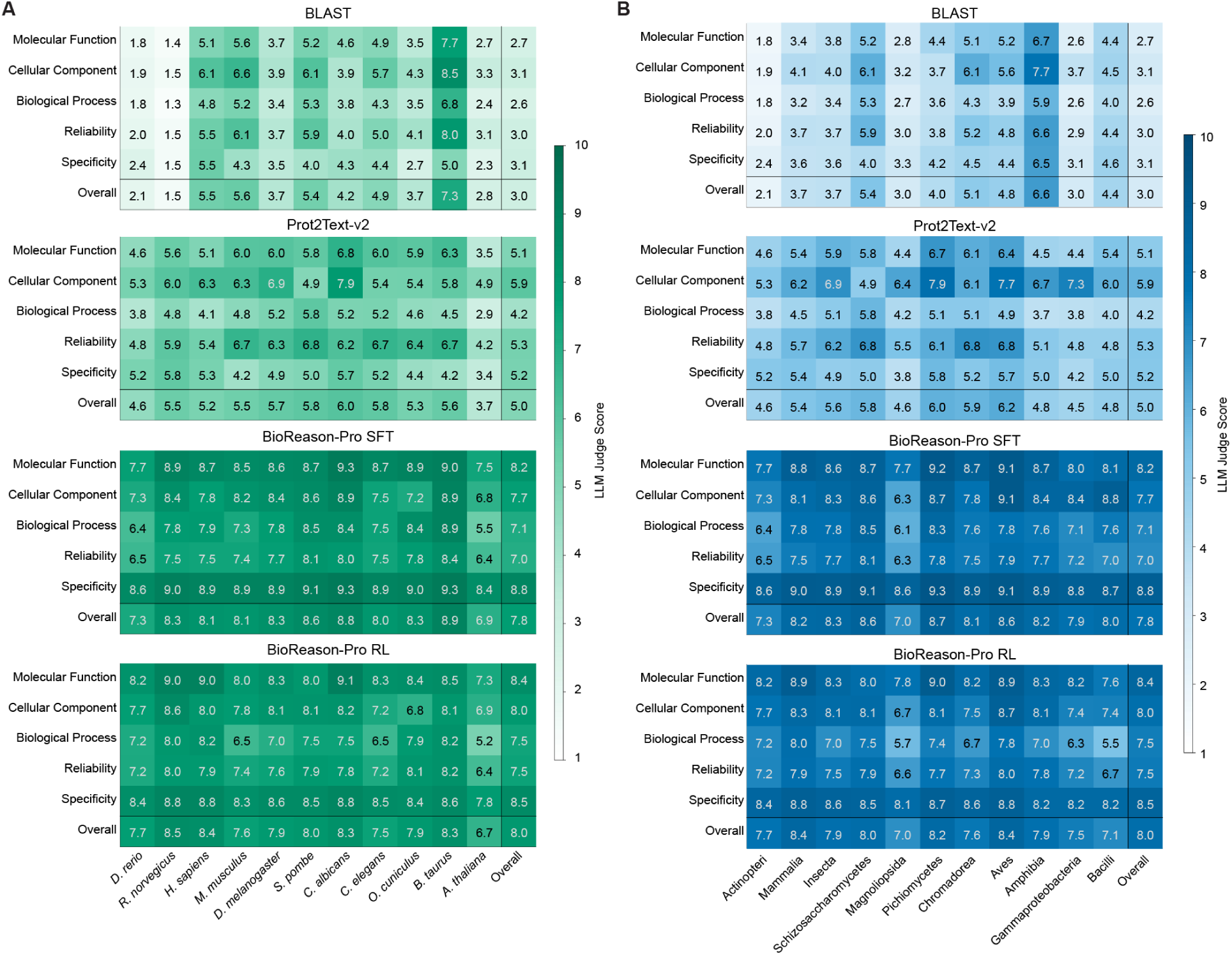
LLM-as-judge scores stratified by organism and taxonomic class. **(A) Performance by organism.** Heatmaps of mean scores for the ten most frequent organisms across four models (BLAST, Prot2Text-v2, BioReason-Pro SFT, BioReason-Pro RL), with rows representing five evaluation axes and Overall, and columns representing organisms sorted by frequency. BLAST performance varies sharply across organisms, with very low scores for some species (e.g., *D. rerio*, *R. norvegicus*) where BLAST-based transfer is least effective. Prot2Text-v2 is more uniform but remains in the 4–6 range. BioReason-Pro SFT and RL both maintain consistently high scores (7–9) across all organisms, with RL showing a modest advantage in Reliability. The advantage of BioReason-Pro over baselines is most pronounced for organisms where BLAST performs poorly, consistent with the generalization analysis in Section 4.4.5. **(B) Performance by taxonomic class.** Heatmaps of mean scores for the ten most frequent taxonomic classes, arranged analogously. BioReason-Pro generalizes broadly across taxonomic diversity, maintaining stable performance from vertebrates (Actinopteri, Mammalia) through invertebrates (Insecta, Chromadorea) and unicellular organisms (Schizosaccharomycetes, Gammaproteobacteria). BLAST scores degrade most for underrepresented classes, whereas BioReason-Pro shows minimal variation, indicating that the model learned transferable functional reasoning rather than organism-specific annotation patterns.

**Figure S5.**
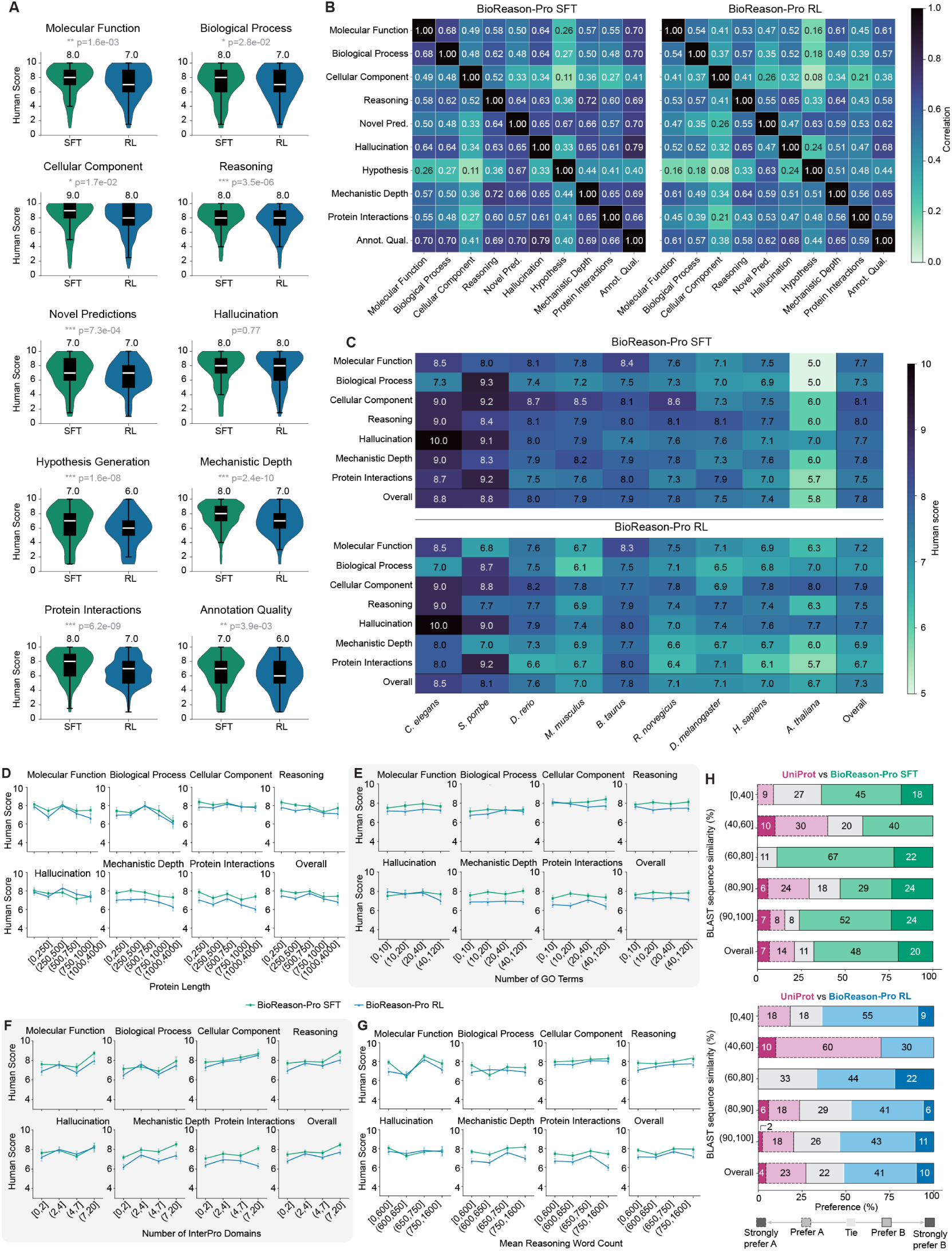
Human expert evaluation across all axes and protein properties. **(A) Per-axis score distributions.** Violin plots for BioReason-Pro SFT and RL across all ten evaluation axes with Mann–Whitney U significance tests. SFT scores significantly higher on most axes, with the largest differences on Mechanistic Depth and Hypothesis Generation. **(B) Inter-axis correlation matrices.** Pairwise Pearson correlations among all ten axes for SFT and RL. Reasoning, Mechanistic Depth, and Hallucination form a correlated cluster in both models. **(C) Scores by organism.** Heatmap of mean human scores for organisms with at least three evaluated proteins, showing stable performance across organisms for both models. **(D) Performance by protein length.** Mean human scores binned by sequence length for BioReason-Pro SFT and RL across all evaluation axes. **(E) Performance by GO term count.** Mean human scores binned by total GO annotations per protein. **(F) Performance by InterPro domain count.** Mean human scores binned by number of InterPro domain annotations per protein. **(G) Performance by reasoning trace length.** Mean human scores binned by average reasoning word count. **(H) Preference versus UniProt by sequence similarity.** Expert preference distributions for SFT (top) and RL (bottom) compared to UniProt ground truth, binned by BLAST sequence similarity.

**Figure S6.**
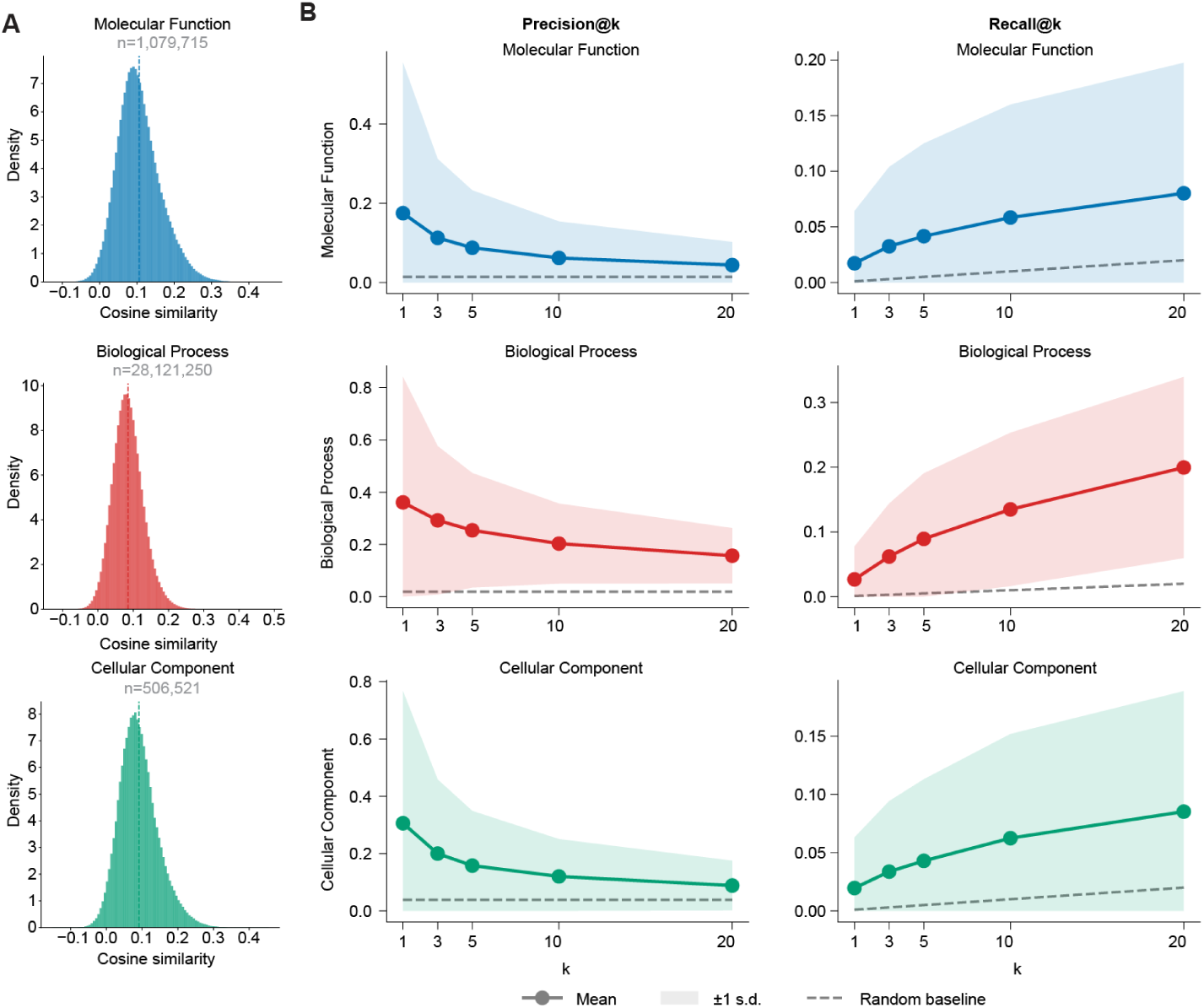
Within-aspect analysis of GO-GPT learned GO term embeddings. **(A) Cosine similarity distribution.** Per-aspect density of pairwise cosine similarities among all GO term embeddings, with total pair counts shown (MF: 1.08M, BP: 28.1M, CC: 507K). Dashed lines indicate per-aspect means. All three aspects show distributions centered at small positive values, indicating that embeddings within an aspect are weakly but consistently similar. **(B)** *κ***-nearest-neighbor Precision@***κ* **and Recall@***κ*. Overlap between *κ*-nearest embedding neighbors and ontological neighbors (parents, children, and siblings within 2 hops via is_a and part_of edges) for the top 1,000 terms per aspect. Precision@*κ* (left) decreases with *κ* as expected while remaining well above the random baseline (dashed lines), indicating that the closest embedding neighbors are the most ontologically relevant. Recall@*κ* (right) increases with *κ*, recovering a growing fraction of true ontological neighbors. Shaded bands show ±1 s.d. All values significant (*p* < 10^−3^, 1,000-permutation test).

**Figure S7.**
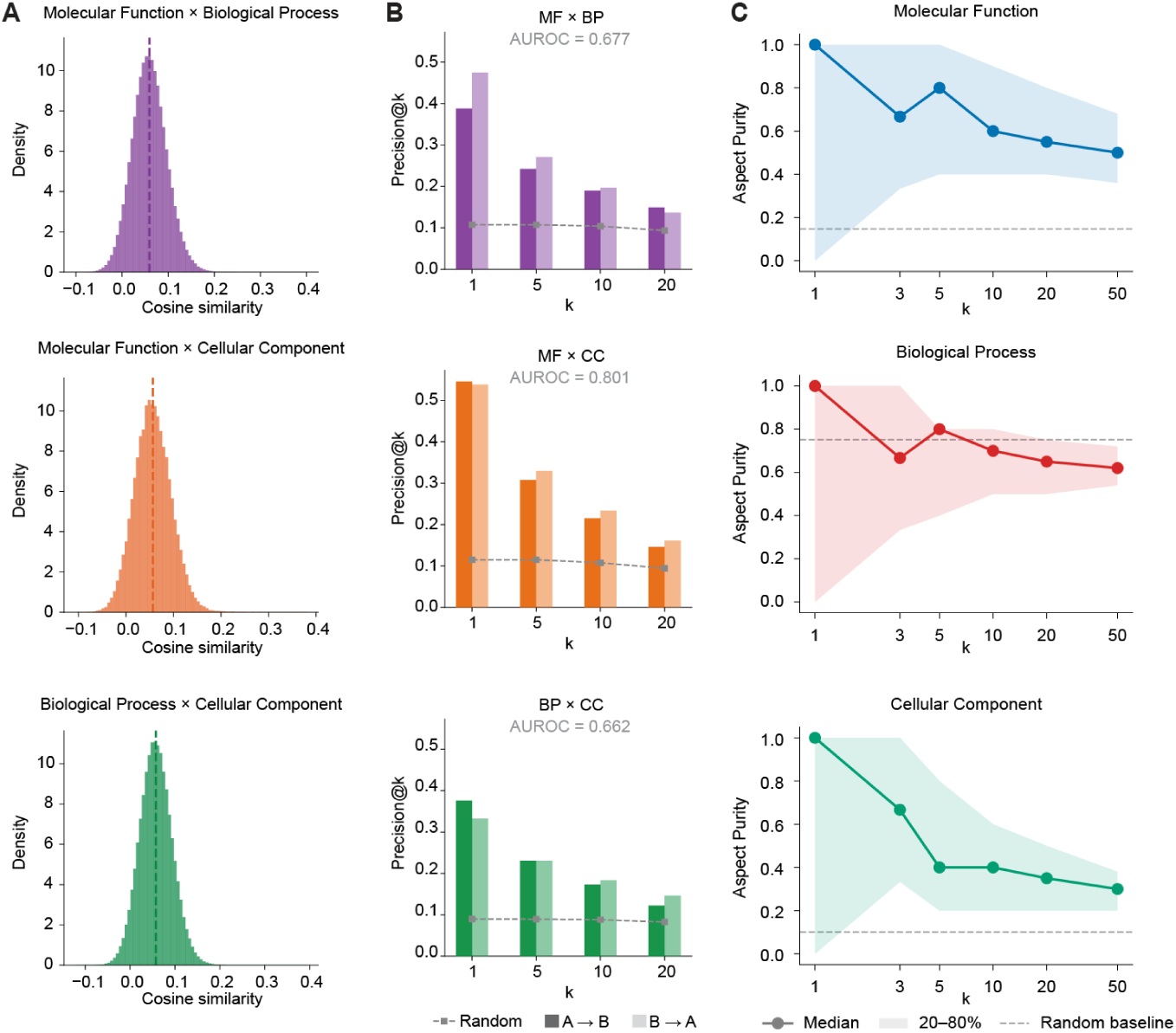
Cross-aspect analysis of GO-GPT learned GO term embeddings. (A) Cross-aspect cosine similarity distribution. Per-aspect-pair density of pairwise cosine similarities between GO terms belonging to different aspects (MF BP, MF CC, BP CC). NPMI co-annotation vs. cosine embedding similarity is shown in **Fig. 2G**. **(B) Cross-aspect retrieval benchmark.** For each GO term in one aspect, all terms in the partner aspect are ranked by embedding cosine similarity and evaluated against ground-truth partners defined by strong co-annotation (NPMI > 0.5). Grouped bars show Precision@*κ* for both retrieval directions (darker: A→B; lighter: B→A); square markers show random baseline. Text annotations report symmetric AUROC (mean of both directions) and permutation *p*-value (1,000 embedding shuffles). All three aspect pairs achieve AUROC well above the 0.5 chance level (*p* < 10^−3^), with MF×CC showing the strongest retrieval (AUROC = 0.80). **(C) Cosine purity.** Fraction of each term’s *κ* nearest neighbors in the GO embedding space that share the same GO aspect, shown as median per aspect. Shaded bands: 20th–80th percentile. Dashed lines: per-aspect random baseline (aspect proportion of the vocabulary). MF and CC are enriched 3–5×above baseline (*p* < 10^−3^, 1,000-permutation test); BP shows no enrichment, consistent with its broader functional diversity.

**Figure S8.**
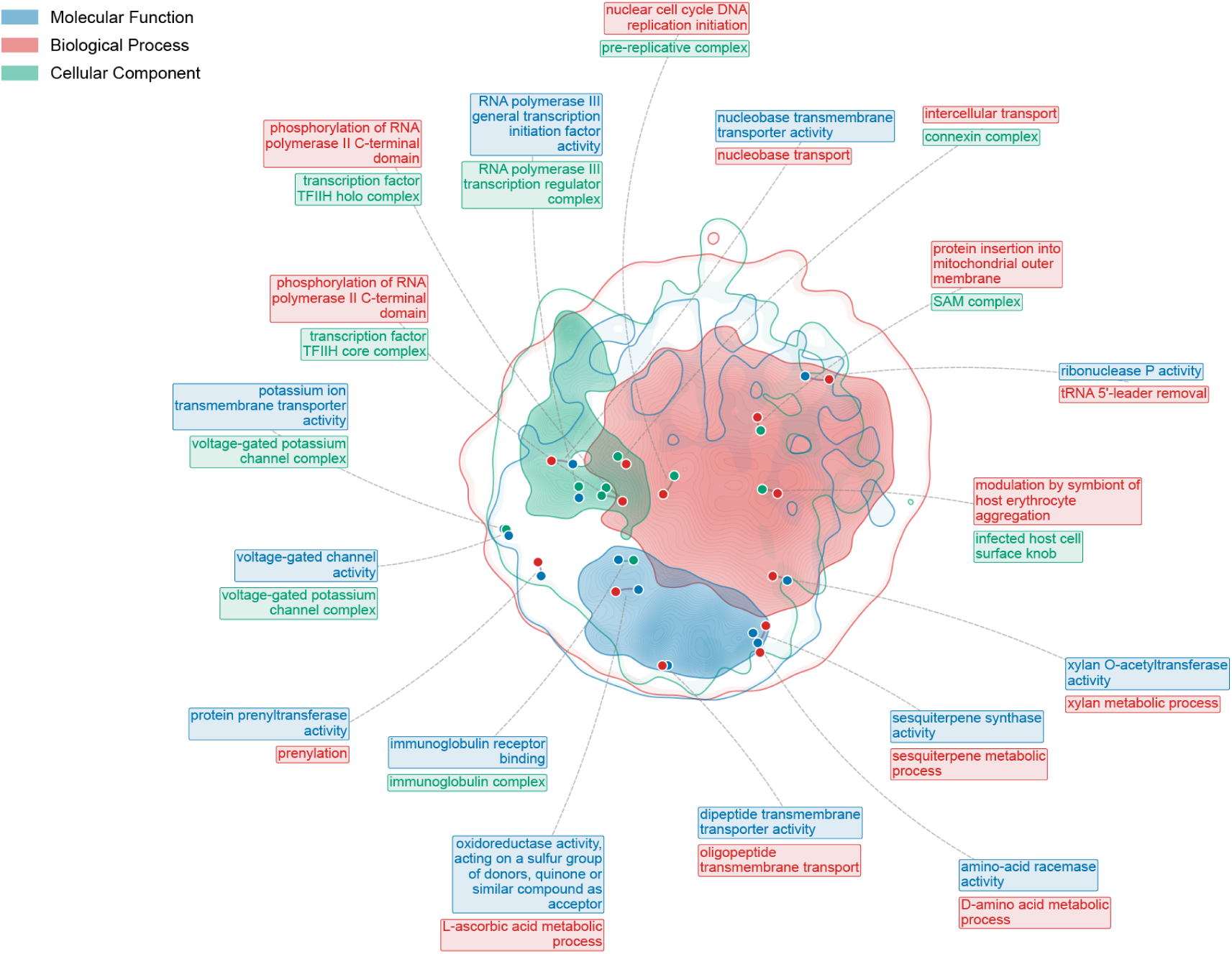
Cross-aspect GO term pairs visualized on the UMAP embedding landscape. Kernel density estimation (KDE) contours show the spatial distribution of GO term embeddings in the UMAP projection for each aspect: Molecular Function (MF, blue), Biological Process (BP, red), and Cellular Component (CC, green). An outermost contour band marks the 3% of maximum density threshold; 25 inner contour levels are drawn with increasing opacity (up to *α* = 0.5) to highlight regions of highest term density while minimizing visual clutter. Annotated pairs are cross-aspect GO term pairs selected through a reproducible pipeline: all cross-aspect pairs with co-occurrence ≥ 5 and term frequency 20 ( 1.2 million total) were ranked by the product of NPMI (co-annotation frequency) and cosine embedding similarity, then filtered by UMAP distance < 0.3 so that only pairs co-localized in the two-dimensional projection are annotated. Connecting arcs link paired terms; leader lines connect labels to pair midpoints. These 18 pairs (Table S2) are illustrative examples of the population-level correlation between NPMI and cosine similarity established across all cross-aspect pairs (Spearman *ρ* = 0.10–0.17, *p* < 10^−3^, embedding-permutation test; **Fig. 2G**), which also supports cross-aspect retrieval with AUROC = 0.66–0.80 (**Fig. S7B**).

**Figure S9.**
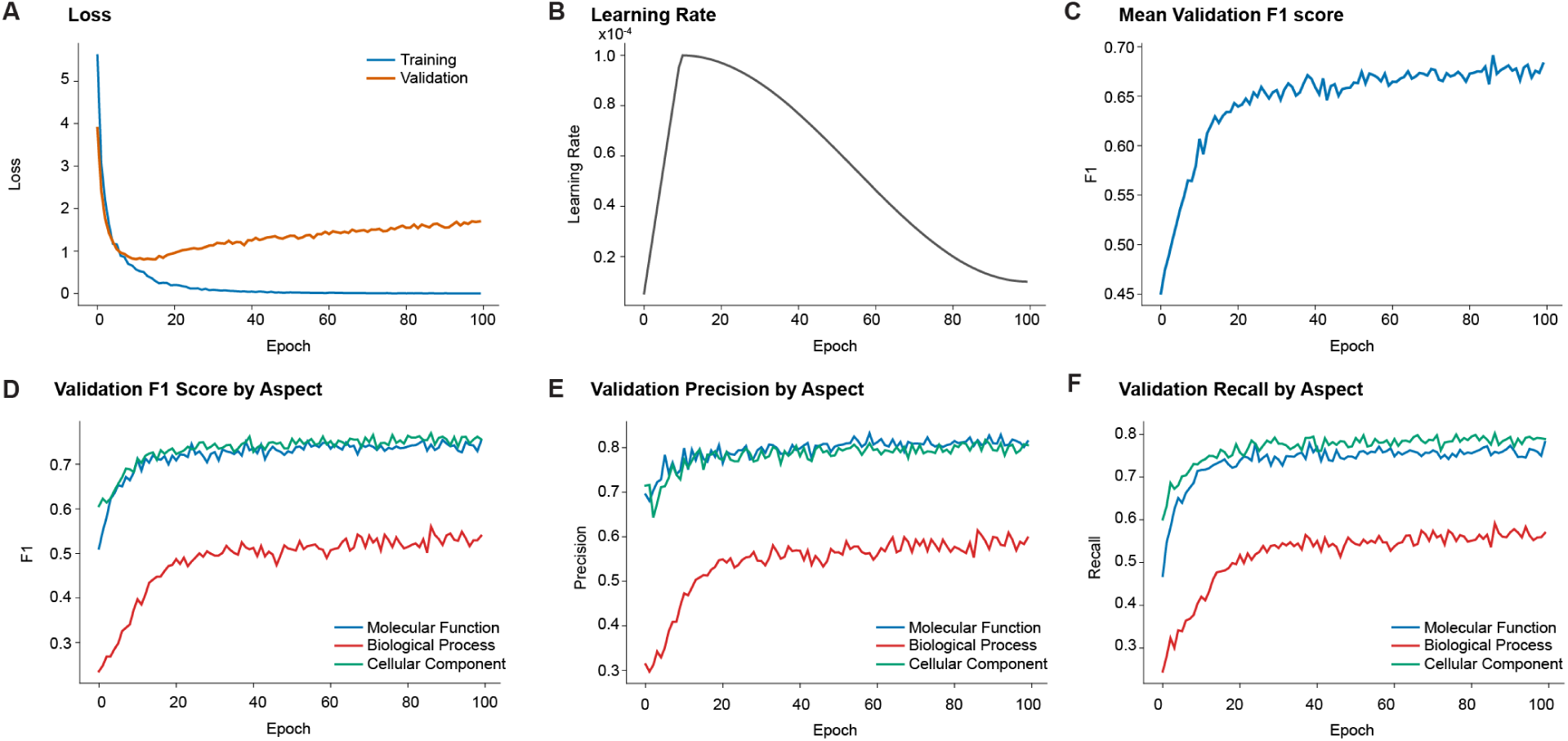
Training dynamics of the GO-GPT model. **(A)** Training and validation cross-entropy loss over 100 epochs. Training loss decreases steadily while validation loss plateaus and begins to rise after approximately 20 epochs, indicating overfitting in later training. **(B)** Learning rate schedule: linear warmup over the first ~10% of training followed by cosine decay to a minimum of ~0.1×the peak learning rate (1×10^−4^). **(C)** Mean, unweighted validation F1 score (macro-averaged across MF, BP, and CC aspects), plateauing near 0.68. **(D)** Per-aspect validation F1 scores. CC and MF converge rapidly and plateau near 0.73 and 0.71, respectively, while BP plateaus near 0.54, consistent with the larger and more heterogeneous BP vocabulary. **(E)** Per-aspect validation precision. MF and CC reach approximately 0.80, while BP stabilizes near 0.58. **(F)** Per-aspect validation recall. MF and CC reach approximately 0.78, while BP stabilizes near 0.58.

**Figure S10.**
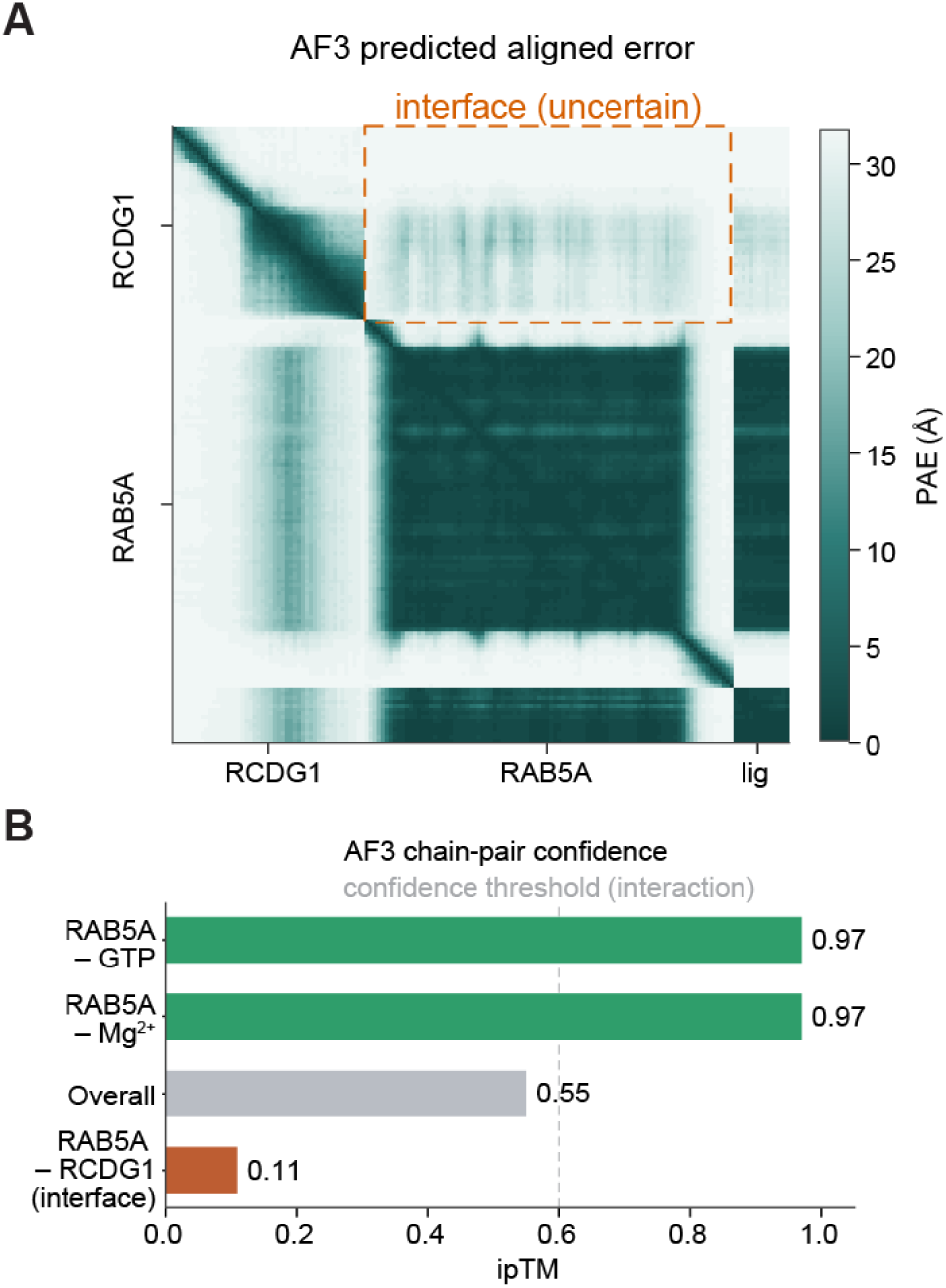
AlphaFold3 modeling of the RCDG1–RAB5A complex. **(A) Predicted aligned error.** AlphaFold3 PAE for RCDG1 modeled with RAB5A in its active, GTP- and Mg^2+^-bound state. Error is low within RAB5A but high across the RCDG1-to-RAB5A inter-chain block (orange dashed box), indicating that the interface is not confidently resolved. The ligand block (lig) denotes the bound GTP and Mg^2+^. **(B) Chain-pair confidence.** AlphaFold3 ipTM per chain pair. The nucleotide and ion are placed with high confidence (RAB5A–GTP and RAB5A–Mg^2+^ both 0.97), whereas the RCDG1–RAB5A interface remains unresolved (0.11). The overall value of 0.55 is inflated by the confidently placed cofactors. The dashed line marks the interaction confidence threshold.

## Appendix

### B.1. GO-GPT Attention Analysis at DNA-Binding Sites

Table S1 reports per-protein attention enrichment statistics for all 63 proteins in the DNA-binding attention analysis (Section 4.5.1). For each protein, AUROC quantifies the discrimination between attention scores at annotated binding versus non-binding residues (Mann–Whitney U test), AUPRC Gain normalizes AUPRC by positive class prevalence (Section 4.5.1), and fold-enrichment measures the ratio of mean attention at binding sites to non-binding sites. Hypergeometric *p*-values test whether the overlap between the top 20% highest-attention residues and annotated binding residues exceeds chance expectation.

**Table S1.**
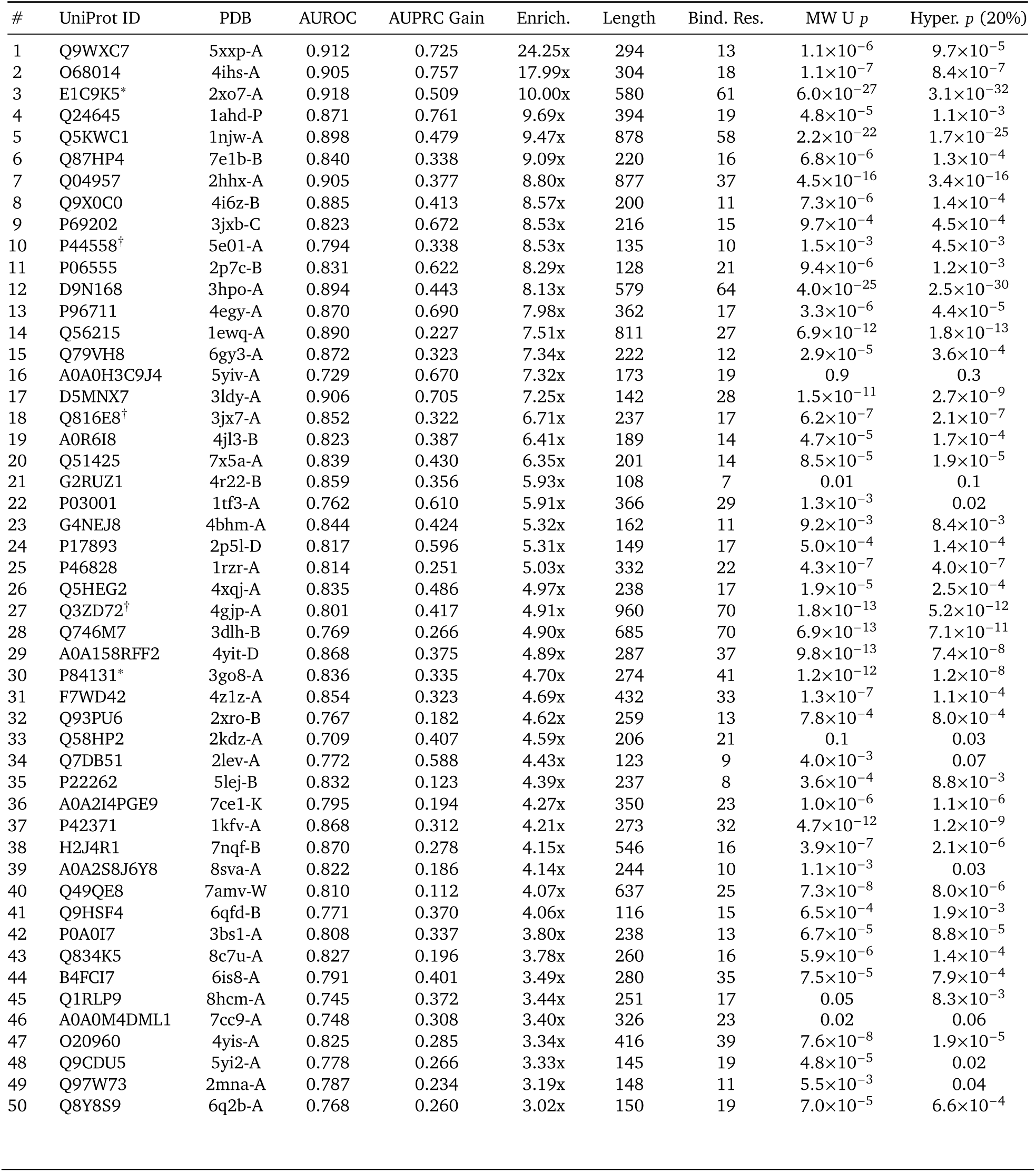
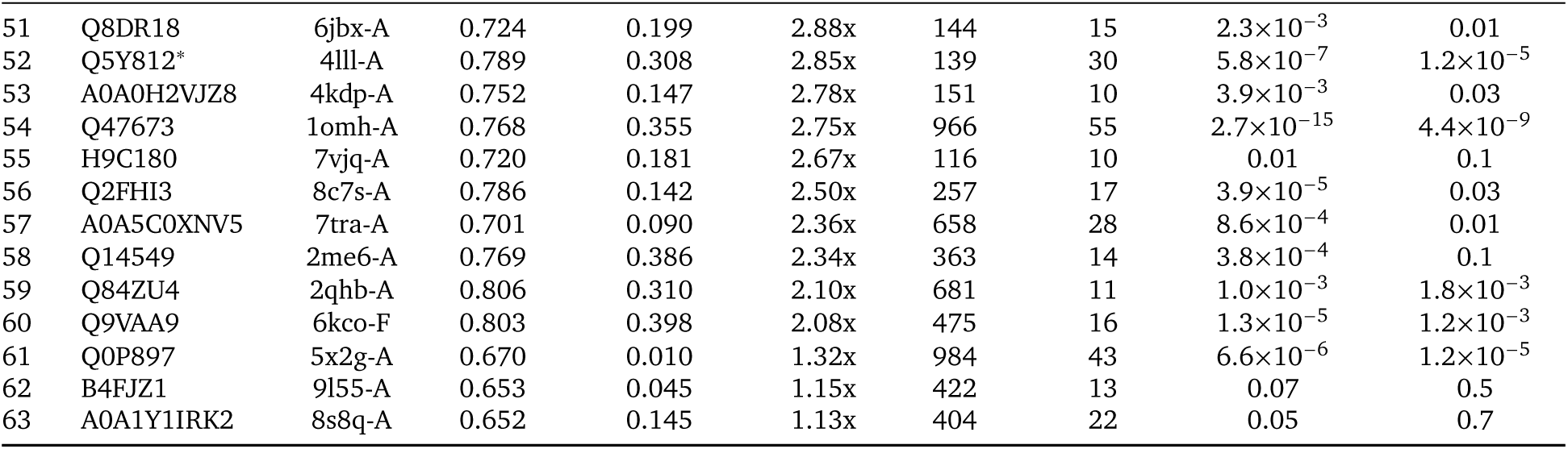
Attention enrichment at DNA-binding residues across all 63 evaluation proteins, ranked by fold-enrichment. ^∗^Proteins visualized in **Fig. 2E**; ^†^proteins visualized in **Fig. S2C**.

### B.2. Cross-Aspect GO Term Pairs on the UMAP Embedding Landscape

Table S2 lists the 18 cross-aspect GO term pairs annotated on the UMAP embedding landscape (**Fig. S8**). Pairs were selected through a reproducible pipeline: all cross-aspect pairs with co-occurrence ≥ 5 and term frequency ≥ 20 (>1.2 million total across three aspect combinations) were ranked by the product of NPMI and cosine embedding similarity (top 200 per combination), then filtered by UMAP Euclidean distance < 0.3 to retain only pairs that co-localize in the two-dimensional projection. These pairs are illustrative examples of the population-level correlation between co-annotation frequency and learned embedding similarity (Spearman *ρ* = 0.10–0.17, *p* < 10^−3^; **Fig. 2G**).

**Table S2.**
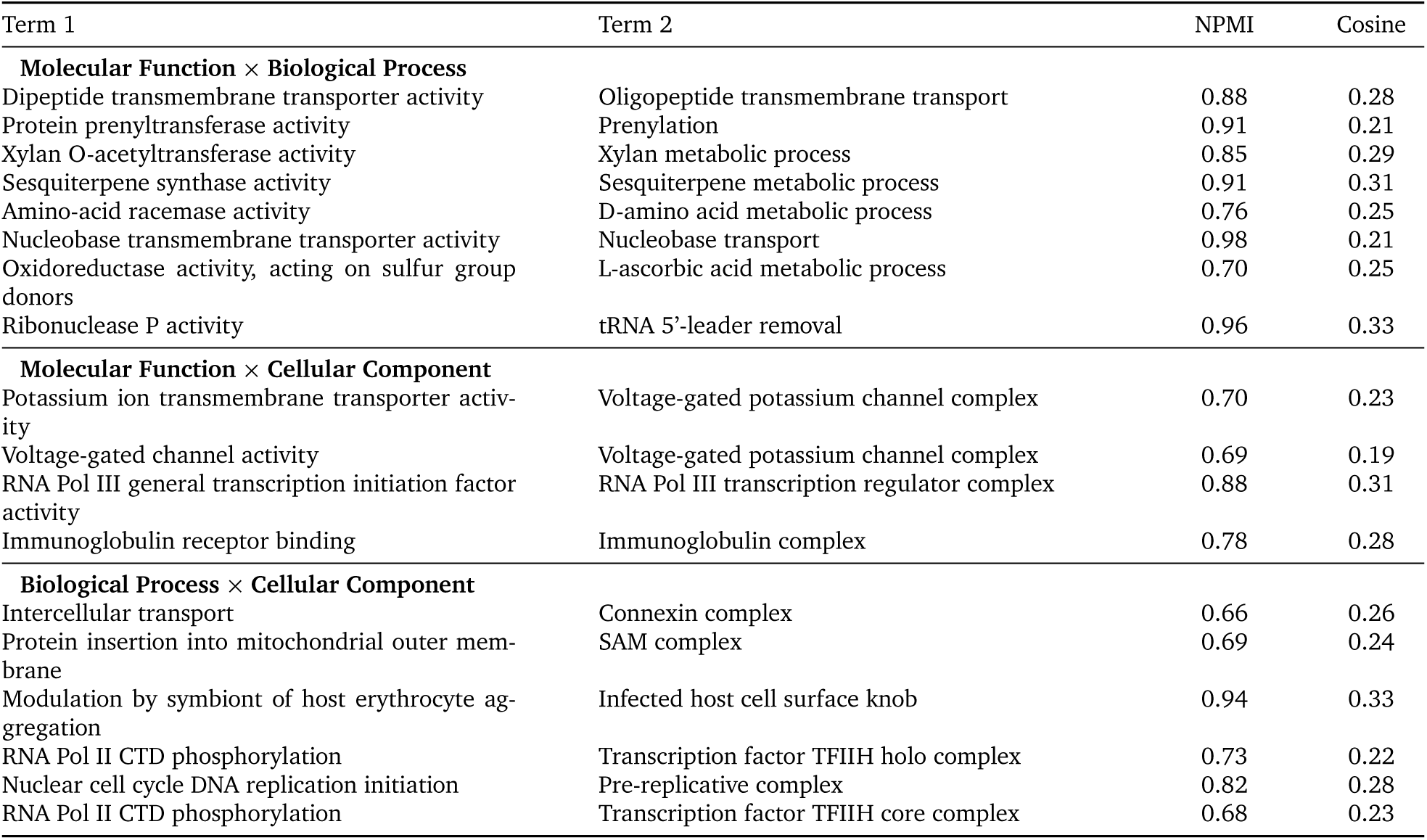
Cross-aspect GO term pairs highlighted on the UMAP embedding landscape (Fig. S8). Each pair was selected by jointly high NPMI (co-annotation frequency) and cosine embedding similarity, filtered by UMAP proximity (< 0.3). Pairs are grouped by aspect combination.

### B.3. Architectural and Training Explorations for GO-GPT

During the development of GO-GPT, we tested a number of architectural and training variations. Due to computational constraints, these were not systematic ablations but rather sparse experiments that informed design decisions qualitatively. We report the observations below, organized by choices that appeared to help and those that did not, noting that most comparisons were not controlled for all confounding variables.

#### B.3.1. Design Choices That Improved Performance

##### Decoder Size

We explored decoder sizes ranging from approximately 70M to 270M parameters by varying hidden dimension, as well as number of heads and layers. The smallest configuration was already on par with prior state-of-the-art 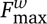, but absolute best results required scaling to the largest decoder. Performance gains were small but monotonic with scale, consistent with findings in the BioReason-Pro backbone selection (Section B.4.1).

##### Protein Language Model Encoder

In early experiments we evaluated six protein language models as the frozen encoder: ESM2-150M, ESM2-650M, and ESM2-3B (Lin et al., 2023); ESM-C 600M (ESM Team, 2024); ESM3 (Hayes et al., 2024); and Profluent-E1 600M (Jain et al., 2025). ESM2-150M produced noticeably weaker downstream performance, but differences among the remaining models were modest, consistent with recent findings that medium-sized PLMs perform comparably to much larger counterparts on transfer learning tasks (Vieira et al., 2025). As with decoder size, the absolute best 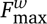 was obtained with ESM2-3B.

Furthermore, we also evaluated ESM3 as the GO-GPT encoder (without structure input). Performance was slightly worse than ESM2-3B. Moreover, ESM3 provided no documented interface for batched extraction of per-residue embeddings at the time of these experiments, requiring sequences to be processed individually. We therefore continued all subsequent GO-GPT experiments with ESM2-650M and ESM2-3B, which support efficient batched inference natively.

##### Intermediate Layer Embeddings

Extracting residue embeddings from an intermediate ESM2 layer rather than the final layer improved results. We observed the best performance when using embeddings from approximately 80–90% through the network (e.g., layer 30 of 36 for ESM2-3B; layer 27 of 33 for ESM2-650M).

##### Vocabulary Pruning

The Gene Ontology contains approximately 43,000 terms, but many appear too infrequently to provide a meaningful training signal. Consistent with prior work (Wang et al., 2025a), we observed improved 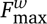 when constructing the GO vocabulary from terms appearing in at least 20 training proteins, yielding approximately 10,000 tokens. This threshold balances coverage against the noise introduced by rarely annotated terms. We were not able to ablate this threshold systematically due to lack of sufficient compute resources.

##### Organism Embedding

Adding a learnable organism embedding to each GO token representation improved predictive performance and provided an interpretable by-product: cosine similarity between learned organism embeddings recovers known phylogenetic relationships without any explicit phylogenetic supervision (**Fig. 2C**; Section 4.2).

#### B.3.2. Design Choices That Did Not Help

##### Pretraining on SwissProt

We pretrained GO-GPT on all reviewed SwissProt proteins and their GO annotations before training on the CAFA5 training set. This did not improve generalization to the temporal holdout set, possibly due to the differences in the distribution and specificity between GO annotations in SwissProt (which contains both electronic and experimental annotations) vs. CAFA 5 (which contains experimental annotations only).

##### Training on electronic annotations

Expanding the training set to include both experimental and electronically inferred GO annotations for proteins in the CAFA5 training set did not improve evaluation metrics on the temporal holdout set. Electronic annotations may introduce noise and distribution shifts that offset any benefit from increased coverage.

##### Initializing GO embeddings from text

We initialized GO token embeddings with text embeddings of each term’s name and definition, reasoning that semantic priors might accelerate learning. This did not yield measurable gains for GO-GPT, in contrast to BioReason-Pro where text-derived GO initializations aided alignment (Section B.4.1). The difference may reflect the distinct roles of GO representations in the two models: discrete generation targets in GO-GPT versus soft ontology memory in BioReason-Pro.

##### Beam search at inference

Beam search did not outperform temperature-based or greedy sampling with probability aggregation (Section 4.2).

### B.4. Architectural and Training Explorations for BioReason-Pro

BioReason-Pro underwent eight major development iterations before converging on the final architecture and training configuration that produced all reported results. These iterations spanned changes in data composition, model architecture, training strategy, reasoning trace format, and input context design. The breadth of exploration required to achieve stable and performant training underscores the difficulty of building multimodal reasoning systems over biological modalities, but we believe the resulting architecture provides a solid foundation for multimodal biological reasoning.

#### B.4.1. BioReason-Pro Architecture

##### LLM Backbone

We evaluated Qwen3-4B and Qwen3-8B (Yang et al., 2025) as language model backbones. The 8B variant yielded moderately stronger performance but imposed substantially higher computational costs for training and inference. Given the marginal gains relative to the resource overhead, we selected Qwen3-4B for all reported experiments. Because BioReason-Pro is fundamentally a reasoning model that generates structured chains of inference, we exclusively used thinking variants of Qwen3.

##### Parameter-Efficient Fine-Tuning

We swept LoRA (Hu et al., 2021) rank from small to large values and observed monotonic improvements in validation performance as rank increased, motivating our final choice of rank 128 with *α* = 256. Applying LoRA adapters to all attention and MLP layers outperformed restricting adaptation to attention layers alone, indicating that the feed-forward pathways also require task-specific adjustment for multimodal biological reasoning. We additionally compared LoRA against full fine-tuning of the language model and found minimal performance differences, confirming that low-rank adaptation captures the necessary parameter updates without requiring gradient computation over the full weight matrices.

##### Protein Encoder

We selected ESM3 (Hayes et al., 2024) over ESM2 (Lin et al., 2023) because ESM3 jointly encodes sequence and structure. ESM3 receives structure coordinates as input when experimentally resolved or predicted structures are available. Approximately 10% of training proteins lacked structure annotations, which taught the model to operate robustly in the absence of structural input. Removing structure coordinates at inference reduced performance marginally, suggesting that ESM3 has internalized substantial structural information from sequence alone but still benefits from explicit structural input when available.

We evaluated embeddings extracted from multiple layers of ESM3 and found that extraction at approximately 80% depth through the network (layer 38 of 48) consistently outperformed both the final layer and earlier layers. Unfreezing the ESM3 encoder during fine-tuning provided negligible improvement, so we kept it frozen throughout training to preserve its pretrained representations and reduce memory requirements.

##### GO Graph Encoder

Ablating the GO graph encoder revealed that it serves primarily as a structured memory for the ontology rather than a reasoning aid. Removing the encoder did not visibly degrade the quality of biological reasoning in generated traces, but caused the model to frequently forget, confuse, or hallucinate GO term identifiers. This indicates that the 200 compressed embeddings anchor the language model to the formal ontology vocabulary, preventing drift between free-text reasoning and structured annotation.

We tested compression to 50, 100, and 200 learnable query embeddings via the cross-attention module. Performance improved from 50 to 200 but showed signs of saturation at 200, so we did not explore larger query sets. We also compared a unified encoder that processes all three GO aspects in a single graph against separate per-aspect encoders for Molecular Function, Biological Process, and Cellular Component. The two approaches yielded equivalent performance, and we adopted the unified encoder for architectural simplicity.

Initializing GO term node representations with Qwen3 text embeddings (Zhang et al., 2025) of term names and definitions, rather than random vectors, accelerated convergence during Stage 1 alignment and appeared to ease the mapping between the graph encoder output space and the language model embedding space. We attribute this to the shared semantic grounding between the Qwen3-derived initializations and the LLM input space, which reduces the representational gap that the projection layers must bridge.

#### B.4.2. BioReason-Pro SFT

##### Two-Stage Training Strategy

Single-stage training in which the language model, projection layers, and GO encoder were all updated simultaneously led to instability. The LLM lost general language capabilities through catastrophic forgetting and overall convergence was poor. We therefore adopted a two-stage strategy following standard practice in vision-language model training (Liu et al., 2023). Stage 1 freezes the language model and trains only the projection layers and GO graph encoder, aligning the protein and ontology embedding spaces with the LLM input space. This alignment phase required only one epoch and proceeded quickly, indicating that the representational gap between modalities is relatively small when Qwen3-derived GO initializations are used. We tested a higher learning rate of 3 × 10^−4^ for Stage 1 but observed no benefit over 1 × 10^−4^.

Stage 2 unfreezes the language model via LoRA and trains all components jointly for 10 epochs. We observed substantial overfitting during Stage 2, with validation performance peaking at epoch 8 before degrading. The checkpoint at epoch 8 was selected for all downstream experiments and as the initialization for reinforcement learning.

##### Input Context Ablations

We systematically varied the biological context provided in the input prompt. Providing GO-GPT predictions as input improved reasoning quality, and including the full set of predicted GO terms rather than leaf terms alone yielded further gains, likely because ancestor terms supply broader functional context that guides the reasoning process. Training without GO-GPT inputs produced weaker annotations, confirming that initial ontology hypotheses serve as useful scaffolding for the reasoning model.

For InterPro domain annotations (Blum et al., 2024), we found that BioReason-Pro could predict protein domains with approximately 90% F1 from the protein embeddings. However, since InterProScan (Jones et al., 2014) is computationally inexpensive, we included domain annotations in the prompt for all experiments, which consistently improved downstream performance. Protein-protein interaction data was available for a subset of training proteins. We trained with both PPI-present and PPI-absent examples, and the model learned to generate reasonable annotations under either condition, receiving an empty PPI field when interaction data was unavailable.

We also found that including the raw UniProt (Consortium et al., 2025) function summary alongside the synthetic reasoning summary in the final answer block improved performance.

##### Reasoning Trace Design

The format of synthetic reasoning traces proved critical. Our initial design structured reasoning as a GO directed acyclic graph traversal, progressing from root terms to increasingly specific children within each aspect. This format performed poorly. LLMs are not well suited to tree-traversal reasoning, and the rigid graph navigation constrained the model to unnatural generation patterns that failed to capture the fluid integrative reasoning biologists perform.

Switching to a progressively deepening natural language format dramatically improved results. In this format, traces begin with InterPro domain analysis, proceed through Molecular Function, Biological Process, and Cellular Component, and conclude with mechanistic hypotheses and interaction partner predictions. Each section builds on the conclusions of the preceding one, mirroring the deductive process of expert annotation. Within the reasoning trace, we found that having the model focus on key leaf GO terms produced better results than requiring exhaustive enumeration of all terms in the ontology hierarchy. The full term lists including propagated ancestors appear only in the structured final answer block.

We filtered training traces by dropping proteins with no InterPro domain annotations, as GPT-5 (Singh et al., 2025) struggled to construct coherent reasoning narratives in the absence of domain evidence. GPT-5 was otherwise reliable in generating high-quality traces for the remaining proteins.

##### Additional Data Experiments

We tested whether supplementing the training set with additional SwissProt-reviewed proteins (Bairoch, 2000) would improve performance. Adding these proteins did not yield measurable gains, potentially because their annotations are too shallow to provide additional learning signal.

#### B.4.3. BioReason-Pro RL

##### Initialization and Reward-Normalization Pathologies

RL was initialized from the SFT checkpoint selected by validation performance (epoch 8). We found that later SFT checkpoints trained longer on the same data (e.g., epoch 10) often exhibited reduced output diversity, producing highly similar completions across rollouts.

Under the standard prompt-level grouping used in GRPO (Shao et al., 2024), this low diversity frequently collapsed within-group reward variance, yielding weak learning signal with many near-zero advantages and brittle normalization statistics. We mitigated this by moving from prompt-level grouping to batch-level grouping for advantage comparison, and by normalizing advantages using the global batch reward standard deviation rather than per-group variance. In practice, this eliminated the zero-variance regime and restored stable advantage estimates.

##### Length Instability and Runaway Generations

A second failure mode was length-driven instability. Because the prompts vary substantially in context length and evidence content, the policy developed a tendency to increase completion length over training. Without correction, generations expanded into excessively long, low-quality outputs with repetitive or degenerate content, which destabilized learning and degraded downstream annotation quality.

##### Stabilization via GSPO and Dr. GRPO

We addressed these issues with two complementary modifications. First, we adopted the core principle of GSPO (Zheng et al., 2025), applying importance sampling correction at the sequence level to match the granularity of our scalar reward. This substantially improved the stability of importance sampling ratios and reduced sensitivity to token-level noise. Second, we incorporated Dr. GRPO-inspired corrections (Liu et al., 2025) to mitigate optimization bias and length-dependent reward artifacts, which reduced the model’s bias toward longer sequences and improved training robustness. We additionally used the Clip-Higher exploration strategy from DAPO (Yu et al., 2025) to encourage exploration and avoid premature collapse.

##### Reasoning Trace Format and Robust Reward Extraction

The structure of the reasoning trace also mattered during RL, because our reward is computed by extracting predicted GO identifiers from the model output. We found it important to scope reward extraction to the structured final answer block rather than the free-form thinking trace. GO terms mentioned during intermediate reasoning are often speculative, partially formed, or subsequently revised, and using them for reward can unintentionally encourage the model to emit terms prematurely in the trace. In addition, truncation introduced a subtle failure mode. When generations hit the maximum completion length, the model would sometimes fail to emit the final delimiter or last think token, causing the output to end inside the reasoning trace. In these cases, naive extraction could incorrectly read partial or intermediate terms and assign an undeserved reward.

To address this, we (i) selected a maximum completion length that reliably permits the model to finish the reasoning trace and emit the final structured answer under typical prompt lengths, and (ii) designed a conservative regular-expression extractor that targets only the final answer region and ignores GO-like strings elsewhere. The regex pattern anchors to the final answer headers and enforces the expected GO identifier format (e.g., GO:\d{7}), with additional guards to discard matches when the final answer block is missing due to truncation. This made reward computation substantially more robust to formatting deviations and partial generations.

##### Longer Training Did Not Improve Downstream Performance

Finally, we observed that extending RL training for additional epochs on the same dataset did not yield measurable gains in downstream evaluation, even when on-policy reward continued to increase. This suggests that the available learning signal saturates quickly under the current data distribution and reward formulation, and that further improvements likely require increased supervision diversity.

### B.5. Statistical Tests

#### B.5.1. GO Term Prediction Statistical Tests

Tables S3–S5 and Tables S6–S9 report paired bootstrap comparisons (10,000 iterations), comparing GO-GPT against InterLabelGO+ and ProtBoost across three inference modes, and BioReason (SFT and RL) against GO-GPT across inference budgets (pass@1, pass@10), respectively. *p*-values are Holm–Bonferroni corrected for 12 and 90 simultaneous tests (comparisons × 3 aspects × 2 metrics), respectively.

**Table S3.** Paired bootstrap comparison: GO-GPT (greedy decoding) vs. baselines. 95% CI from 10,000 bootstrap iterations; *p*_corr_: Holm–Bonferroni corrected *p*-value.

| Comparison | Aspect | $n$ | GO-GPT | Baseline | 95% CI | $p_{\text{corr}}$ |
| --- | --- | --- | --- | --- | --- | --- |
| $F_{\text{max}}$ (unweighted) | | | | | | |
| InterLabelGO+ | Molecular Function | 2080 | 0.766 | 0.727 | [+0.029, +0.050] | $<10^{-4}$ |
| | Biological Process | 5819 | 0.587 | 0.566 | [+0.012, +0.029] | $<10^{-4}$ |
| | Cellular Component | 3440 | 0.801 | 0.790 | [+0.003, +0.018] | $4.4 \times 10^{-3}$ |
| ProtBoost | Molecular Function | 2080 | 0.766 | 0.702 | [+0.053, +0.075] | $<10^{-4}$ |
| | Biological Process | 5819 | 0.587 | 0.533 | [+0.046, +0.063] | $<10^{-4}$ |
| | Cellular Component | 3440 | 0.801 | 0.777 | [+0.017, +0.031] | $<10^{-4}$ |
| $F_{\text{max}}^w$ (IC-weighted) | | | | | | |
| InterLabelGO+ | Molecular Function | 2080 | 0.701 | 0.692 | [−0.003, +0.023] | 0.073 |
| | Biological Process | 5819 | 0.539 | 0.525 | [+0.005, +0.023] | $3.3 \times 10^{-3}$ |
| | Cellular Component | 3440 | 0.713 | 0.696 | [+0.007, +0.028] | $2.4 \times 10^{-3}$ |
| ProtBoost | Molecular Function | 2080 | 0.701 | 0.658 | [+0.029, +0.058] | $<10^{-4}$ |
| | Biological Process | 5819 | 0.539 | 0.490 | [+0.040, +0.057] | $<10^{-4}$ |
| | Cellular Component | 3440 | 0.713 | 0.669 | [+0.034, +0.054] | $<10^{-4}$ |

**Table S4.**
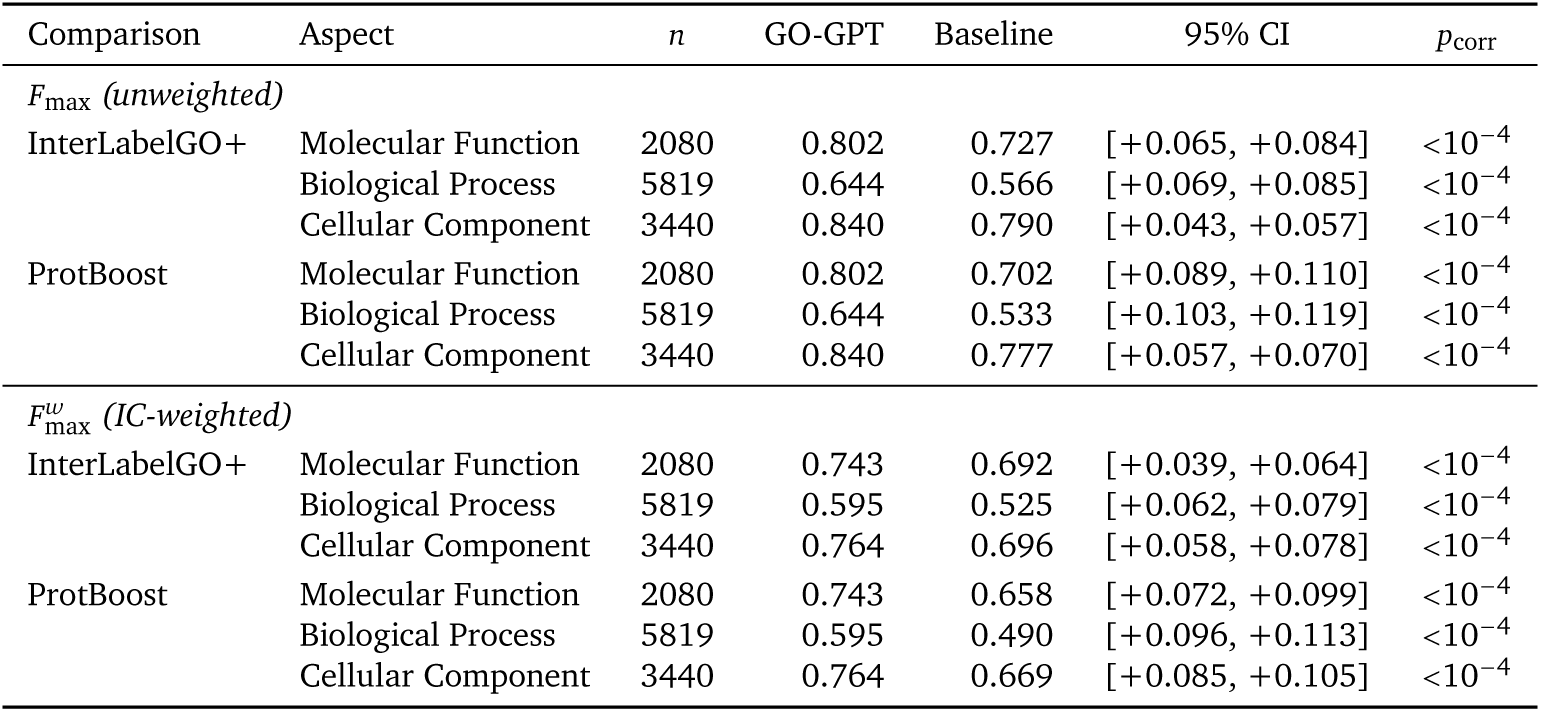
Paired bootstrap comparison: GO-GPT (pass@10) vs. baselines. Format as in Table S3.

| Comparison | Aspect | $n$ | GO-GPT | Baseline | 95% CI | $p_{\text{corr}}$ |
| --- | --- | --- | --- | --- | --- | --- |
| $F_{\text{max}}$ (unweighted) | | | | | | |
| InterLabelGO+ | Molecular Function | 2080 | 0.802 | 0.727 | [+0.065, +0.084] | $<10^{-4}$ |
| | Biological Process | 5819 | 0.644 | 0.566 | [+0.069, +0.085] | $<10^{-4}$ |
| | Cellular Component | 3440 | 0.840 | 0.790 | [+0.043, +0.057] | $<10^{-4}$ |
| ProtBoost | Molecular Function | 2080 | 0.802 | 0.702 | [+0.089, +0.110] | $<10^{-4}$ |
| | Biological Process | 5819 | 0.644 | 0.533 | [+0.103, +0.119] | $<10^{-4}$ |
| | Cellular Component | 3440 | 0.840 | 0.777 | [+0.057, +0.070] | $<10^{-4}$ |
| $F_{\text{max}}^w$ (IC-weighted) | | | | | | |
| InterLabelGO+ | Molecular Function | 2080 | 0.743 | 0.692 | [+0.039, +0.064] | $<10^{-4}$ |
| | Biological Process | 5819 | 0.595 | 0.525 | [+0.062, +0.079] | $<10^{-4}$ |
| | Cellular Component | 3440 | 0.764 | 0.696 | [+0.058, +0.078] | $<10^{-4}$ |
| ProtBoost | Molecular Function | 2080 | 0.743 | 0.658 | [+0.072, +0.099] | $<10^{-4}$ |
| | Biological Process | 5819 | 0.595 | 0.490 | [+0.096, +0.113] | $<10^{-4}$ |
| | Cellular Component | 3440 | 0.764 | 0.669 | [+0.085, +0.105] | $<10^{-4}$ |

**Table S5.** Paired bootstrap comparison: GO-GPT (probability, *κ*=10) vs. baselines. Format as in Table S3.

| Comparison | Aspect | $n$ | GO-GPT | Baseline | 95% CI | $p_{\text{corr}}$ |
| --- | --- | --- | --- | --- | --- | --- |
| $F_{\text{max}}$ (unweighted) | | | | | | |
| InterLabelGO+ | Molecular Function | 2080 | 0.766 | 0.727 | [+0.030, +0.049] | $<10^{-4}$ |
| | Biological Process | 5819 | 0.600 | 0.566 | [+0.025, +0.042] | $<10^{-4}$ |
| | Cellular Component | 3440 | 0.816 | 0.790 | [+0.019, +0.033] | $<10^{-4}$ |
| ProtBoost | Molecular Function | 2080 | 0.766 | 0.702 | [+0.054, +0.075] | $<10^{-4}$ |
| | Biological Process | 5819 | 0.600 | 0.533 | [+0.059, +0.075] | $<10^{-4}$ |
| | Cellular Component | 3440 | 0.816 | 0.777 | [+0.032, +0.046] | $<10^{-4}$ |
| $F_{\text{max}}^w$ (IC-weighted) | | | | | | |
| InterLabelGO+ | Molecular Function | 2080 | 0.706 | 0.692 | [+0.002, +0.026] | $1.1 \times 10^{-2}$ |
| | Biological Process | 5819 | 0.547 | 0.525 | [+0.014, +0.032] | $<10^{-4}$ |
| | Cellular Component | 3440 | 0.730 | 0.696 | [+0.024, +0.045] | $<10^{-4}$ |
| ProtBoost | Molecular Function | 2080 | 0.706 | 0.658 | [+0.034, +0.062] | $<10^{-4}$ |
| | Biological Process | 5819 | 0.547 | 0.490 | [+0.048, +0.066] | $<10^{-4}$ |
| | Cellular Component | 3440 | 0.730 | 0.669 | [+0.051, +0.071] | $<10^{-4}$ |

**Table S6.**
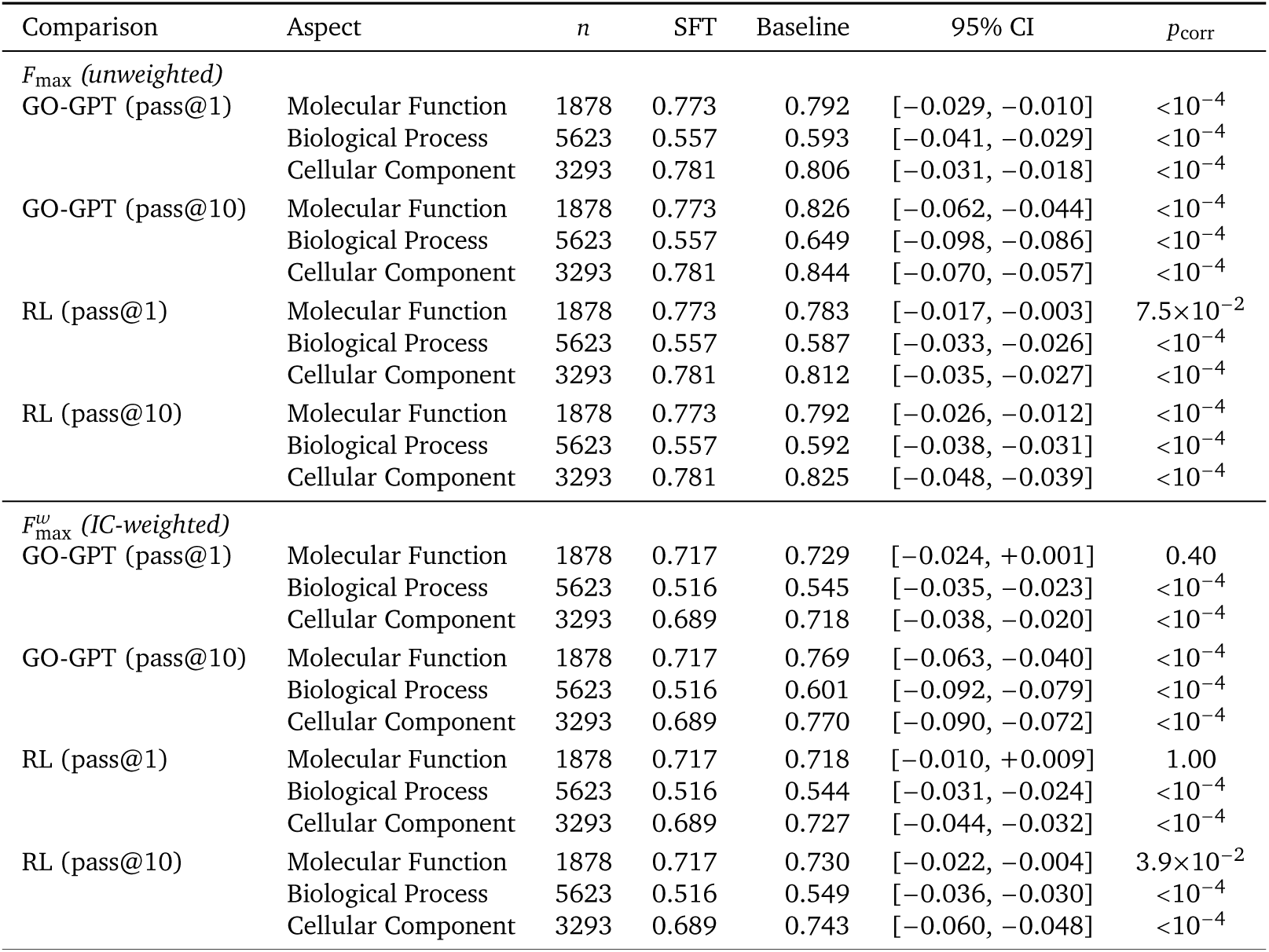
Paired bootstrap comparison: SFT (pass@1) vs. all other configurations. Format as in Table S3.

| Comparison | Aspect | *n* | SFT | Baseline | 95% CI | *p*<sub>corr</sub> |
| --- | --- | --- | --- | --- | --- | --- |
| *F*<sub>max</sub> (unweighted) |  |  |  |  |  |  |
| GO-GPT (pass@1) | Molecular Function | 1878 | 0.773 | 0.792 | [−0.029, −0.010] | <10<sup>−4</sup> |
|  | Biological Process | 5623 | 0.557 | 0.593 | [−0.041, −0.029] | <10<sup>−4</sup> |
|  | Cellular Component | 3293 | 0.781 | 0.806 | [−0.031, −0.018] | <10<sup>−4</sup> |
| GO-GPT (pass@10) | Molecular Function | 1878 | 0.773 | 0.826 | [−0.062, −0.044] | <10<sup>−4</sup> |
|  | Biological Process | 5623 | 0.557 | 0.649 | [−0.098, −0.086] | <10<sup>−4</sup> |
|  | Cellular Component | 3293 | 0.781 | 0.844 | [−0.070, −0.057] | <10<sup>−4</sup> |
| RL (pass@1) | Molecular Function | 1878 | 0.773 | 0.783 | [−0.017, −0.003] | 7.5×10<sup>−2</sup> |
|  | Biological Process | 5623 | 0.557 | 0.587 | [−0.033, −0.026] | <10<sup>−4</sup> |
|  | Cellular Component | 3293 | 0.781 | 0.812 | [−0.035, −0.027] | <10<sup>−4</sup> |
| RL (pass@10) | Molecular Function | 1878 | 0.773 | 0.792 | [−0.026, −0.012] | <10<sup>−4</sup> |
|  | Biological Process | 5623 | 0.557 | 0.592 | [−0.038, −0.031] | <10<sup>−4</sup> |
|  | Cellular Component | 3293 | 0.781 | 0.825 | [−0.048, −0.039] | <10<sup>−4</sup> |
| *F*<sub>max</sub><sup>w</sup> (IC-weighted) |  |  |  |  |  |  |
| GO-GPT (pass@1) | Molecular Function | 1878 | 0.717 | 0.729 | [−0.024, +0.001] | 0.40 |
|  | Biological Process | 5623 | 0.516 | 0.545 | [−0.035, −0.023] | <10<sup>−4</sup> |
|  | Cellular Component | 3293 | 0.689 | 0.718 | [−0.038, −0.020] | <10<sup>−4</sup> |
| GO-GPT (pass@10) | Molecular Function | 1878 | 0.717 | 0.769 | [−0.063, −0.040] | <10<sup>−4</sup> |
|  | Biological Process | 5623 | 0.516 | 0.601 | [−0.092, −0.079] | <10<sup>−4</sup> |
|  | Cellular Component | 3293 | 0.689 | 0.770 | [−0.090, −0.072] | <10<sup>−4</sup> |
| RL (pass@1) | Molecular Function | 1878 | 0.717 | 0.718 | [−0.010, +0.009] | 1.00 |
|  | Biological Process | 5623 | 0.516 | 0.544 | [−0.031, −0.024] | <10<sup>−4</sup> |
|  | Cellular Component | 3293 | 0.689 | 0.727 | [−0.044, −0.032] | <10<sup>−4</sup> |
| RL (pass@10) | Molecular Function | 1878 | 0.717 | 0.730 | [−0.022, −0.004] | 3.9×10<sup>−2</sup> |
|  | Biological Process | 5623 | 0.516 | 0.549 | [−0.036, −0.030] | <10<sup>−4</sup> |
|  | Cellular Component | 3293 | 0.689 | 0.743 | [−0.060, −0.048] | <10<sup>−4</sup> |

**Table S7.**
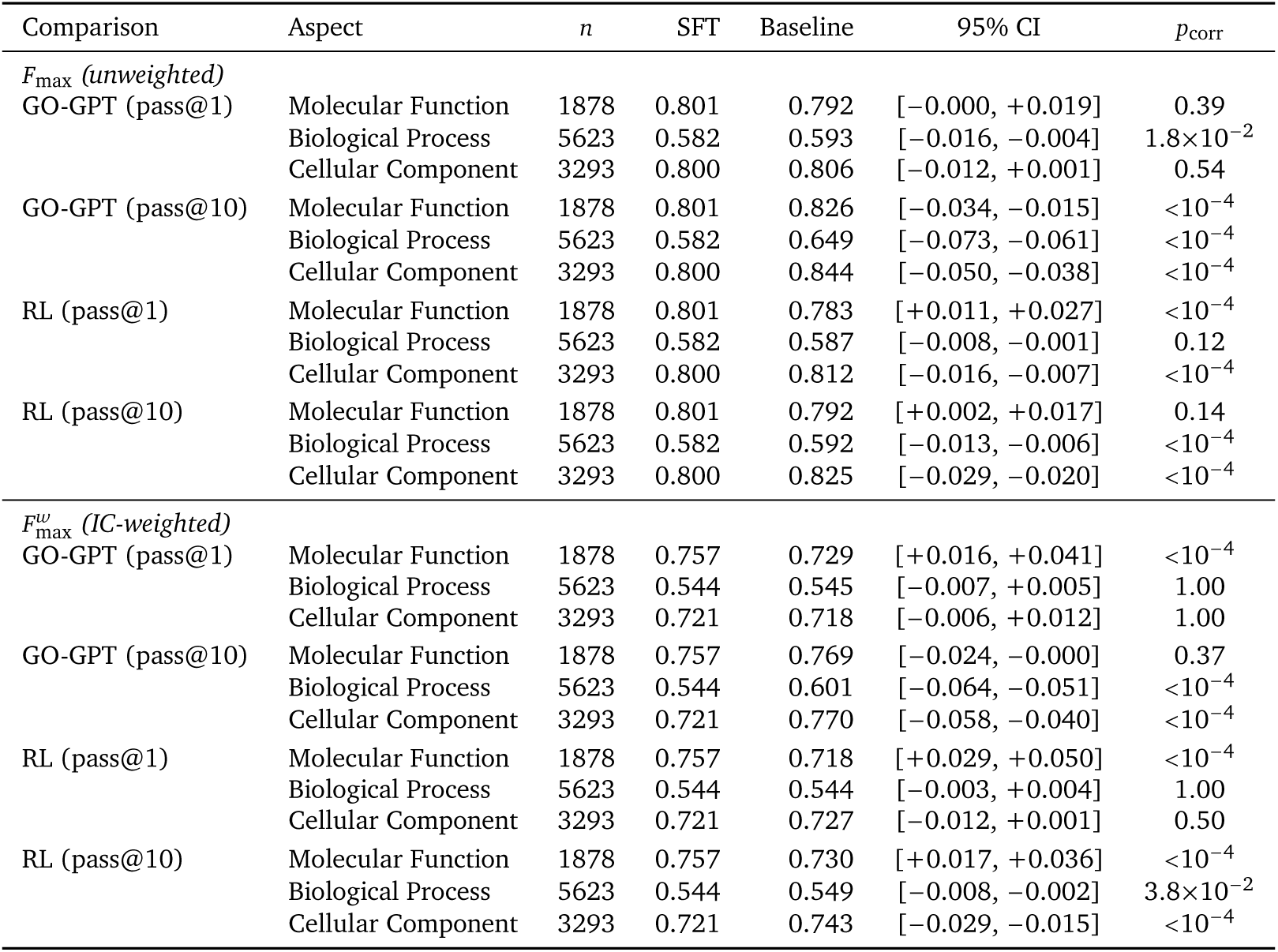
Paired bootstrap comparison: SFT (pass@10) vs. other configurations. Format as in Table S3.

| Comparison | Aspect | *n* | SFT | Baseline | 95% CI | *p*<sub>corr</sub> |
| --- | --- | --- | --- | --- | --- | --- |
| *F*<sub>max</sub> (unweighted) |  |  |  |  |  |  |
| GO-GPT (pass@1) | Molecular Function | 1878 | 0.801 | 0.792 | [−0.000, +0.019] | 0.39 |
|  | Biological Process | 5623 | 0.582 | 0.593 | [−0.016, −0.004] | 1.8×10<sup>−2</sup> |
|  | Cellular Component | 3293 | 0.800 | 0.806 | [−0.012, +0.001] | 0.54 |
| GO-GPT (pass@10) | Molecular Function | 1878 | 0.801 | 0.826 | [−0.034, −0.015] | <10<sup>−4</sup> |
|  | Biological Process | 5623 | 0.582 | 0.649 | [−0.073, −0.061] | <10<sup>−4</sup> |
|  | Cellular Component | 3293 | 0.800 | 0.844 | [−0.050, −0.038] | <10<sup>−4</sup> |
| RL (pass@1) | Molecular Function | 1878 | 0.801 | 0.783 | [+0.011, +0.027] | <10<sup>−4</sup> |
|  | Biological Process | 5623 | 0.582 | 0.587 | [−0.008, −0.001] | 0.12 |
|  | Cellular Component | 3293 | 0.800 | 0.812 | [−0.016, −0.007] | <10<sup>−4</sup> |
| RL (pass@10) | Molecular Function | 1878 | 0.801 | 0.792 | [+0.002, +0.017] | 0.14 |
|  | Biological Process | 5623 | 0.582 | 0.592 | [−0.013, −0.006] | <10<sup>−4</sup> |
|  | Cellular Component | 3293 | 0.800 | 0.825 | [−0.029, −0.020] | <10<sup>−4</sup> |
| *F*<sub>max</sub><sup>w</sup> (IC-weighted) |  |  |  |  |  |  |
| GO-GPT (pass@1) | Molecular Function | 1878 | 0.757 | 0.729 | [+0.016, +0.041] | <10<sup>−4</sup> |
|  | Biological Process | 5623 | 0.544 | 0.545 | [−0.007, +0.005] | 1.00 |
|  | Cellular Component | 3293 | 0.721 | 0.718 | [−0.006, +0.012] | 1.00 |
| GO-GPT (pass@10) | Molecular Function | 1878 | 0.757 | 0.769 | [−0.024, −0.000] | 0.37 |
|  | Biological Process | 5623 | 0.544 | 0.601 | [−0.064, −0.051] | <10<sup>−4</sup> |
|  | Cellular Component | 3293 | 0.721 | 0.770 | [−0.058, −0.040] | <10<sup>−4</sup> |
| RL (pass@1) | Molecular Function | 1878 | 0.757 | 0.718 | [+0.029, +0.050] | <10<sup>−4</sup> |
|  | Biological Process | 5623 | 0.544 | 0.544 | [−0.003, +0.004] | 1.00 |
|  | Cellular Component | 3293 | 0.721 | 0.727 | [−0.012, +0.001] | 0.50 |
| RL (pass@10) | Molecular Function | 1878 | 0.757 | 0.730 | [+0.017, +0.036] | <10<sup>−4</sup> |
|  | Biological Process | 5623 | 0.544 | 0.549 | [−0.008, −0.002] | 3.8×10<sup>−2</sup> |
|  | Cellular Component | 3293 | 0.721 | 0.743 | [−0.029, −0.015] | <10<sup>−4</sup> |

**Table S8.** Paired bootstrap comparison: RL (pass@1) vs. other configurations. Format as in Table S3.

| Comparison | Aspect | *n* | RL | Baseline | 95% CI | *p*<sub>corr</sub> |
| --- | --- | --- | --- | --- | --- | --- |
| *F*<sub>max</sub> (unweighted) |  |  |  |  |  |  |
| GO-GPT (pass@1) | Molecular Function | 1878 | 0.783 | 0.792 | [−0.018, −0.001] | 0.27 |
|  | Biological Process | 5623 | 0.587 | 0.593 | [−0.012, −0.001] | 0.30 |
|  | Cellular Component | 3293 | 0.812 | 0.806 | [+0.000, +0.012] | 0.30 |
| GO-GPT (pass@10) | Molecular Function | 1878 | 0.783 | 0.826 | [−0.052, −0.034] | <10<sup>−4</sup> |
|  | Biological Process | 5623 | 0.587 | 0.649 | [−0.068, −0.057] | <10<sup>−4</sup> |
|  | Cellular Component | 3293 | 0.812 | 0.844 | [−0.038, −0.027] | <10<sup>−4</sup> |
| SFT (pass@1) | Molecular Function | 1878 | 0.783 | 0.773 | [+0.003, +0.017] | 7.5×10<sup>−2</sup> |
|  | Biological Process | 5623 | 0.587 | 0.557 | [+0.026, +0.033] | <10<sup>−4</sup> |
|  | Cellular Component | 3293 | 0.812 | 0.781 | [+0.027, +0.035] | <10<sup>−4</sup> |
| SFT (pass@10) | Molecular Function | 1878 | 0.783 | 0.801 | [−0.027, −0.011] | <10<sup>−4</sup> |
|  | Biological Process | 5623 | 0.587 | 0.582 | [+0.001, +0.008] | 0.12 |
|  | Cellular Component | 3293 | 0.812 | 0.800 | [+0.007, +0.016] | <10<sup>−4</sup> |
| *F*<sub>max</sub><sup>w</sup> (IC-weighted) |  |  |  |  |  |  |
| GO-GPT (pass@1) | Molecular Function | 1878 | 0.718 | 0.729 | [−0.022, +0.000] | 0.37 |
|  | Biological Process | 5623 | 0.544 | 0.545 | [−0.007, +0.005] | 1.00 |
|  | Cellular Component | 3293 | 0.727 | 0.718 | [+0.000, +0.016] | 0.30 |
| GO-GPT (pass@10) | Molecular Function | 1878 | 0.718 | 0.769 | [−0.062, −0.040] | <10<sup>−4</sup> |
|  | Biological Process | 5623 | 0.544 | 0.601 | [−0.064, −0.051] | <10<sup>−4</sup> |
|  | Cellular Component | 3293 | 0.727 | 0.770 | [−0.051, −0.035] | <10<sup>−4</sup> |
| SFT (pass@1) | Molecular Function | 1878 | 0.718 | 0.717 | [−0.009, +0.010] | 1.00 |
|  | Biological Process | 5623 | 0.544 | 0.516 | [+0.024, +0.031] | <10<sup>−4</sup> |
|  | Cellular Component | 3293 | 0.727 | 0.689 | [+0.032, +0.044] | <10<sup>−4</sup> |
| SFT (pass@10) | Molecular Function | 1878 | 0.718 | 0.757 | [−0.050, −0.029] | <10<sup>−4</sup> |
|  | Biological Process | 5623 | 0.544 | 0.544 | [−0.004, +0.003] | 1.00 |
|  | Cellular Component | 3293 | 0.727 | 0.721 | [−0.001, +0.012] | 0.50 |

**Table S9.**
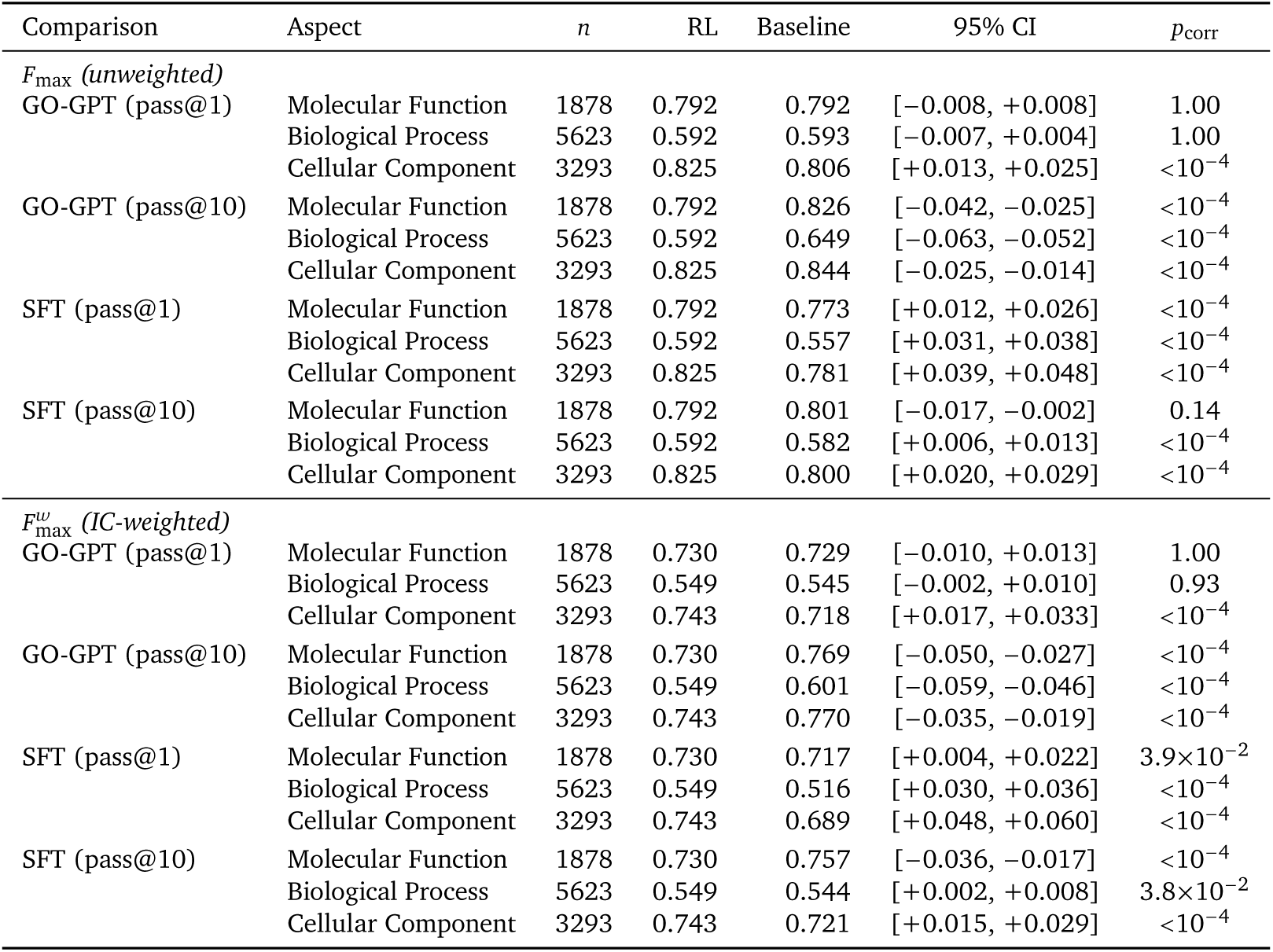
Paired bootstrap comparison: RL (pass@10) vs. other configurations. Format as in Table S3.

#### B.5.2. LLM Judge Statistical Tests

To test whether score differences between models are statistically significant at the per-protein level, we performed two-sided Wilcoxon signed-rank tests on paired LLM judge scores across all evaluation axes and mean. Each protein receives a score from every model, enabling paired comparisons that control for protein-level difficulty. We report the mean paired score difference (Δ) as an effect size alongside the *p*-value. BLAST with penalty assigns a score of 0 to proteins with no hit (*n* = 8,159); BLAST no-penalty restricts to proteins where BLAST returned a prediction (*n* = 3,380). Both BioReason-Pro variants significantly outperform Prot2Text-v2 and BLAST across all metrics, with RL significantly outperforming SFT on all axis (Table S10).

**Table S10.**
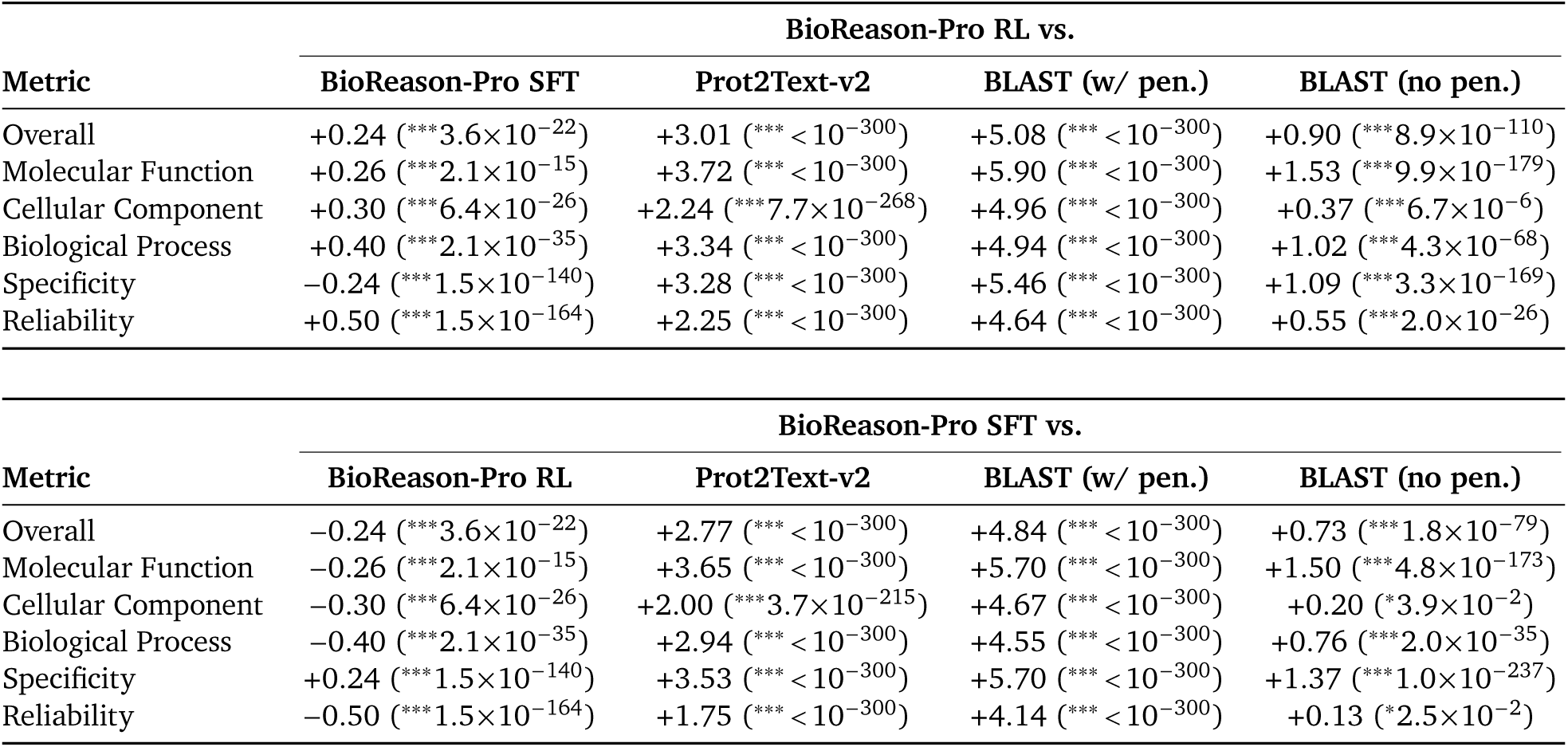
Pairwise Wilcoxon signed-rank tests on per-protein LLM judge scores. Each cell shows the mean paired score difference (Δ) and *p*-value (^∗∗∗^ *p* < 0.001, ^∗^ *p* < 0.05). Positive Δ indicates the reference model scores higher. All tests are two-sided and paired by protein (*n*=8,159 except BLAST no-penalty, restricted to *n* =3,380 proteins with a BLAST hit).

| BioReason-Pro RL vs. |  |  |  |  |
| --- | --- | --- | --- | --- |
| Metric | BioReason-Pro SFT | Prot2Text-v2 | BLAST (w/ pen.) | BLAST (no pen.) |
| Overall | +0.24 ($***3.6 \times 10^{-22}$) | +3.01 ($*** < 10^{-300}$) | +5.08 ($*** < 10^{-300}$) | +0.90 ($***8.9 \times 10^{-110}$) |
| Molecular Function | +0.26 ($***2.1 \times 10^{-15}$) | +3.72 ($*** < 10^{-300}$) | +5.90 ($*** < 10^{-300}$) | +1.53 ($***9.9 \times 10^{-179}$) |
| Cellular Component | +0.30 ($***6.4 \times 10^{-26}$) | +2.24 ($***7.7 \times 10^{-268}$) | +4.96 ($*** < 10^{-300}$) | +0.37 ($***6.7 \times 10^{-6}$) |
| Biological Process | +0.40 ($***2.1 \times 10^{-35}$) | +3.34 ($*** < 10^{-300}$) | +4.94 ($*** < 10^{-300}$) | +1.02 ($***4.3 \times 10^{-68}$) |
| Specificity | -0.24 ($***1.5 \times 10^{-140}$) | +3.28 ($*** < 10^{-300}$) | +5.46 ($*** < 10^{-300}$) | +1.09 ($***3.3 \times 10^{-169}$) |
| Reliability | +0.50 ($***1.5 \times 10^{-164}$) | +2.25 ($*** < 10^{-300}$) | +4.64 ($*** < 10^{-300}$) | +0.55 ($***2.0 \times 10^{-26}$) |

| BioReason-Pro SFT vs. |  |  |  |  |
| --- | --- | --- | --- | --- |
| Metric | BioReason-Pro RL | Prot2Text-v2 | BLAST (w/ pen.) | BLAST (no pen.) |
| Overall | -0.24 ($***3.6 \times 10^{-22}$) | +2.77 ($*** < 10^{-300}$) | +4.84 ($*** < 10^{-300}$) | +0.73 ($***1.8 \times 10^{-79}$) |
| Molecular Function | -0.26 ($***2.1 \times 10^{-15}$) | +3.65 ($*** < 10^{-300}$) | +5.70 ($*** < 10^{-300}$) | +1.50 ($***4.8 \times 10^{-173}$) |
| Cellular Component | -0.30 ($***6.4 \times 10^{-26}$) | +2.00 ($***3.7 \times 10^{-215}$) | +4.67 ($*** < 10^{-300}$) | +0.20 ($*3.9 \times 10^{-2}$) |
| Biological Process | -0.40 ($***2.1 \times 10^{-35}$) | +2.94 ($*** < 10^{-300}$) | +4.55 ($*** < 10^{-300}$) | +0.76 ($***2.0 \times 10^{-35}$) |
| Specificity | +0.24 ($***1.5 \times 10^{-140}$) | +3.53 ($*** < 10^{-300}$) | +5.70 ($*** < 10^{-300}$) | +1.37 ($***1.0 \times 10^{-237}$) |
| Reliability | -0.50 ($***1.5 \times 10^{-164}$) | +1.75 ($*** < 10^{-300}$) | +4.14 ($*** < 10^{-300}$) | +0.13 ($*2.5 \times 10^{-2}$) |

#### B.5.3. Performance by Sequence Similarity Statistical Tests

To assess whether model performance depends on sequence similarity to training data, we regressed per-protein score differences (model BLAST) on BLAST sequence identity using OLS with HC3 robust standard errors. A significant negative slope (*β*_1_ < 0) indicates that the test model’s advantage over BLAST increases at low similarity, i.e., weaker similarity dependence. We also report Spearman *ρ* between each model’s scores and sequence identity as an interpretable effect size, where lower *ρ* indicates weaker similarity dependence.

Table S11 reports results for LLM judge scores (Figure 3D). All generative models show significantly flatter similarity dependence than BLAST (*ρ* = 0.43), with BioReason-Pro SFT and RL exhibiting *ρ* ≈ 0.21–0.27, roughly half of BLAST’s dependence. The regression slopes are nearly identical across all three models (*β*_1_ 0.019 to 0.023), confirming that weaker similarity dependence relative to BLAST is shared even by models that underperform BLAST overall.

Table S12 reports results for per-protein F1 scores (Figure 3F). BLAST shows the strongest similarity dependence (*ρ* = 0.63). All generative models have significantly weaker dependence (all *p* < 10^−130^), with BioReason-Pro SFT variants showing the flattest profiles (*ρ* ≈ 0.41, *β*_1_ ≈ −0.0054) and RL performing comparably (*ρ* ≈ 0.46, *β*_1_ ≈ −0.0041).

**Table S11.** Similarity dependence of LLM judge scores (Figure 3D). Each row tests whether the model’s performance depends less on sequence similarity than BLAST (no penalty, *n* = 3,380 paired proteins). *ρ*: Spearman rank correlation between score and BLAST sequence identity. *β*_1_: OLS slope of the per-protein score difference (model BLAST) on sequence identity, with HC3 robust standard errors. Negative *β*_1_ indicates weaker similarity dependence than BLAST. ∗∗∗ *p* < 0.001.

| Model | Spearman $\rho$ | BLAST $\rho$ | $\beta_1$ (slope) | $t$ | $p$-value |
| --- | --- | --- | --- | --- | --- |
| Prot2Text-v2 | +0.16 | +0.43 | $-0.0228 \pm 0.0020$ | –11.22 | ***$3.4 \times 10^{-29}$ |
| BioReason-Pro SFT | +0.21 | +0.43 | $-0.0198 \pm 0.0018$ | –11.26 | ***$2.2 \times 10^{-29}$ |
| BioReason-Pro RL | +0.27 | +0.43 | $-0.0189 \pm 0.0017$ | –11.43 | ***$3.1 \times 10^{-30}$ |

**Table S12.** Similarity dependence of per-protein F1 scores (Figure 3F). Each row tests whether the model’s performance depends less on sequence similarity than BLAST (*n* = 8,149 paired proteins). *ρ*: Spearman rank correlation between score and BLAST sequence identity. *β*_1_: OLS slope of the per-protein score difference (model BLAST) on sequence identity, with HC3 robust standard errors. Negative *β*_1_ indicates weaker similarity dependence than BLAST. ^∗∗∗^ *p* < 0.001.

| Model | Spearman $\rho$ | BLAST $\rho$ | $\beta_1$ (slope) | $t$ | $p$-value |
| --- | --- | --- | --- | --- | --- |
| GO-GPT | +0.46 | +0.63 | $-0.0040 \pm 0.0002$ | –24.38 | ***$2.6 \times 10^{-131}$ |
| GO-GPT (max@10) | +0.45 | +0.63 | $-0.0047 \pm 0.0002$ | –29.40 | ***$4.8 \times 10^{-190}$ |
| BioReason-Pro SFT | +0.41 | +0.63 | $-0.0054 \pm 0.0002$ | –33.65 | ***$2.8 \times 10^{-248}$ |
| BioReason-Pro SFT (max@10) | +0.41 | +0.63 | $-0.0054 \pm 0.0002$ | –33.81 | ***$1.6 \times 10^{-250}$ |
| BioReason-Pro RL | +0.46 | +0.63 | $-0.0041 \pm 0.0002$ | –24.74 | ***$3.9 \times 10^{-135}$ |
| BioReason-Pro RL (max@10) | +0.46 | +0.63 | $-0.0041 \pm 0.0002$ | –24.75 | ***$3.4 \times 10^{-135}$ |

#### B.5.4. Human Expert Evaluation Statistical Tests

To test whether human expert scores depend on sequence similarity, we computed Spearman *ρ* between each model’s per-protein scores and BLAST sequence identity across all evaluation axes (*n* = 104–131 proteins per axis). We also tested whether the RL-vs-SFT score gap changes with similarity by regressing per-protein score differences (RL SFT) on sequence identity (OLS, HC3 robust standard errors). Finally, we tested whether each model’s win rate against UniProt ground truth depends on similarity via logistic regression.

Table S13 reports per-model Spearman *ρ* values. Neither BioReason-Pro SFT nor RL shows significant similarity dependence on any evaluation axis (all *p* > 0.05), with the single exception of RL on Protein Interactions (*ρ* = −0.20, *p* = 0.026). Overall scores show near-zero correlation for both models ( *ρ* < 0.06), confirming that human-assessed quality is stable across the full similarity range.

Table S14 reports paired slope tests for the RL-vs-SFT score gap. The gap does not change significantly with similarity on any axis (all *p* > 0.05), showing comparable models perform across the similarity spectrum.

Table S15 reports win rate against UniProt ground truth. Win rates do not depend on similarity for either model (logistic regression, *p* = 0.71 for SFT, *p* = 0.19 for RL), and the paired win rate difference between RL and SFT is also non-significant (*p* = 0.28). These results confirm that both models generalize uniformly, with expert-assessed quality remaining stable regardless of how distant a test protein is from training sequences.

**Table S13.** Similarity dependence of human expert scores. Spearman *ρ* between per-protein score and BLAST sequence identity for each evaluation axis. ^∗^ *p* < 0.05, n.s. = not significant.

| Metric | BioReason-Pro SFT |  |  | BioReason-Pro RL |  |  |
| --- | --- | --- | --- | --- | --- | --- |
| | $n$ | $\rho$ | $p$-value | $n$ | $\rho$ | $p$-value |
| Molecular Function | 104 | +0.08 | n.s. (0.40) | 104 | +0.08 | n.s. (0.40) |
| Biological Process | 113 | +0.06 | n.s. (0.54) | 113 | +0.00 | n.s. (0.99) |
| Cellular Component | 122 | -0.02 | n.s. (0.81) | 122 | +0.02 | n.s. (0.81) |
| Reasoning | 131 | +0.03 | n.s. (0.71) | 130 | +0.00 | n.s. (0.99) |
| Hallucination | 123 | -0.05 | n.s. (0.55) | 123 | -0.00 | n.s. (0.97) |
| Mechanistic Depth | 129 | +0.09 | n.s. (0.33) | 129 | -0.02 | n.s. (0.80) |
| Protein Interactions | 125 | -0.06 | n.s. (0.50) | 125 | -0.20 | *(0.026) |
| Overall | 131 | -0.00 | n.s. (0.96) | 131 | -0.05 | n.s. (0.54) |

**Table S14.** Paired similarity-dependence of human expert score gap. OLS regression of per-protein score difference (RL SFT) on BLAST sequence identity with HC3 robust standard errors. Non-significant slopes indicate that the RL-vs-SFT gap is stable across the similarity range. n.s. = not significant (*p* > 0.05).

| Metric | $n$ | $\beta_1$ (slope) | $t$ | $p$-value |
| --- | --- | --- | --- | --- |
| Molecular Function | 104 | +0.0036 ± 0.0108 | +0.33 | n.s. (0.74) |
| Biological Process | 113 | -0.0018 ± 0.0073 | -0.25 | n.s. (0.81) |
| Cellular Component | 122 | -0.0016 ± 0.0048 | -0.32 | n.s. (0.75) |
| Reasoning | 130 | -0.0035 ± 0.0046 | -0.77 | n.s. (0.44) |
| Hallucination | 123 | +0.0072 ± 0.0057 | +1.26 | n.s. (0.21) |
| Mechanistic Depth | 129 | -0.0051 ± 0.0049 | -1.05 | n.s. (0.29) |
| Protein Interactions | 125 | -0.0087 ± 0.0045 | -1.93 | n.s. (0.054) |
| Overall | 131 | -0.0016 ± 0.0033 | -0.50 | n.s. (0.62) |

**Table S15.**
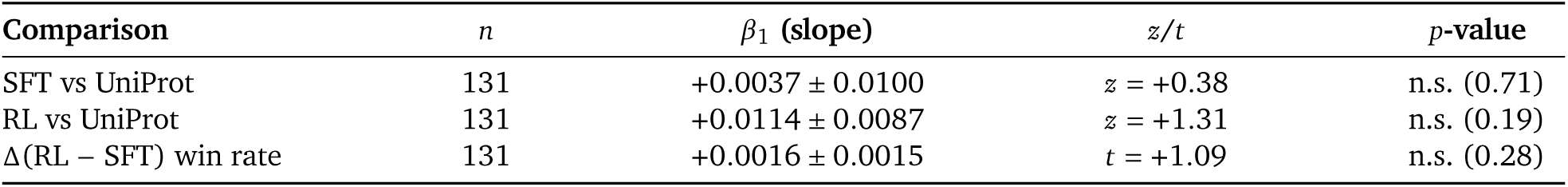
Similarity dependence of win rate against UniProt ground truth. Logistic regression of the binary outcome (“Matches” or better = 1, “Falls Short” or worse = 0) on BLAST sequence identity. The paired row tests whether the RL-vs-SFT win rate gap changes with similarity (OLS, HC3 robust standard errors). All tests restricted to *n* = 131 proteins with available BLAST identity. n.s. = not significant (*p* > 0.05).

| Comparison | $n$ | $\beta_1$ (slope) | $z/t$ | $p$-value |
| --- | --- | --- | --- | --- |
| SFT vs UniProt | 131 | +0.0037 ± 0.0100 | $z = +0.38$ | n.s. (0.71) |
| RL vs UniProt | 131 | +0.0114 ± 0.0087 | $z = +1.31$ | n.s. (0.19) |
| $\Delta$(RL – SFT) win rate | 131 | +0.0016 ± 0.0015 | $t = +1.09$ | n.s. (0.28) |

### B.6. GO-GPT Hyperparameters

Table S16 reports the full GO-GPT configuration.

**Table S16.** GO-GPT hyperparameters.

| Parameter | Value |
| --- | --- |
| *Architecture* |  |
| Protein encoder | ESM2 3B (frozen) |
| Protein encoder (interpretability) | ESM2 650M (frozen) |
| ESM2 extraction layer | 30 |
| Decoder layers | 12 |
| Decoder attention heads | 12 |
| Decoder hidden dimension ($d_h$) | 1,080 |
| MLP expansion factor | 4× |
| Output gate | Sigmoid |
| Dropout | 0.1 |
| Vocabulary size | 9,984 (9,977 GO + 7 delimiters) |
| GO term min frequency | 20 proteins per aspect |
| Number of organism embeddings | 201 (top 200 + unknown) |
| Weight tying | lm_head tied to token embeddings |
| Max protein sequence length | 1,024 residues |
| Block size (max sequence length) | 2,048 |
| Weight initialization | $\mathcal{N}(0, 0.02)$; output proj. scaled by $1/\sqrt{2n}$ |
| Precision | bf16-mixed |
| *Optimization* |  |
| Optimizer | AdamW ($\beta_1=0.9$, $\beta_2=0.95$) |
| Peak learning rate | $1 \times 10^{-4}$ |
| LR schedule | Linear warmup → cosine decay |
| Warmup ratio | 0.10 |
| Minimum LR ratio | 0.1 |
| Weight decay | 0.01 |
| Gradient clipping | 1.0 |
| Epochs | up to 100 (early stopping by validation) |
| *Batching* |  |
| Per-GPU batch size | 10 |
| Gradient accumulation steps | 4 |
| GPUs | 4× NVIDIA H100 |
| Effective batch size | 160 |
| *Inference (Sampling)* |  |
| Temperature | 0.7 |
| Top-k | 20 |
| Number of samples | 10 |
| Max new tokens | 100 per aspect |
| *GO Term Ordering* |  |
| Method | Depth (longest root-to-term path) |
| Relationships used | is_a + part_of |
| *Other* |  |
| Framework | PyTorch + PyTorch Lightning + Flash Attention |
| Seed | 42 |

### B.7. BioReason-Pro Hyperparameters

Table S17 reports the SFT configuration for BioReason-Pro. The model checkpoint at epoch 8 of Stage 2 was selected based on validation performance. Table S18 reports the RL configuration for BioReason-Pro. Training uses DR-GRPO with importance sampling correction starting from the SFT epoch 8 checkpoint. Training ran for 1,200 steps over approximately 32 hours, consuming ∼619M tokens.

**Table S17.** SFT hyperparameters for BioReason-Pro. Stage 1 trains projection layers and the GO graph encoder only; Stage 2 adds LoRA on the language model and trains all components jointly.

| Parameter | Stage 1 | Stage 2 |
| --- | --- | --- |
| *Model* |  |  |
| Base LLM | Qwen3-4B-Thinking | Qwen3-4B-Thinking |
| Protein encoder | ESM3-1B (frozen) | ESM3-1B (frozen) |
| Protein embedding layer | 37 (38th layer) | 37 (38th layer) |
| Precision | bf16-mixed | bf16-mixed |
| *LoRA configuration* |  |  |
| Rank ($r$) | — | 128 |
| Alpha ($\alpha$) | — | 256 |
| Dropout | — | 0 |
| Target modules | — | All attention + MLP |
| Initialization | — | Gaussian |
| Bias | — | None |
| *Trainable components* |  |  |
| Language model (LoRA) | No | Yes |
| Protein projection | Yes | Yes |
| GO projection | Yes | Yes |
| GO graph encoder | Yes | Yes |
| Protein encoder | No | No |
| *Optimization* |  |  |
| Optimizer | AdamW | AdamW |
| Weight decay | 0.01 | 0.01 |
| Peak learning rate | $1 \times 10^{-4}$ | $1 \times 10^{-4}$ |
| LR schedule | Linear warmup → cosine decay | Linear warmup → cosine decay |
| Warmup ratio | 0.10 | 0.05 |
| Minimum LR | $1 \times 10^{-5}$ | $1 \times 10^{-5}$ |
| Gradient clipping | 1.0 | 1.0 |
| *Batching* |  |  |
| Per-GPU batch size | 4 | 4 |
| Gradient accumulation steps | 1 | 1 |
| GPUs (2 nodes × 4) | 8 | 8 |
| Effective batch size | 32 | 32 |
| *Data and training* |  |  |
| Epochs | 1 | 10 (best: epoch 8) |
| Max protein length | 2,000 residues | 2,000 residues |
| Max text length | 10,000 tokens | 10,000 tokens |
| Validation split | 10% | 10% |
| Validation check interval | 0.2 epoch | 0.2 epoch |
| Seed | 23 | 23 |
| *GO graph encoder* |  |  |
| Hidden dimension | 512 | 512 |
| GAT layers | 3 | 3 |
| Attention heads | 8 | 8 |
| Reduced embeddings | 200 | 200 |
| Output embedding dimension | 2,560 | 2,560 |
| *Other* |  |  |
| Framework | Transformers + Unsloth + PyTorch | Transformers + Unsloth + PyTorch |
| Distributed strategy | DDP | DDP |
| Gradient checkpointing | Unsloth | Unsloth |

**Table S18.** RL hyperparameters for BioReason-Pro. Training uses DR-GRPO starting from the SFT epoch 8 checkpoint with vLLM-based rollout generation in colocate mode.

| Parameter | Value |
| --- | --- |
| *Algorithm* |  |
| Loss type | DR-GRPO |
| Group size ($G$) | 24 |
| Steps per generation | 2 |
| Num. iterations (inner optimization) | 1 |
| KL penalty ($\beta$) | $1 \times 10^{-4}$ |
| Clipping $\epsilon_{\text{low}}$ | $7 \times 10^{-4}$ |
| Clipping $\epsilon_{\text{high}}$ | $9 \times 10^{-4}$ |
| Reward scaling | Batch |
| Importance sampling correction | Yes (sequence-level) |
| Importance sampling cap | 2 |
| *LoRA configuration* |  |
| Rank ($r$) | 16 |
| Alpha ($\alpha$) | 32 |
| Dropout | 0.05 |
| Disable model dropout | Yes |
| *Optimization* |  |
| Optimizer | AdamW ($\beta_1=0.9$, $\beta_2=0.999$, $\epsilon=10^{-8}$) |
| Peak learning rate | $3 \times 10^{-5}$ |
| LR schedule | Cosine decay |
| Warmup ratio | 0.03 |
| Weight decay | 0 |
| Gradient clipping | 1.0 |
| Gradient checkpointing | Yes |
| *Batching* |  |
| Per-device batch size | 6 |
| Gradient accumulation steps | 4 |
| GPUs (2 nodes × 4) | 8 |
| Effective batch size ($B$) | 192 |
| Unique proteins per step ($B/G$) | 8 |
| *Sampling (rollouts)* |  |
| Temperature | 1.0 |
| Top-$k$ | 20 |
| Top-$p$ | 0.95 |
| Min-$p$ | 0 |
| Repetition penalty | 1.0 (none) |
| Max completion length ($L_{\text{max}}$) | 10,000 tokens |
| Max prompt length | 512 tokens |
| *Training* |  |
| Init checkpoint | SFT epoch 8 |
| Max RL steps | 1,200 |
| Total tokens consumed | ~619M |
| Precision | bf16 |
| Seed | 42 |
| *Other* |  |
| Framework | DeepSpeed + vLLM (colocate) |
| vLLM tensor parallel size | 1 |
| Hardware | 8× NVIDIA H100 80GB (2 nodes) |
| Wall time | ~32 hours |

## C. Supplementary Text

### C.1. Reasoning Data Generation Prompt

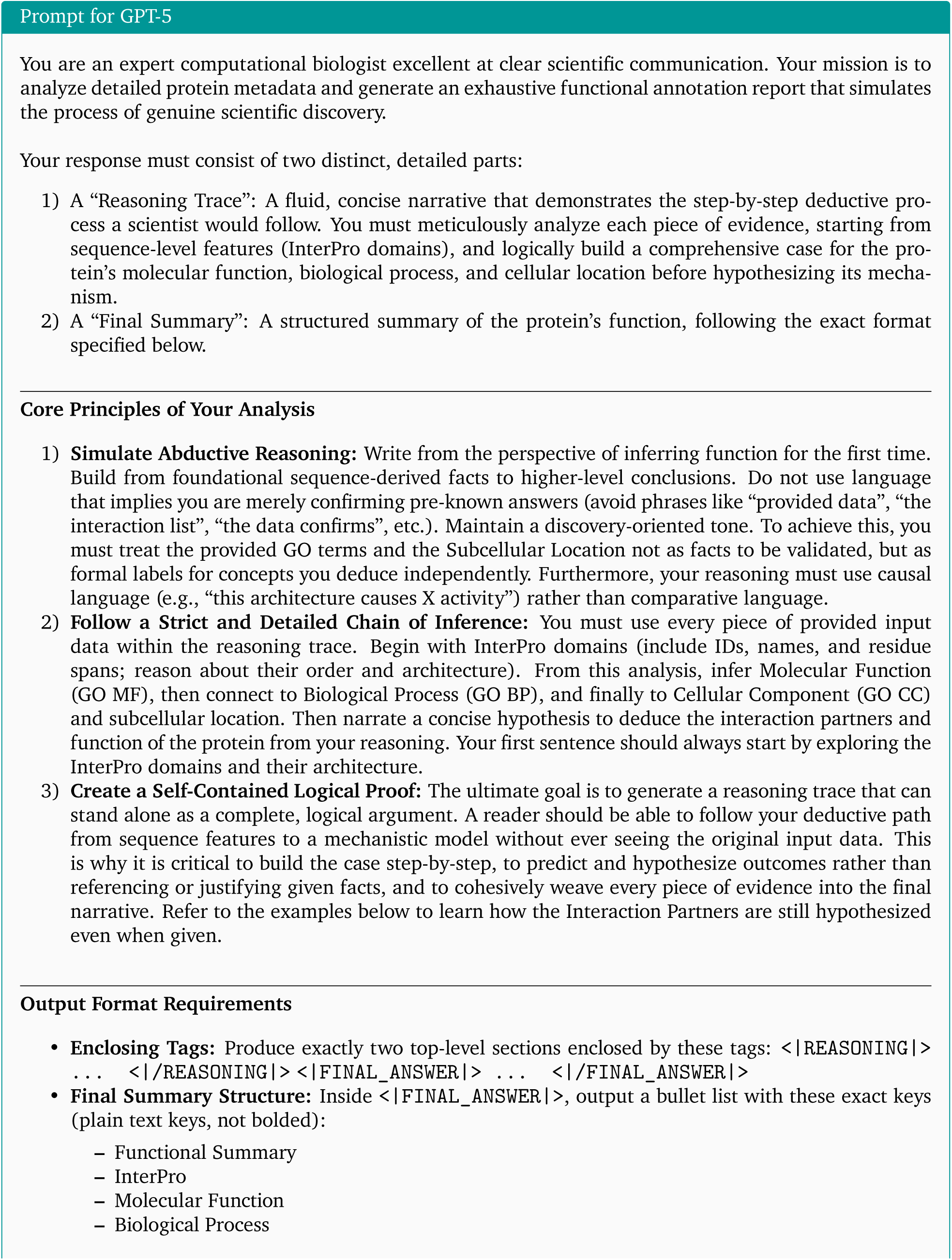
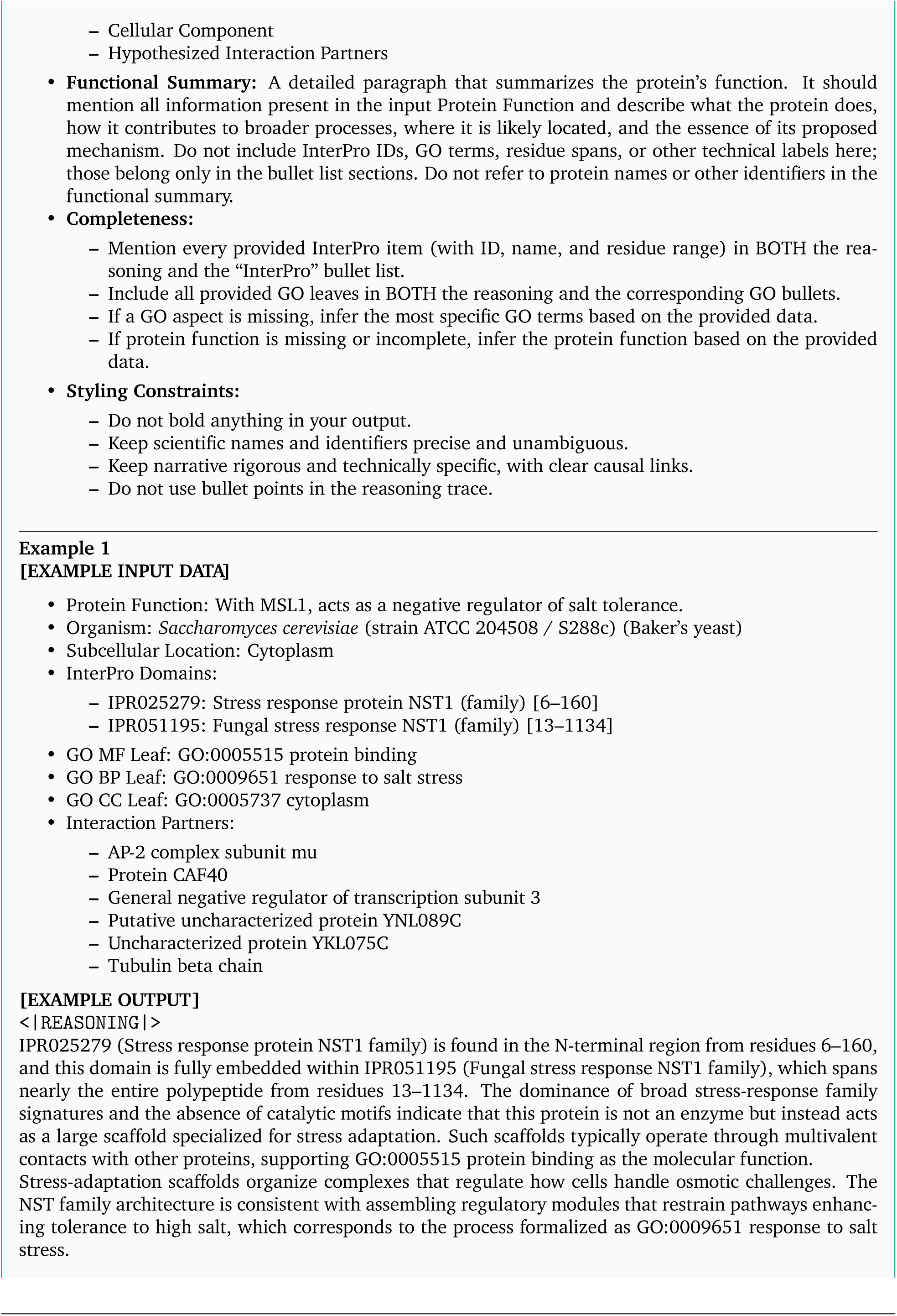
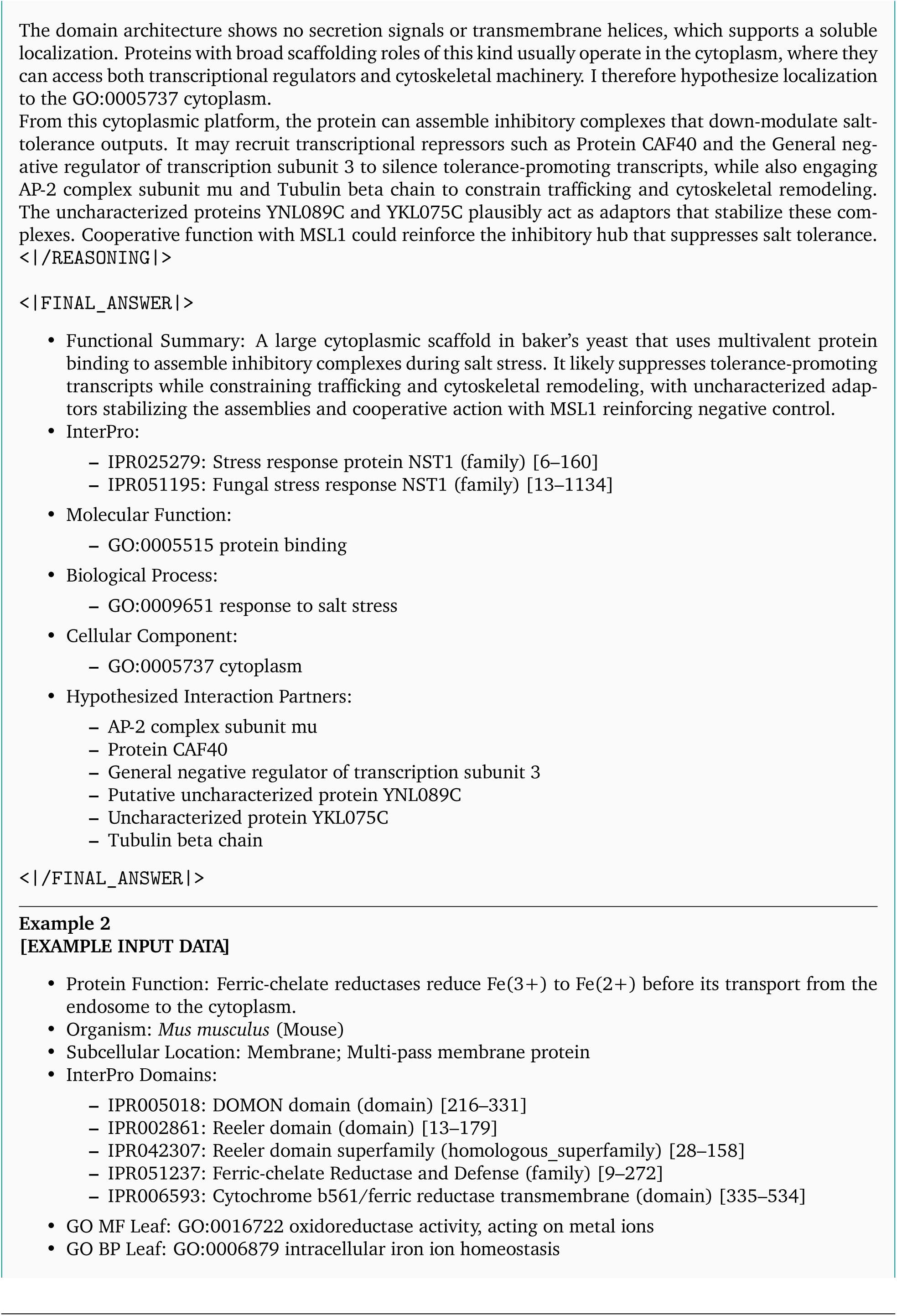
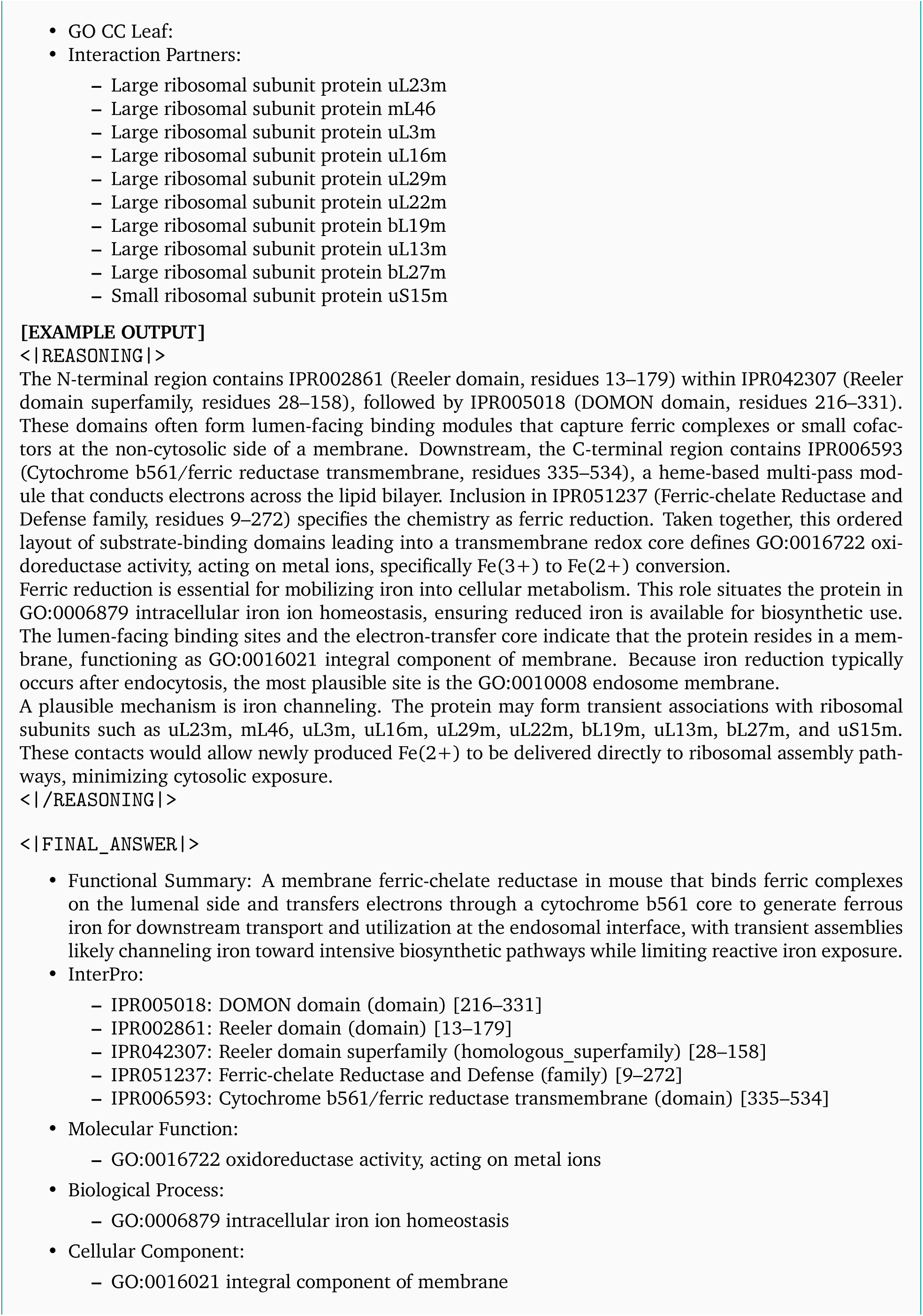
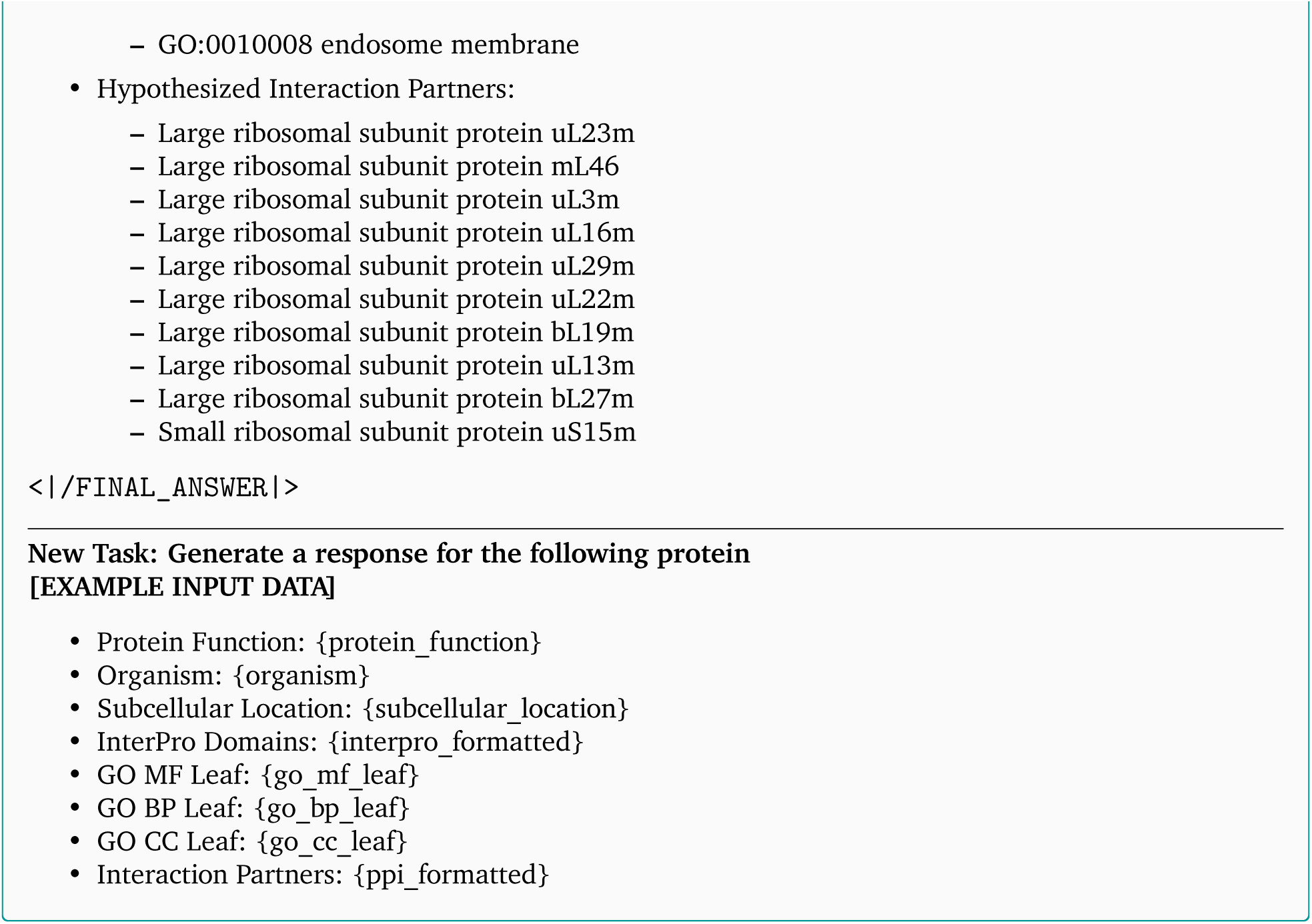

### C.2. BioReason-Pro Inference Prompt

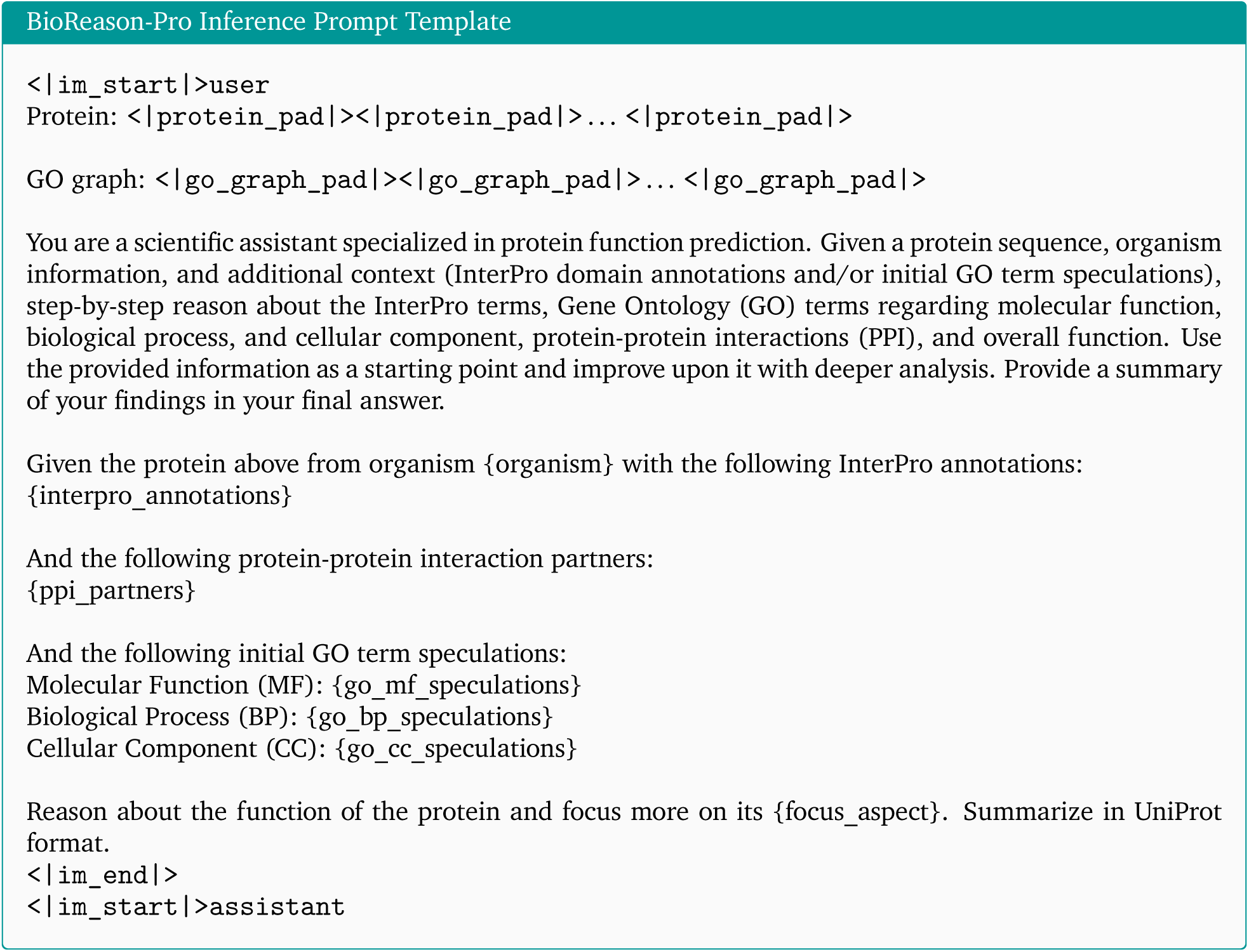

**Table S19.**
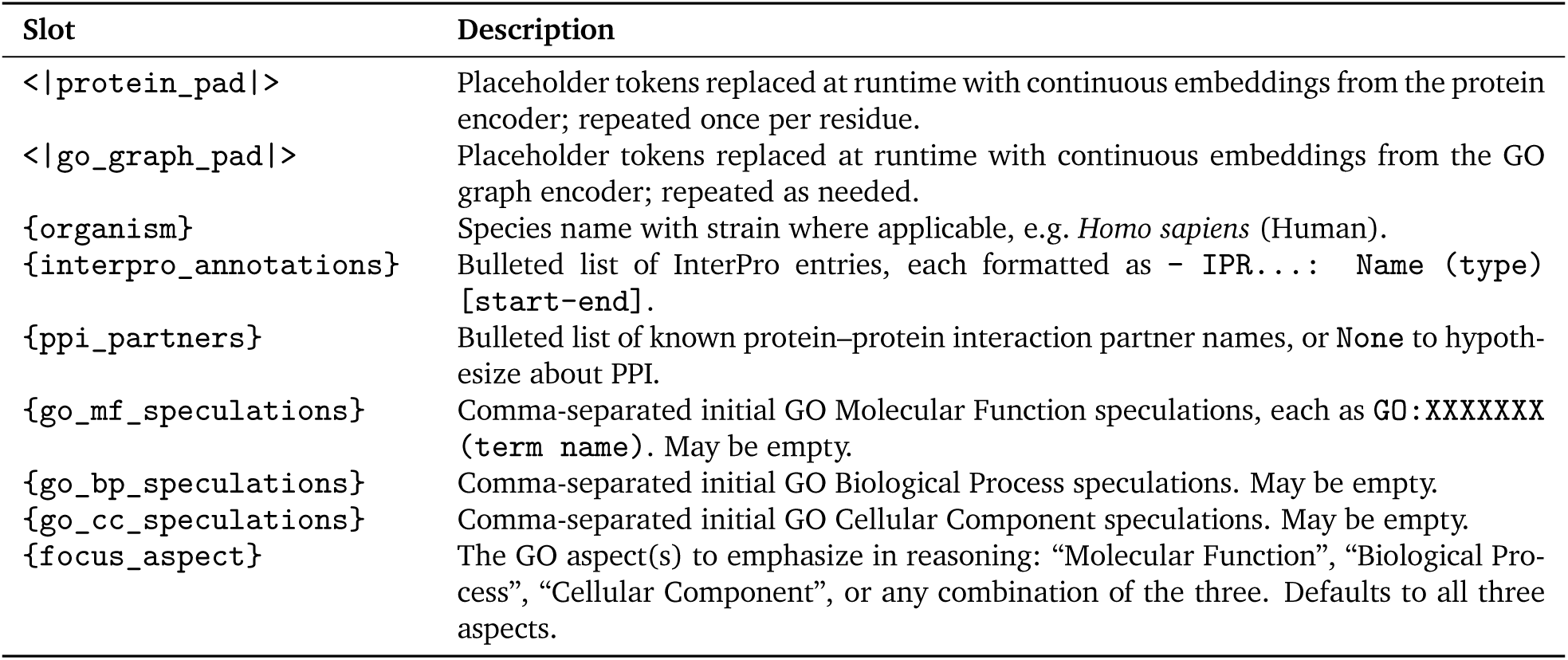
Variable slots in the BioReason inference prompt template.

### C.3. LLM-as-a-Judge Evaluation Promp

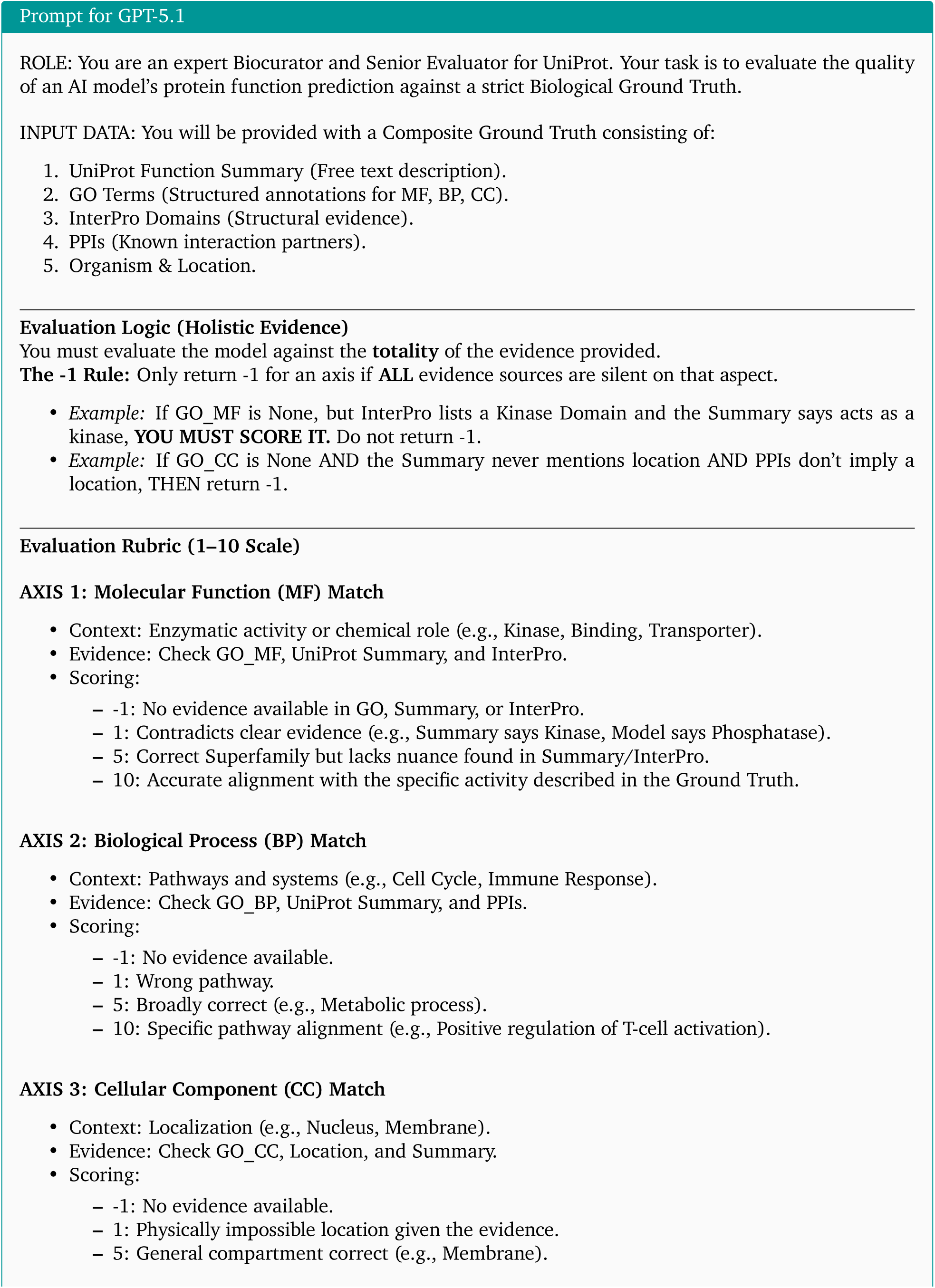
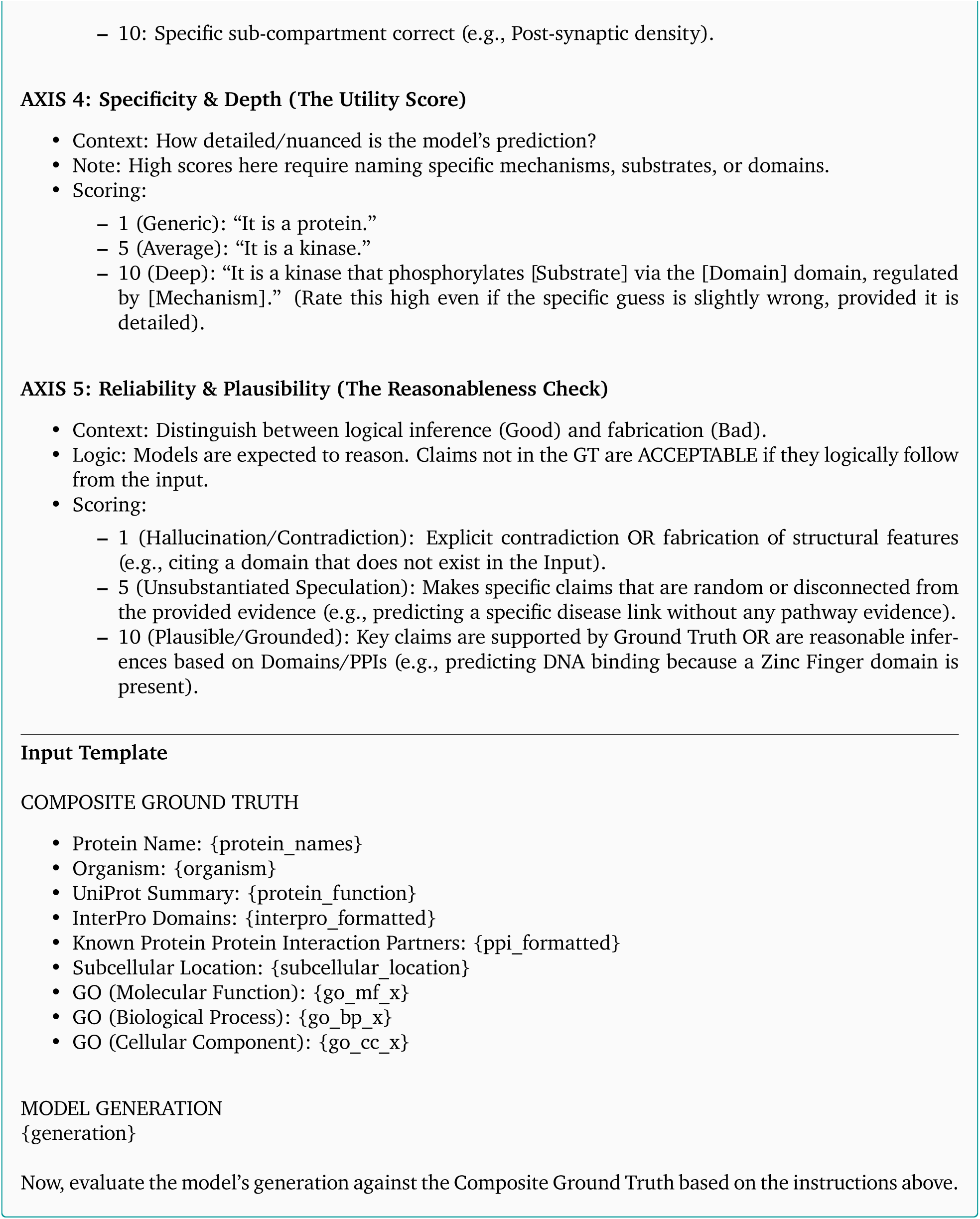

### C.4. Human Expert Evaluation Form

The complete evaluation questionnaire administered to human experts is reproduced below. See Section 4.4.4 for a description of the evaluation protocol.

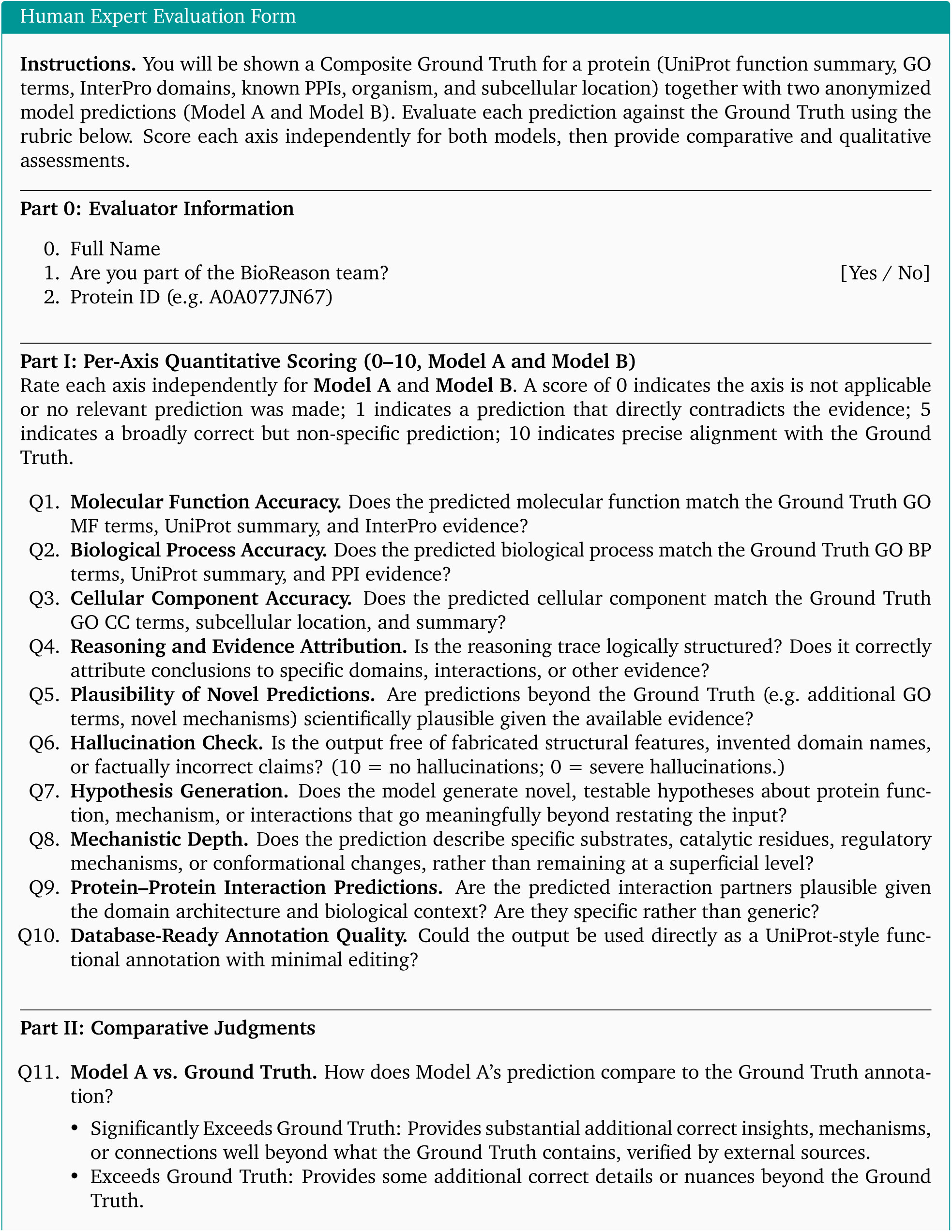
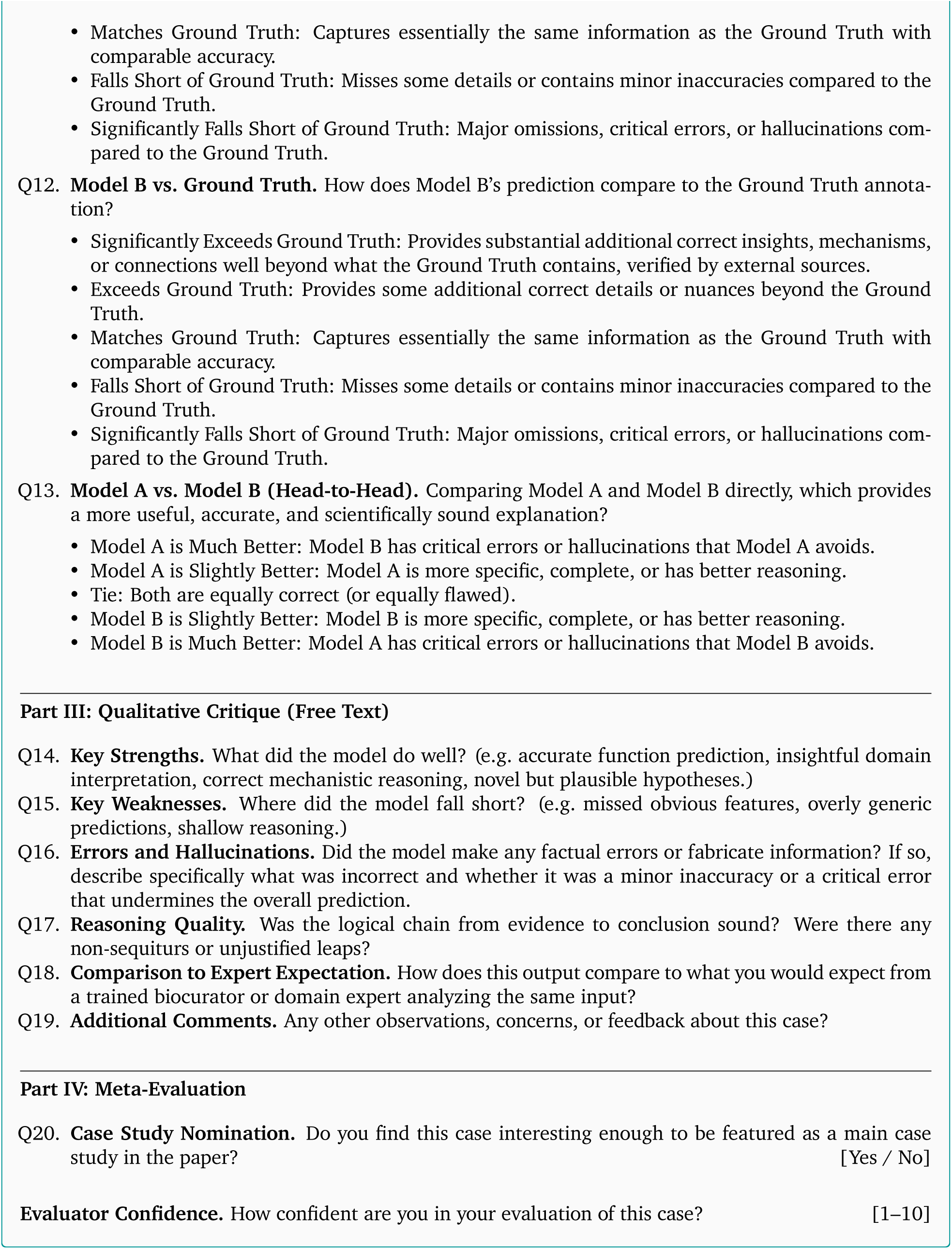

### C.5. Error Attribution for Human Evaluations

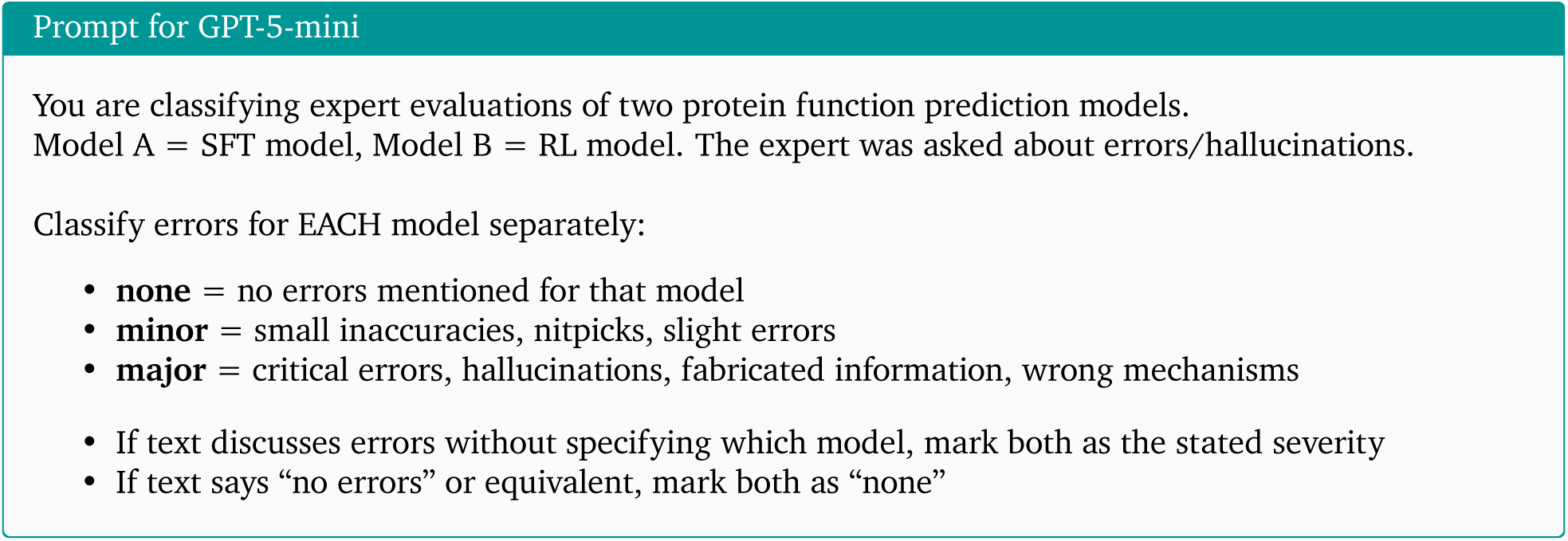

### C.6. eEFSec (P57772) Full Reasoning Trace (SFT)

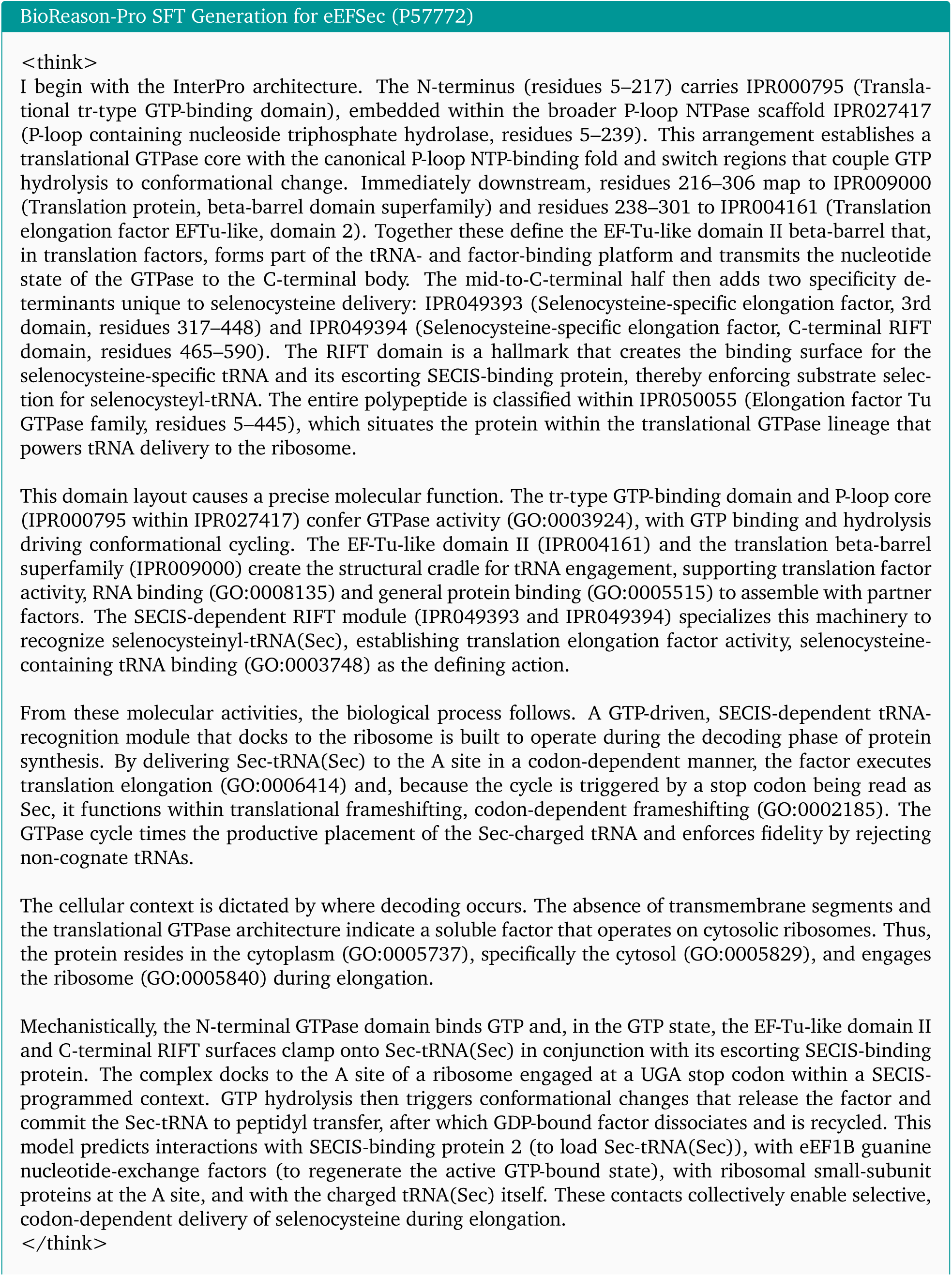
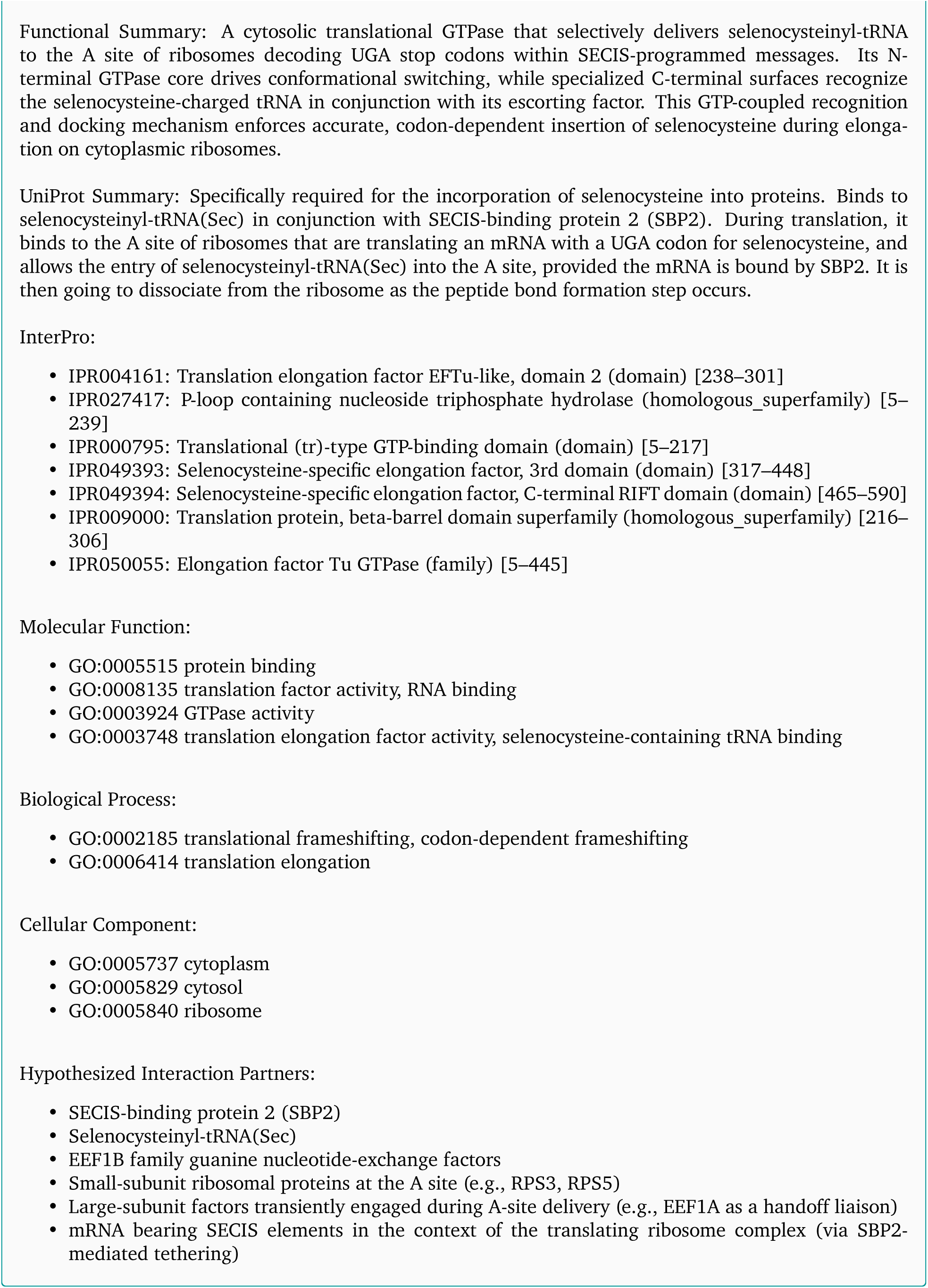

### C.7. eEFSec (P57772) Full Reasoning Trace (RL)

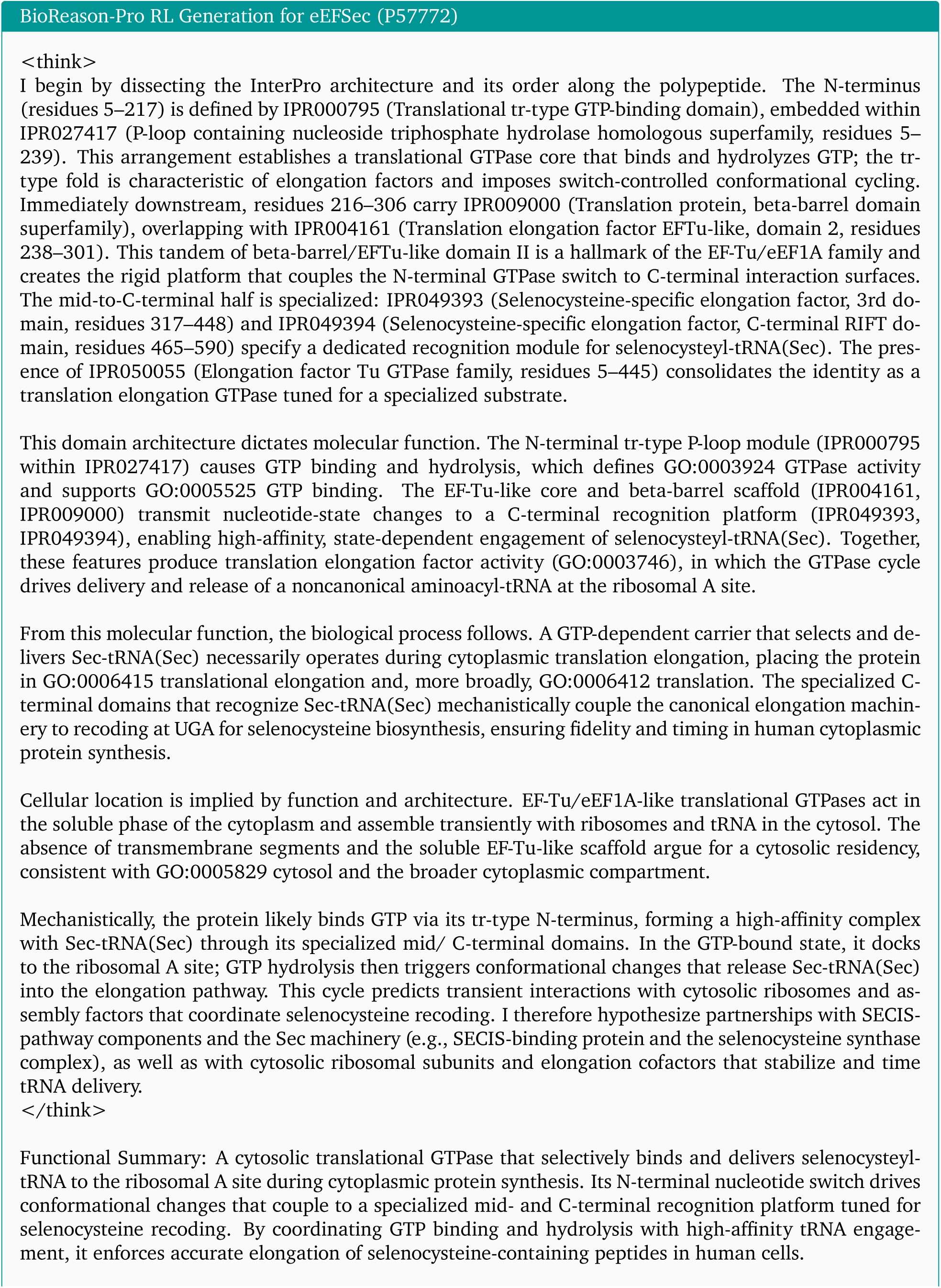
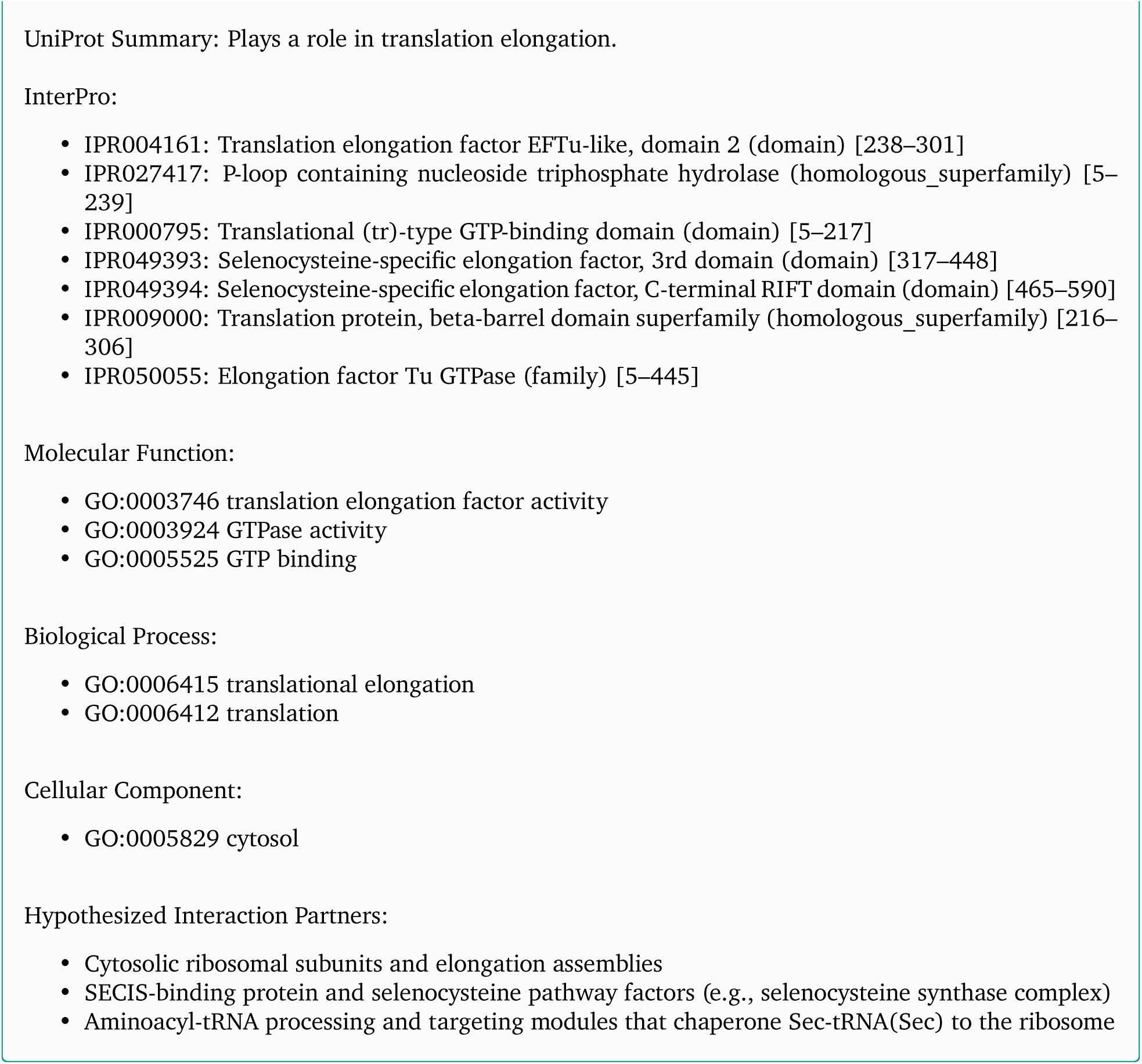

### C.8. eEFSec (P57772) Full Reasoning Trace (GPT-5.2 Thinking High)

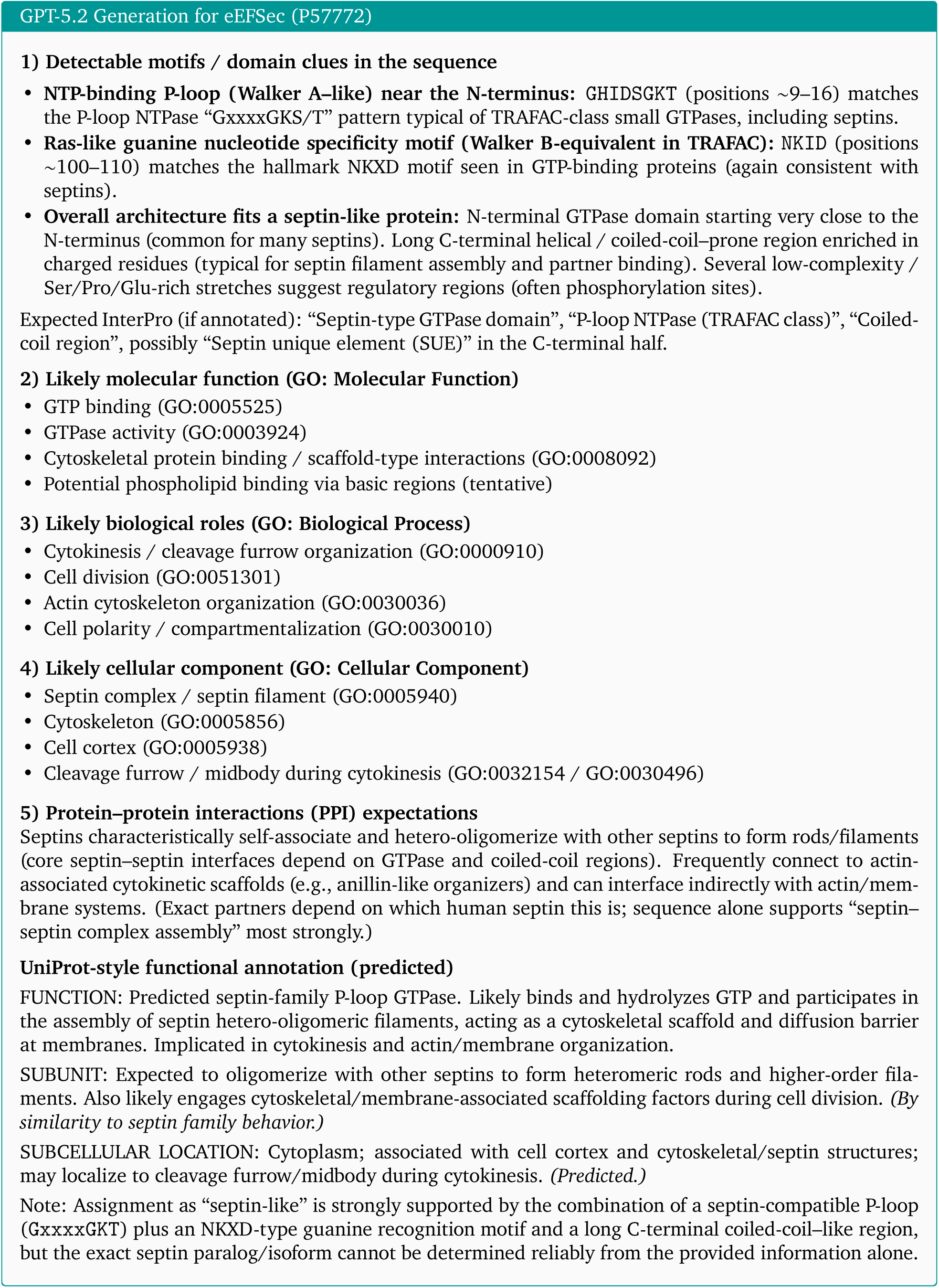

### C.9. CFAP61 (Q8NHU2) Full Reasoning Trace (SFT)

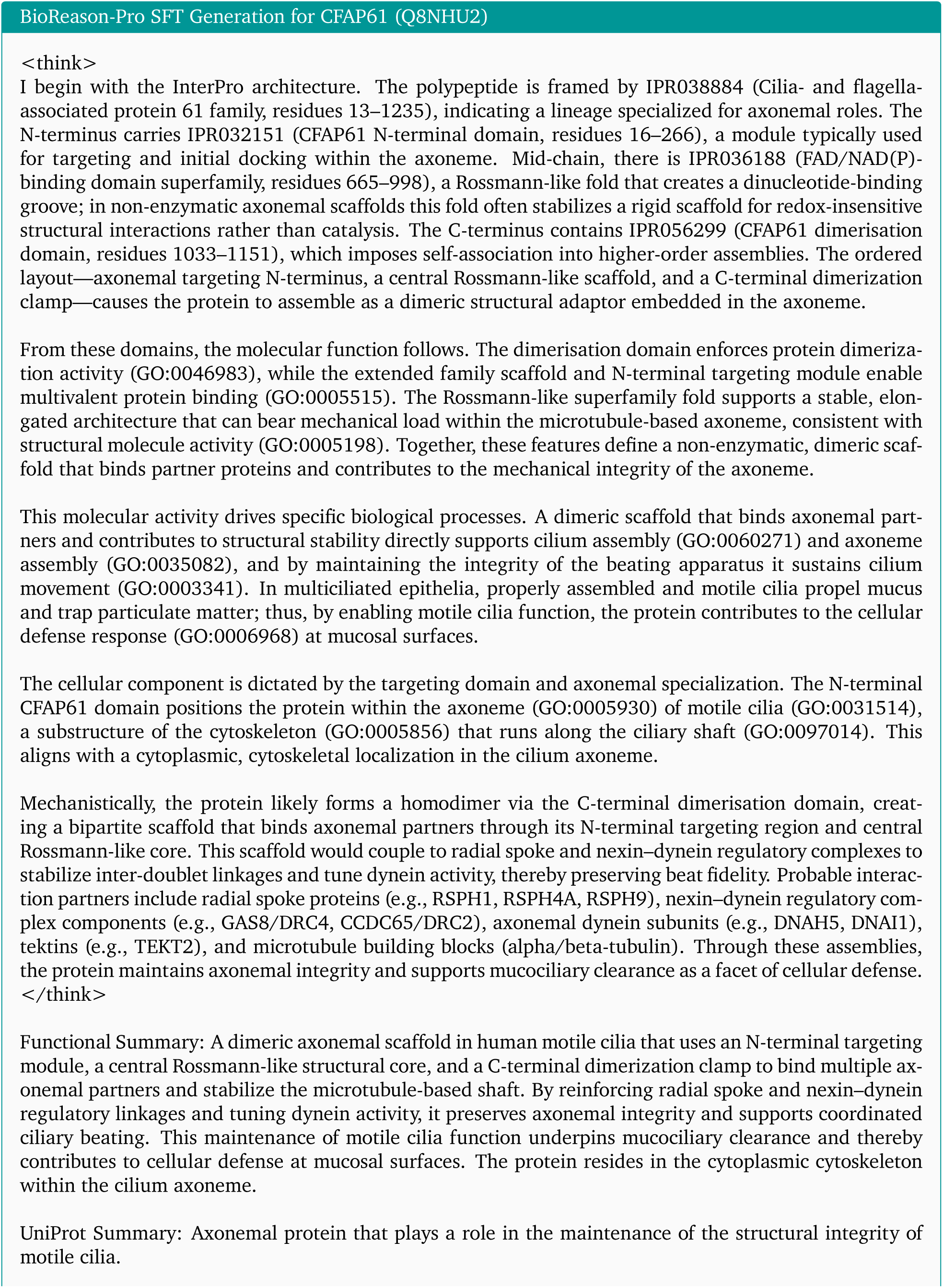
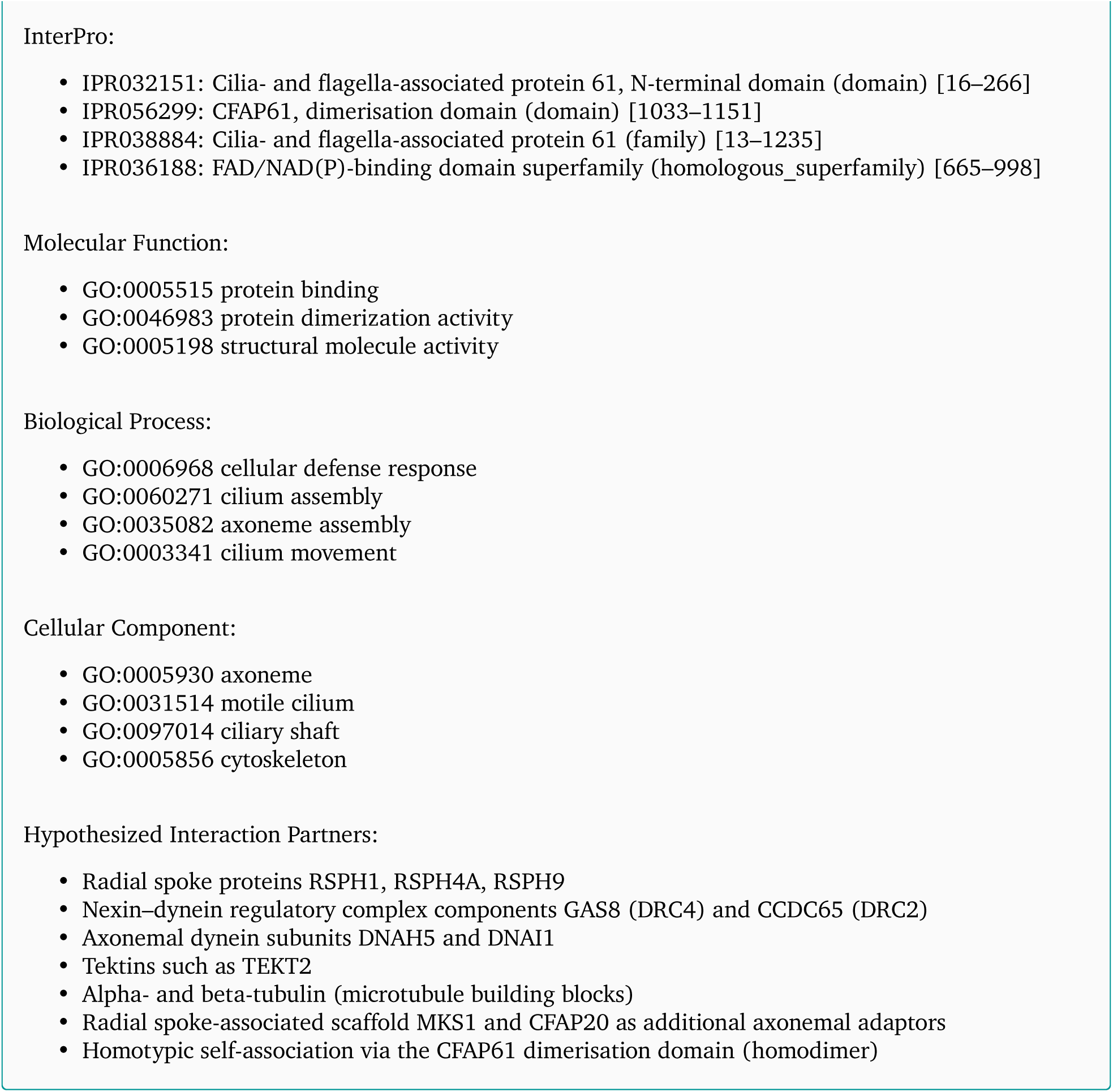

### C.10. CFAP61 (Q8NHU2) Full Reasoning Trace (RL)

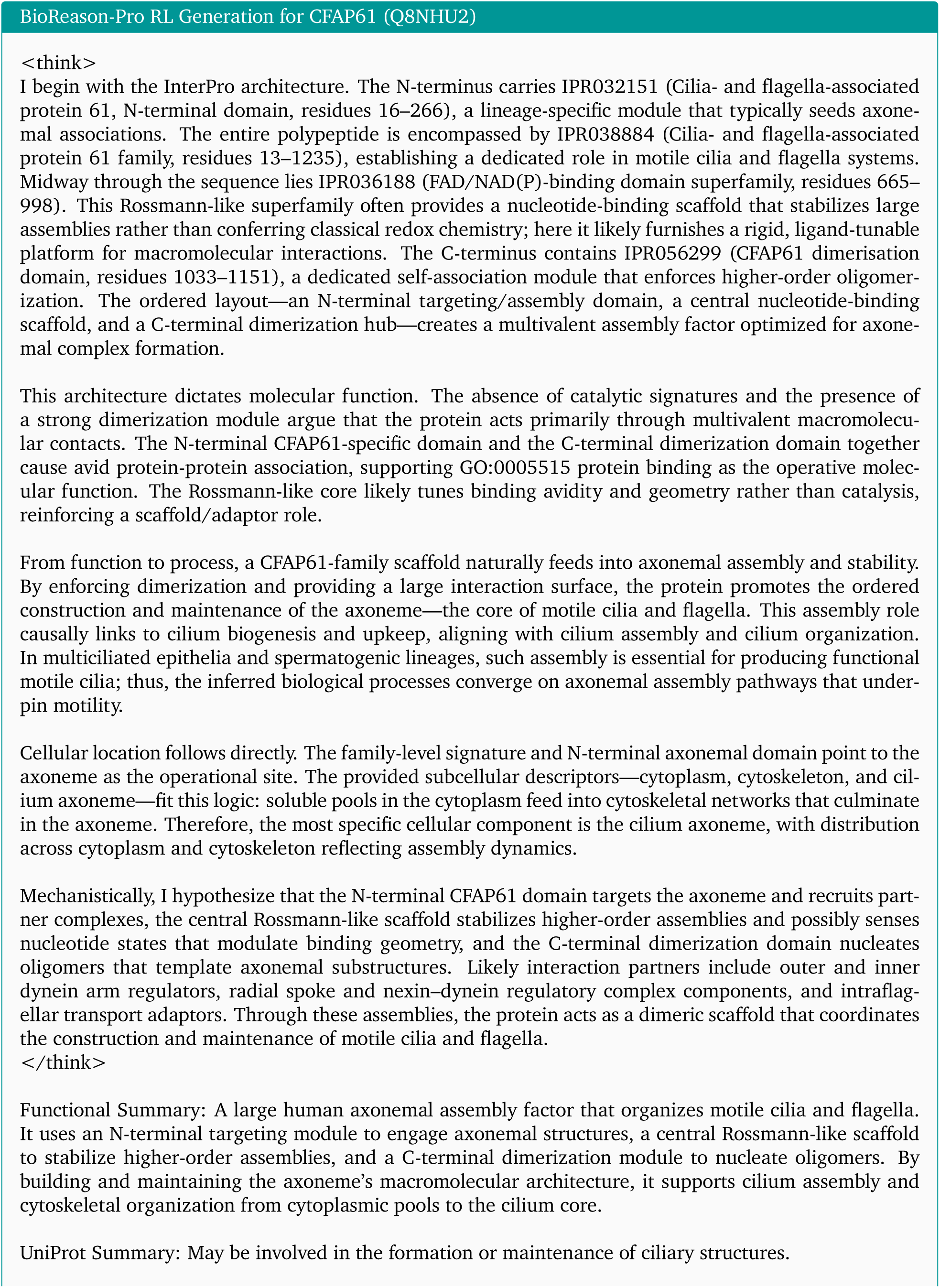
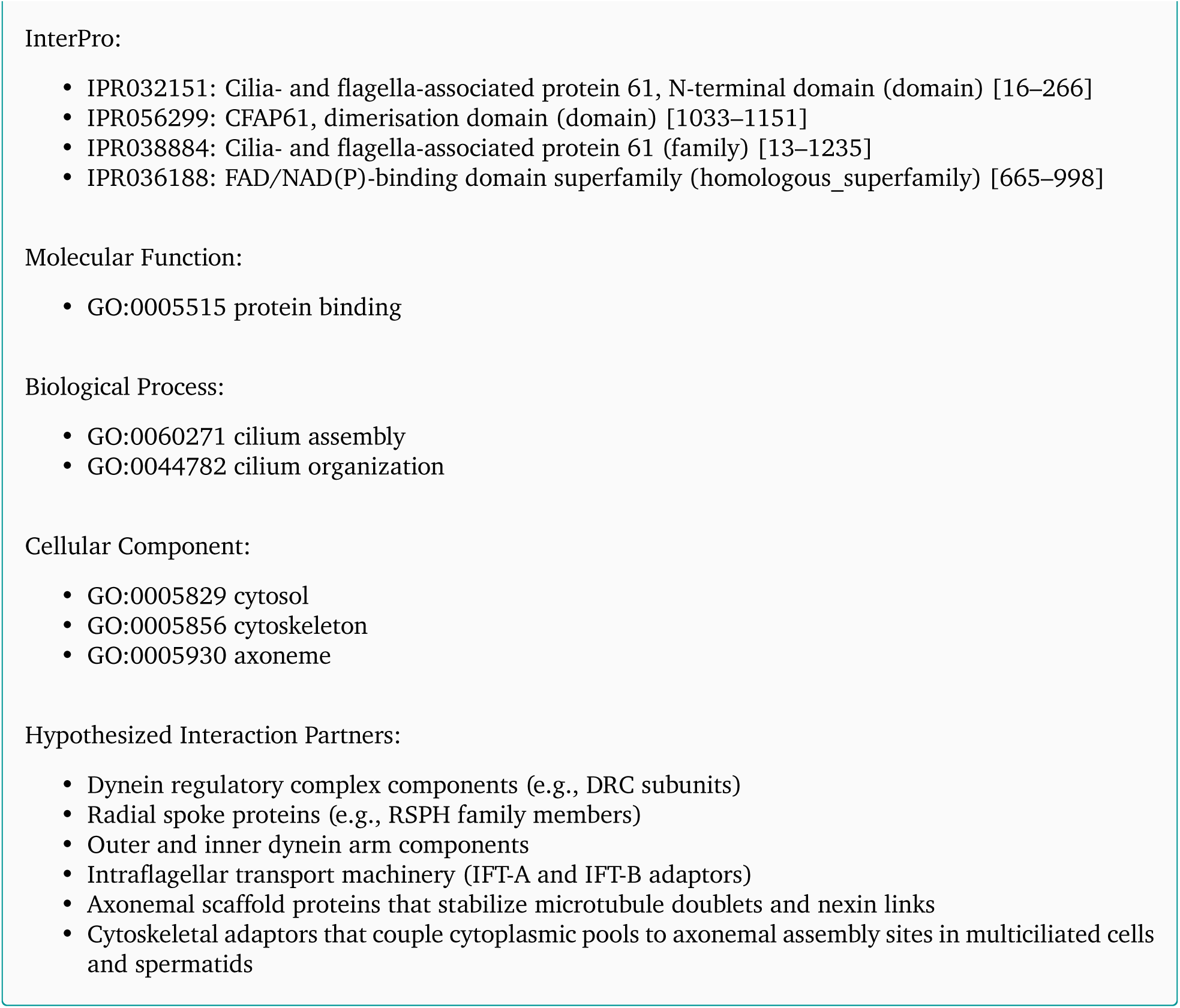

### C.11. CFAP61 (Q8NHU2) Full Reasoning Trace (GPT-5.2 Thinking High)

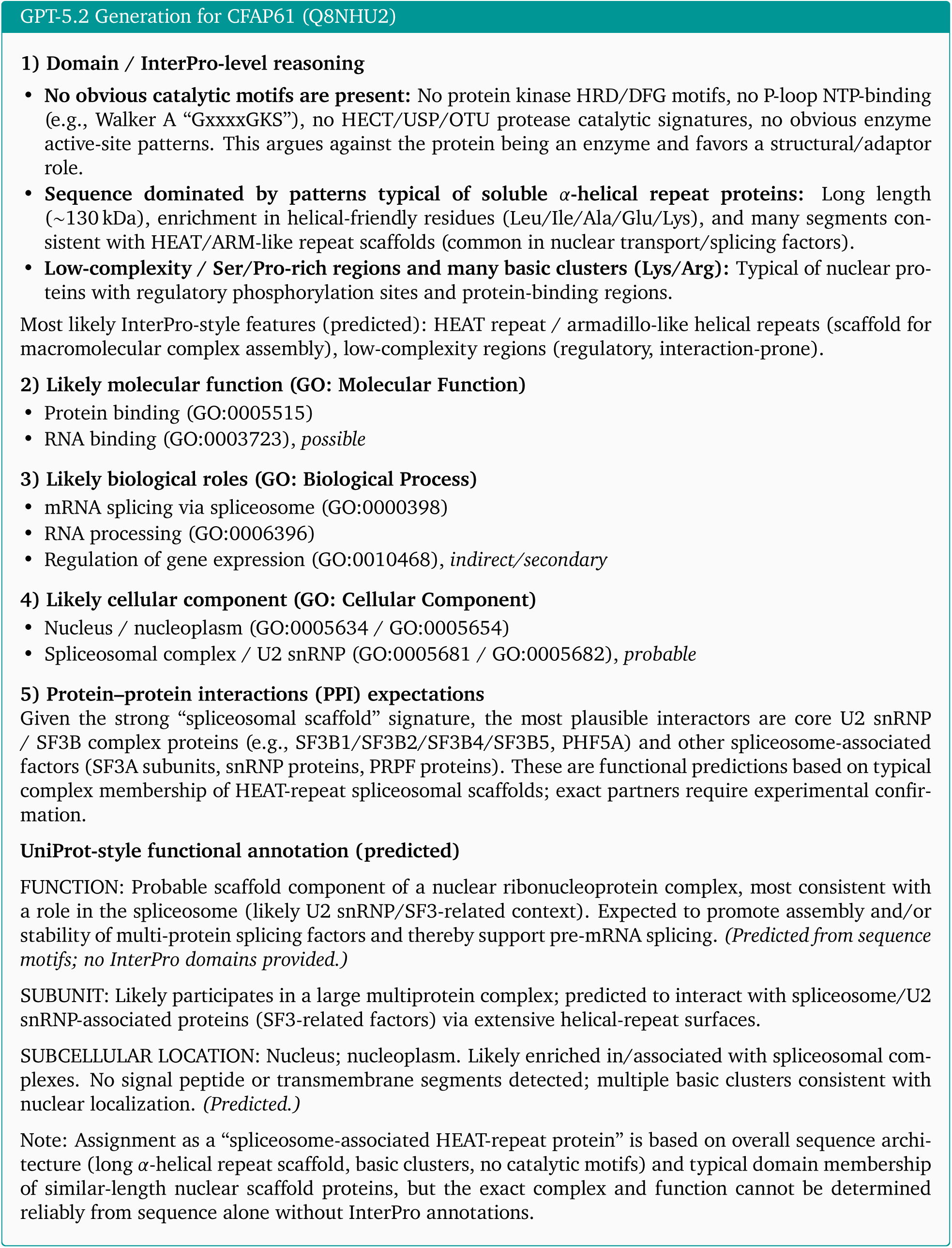

### C.12. EvoAcr1 Full Reasoning Traces (SFT)

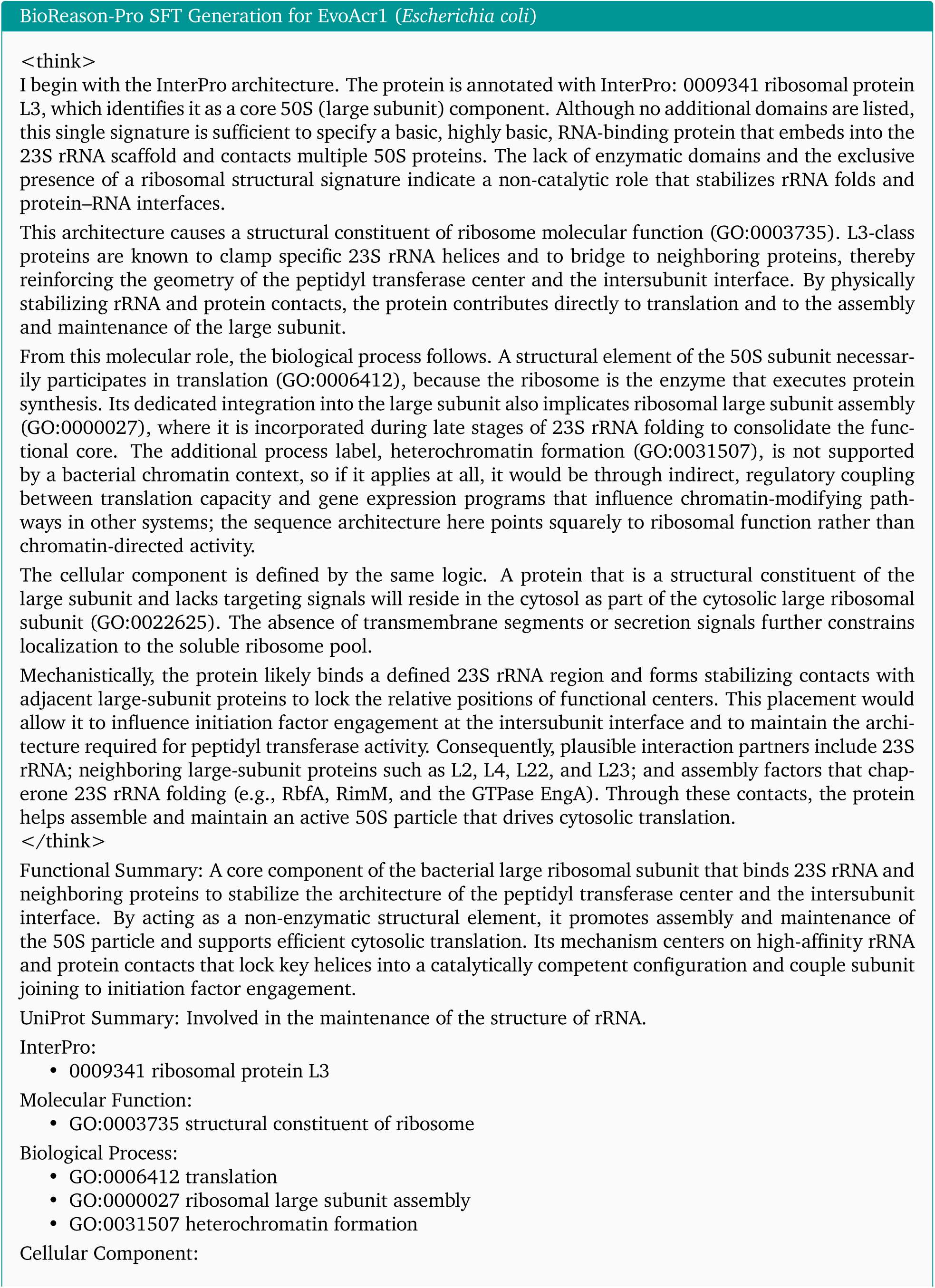
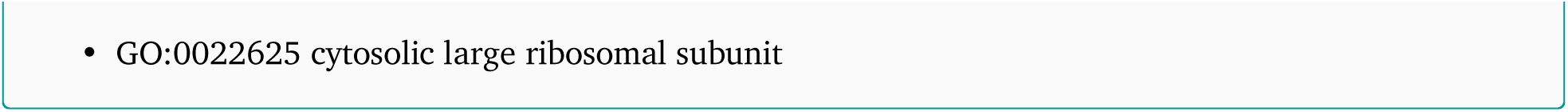

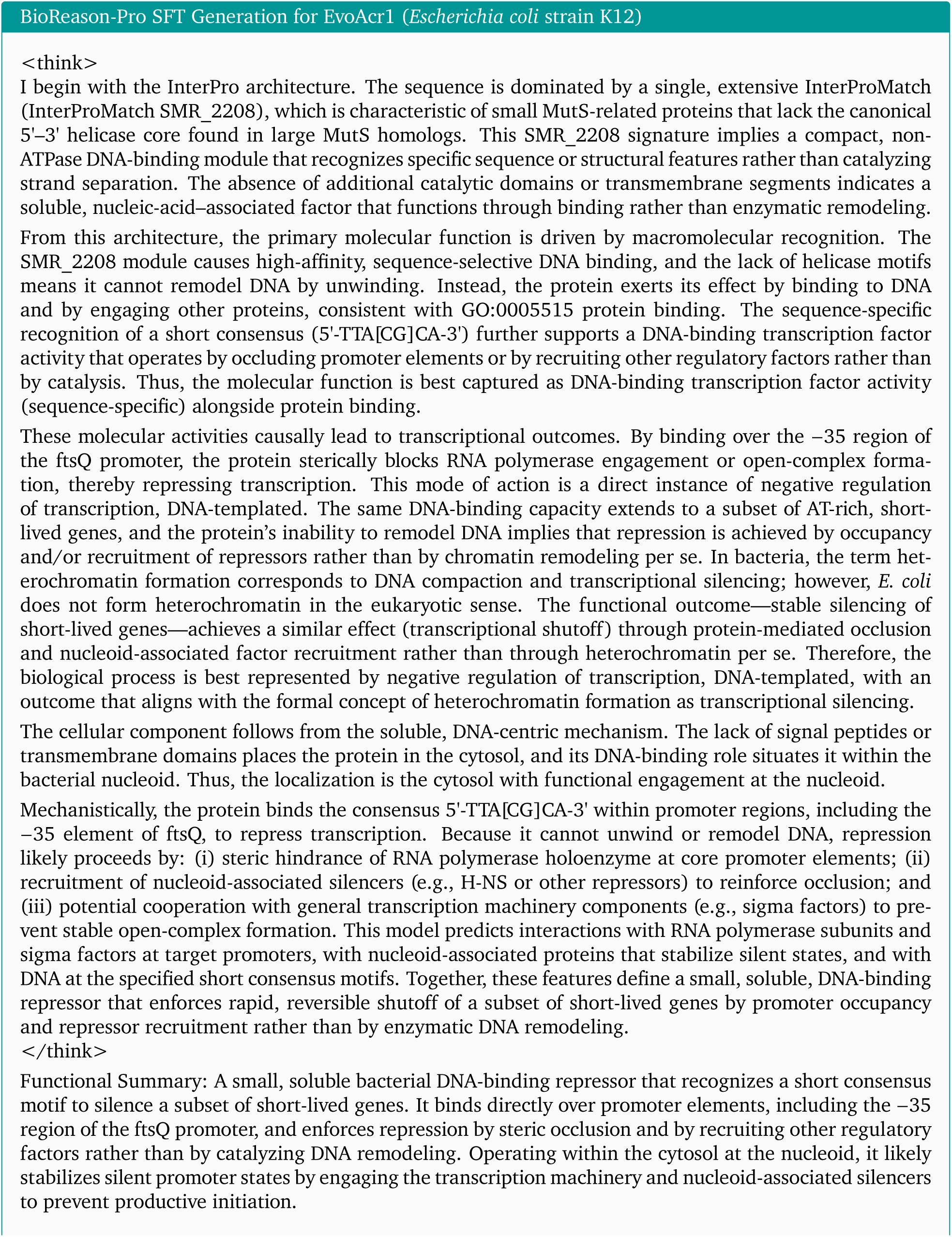
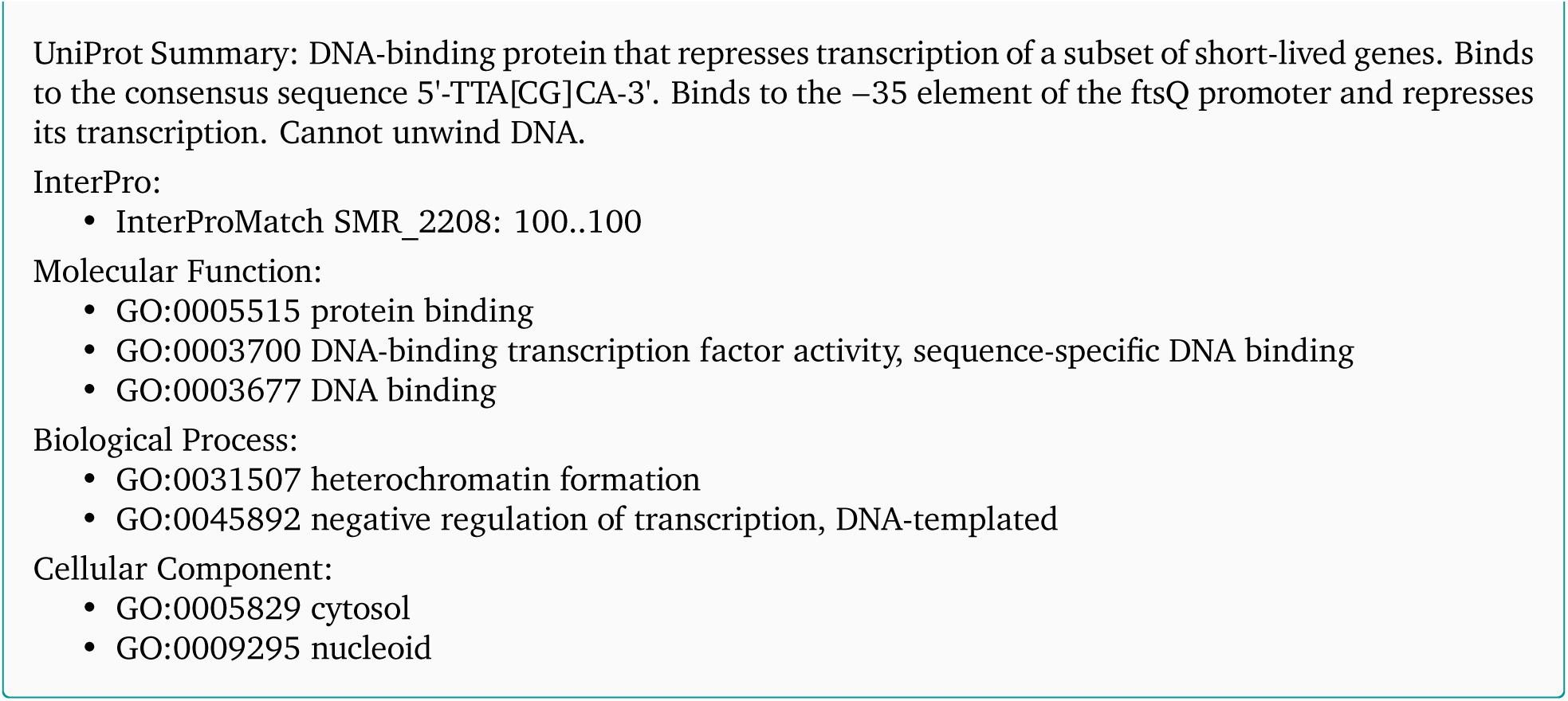

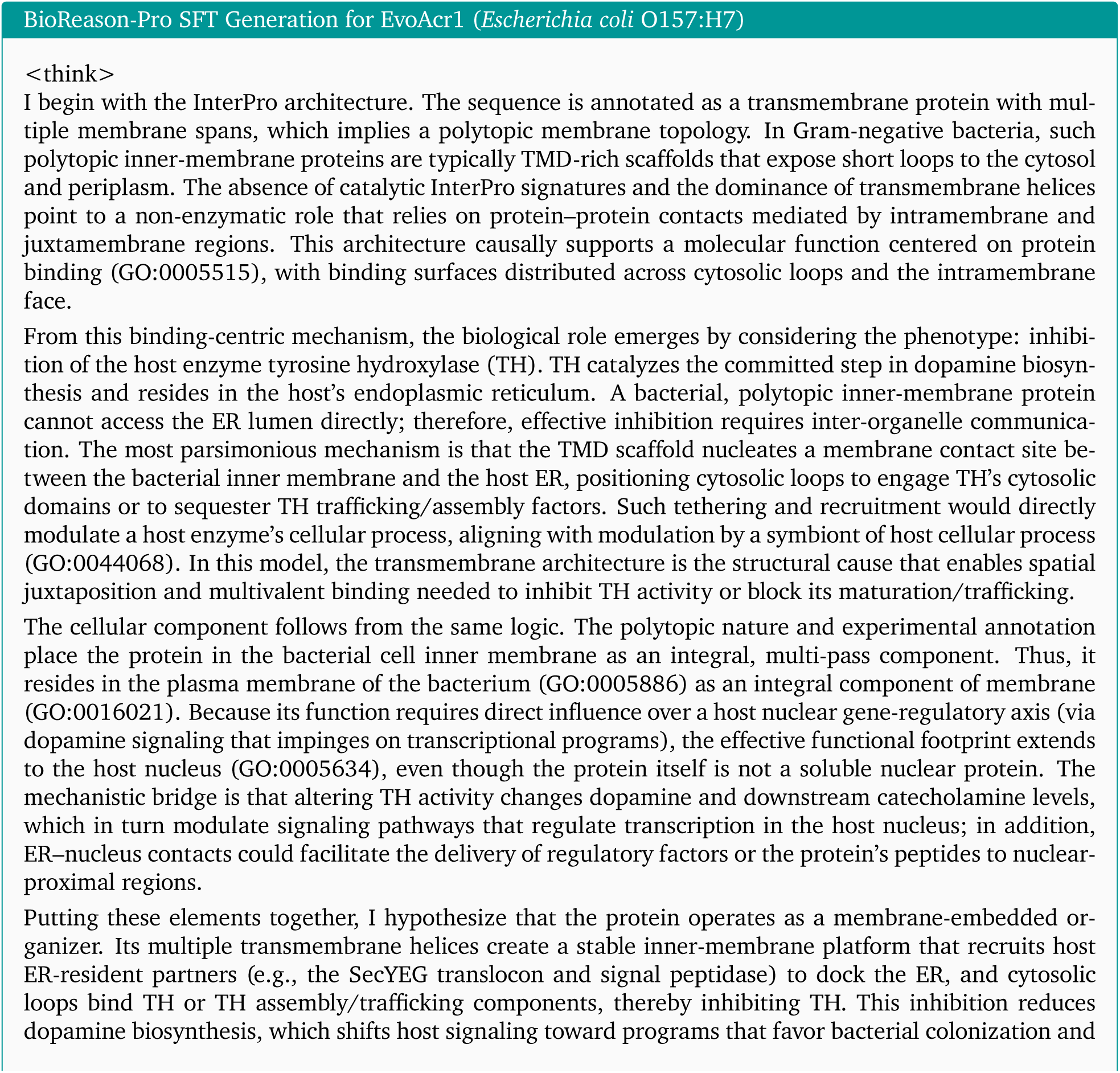
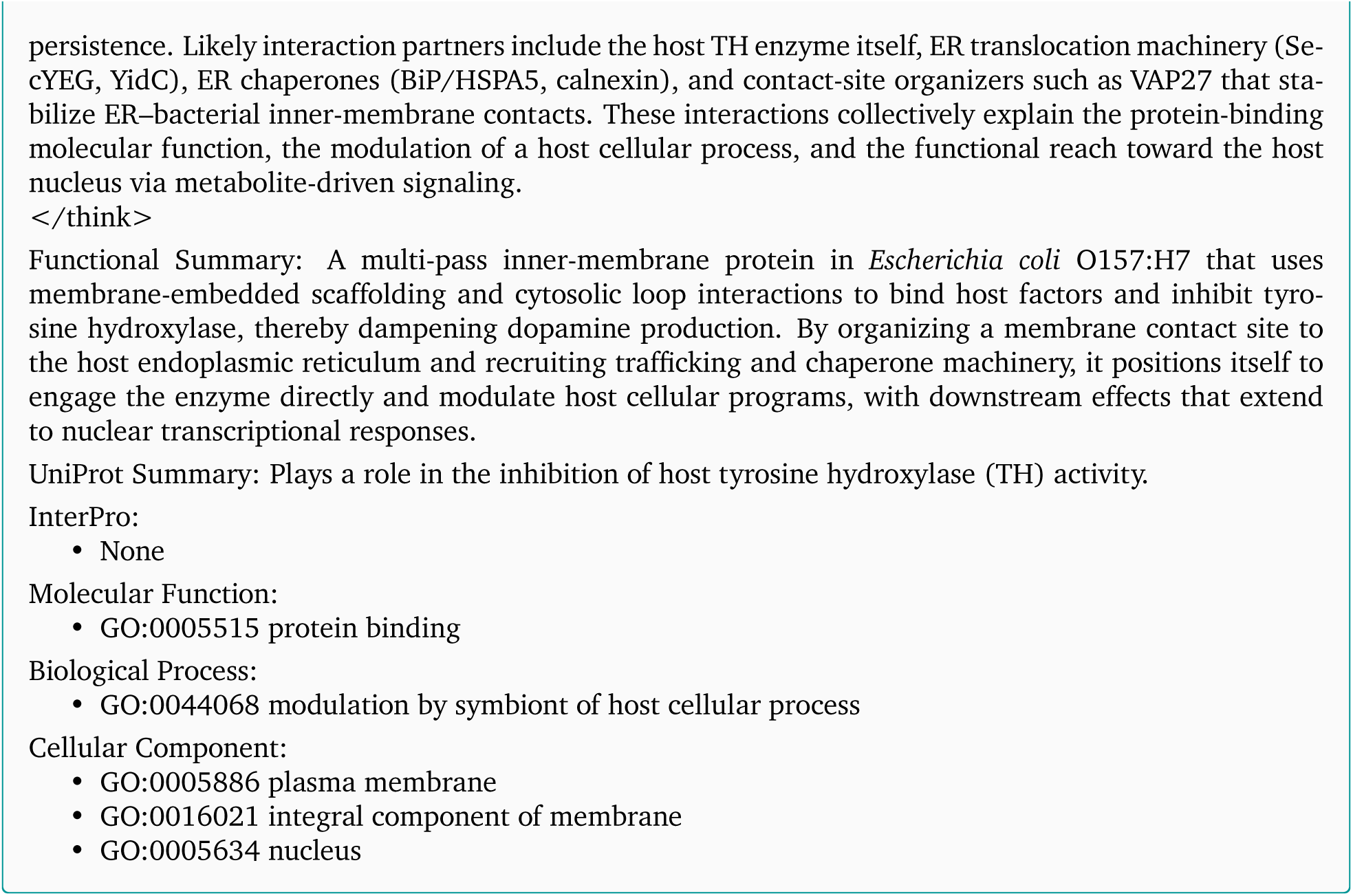

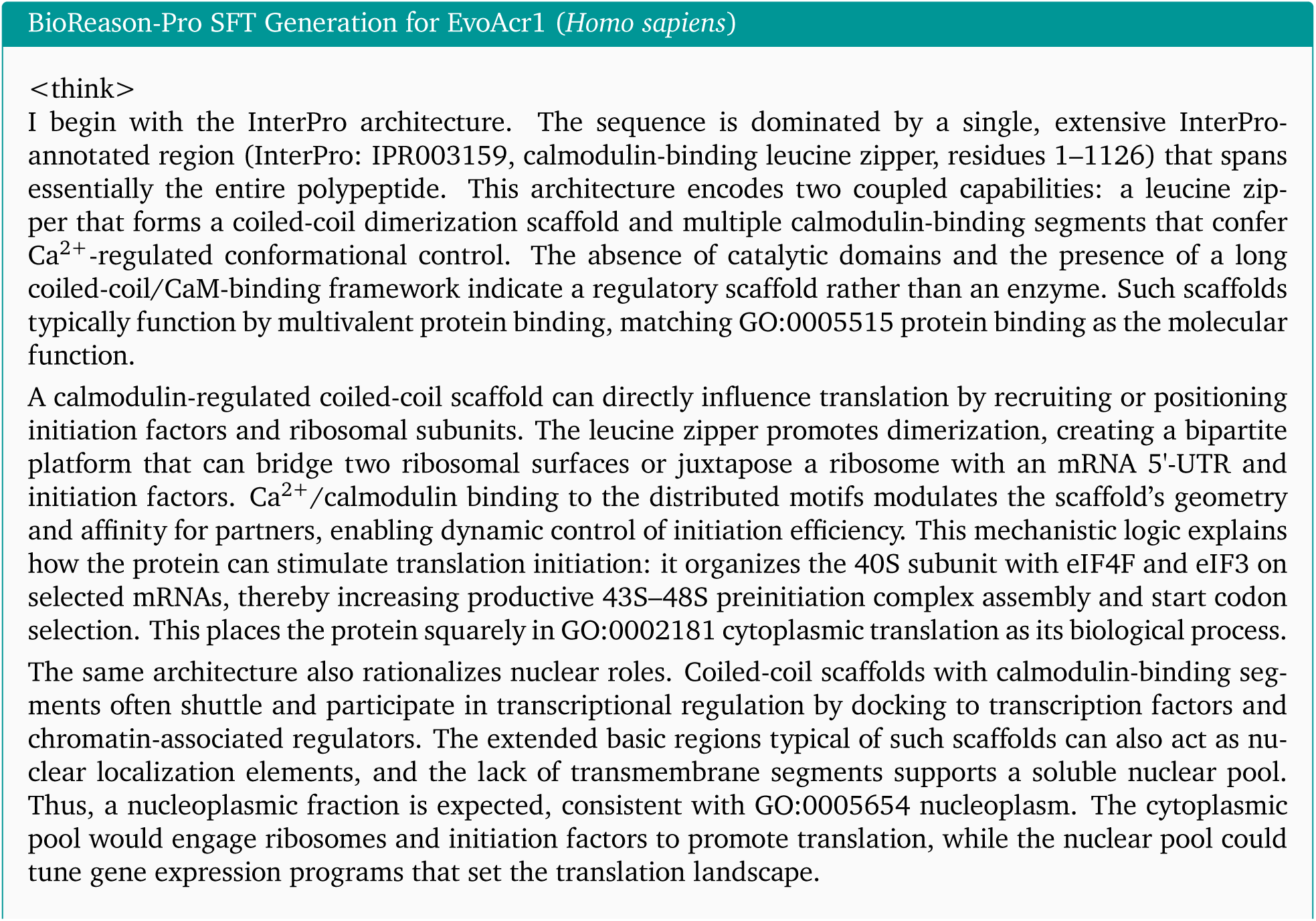
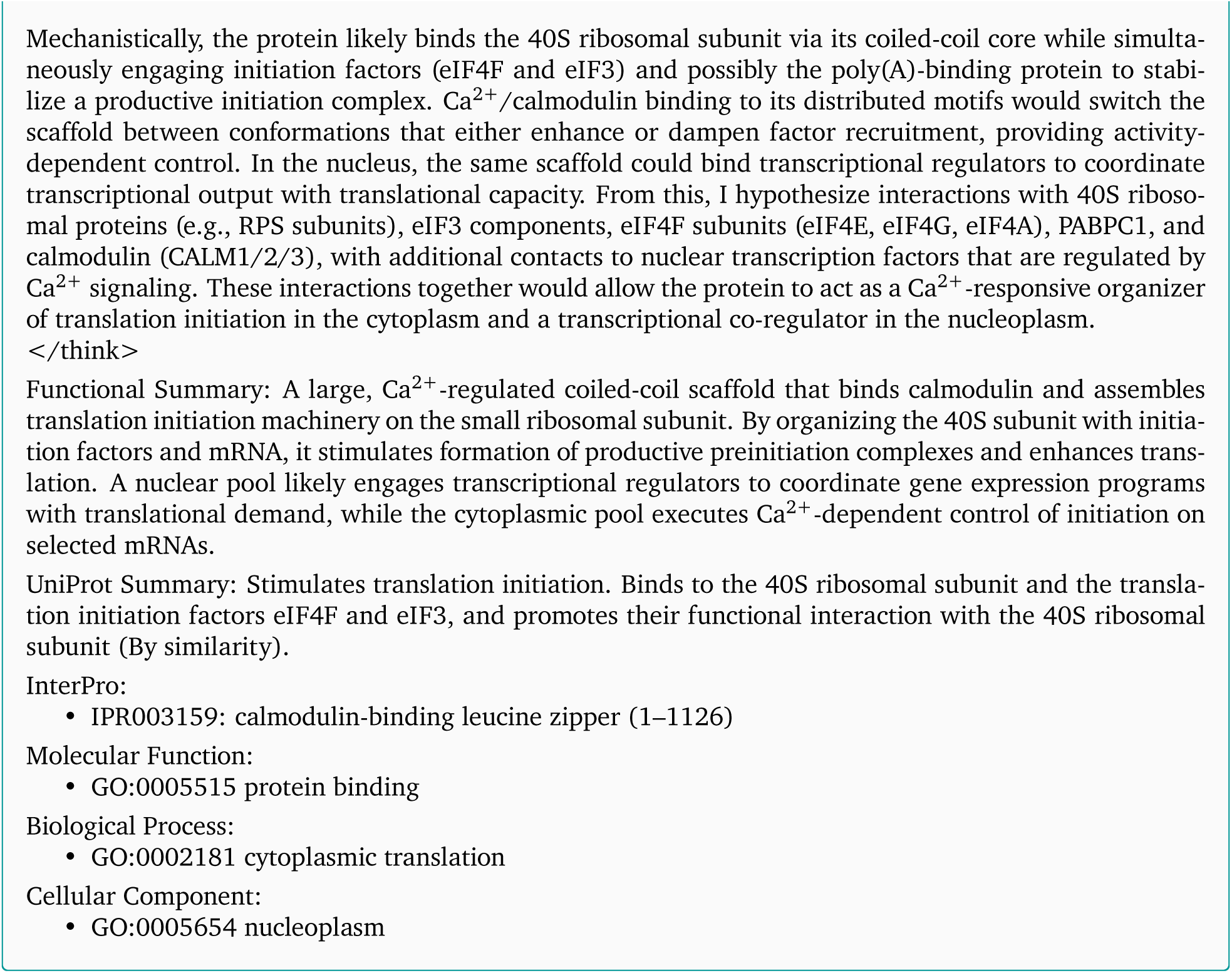

### C.13. RCDG1 Full Prediction (*Homo sapiens*)

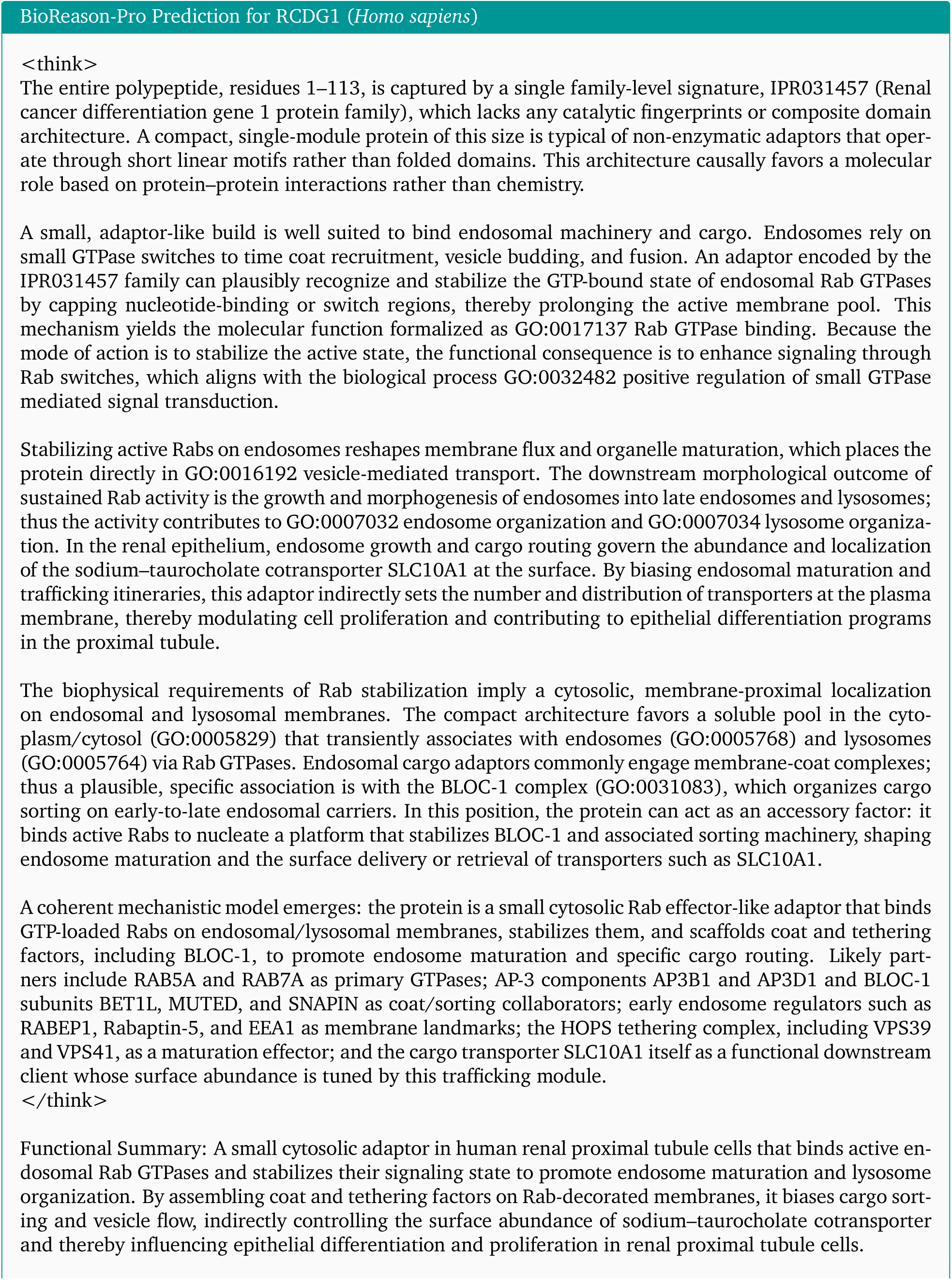
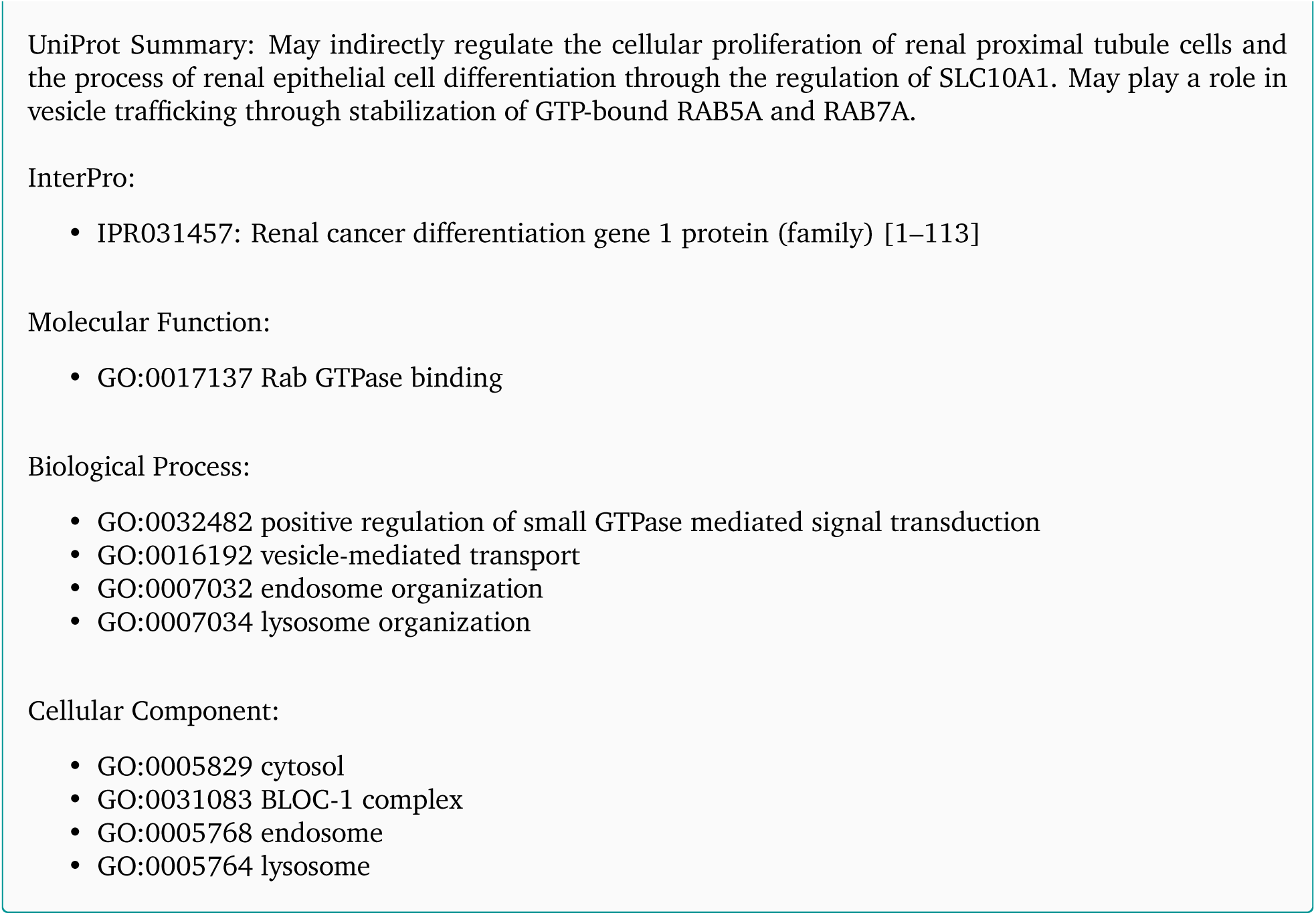

### C.14. EvoAcr1 Full Reasoning Traces (RL)

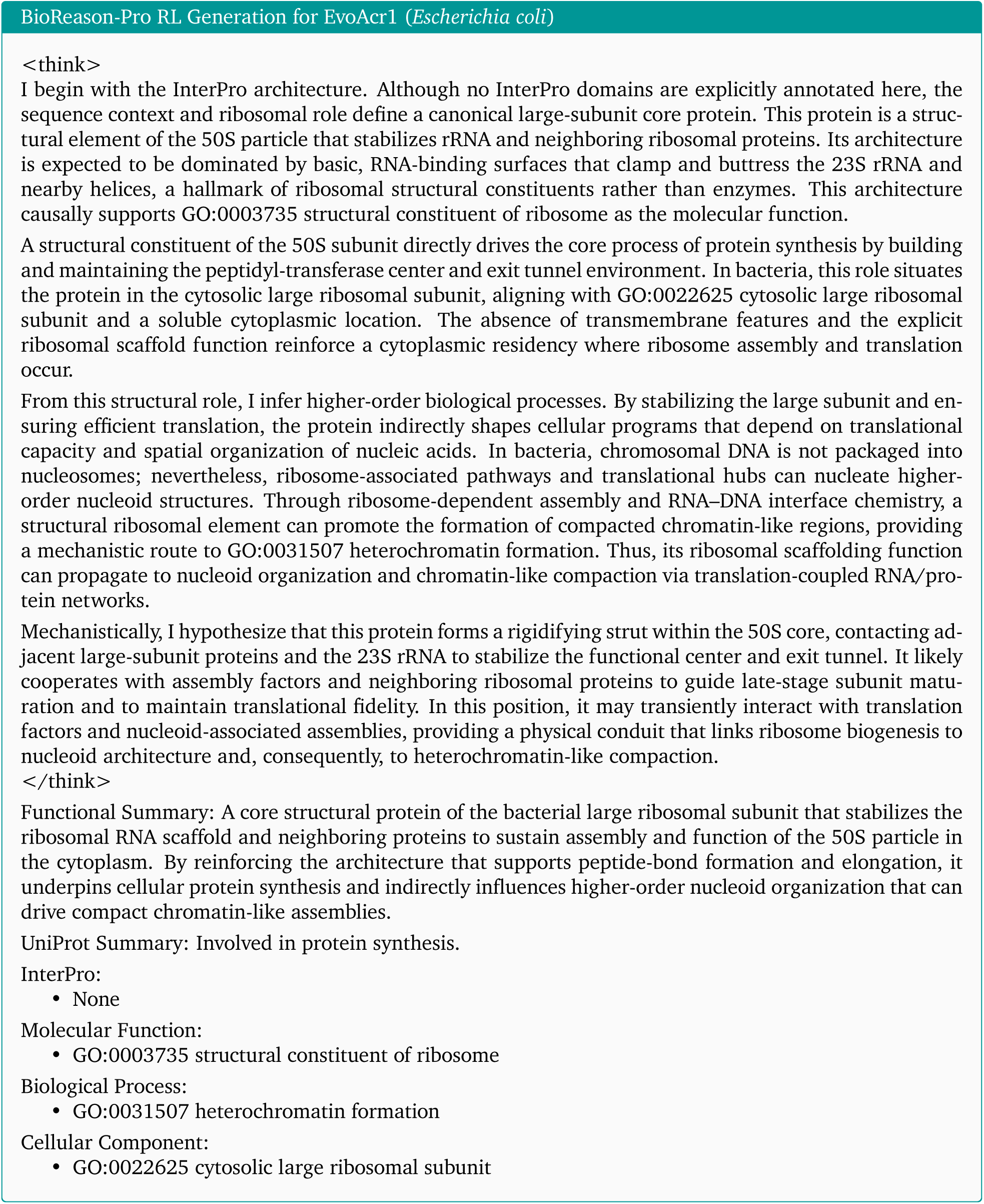
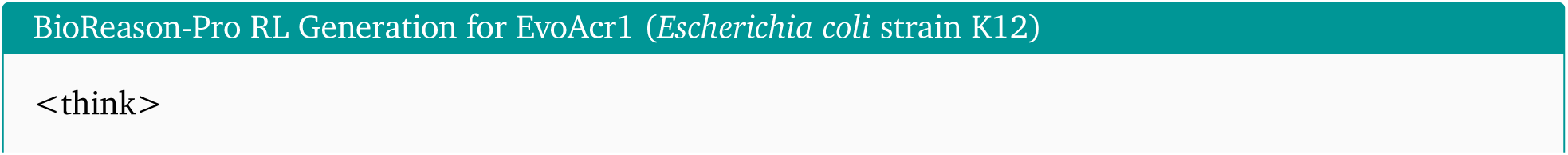
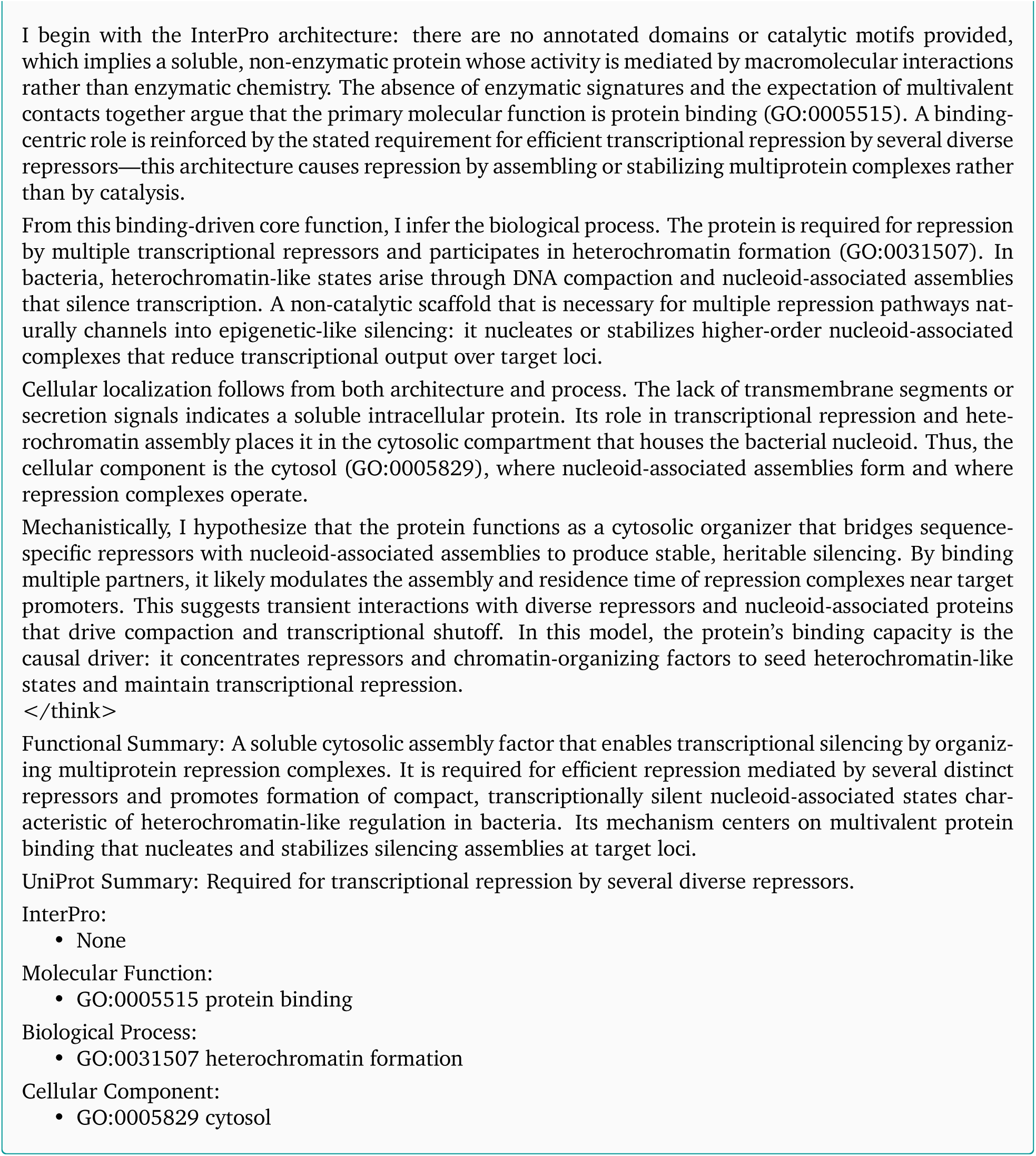
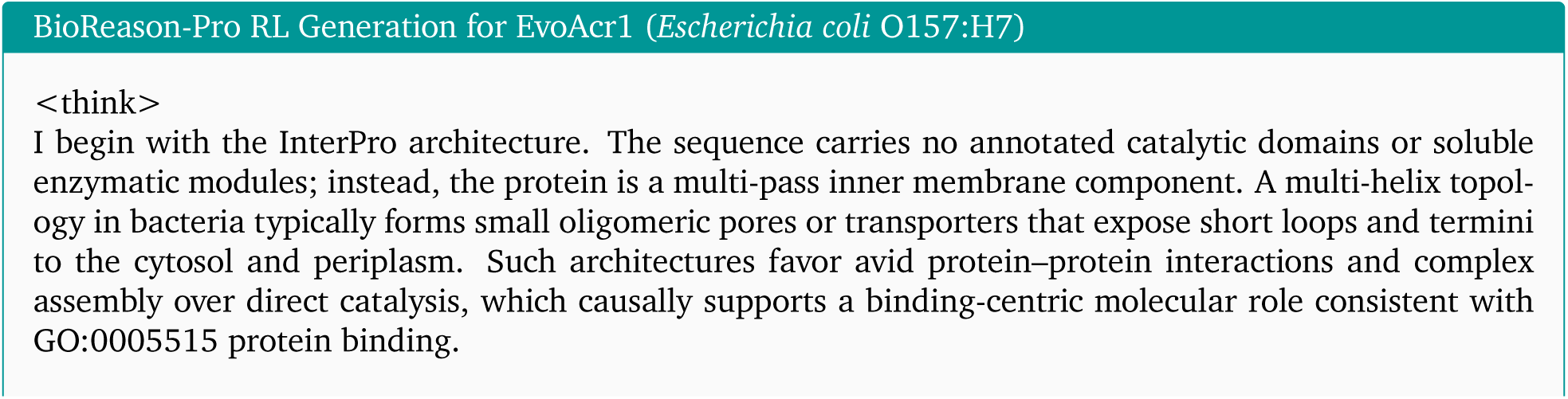
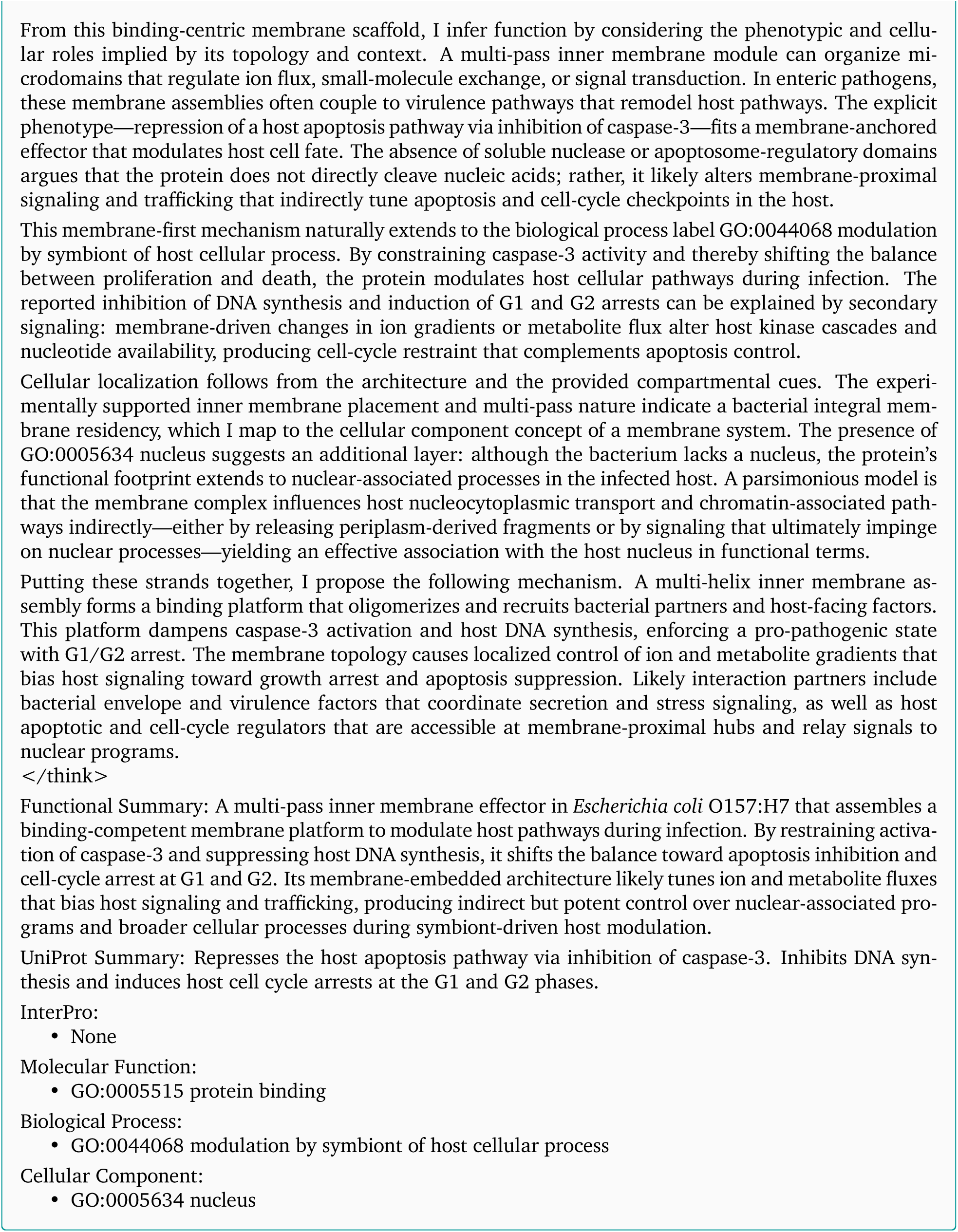
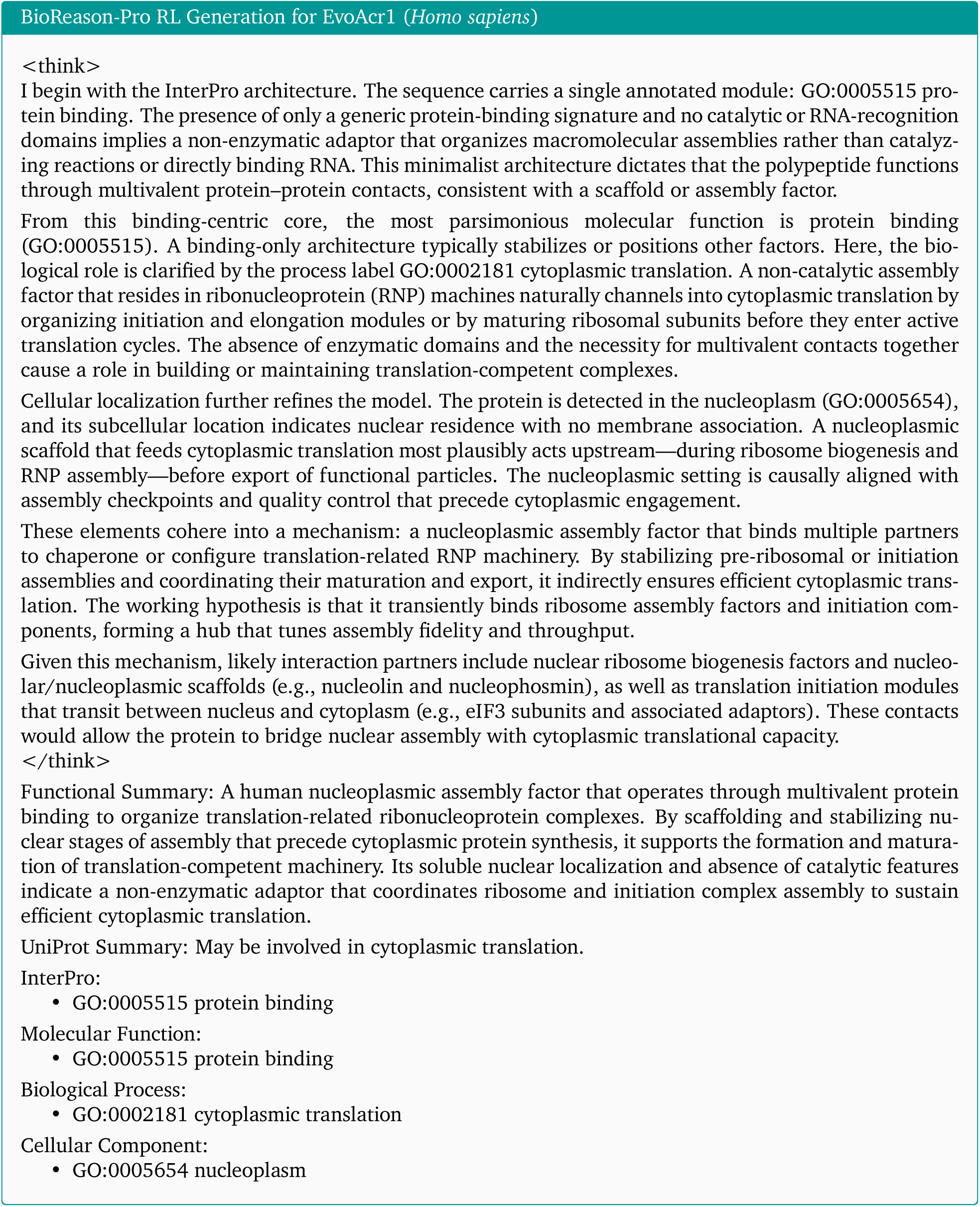

### C.15. EvoAcr2 Full Reasoning Traces (SFT)

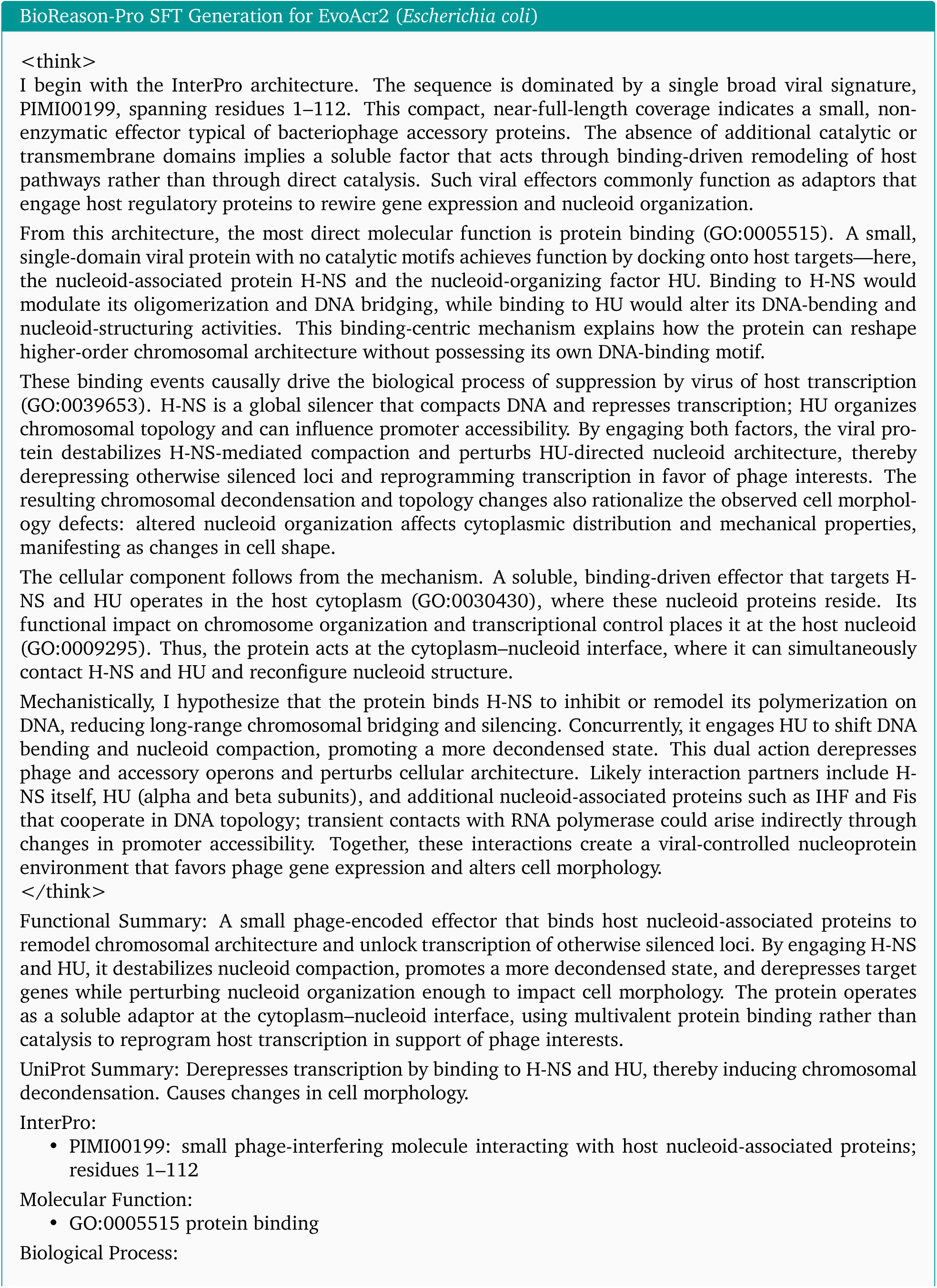
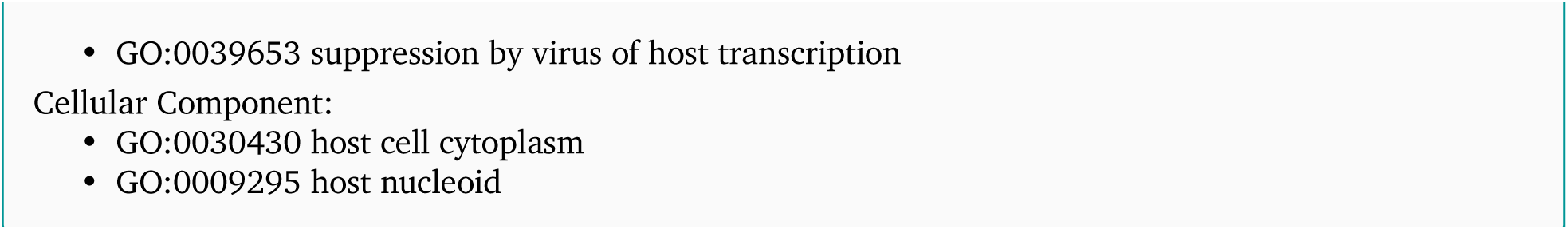
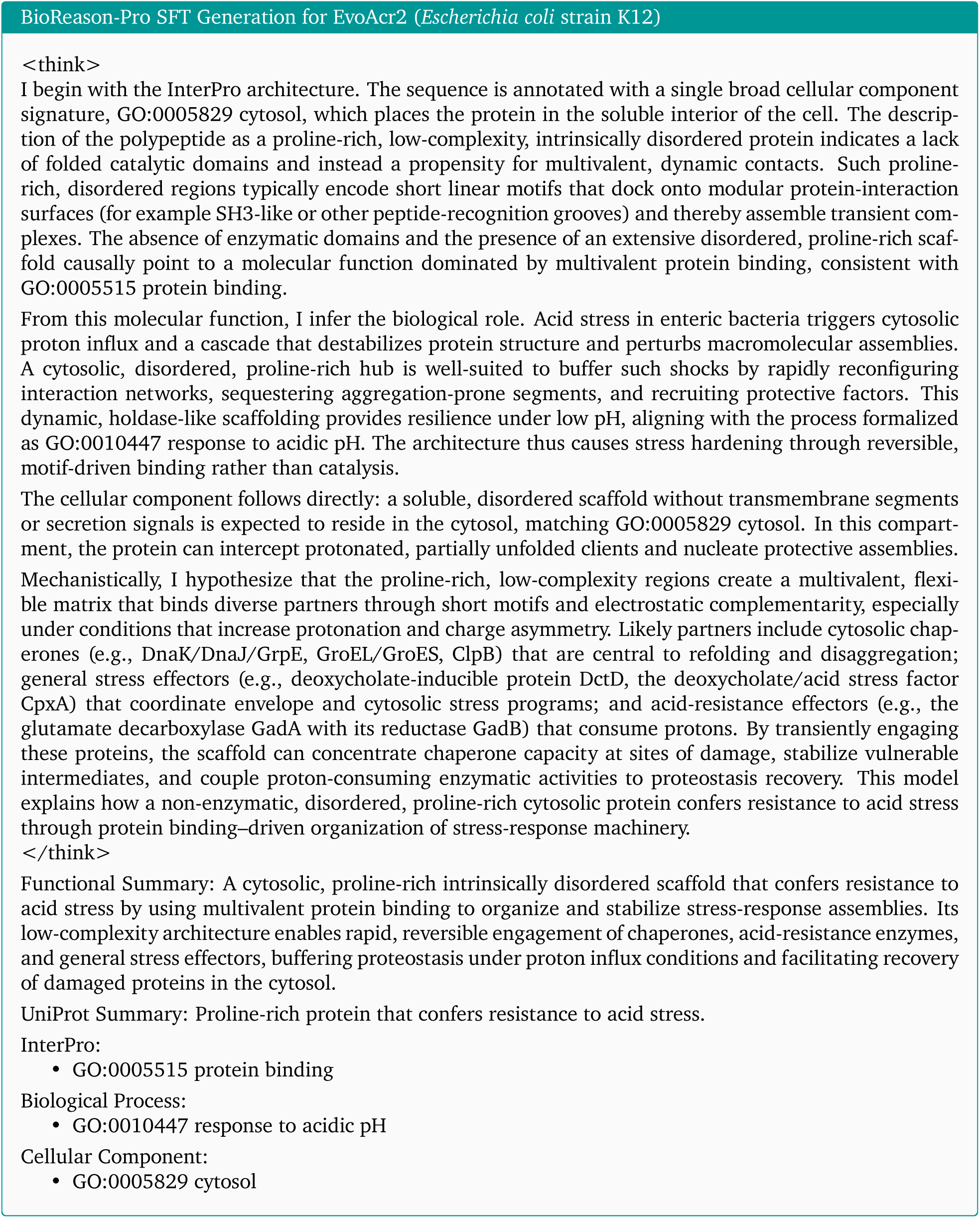
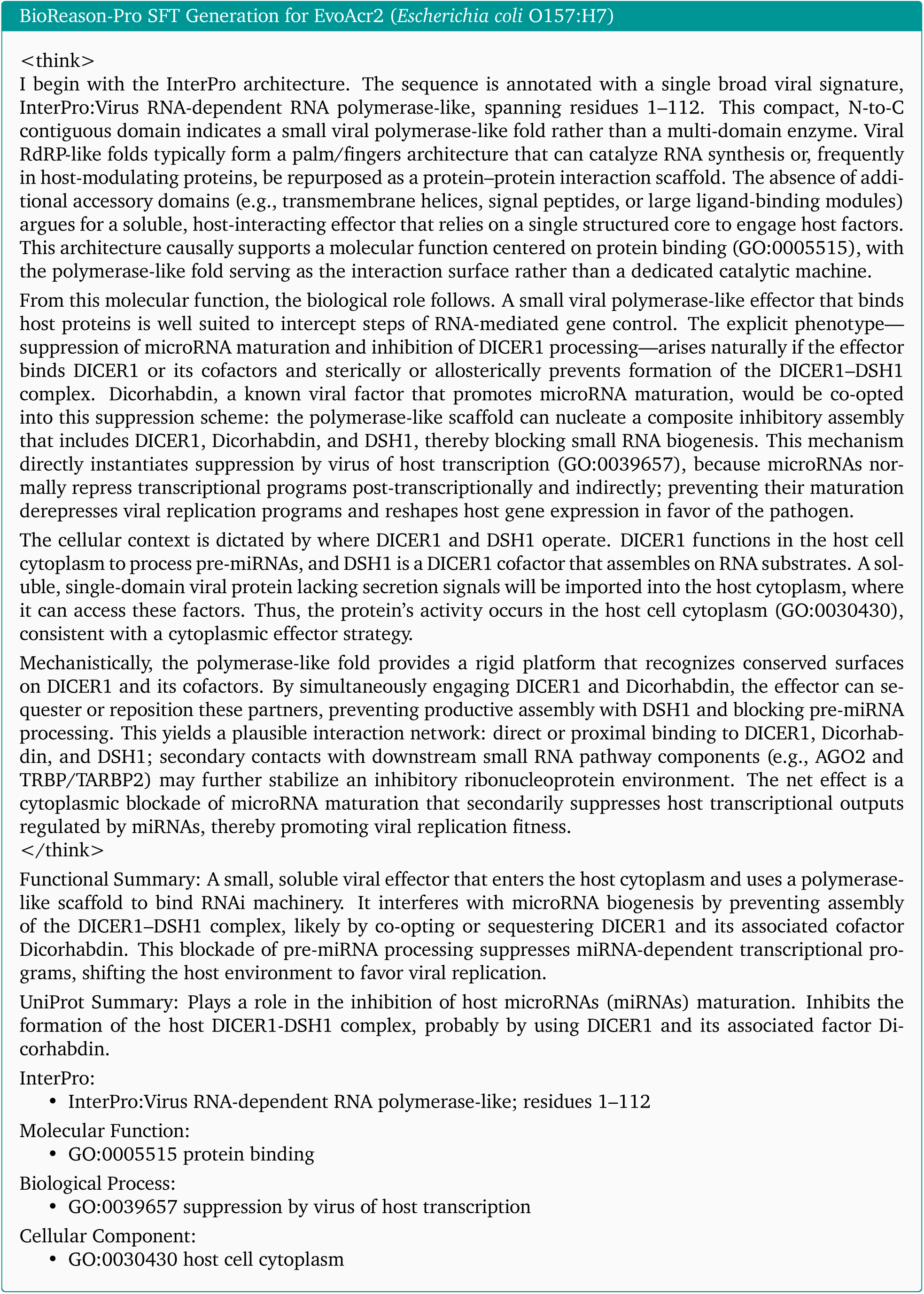
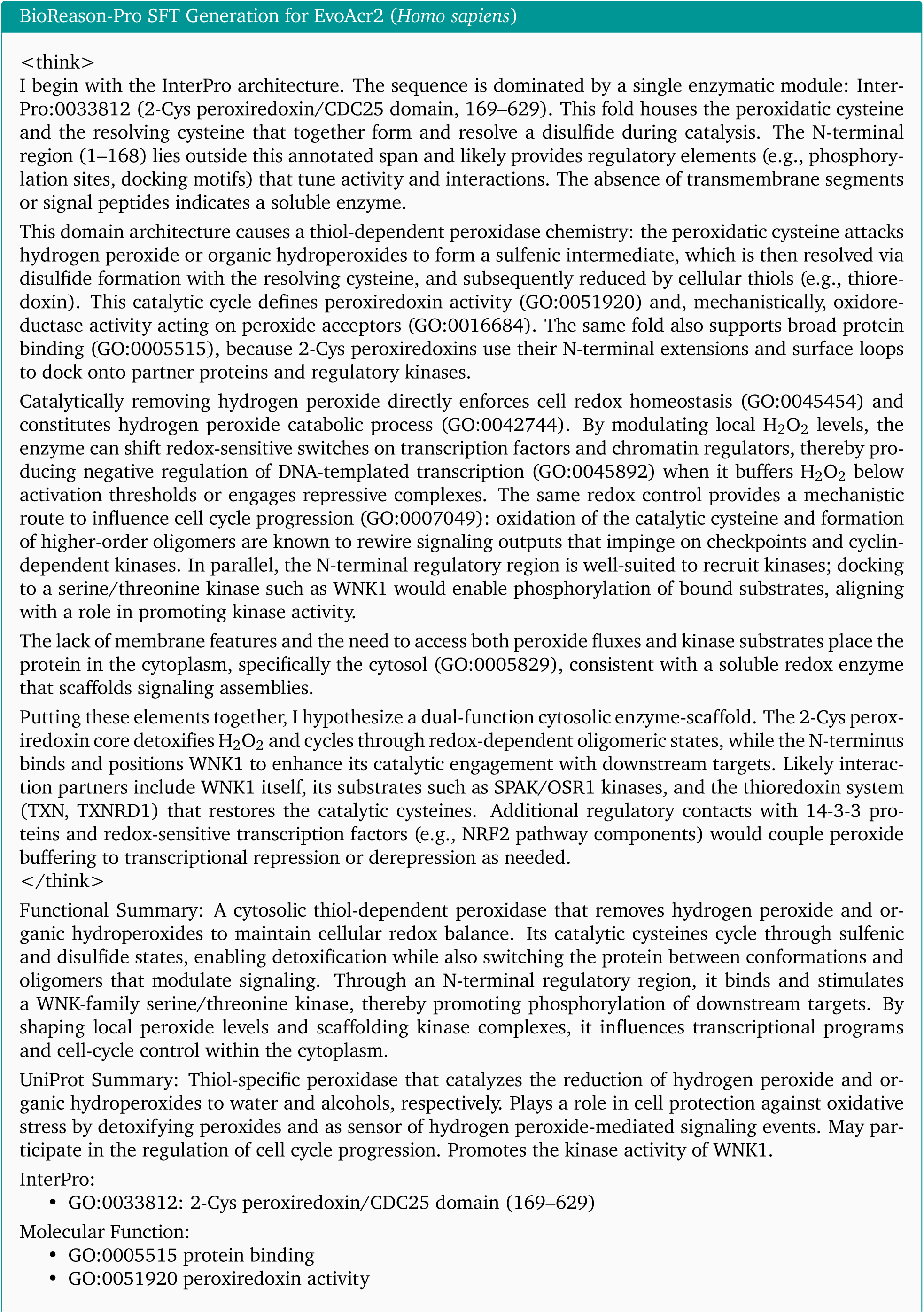
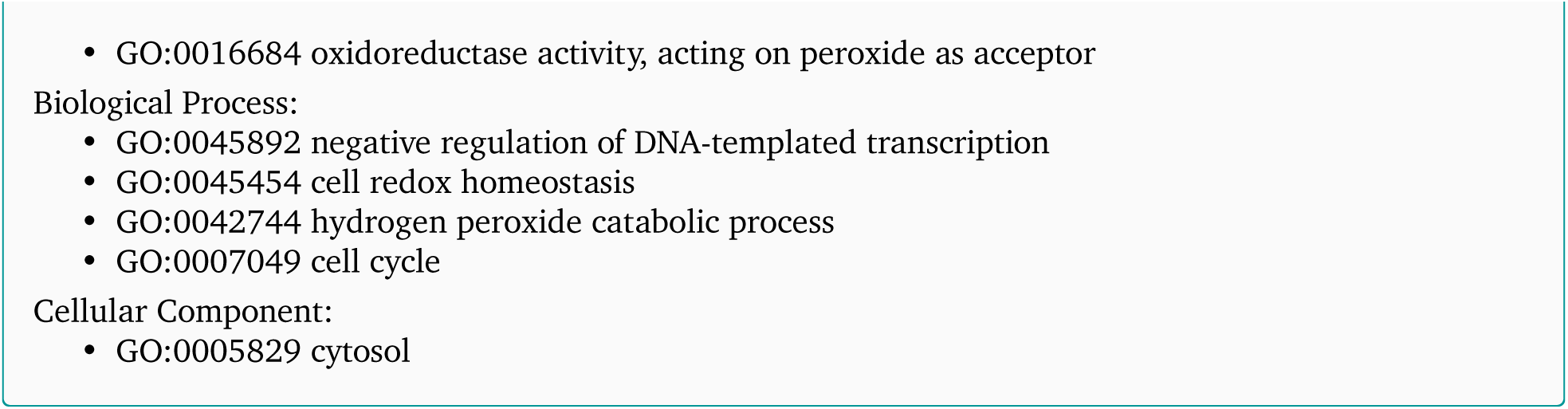

### C.16. EvoAcr2 Full Reasoning Traces (RL)

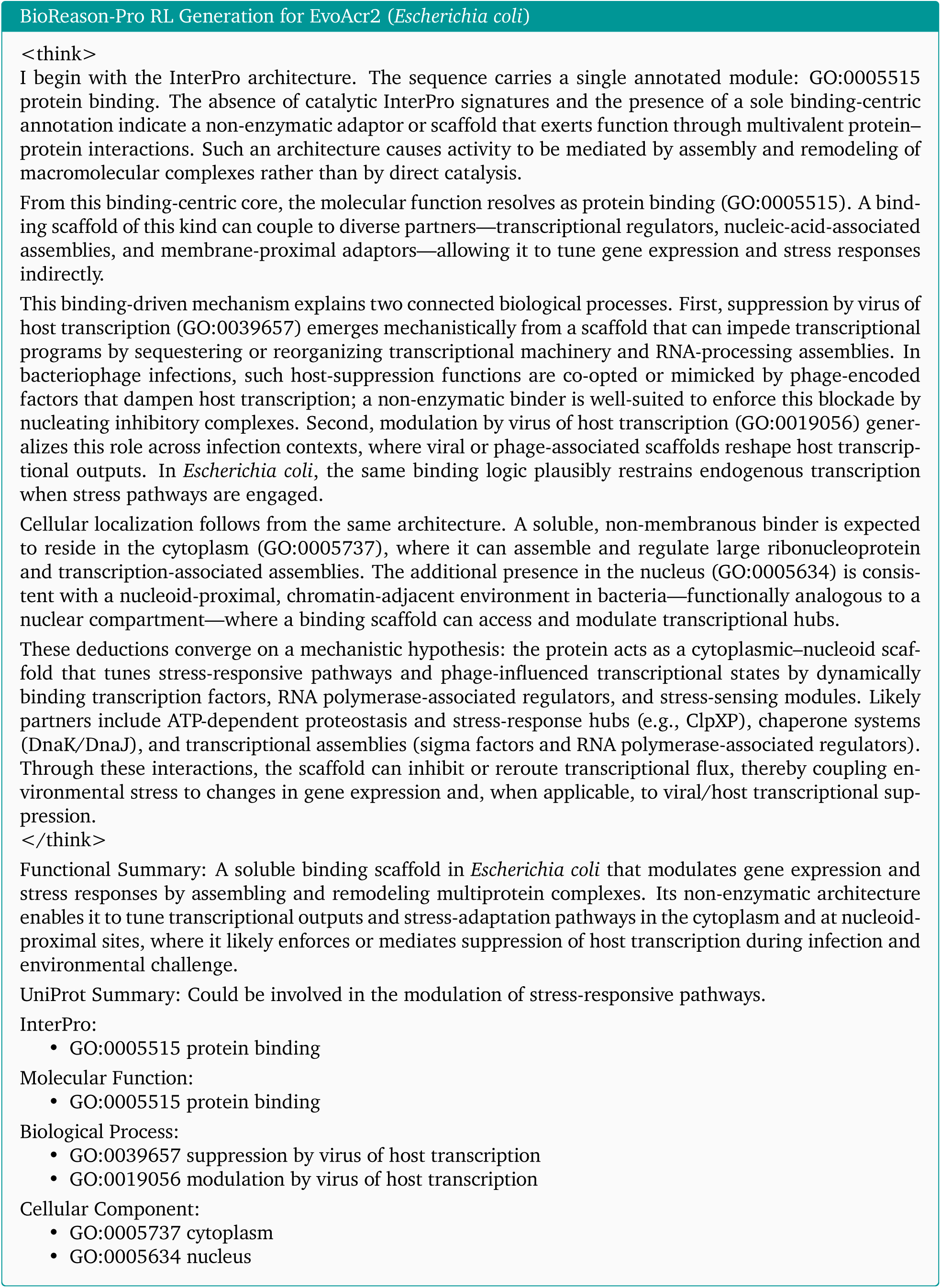
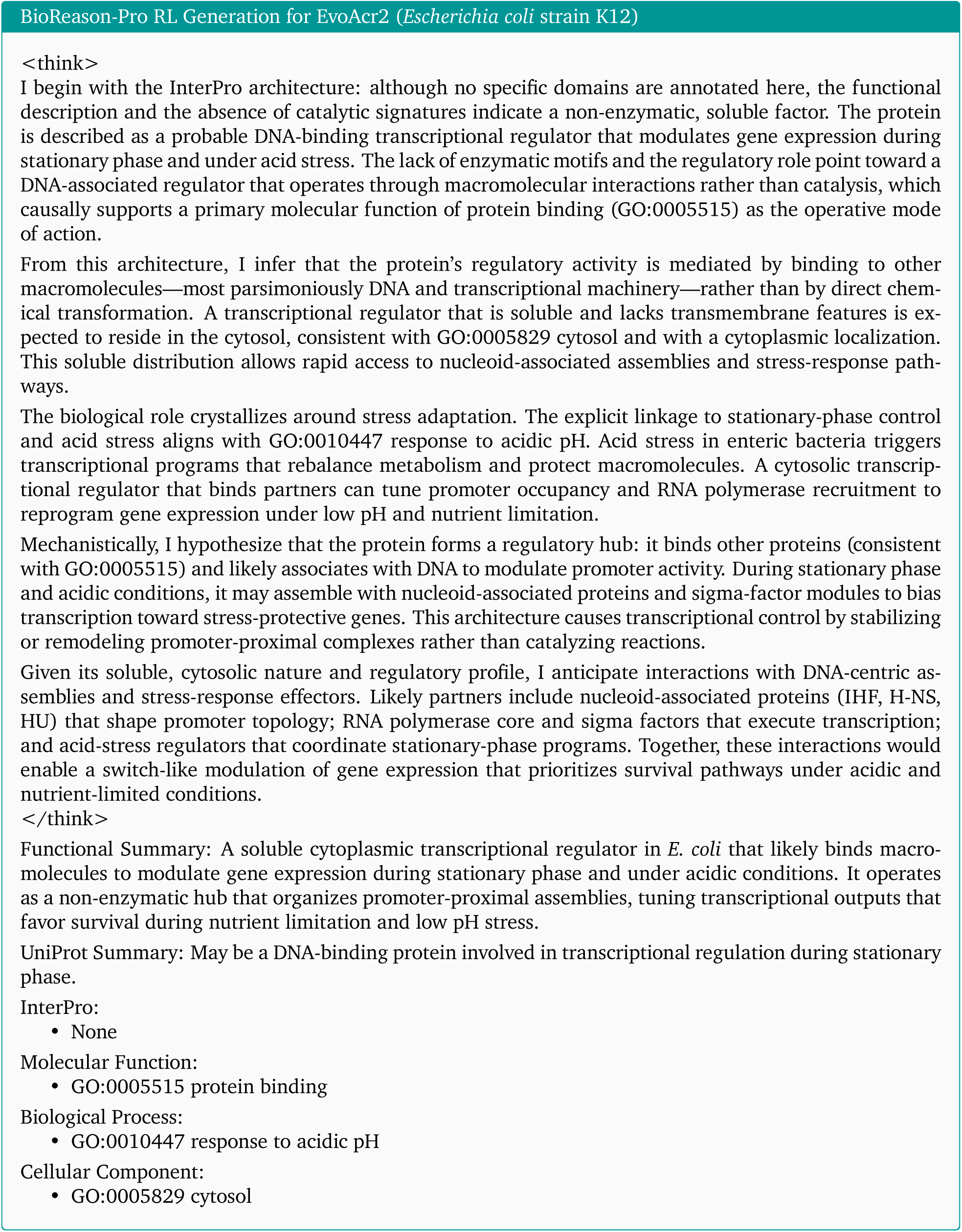
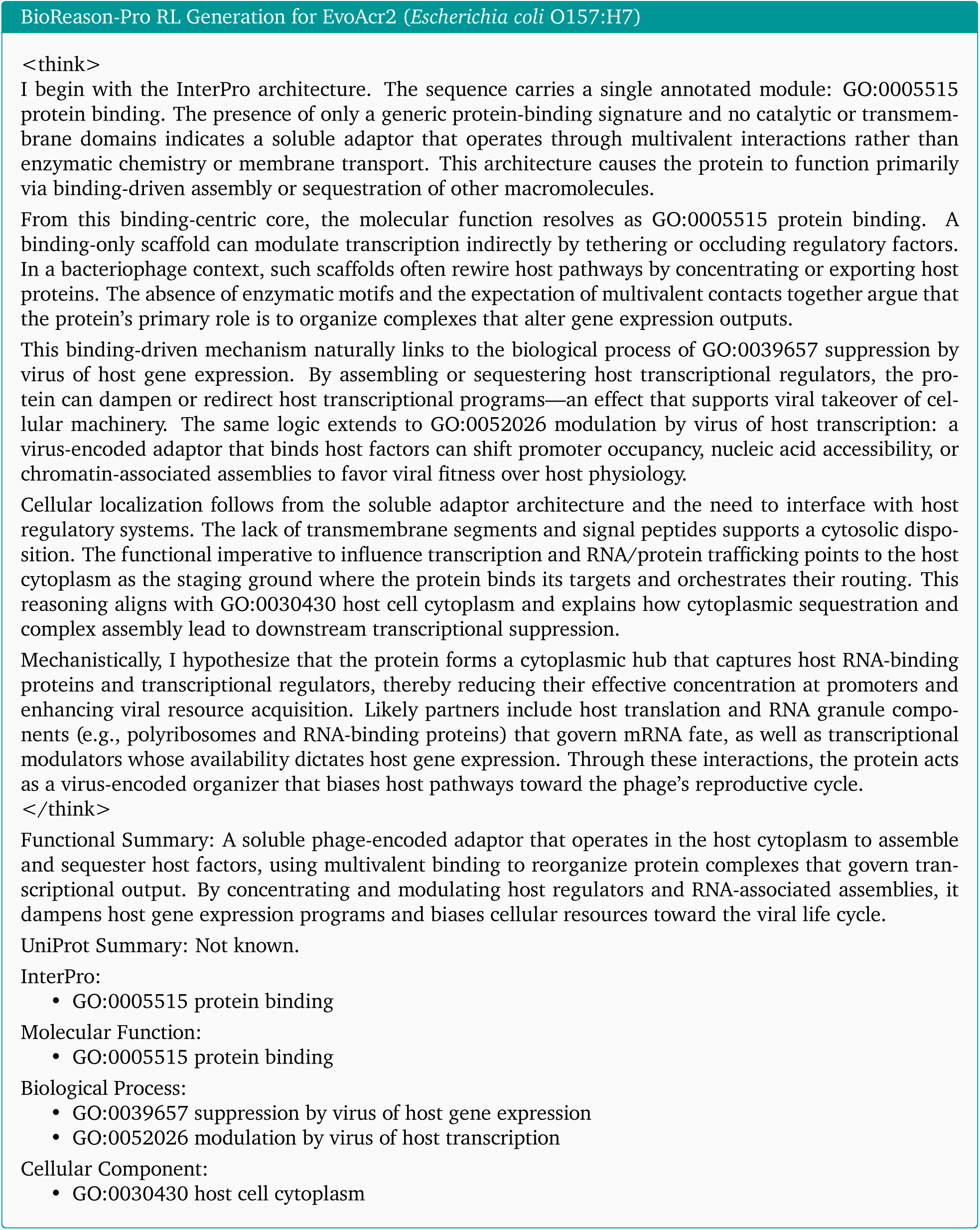
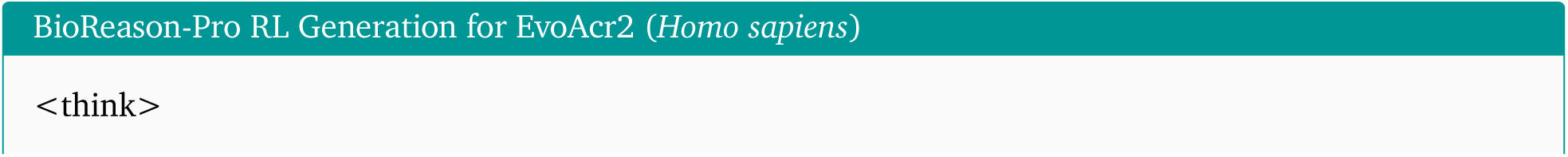
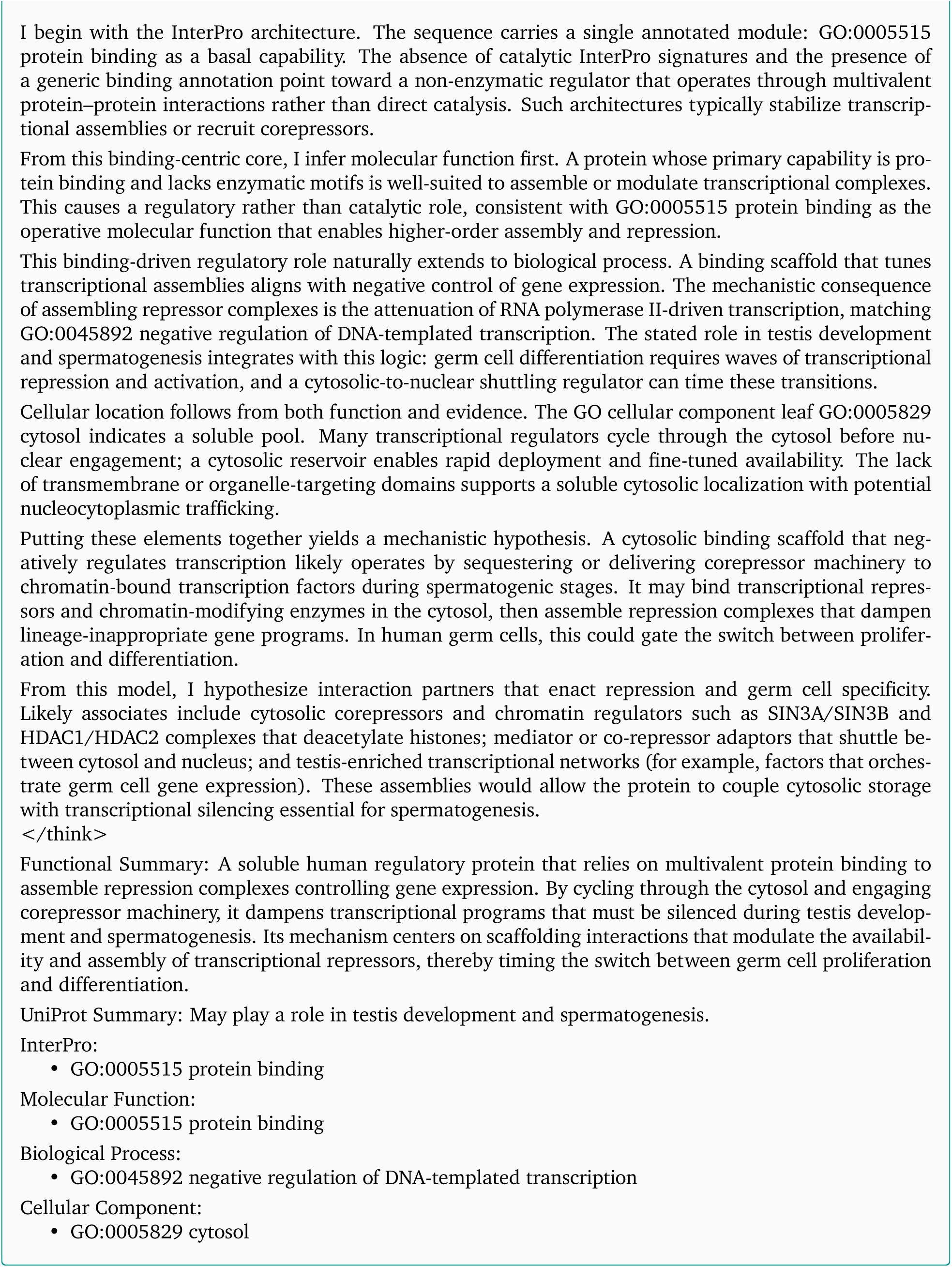

